# Complete genome sequence assembly elucidates evolution and regulation of 5S rDNA loci in the greater duckweed *Spirodela polyrhiza*

**DOI:** 10.1101/2025.01.26.634961

**Authors:** Anton Stepanenko, Veit Schubert, Guimin Chen, Olena Kishchenko, Todd P. Michael, Eric Lam, Maria Hrmova, Ingo Schubert, Nikolai Borisjuk

**Author notes:** Corresponding author: Nikolai Borisjuk.

## Abstract

We resolved the molecular architecture of the 5S rDNA loci in an aquatic monocot *Spirodela polyrhiza*. Measurements of fluorescence *in situ* hybridization signals revealed two loci with 5S rDNA clusters. A combination of the extra-long DNA reads and conventional sequencing of 5S rDNA repeats allowed the assembling of complete loci sequences located on ChrSp6 and ChrSp13. The homologous chromosomes of the ChrSp6 locus contain clusters of 40 and >60 copies of 5S rDNA repeat units with intergenic spacer (IGS) of 400 bp. The ChrSp13 locus is represented by 5S rDNA clusters of rRNA genes spaced by IGSs of 1,056 or 1,069 bp arranged in two sub-clusters, suggesting a different rate of repeat homogenization between loci. The G/C-rich 5S rDNA arrays in both loci are embedded in A/T-enriched chromosome regions with possible regulatory functions. The TATA-like boxes of the 5S rDNA repeat on ChrSp6 and ChrSp13 exhibit different affinities for the TATA-binding protein in 3D modeling of protein/DNA interactions, suggesting a locus-specific regulation of rRNA transcription. Our findings shed light on the molecular architecture of the 5S rDNA loci, which could foster an innovative era for the principles of evolution and regulation of rRNA genes in plants.

---

The 5S nuclear ribosomal DNA (5S rDNA) loci represent a universal eukaryotic multigene family transcribed into 5S ribosomal RNA (5S rRNA) and together with the 35S nuclear rDNA encoding 18S, 5.8S and 25S rRNA, play a key role in ribosome biogenesis and functionality^1^. To ensure a sufficient supply of rRNA, which comprises more than 60% of total RNA in eukaryotic cells^2^, the 5S and 35S rDNA loci usually consist of hundreds to thousands of copies of tandemly arranged gene units. For example, the model plant *Arabidopsis* has nearly 700 copies of 35S rRNA genes^3^ and 800 to 4,800 gene copies encoding 5S rRNAs^4^, while maize has between 5,000 and 14,000 copies of 35S rRNA genes and 2,000-5,000 copies of 5S rRNA genes^5^. The rDNA has been broadly used to study plant biodiversity, systematics, and evolution because of its high copy number and specific structural features containing highly conserved gene coding regions spaced by variable intergenic sequences (IGSs)^6^.

The 5S and 35S rDNA arrays are usually localized on separate loci, and their high copy number allowed visualization of the gene clusters by chromosome *in situ* hybridization in many plant species^7^, indicating that 5S rDNA loci usually reside on one or two chromosome pairs. Due to concerted evolution promoting homogenization of repeated DNA units^8^, the rDNA repeats within the loci are highly homogeneous albeit with some variations. A unit-to-unit variation of the 5S rDNA repeats has been reported for several organisms, including plants^9^. For some plants, it has been shown that such variations could have functional consequences *via* differential 5S rRNA variant expression^10,11^ or general genome regulation^4,12^.

Researchers gained most of the current knowledge on the molecular organization, evolution, and functionality of the rDNA loci through cytogenetics and sequence analyses of random individual gene units. The recent advances in long-read sequencing technologies, especially by the Pacific Biosciences (PacBio) provided highly accurate reads of 10-30 kb from individual DNA molecules, while the Oxford Nanopore Technologies (ONT) enabled ultra-long reads of hundreds of kilobases^13,14^. This has led to successful assemblies of portions of the continuous rDNA arrays in wheat^15^ and *Arabidopsi*s^16^. However, the reading through entire rDNA loci has not yet been achieved because the large repeat arrays cover up to millions of base pairs.

Duckweeds are aquatic monocotyledonous plants that comprise the cosmopolitan *Lemnaceae* family with five genera and 36 species. These plants exhibit a simple morphology, predominantly reproduce through clonal propagation and possess the fastest growth rate among all flowering plants^17,18^. In addition to having a minimal set of ∼19,000 protein-encoding genes, the Greater Duckweed *Spirodela polyrhiza* has the fewest 35S rDNA repeats (∼100) reported so far for a flowering plant^19,20^. The follow-up studies demonstrated a reduced amount of 5S rDNA ranging between 46 and 220 copies per genome in *S. polyrhiza* accessions^21,22^, suggesting that *S. polyrhiza* is a good model to study the whole 5S rDNA array(s) in plants.

Here, we took advantage of the low 5S rDNA gene copy number in the *S. polyrhiza* genome, to resolve its full-length loci at the nucleotide level. We combined high-resolution locus-specific fluorescence *in situ* hybridization (FISH), analyses of the extra-long ONT reads supplemented with gene validation by conventional sequencing of multiple clones containing 5S rDNA units, and 3D modeling of DNA-protein interactions.

## Results

### S. polyrhiza 5S rDNA gene sequences and FISH visualization with locus-specific IGS probes

Following the strategy applied for the characterization of 5S rRNA genes in genomes of the aquatic plants *Pistia strateotis, Landoltia punctata* and ecotypes of *S. polyrhiza*^9,22,23^, we cloned and sequenced multiple PCR-amplified 5S rDNA units and estimated the genes copy number by real-time quantitative PCR (RT-qPCR) of *S. polyrhiza* strain Sp9509^20^. PCR amplification using 5S rRNA gene-specific primers for *S. polyrhiza* produced expected fragments of about 1.1 and 0.5 kb (Fig. 1a). The fragments, separately isolated from the gel were cloned into a plasmid vector, and twenty individual clones, each containing either the 0.5 kb (Sp5S-S) or 1.1 kb (Sp5S-L) fragments, were sequenced. The analysis of 20 Sp5S-S clone sequences demonstrated that all contained a conserved unit of 119 bp encoding the 5S rRNA with no nucleotide variation. In contrast, a total of 22 SNPs (11 T↔C, 8 A↔G transitions and three nucleotide transversions) were detected within the IGS sequences (Supplementary Fig. 1).

**Fig. 1.**
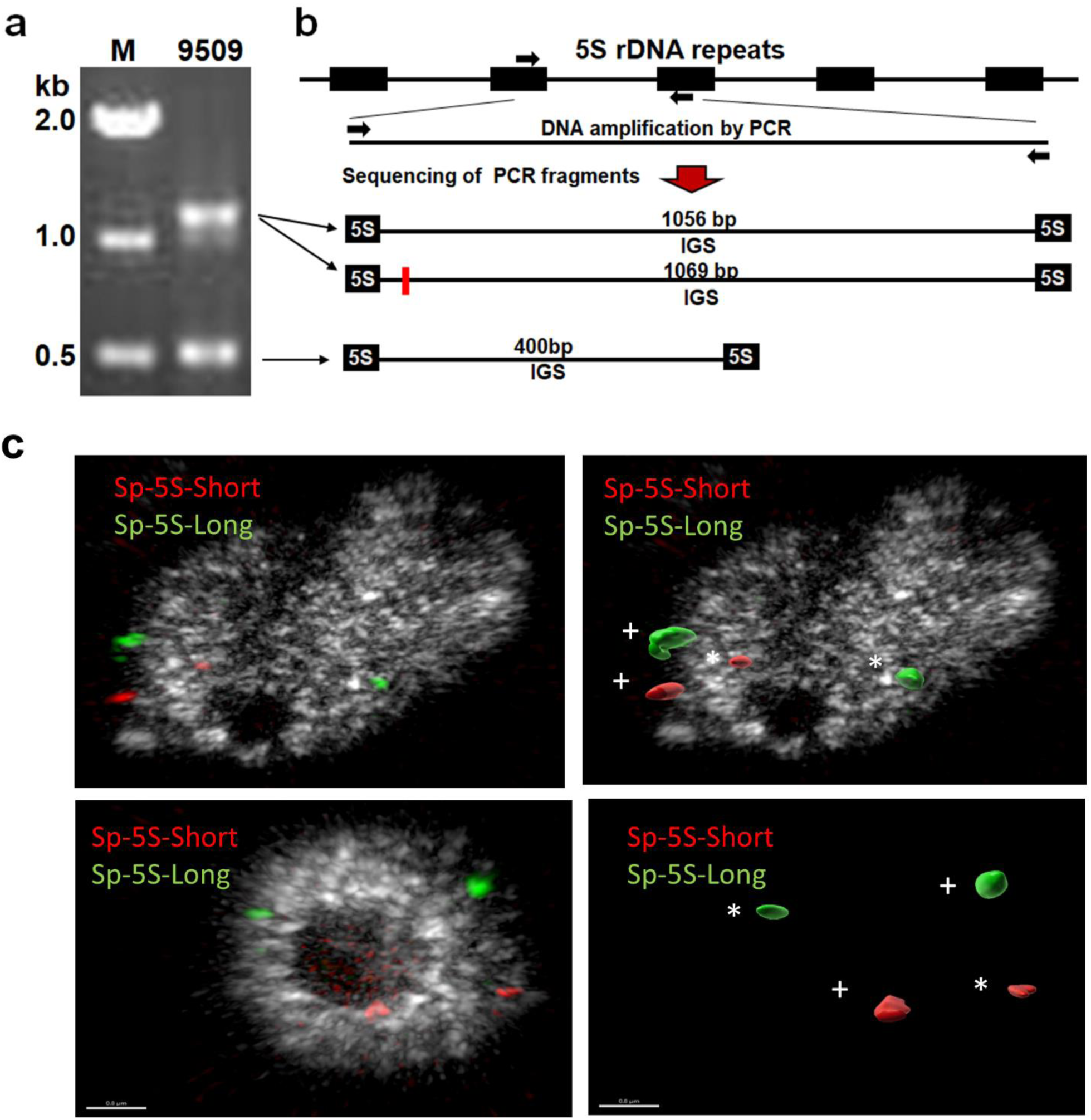
Molecular and chromosomal variation of 5S rDNA in the *S. polyrhiza* accession Sp9509. **a** PCR fragments amplified with primers specific for 5S rRNA genes; M: DNA size marker, 9509: amplification products from genomic DNA of Sp9509. **b** Three variants of 5S rDNA repeat units; black arrows (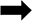) represent 5S rDNA primers, red box (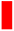) marks the position of a 13 bp insertion, NTS: non-transcribed intergenic spacer. **c** FISH signals (left) and volumes after surface rendering of the signals in 3D structured illumination microscopy (3D-SIM) image stacks (right) indicate that the amounts of 5SrDNA- short and 5SrDNA-long differ between the homologous chromosomes (See also Supplementary Movie 1). * Indicates smaller signals; + indicates larger signals.

Nineteen clones with the larger PCR fragment in addition to the 119 bp encoding 5S rRNA contained 1,056 bp IGSs (Fig. 1b). One clone contained the 5S rDNA unit with an IGS of 1,069 bp, which differed from the remainder by a 13 bp insertion downstream of the conserved translation termination motif TTTT. The IGS regions had nine T↔C, seven A↔G transitions and five nucleotide transversions (Supplementary Fig. 2). The IGS sequences from Sp5S-S and Sp5S-L clones showed almost no sequence similarity to each other, except weak homology within 30 bp stretches proximal to the gene. The RT-qPCR estimation resulted in 147±34 copies of 5S rRNA gene per haploid genome of Sp9509.

We applied FISH and in-depth analysis of long ONT sequence reads containing clusters of 5S rDNA repeats to get insights into the detailed arrangement of loci, with a particular interest in how different types of the 5S rDNA units are arranged within the loci. The visualization of the 5S rDNA loci was performed using IGS-based DNA FISH probes specific for Sp5S-S and Sp5S-L repeat units labeled by red and green fluorescence emitting dyes, respectively. The examination of the FISH signals at the ultrastructural level *via* high-resolution 3D-SIM revealed two pairs of separated red and green spots for the Sp5S-S and Sp5S-L repeats respectively, indicating the localization of both loci on different chromosomal sites (Fig. 1c; Supplementary Movie 1). Measurements of multiple FISH images revealed an average of two-fold and three-fold differences in the Sp5S-S and Sp5S-L repeat signal volume, respectively (Supplementary Tables 1, 2). These results suggest that the allelic loci contain a different number of 5S rDNA copies.

### Sequence assembly of the 5S rDNA loci

The search of the ONT library provided a set of reads containing the clusters of Sp5S-S or Sp5S-L repeat units, with certain reads containing adjacent sequences without homology to rDNA. No sequence was found containing both types of the 5S rDNA units, supporting that these clusters represented two distinct chromosomal loci. Although the sequence quality through the rDNA repeats in the long ONT reads was rather low, the reads showed high variation between the units without a single sequence corresponding fully to the sequences obtained by conventional sequencing of individual PCR clones (Supplementary Figs. 1, 2), using multiple short nucleotide sequences targeting the conservative region of the 5S rRNA gene allowed confident identification of the gene copies and the borders of adjacent chromosomal regions. The selected 22 reads with shorter repeat type ranged between 16 and 58 kb in size and contained 6-65 tandemly arranged 5S rDNA repeats with or without 5’-/3’-end flanking chromosome sequences (Supplementary Table 3). Figure 2a shows a set of representative long ONT reads that allowed mapping of the rDNA loci, with one of the reads, 3ab0, containing 40 copies of 5S rDNA genes bordered with both 5’-end and 3’-end sequences, and one read, fe9e, containing 65 gene copies. This agrees with the FISH data of unequal copy numbers between the homologous chromosomes as revealed by hybridization with the IGS probe specific for 5S-Sp-S (Fig. 1c). We conclude that one of the homologous chromosomes contained 40 copies of 5S-Sp-S and the other had more than 65 copies.

**Fig. 2.**
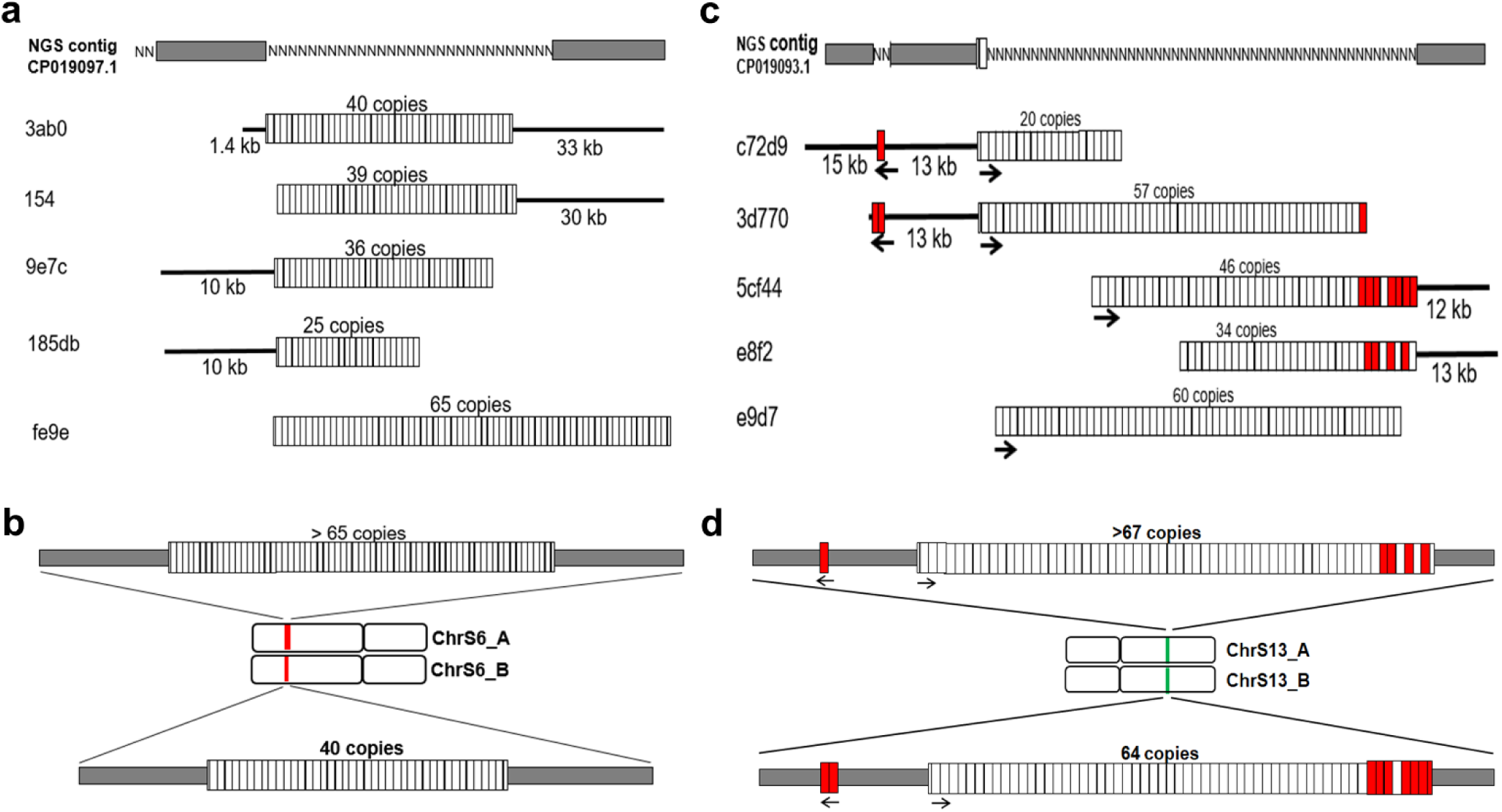
Assembling **the 5S rDNA loci of *S. polyrhiza*, Sp9509. a** Schematic representation of NGS contig and selected representative ONT reads used for assembling sequences composed of short Sp5S-S type 5S rDNA repeats (3ab0, 154, 9e7c, 185db and fe9e represent corresponding reads 3ab0b3f8-9e42-499a- a733-e7f7ef5a88e3; tig00000154; 9e7ce0b7-2939-4dbb-bae9-bd122f577594; 185db047-daf5-4418-bfc1- 3146a719f062 and fe9ebfe0-b4ab-4f96-bb7f-b8aefc7a02d8 listed in Supplementary Table 3). **b** Schematic representation of 5S rDNA containing Sp5S-S type gene repeats at homologous chromosomes ChrSp6_A and ChrSp6_B. **c** Schematic representation of NGS contig and selected representative ONT reads used for assembling sequences composed of long Sp5S-L type 5S rDNA repeats (c72d9, 3d770, 5cf44, e8f2 and e9d7 correspond to the readsc72d9c80-f704-4d3d-b030-25b9f213fe3c; 3d770410-1945-4622-afd8- 7a3257e79119; 5cf44779-0cda-4a54-a602-c253f27d29a6; e8f2492e-3138-4f7f-bf33-7c0c23f3e617 and e9d7a5e2-5a84-4615-a68c-f902b4cce5d4, listed in Supplementary Table 4). **D.** Schematic representation of 5S rDNA containing Sp5S-L type gene repeats at homologous chromosomes ChrSp13_A and ChrSp13_B. Vertical boxes mark 5S rDNA repeated units, clear boxes in Fig. 2C mark Sp5S-L type 5S rDNA repeats with IGS of 1056bp (no-13bp insert) and red boxes mark Sp5S-L type 5S rDNA repeats with IGS of 1069bp (with 13bp insert); black lines represent raw ONT sequences bordering rDNA clusters; grey horizontal boxes mark consensus border sequences surrounding 5S rDNA clusters; arrows mark direction of 5S rDNA transcription.

Sequences bordering the rDNA clusters at the 5’- and 3’-ends aligned well with different long reads (Supplementary Table 3). These sequences also showed high homology to corresponding sequences outside the 5S rDNA cluster of contig 45l assigned to ChrSp6^21^ and to the Next-Generation Sequencing (NGS) contig CP019097.1 of the whole *S. polyrhiza* genome assembly available at NCBI taxid: 29656. The integration of these data allowed the building of a cohesive map of the ChrSp6 rDNA locus with more than 65 copies of Sp5S-S rDNA units located at one homolog (ChrSp6_A) and 40 copies at the homologous chromosome ChrSp6_B (Fig. 2b).

Analysis of 29 selected long reads containing the longer 5S rDNA repeats showed the sequences with clusters of three to 60 gene copies, with many of the sequences having extended non-rDNA regions of locus borders (Supplementary Table 4). In contrast to one type of the border sequences surrounding the 5S rDNA cluster of Sp5S-S on ChrSp6, the borders of the Sp5S-L clusters showed two types of sequence arrangements, suggesting that the homologous chromosomes differ regarding the regions flanking the main 5S rDNA cluster. Among the selected ONT reads we found five sequences containing one additional 5S rRNA gene upstream of the major rDNA cluster, and five sequences with two genes located at the same distance of 13 kb upstream of the major cluster. These small gene clusters are in reverse orientation to the major 5S rDNA array and contain the IGS spacer with a 13 bp insert characteristic for one out of twenty sequenced PCR clones (Fig. 1b, Supplementary Fig. 2); these two types of 5’-border structures are correspondingly represented in Fig. 2c by ONT sequences c72d9 and 3d770. The 3’- borders of the major 5S rDNA array differed by the arrangement of the Sp5S-L units with and without the 13 bp insert in their IGS. In one type of arrangement, found in five ONT sequences (Supplementary Table 4) and represented by the sequence 5cf44 in Fig. 2c, a range of 5S gene units without 13 bp IGS followed by three units with the insert, then one copy with no insert and four terminal copies with the insert. The 3’-end termini in the second arrangement, found in three ONT sequences (Supplementary Table 4) revealed two units with the insert, one unit without an insert and two blocks of insert/no insert, as represented by sequence e8f2 (Fig. 2c). These two types of arrangements of the 3’-end cluster termini were also observed in the ONT assembly utg000019l^21^ and the assembly Gi/13 (https://genomevolution. org/coge/GenomeInfo.pl?gid=51364). Except for these, no other sequence variations were detected within the loci border regions among the ONT reads.

The sequences bordering rDNA clusters in ONT reads aligned well with each other and with the corresponding sequences of the genomic assembly CPO19093.1 (Taxid: 29656) of *S. polyrhiza* chromosome 1, later revised to be chromosome ChrSp13^21^, assembly Gi/13 and the contig utg000019l.

The immediate border sequences of the 5S rDNA clusters of ChrSp6 and ChrSp3 were validated through sequencing ∼1 kb DNA PCR fragments amplified by primers, one of which is homologous to chromosomal sequences closely located, but not overlapping with rDNA (based on the combined sequence data of the long reads and Sp9509 genome assembly) and the other specific for the sequence encoding 5S rRNA. Nucleotide alignments of multiple clones representing borders of the ChrSp6 5S rDNA cluster revealed a single sequence, bordering the cluster at 5’- and 3’- termini, meaning no variations between the ChrSp6_A and ChrSp6_B homologs. The sequences demonstrated that the 5S rDNA array started upstream of the first 5S rRNA gene with 272 bp of the NTS (Supplementary Fig. 3) and ended with a 140 bp sequence of NTS downstream of the last 5S rRNA gene (Supplementary Fig. 4). The 10 kb upstream and downstream consensus sequences gained from ONT and NGS reads, manually validated sequences of the 5S rDNA repeats and immediate cluster borders were integrated into the whole ChrSp6_B locus sequence of 40,879 bp (Supplementary Fig. 5).

Sequence alignments of independent clones representing the 5’-end of the major cluster on ChrSp13 showed that it starts with a 255 bp fragment of perfect homology to the corresponding NTS region upstream of the first 5S rRNA gene (Supplementary Fig. 6). Sequences of the clones representing the locus’ 3’-end, confirmed that this locus is represented by two different 5S rDNA arrays: one terminated with an NTS sequence of 841 bp containing the 13 bp insert and another with the NTS of 828 bp without that insert (Supplementary Fig. 7). The sequence verification of the 5S rRNA gene clusters in reverse orientation agreed with the ONT reads, confirming the two types of arrangements: one with a single intact 5S rDNA copy and the second one with two copies of the gene separated by an NTS of 1,069 bp. Both clusters started with the 969 bp portion of NTS, featuring several single nucleotide polymorphisms (SNPs) and a 23 bp deletion including the whole GAGA stretch. The clusters ended by an NTS sequence of 844 bp containing the GAGA stretch enlarged from 8 to 16 GA-dinucleotides (Supplementary Fig. 8).

Albeit we found no read containing 5’- and 3’- borders of the major 5S rDNA cluster, the number of rDNA units could be defined at least for one of the homologues, based on the composition of the ONT read d3770. Thus, the locus with an upstream cluster of two reversed rDNA units fitted 64 gene copies within the major cluster, counting a stretch of 56 repeat units with no 13 bp insert complemented with the cluster of eight 5S rDNA terminal units, represented correspondingly in the ONT read 5cf44 (Fig. 2c) and utg000019l. Considering the ONT read e9d7, which contained 60 copies of Sp5S-L gene units with no 13 bp insert, the sister locus with one upstream reversed rDNA copy is bigger and contains at least 67 gene copies (60 plus 7 units of 3’-terminal part represented in e8f2) (Fig. 2d).

Integration of data for the available long reads with conventional sequences validating the 5S rDNA units and borders of the two revealed gene clusters, allowed us to build a reliable sequence of the 5S rDNA locus associated with Sp9509 ChrSp13_B. The total locus sequence of 110,911 bp covers two and sixty tandem 5S rDNA gene units in opposite orientations, separated from each other by 12,359 bp, with an additional 10 kb of chromosomal DNA upstream and downstream of the rDNA arrays (Supplementary Fig. 9).

### AT-rich DNA domains homologous to upstream regions of 5S rDNA clusters are widely distributed throughout the S. polyrhiza genome

Determination of the base composition revealed a striking difference between the 5S rDNA clusters and their loci borders, with the G/C content of the rDNA cluster averaging 65.5%, compared to 35% and 36% of the upstream and downstream 10 kb borders of the ChrSp6_B locus, and very similar G/C distribution for the ChrSp13_B locus (Fig. 3).

**Fig. 3.**
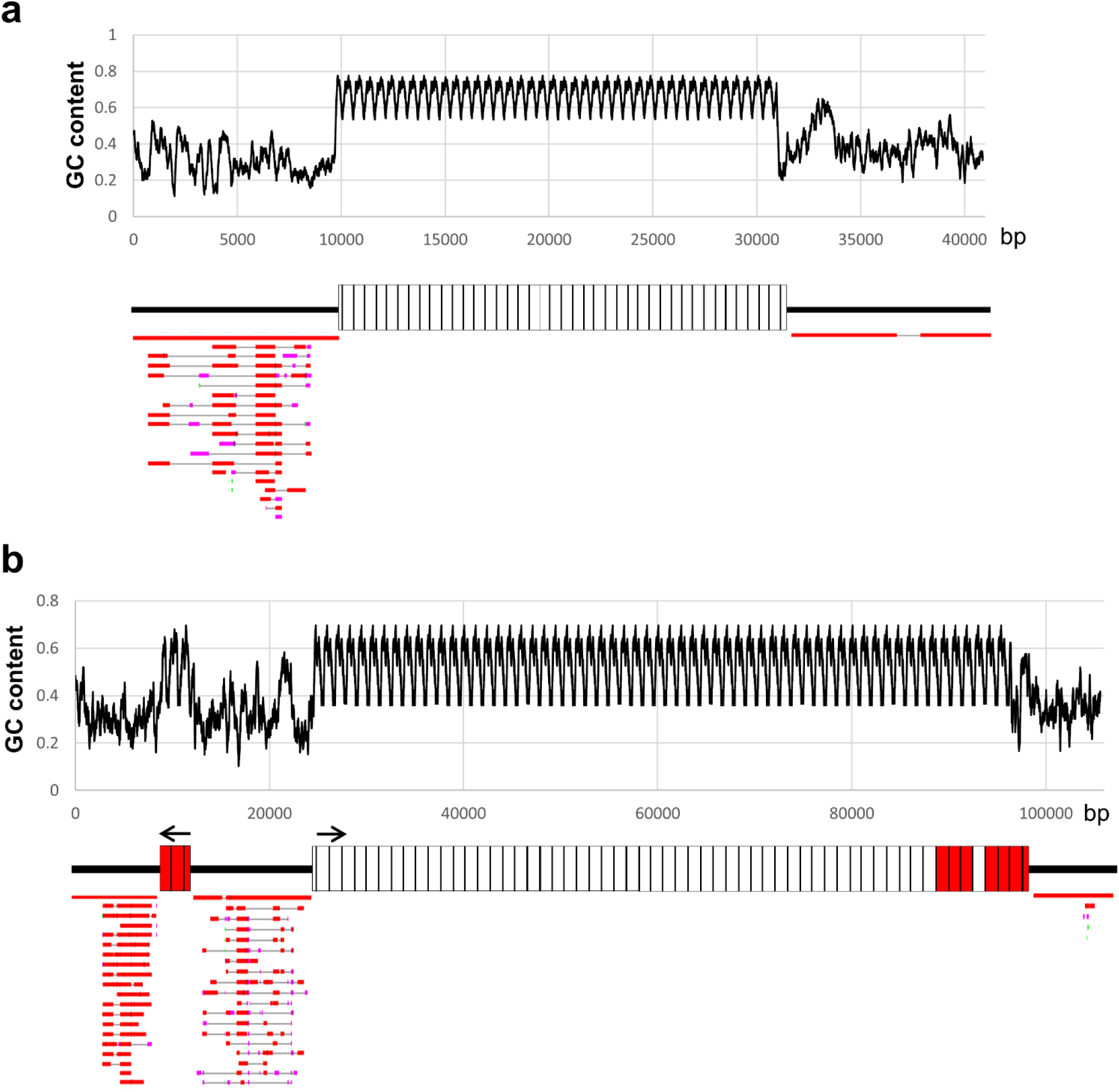
**G/C plot of the 5S rDNA loci and alignment of borders sequences revealing homologous regions on Sp9509 chromosomes**. **a** G/C plot of the 5S rDNA locus on ChrSp6_B (top) and DNA blast hits of the border elements mapped to the set of all 20 *S. polyrhiza* chromosomes (bottom). **b** G/C plot of 5S rDNA loci located at ChrSp13_B (top) and DNA blast hits of the border elements mapped to the set of 20 *S. polyrhiza* chromosomes (bottom).

Running the BLAST search for the DNA regions upstream of the 5S rDNA arrays on ChrSp6 and ChrSp13 revealed multiple homologies with sequences specific for all 20 chromosome assemblies of *S. polyrhiza* represented in NCBI Taxid: 296656. In contrast, the downstream 10 kb regions of ChrSp13 and ChrSp6 5S rDNA loci did not reveal any significant homologies to sequences outside of the loci (Fig. 3). Moreover, compared to the GC-rich rDNA arrays, these AT-enriched border regions, contained multiple 10 bp or longer DNA domains with an A/T content >90% (Supplementary Figs. 5, 9), the motifs typical for replication origins and/or transcriptional enhancers^24^.

### Potential regulatory motifs of the 5S rDNA loci at the level of individual 5S rDNA units

Transcription of 5S rDNA in plants relies on the interaction of RNA polymerase III with three highly conserved DNA motifs within the 5S rDNA sequence^25^, defined as A-Box, IE and C-Box. All these regulatory elements are present in the *S. polyrhiza* 5S rDNA (Fig. 4). In addition, gene activity of the 5S rDNA units is modulated by external *cis*-elements, located upstream of the gene, and the conserved TTTT motif of the transcription terminator downstream of the 5S rRNA gene sequence. The DNA regions adjacent to 5’- and 3’-ends of the 5S rRNA gene on ChrSp6 and ChrSp13, show sequence differences that might influence locus-specific gene activity. First, the short rDNA repeat units on ChrSp6 contain a C at position - 1, the most common nucleotide at this position for higher plants^25,26^, while the long repeat units on ChrSp13 have at that position a T base. Both short and long *S. polyrhiza* 5S rDNA IGS variants have a duplicated GC between nucleotides -14 and -11, instead of a single GC at position -13 -12, common for the majority of plant species. As in many other plant species, a TATA-like motif is located at nucleotides -28 -23, differing in two out of six nucleotides between the Sp5S-S and Sp5S-L (Fig. 4a). Considering that this motif is crucial for the modulation of plant 5S rRNA gene expression through interactions with the TATA-Binding Protein (TBP), a DNA-binding factor common for all three major RNA polymerase complexes, Pol I, Pol II and Pol III^27,28^, such a difference in a key motif signature suggests potential differential regulation of the ChrSp6 and ChrSp13 loci. There is also a noticeable variation in the region downstream of the conserved TTTT motif of the transcription terminator (Fig. 4a).

**Fig. 4.**
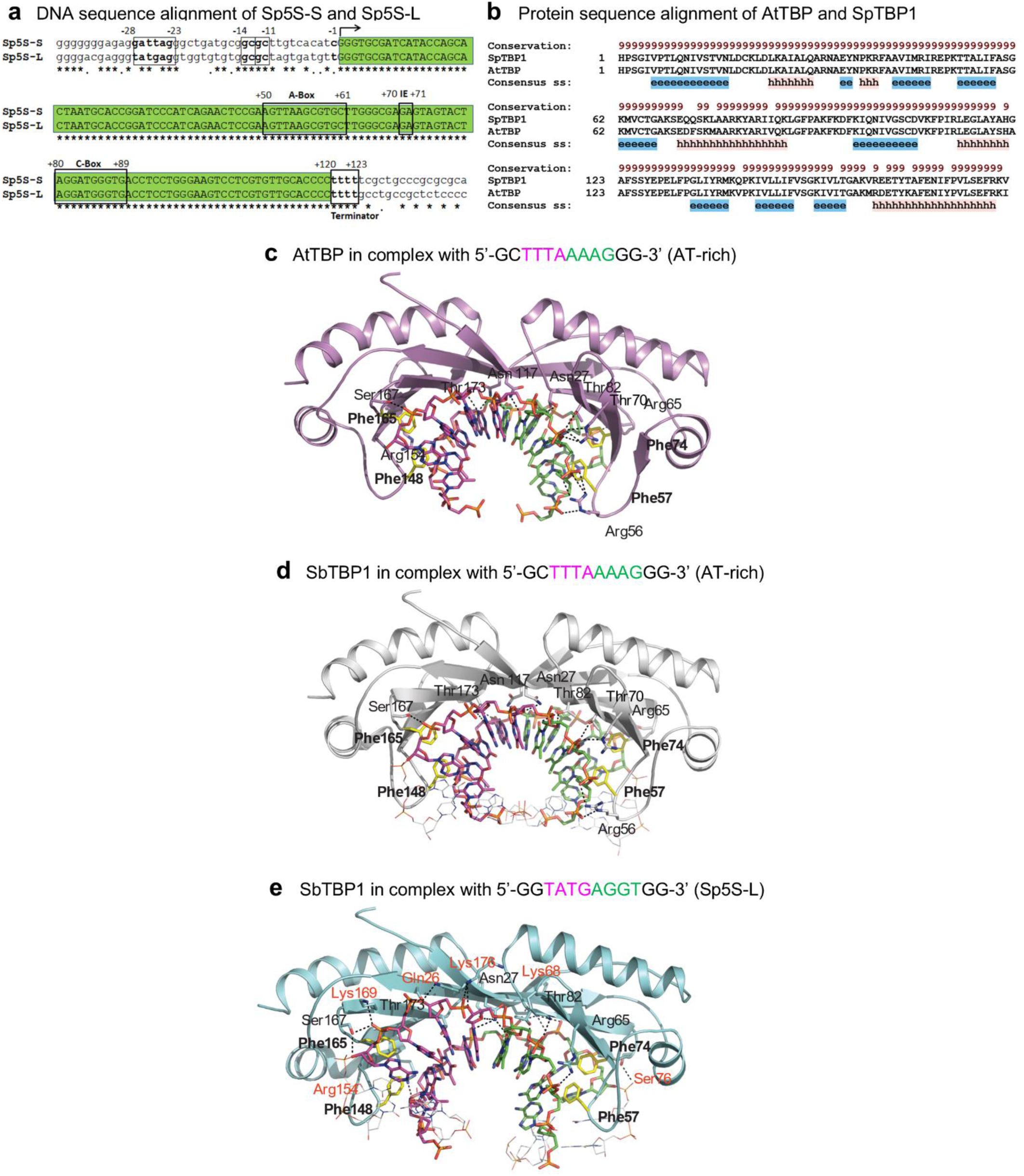
3D modeling of *A. thaliana* and *S. polyrhiza* TBPs interactions with TATA-like DNA motifs. **a** Alignment of DNA motifs upstream of the 5S rRNA gene units in Sp9509. Sequences encoding 5S rRNA are highlighted in green; DNA motifs important for gene regulation are boxed; asterisks mark identical base pairs. **b** Sequence alignment of *S. polyrhiza* TBP1 (SpTBP1) and *A. thaliana* TBP (AtTBP) by ProMals3D^32^. α-Helical and β-sheet secondary structural elements (ss) are indicated with ‘h’ (pink) and ‘e’ (blue) below the sequences. The absolute degree of conservation is marked by ‘9’ (brown) above the sequences; **c** AtTBP in complex with the canonical TATA-like motif 5’-GC<u>TTTAAAAG</u>GG-3’; **d** SpTBP1 in complex with TATA-like 5’-GC<u>TTTAAAAG</u>GG-3’; **e**SpTBP1 in complex with Sp5S-L 5’-GG<u>TATGAGGT</u>GG-3’ motifs. Ribbon representations illustrate pseudo-dimeric folds (AtTBP in magenta; SpBP1 in grey and cyan), and residues interacting with DNA in sticks colored in magenta-cpk for C-atoms (coding strands of TTTA and TATG motifs) and green-cpk for C-atoms (coding strands of AAAG and AGGT motifs). Interactions between residues and DNA are shown in black dashed lines with separations of 2.7 Å-3.5 Å. Additional SpTBP1 residues recognizing Sp5S-L compared to canonical TATA-like motifs are highlighted in red. Positionally identical residues involved in DNA binding by AtBP and SpTBP1 are: Asn27, Arg56, Arg65, Thr70 and Thr82 from the 1^st^ pseudo-monomer, and Asn117, Arg154, Ser167 and Thr173 from the 2^nd^ pseudo-monomer. Notably, the dispositions of Phe57, Phe74, Phe148 and Phe165 side-chains are alike in both complexes; these interstitial residues contribute to the kinking of DNA around regions surrounding T and A bases^33^. Conversely, the Sp5S-L (5’-GGTATGAGGTGG-3’) motif is bound differently by SpTBP1, involving Asn27, Arg65 and Thr82 from the 1^st^ pseudo-monomer, and Ser167 and Thr173 from the 2^nd^ pseudo-monomer. These new interacting residues form a second ‘assembly row’ of binders and include Gln26, Lys68 and Ser76 from the 1^st^ pseudo-monomer, and Arg154, Lys169 and Lys176 from the 2^nd^ pseudo-monomer. Notably, except Gln26 and Ser76, these new interactors contain a positively charged and flexible Lys residue.

To estimate the functionality of the two TATA-like motif variants, we modeled the interactions of these DNA motifs with the cognate TBP sequence in the Greater Duckweed genome. Fifty models of *S. polyrhiza* TBP1 in complex with the Sp5S-L motifs were constructed, and the model with the lowest evaluation scores of the Modeler Objective Function^29^, the Discrete Optimized Protein Energy^30^ parameter, and the Prosa2003 energy function^31^ was selected (Fig. 4).

According to the available whole genome sequence (Taxid: 29656), *S. polyrhiza* contains two genes encoding TBP, which we confirmed by cloning and sequencing the corresponding PCR fragments. The deduced proteins, designated as *S. polyrhiza* TBP1 and *S. polyrhiza* TBP2 (Supplementary Fig. 10), shared high sequence homology, and the SpTBP1 protein was selected for further computational 3D molecular modelling. The protein sequence alignment of the *Arabidopsis* TBP with SpTBP1 indicated a high similarity (91%, Fig. 4b). Indeed, both proteins folded into pseudo-dimer structures with nearly identical secondary structural elements (α-helices and β-sheet strands) and formed similar binding complexes with the AT-rich motif 5’-GCTTTAAAAGGG-3’, where the concave sides of AtTBP and SpTBP proteins fitted well inside minor grooves of DNA with the bent sugar/phosphate backbones (Figs. 4c-e). Testing the SpTBP1 interactions with the TATA-like boxes of the duckweed long (Sp5S-L: 5’-GG**TATGAGG**<u>T</u>GG) and short (Sp5S-S: 5’-AG**GATTAGGG**CT) 5S gene variants, suggested a protein binding site for Sp5S-L, whereas the three guanines at the 3’-end of Sp5S-S made this motif too rigid for bending and less favorable to bind SpTBP1.

## Discussion

Although genome information gained from Next Generation Sequencing is abundant for many plant species, almost in all cases the rDNA data, representing a significant portion of the genomes, are missing due to the difficulty of assembling long regions of tandemly repeated DNA units. Even with the advanced extra-long, single-molecule sequencing techniques^14^, deciphering sequences of rDNA loci of ten to hundred million nucleotides remains challenging, and the locus characterization in terms of differences between haplotypes is out of reach. The Greater Duckweed, *Spirodela polyrhiza*, with its highly reduced number of both 35S and 5S rDNA repeats^9,20,21^ offered a unique opportunity to gain insights into the molecular structure of the rDNA loci.

Our efforts demonstrated importance of the complex approach using high-resolution FISH, long-read ONT and conventional sequencing to resolve the sequence complexity and architecture of the 5S rDNA loci in Sp9509^20^. In accordance with FISH results, the analysis of the long ONT reads revealed two variants of 5S rDNA arrangements on both ChrSp6 and ChrSp13 chromosomal loci. It showed that one rDNA cluster of ChrSp6 is composed of 40 copies of 5S-Sp-S repeats, while the second contains contained more than 65 repeats (Fig. 2), confirming the copy number variation between the two homologues of both loci. We can estimated that the ChrSp6 locus contained about 120 copies of 5S rDNA units per diploid genome.We can only speculate that based on the available FISH signal measurements (Supplementary Table 2) the number could be two to three times higher. Therefore, the copy number of 5S-Sp-L repeats per the ChrSp13 locus is between 200 and 270. Summing up the 5S rDNA repeats on the ChrSp6 and ChrSp13 loci resulted in a total gene count of 320 to 390 copies per *S. polyriyza* diploid genome, which is within the range of our RT-qPCR estimate of 147±34 copies per haploid genome. For loci sequence accuracy, it was crucial to complement the long ONT read with conventional sequencing of multiple PCR fragments, because of the rather poor quality of the ONT sequences through the G/C rich 5S rDNA repeats.

Apart from providing the first evidence for a higher plant of rDNA array variations between homologous loci, our data demonstrate different evolutionary dynamics for the 5S rDNA loci located at chromosomes ChrSp6 and ChrSp13, with faster structural changes occurring in the locus composed of longer, 5S-Sp-L type repeats on ChrSp13. This conclusion agrees well with the results of comparative sequencing of the short and long variants of 5S rDNAin six geographic ecotypes of *S. polyrhiza* from Europe and Asia^22^. It is also in line with the sequence study of two separate 5S rDNA loci on bread wheat chromosome 5BS showing that one of them is composed of diverged units with multiple insertions of transposable elements, while another contained highly homogenized tandem gene array^15^.

Deciphering the whole structure and sequencing its components also provided some insights into loci functioning. Analysis of the 5’- and 3’- ends of the 5S rDNA arrays in both loci, showed that they start and terminate by IGS sequences that contain the upstream TATA-like box and the downstream transcription termination signal correspondingly (Supplementary Fig. 5, 9), enabling the functionality of the adjacent gene units. Although the 5S rRNA gene sequences and therefore gene products do not differ between the loci, the 5S-Sp-S and 5S-Sp-L type IGS sequences show nucleotide variations within the genes’ upstream region containing regulatory motifs such as TATA-like boxe (Fig. 4). The assumption that those sequence differences might contribute to differential regulation of the two loci was tested by 3D modeling of the interactions of locus-specific TATA-like motifs with the TATA-binding protein, postulated to be an integral part of the 5S rDNA transcriptional complex^27,28^. The 3D protein modeling supported *S. polyrhiza* TBP binding with the TATA-like motif of the rDNA units located at the ChS13 locus and predicted that binding with the corresponding motif of rDNA units on the ChSp6 locus was less favorable because of the higher rigidity of the TATA-like box of the 5S-Sp-S type repeats. This finding, together with our observation of a clear tendency of 5S-Sp-S type being preferentially associated with genotypes from tropical areas and the 5S-Sp-L type predominating in populations adapted to cooler climatic zones^22^, suggests a well-orchestrated mechanism for fine-tuning of Pol III-mediated 5S rDNA expression in response to environmental conditions.

Additional potential for differential regulation of the loci might be contributed by the stretches of GAGA-motifs for GAGA-binding proteins (GBPs), an evolutionary conserved system involved in the regulation of multiple genome functions^34^. The stretch of eight GA dinucleotides is a characteristic feature of the long but not the short IGS (Supplementary Figs. 1, 2), with the long GA_n_ stretches also present in the border sequences (Supplementary Fig. 5, 9).

Another factor to be considered with respect to locus regulation are the features of the sequences upstream of the 5S rDNA clusters on both ChrSp6 and ChrSp13. These sequences contain numerous A/T stretches (Supplementary Figs. 5, 9) and show homology to similar DNA elements distributed along all 20 chromosomes of *S. polyrhiza* (Fig. 3). Similar A/T stretches, resembling the motifs of yeast autonomously replicating sequences (ARS), were also found in the IGS of tobacco 35S rDNA and shown to regulate gene copy number and transcription^24^.

Locus-specific silencing of rRNA genes limiting the pool of transcribed rDNA units to the actual requirement is a well known mechanism described for many plant species with redundant rDNA copies^4,10,11,35,36^ . Considering that *S. polyrhiza* has an unusually low number of 5S rDNA copies, possibly due to general genome shrinkage^20^, we hypothesize that ChrSp6 and ChrSp13 loci are active probably depending on environmental conditions. Arguments supporting this assumption are: i) both loci harbor intact, highly homogenized gene units with identical coding sequences and IGS regions with potential motifs for differential transcription regulation, ii) the proximal chromosome sequences upstream of the Sp-5S-S and Sp-5S-L clusters have a similar A/T rich architecture, characteristic for open chromatin regions involved in transcription/replication, iii) there is a tendency of selection for genomes with Sp-5S-S types in the warm regions (around the equator), and for Sp-5S-L types in the temperate areas.

In conclusion, the reported data on 5S rRNA genes in *S. polyrhiza* present the first complete structure of plant 5S rDNA architecture resolved at the nucleotide level. The two loci localized to separate chromosomes contain head-to-tail repeat units with identical sequences of the 5S rRNA gene intertwined with locus-specific IGSs that differed in length and nucleotide sequence. Our study also demonstrated haplotype specificity of 5S rDNA arrangements manifested in copy number variation between homologous chromosomes for both loci and sequence divergence for the locus with longer type rDNA repeats. Based on the level of rDNA repeat homogenization within each locus, the study demonstrated different evolutionary dynamics of the 5S rDNA between the two loci, confirming earlier findings in other plant species^15,37^. We also show that the units with variant IGSs are clustered within a locus in the manner reminiscing clustering of 35S rDNA units in Arabidopsis^38^ and suggested a role of the proximal chromosomal domain in locus regulation. Collectively, the obtained data offer new insights into the structural organization and regulation of 5S rDNA loci at the chromosomal and individual gene levels.

## Experimental Procedures

### Plant material and DNA preparation

The total DNA of the greater duckweed, *Spirodela polyrhiza* was isolated from strain 9509 (Sp9509), aseptically grown and maintained at the Rutgers Duckweed Stock Cooperative (http://www.ruduckweed.org). For generating extra-long Oxford Nanopore Technology (ONT) reads, the DNA was isolated from 5 g of flash-frozen fronds as described by Hoang et al.^21^. For PCR amplification, the genomic DNA of *S. polyrhiza* was prepared using a modified cetyltrimethyl ammonium bromide (CTAB) protocol adapted for midi-scale^20^.

### Fluorescence in situ hybridization (FISH)

Nuclei for FISH were prepared according to Ahmadli et al.^39^. The appropriate amounts of fresh *S. polyrhiza* fronds were fixed for 25 min under vacuum in a desiccator in 4% v/v ice-cold formaldehyde in Tris buffer (10 mM Tris–HCl pH 7.5, 10 mM Na2-EDTA, 100 mM NaCl, 0.1% v/v Triton X-100, pH 7.5) before twice washing in Tris buffer. The fixed fronds were chopped with a sharp razor blade in ice-cold nuclei isolation buffer (15 mM Tris–HCl, 2 mM Na2EDTA, 0.5 mM Spermin, 80 mM KCl, 20 mM NaCl, 15 mM Mercaptoethanol, 0.1% v/v Triton X100, pH 7.5). The nuclei, filtered and stained with 1 μg/ml 40,6- diamidino-2-phenylindole (DAPI), were sorted using the BD Influx Cell Sorter (BD Biosciences, Heidelberg, Germany). Sorted nuclei in sucrose buffer (100 mM Tris–HCl, 50 mM KCl, 2 mM MgCl2, 0.05% v/v Tween 20, and 5% w/v sucrose, pH 7.9) were pipetted onto microscope slides and dried overnight.

The DNA probes specific for Sp5S-S and Sp5S-L repeats were prepared following the procedure described by Hoang and Schubert^40^. First, the genomic DNA isolated from *S. polyrhiza*, Sp9509 was used as the template for PCR amplification of the IGS regions with pairs of primers specific to the two types of 5S rDNA units (Supplementary Table 5). After purification, the obtained PCR products were used as templates for labeling with specific fluorescent dyes. The IGS fragment specific for Sp5S-S was subjected to nick-translation using the AF594 NT Labeling Kit and the fragment specific for Sp5S-L was labelled with the Atto488 NT Labeling Kit (Jena Bioscience, Jena, Germany). After precipitation with 96 % v/v ethanol, probe pellets were dissolved in 100 μl hybridization buffer (50% v/v formamide, 20% w/v dextran sulfate in 2 × SSC, pH 7).

Ten μL of each probe was pre-denatured at 95°C for five minutes and cooled on ice for 10 minutes before being added to the slide. Sorted nuclei on slides were denatured on a hot plate at 80 °C for three min after adding 10 μl pre-denatured probe and then incubated in a moist chamber at 37 °C for at least 16 h. Post-hybridization washing and signal detection were carried out according to Lysak et al.^41^

### Super-resolution microscopy, FISH signal volume and intensity measurements

To detect the ultrastructural chromatin organization of nuclei at a resolution of ∼120 nm (super-resolution achieved with a 488 nm laser excitation), spatial structured illumination microscopy (3D-SIM) was performed with a 63×/1.4 Oil Plan-Apochromat objective of an Elyra 7 microscope system and the software ZENBlack (Carl Zeiss GmbH). Images were captured separately for each fluorochrome using the 561, 488, and 405 nm laser lines for excitation and appropriate emission filters^42^. 3D rendering to measure the FISH signal volumes and a spatial animation (Supplementary Movie 1) was done based on the SIM image stacks using the Imaris 9.7 (Bitplane) tool ‘Surface’. The signal intensity values were calculated within these volumes.

### Selection and analysis of ONT reads containing clusters of 5S rDNA repeats

The long reads with 5S rDNA repeats were selected by searching the ON library Sp9509_Oxford_v3 deposited and accessible in the European Nucleotide Archive (PRJEB27612) with six nucleotide sequences (5’-CTTGGGCGAGAGTAGTACTAGG, 5’-CACTAATGCACCGGATCCC, 5’-GGGTGCGATCATACCAGCAC, 5’-CACGCTTAACTTCGGAGTTCTG, 5’-GCAACACGAGGACTTCCCA, and 5’-GGGTGCAA-CACGAGGACTTC) complemental to the sequence of *S. polyrhiza* 5S rRNA gene^22^. The search produced numerous sequences with various copies of targeted 5S rRNA genes separated either by a short (∼400 bp) or longer (∼1000 bp) spacer (Supplementary Tables 3, 4). From that set of ONT reads, four sequences of each class were prioritized for further analysis, taking into account a maximal coverage of 5S rDNA units and the availability of the corresponding 5’ and 3’ border sequences. As a reference genome for border sequences surrounding the 5S rDNA clusters, in addition to Sp9509_Oxford_v3 (available on GoGe with the genome ID of 51364 (https://genomevolution. org/coge/GenomeInfo.pl?gid=51364), we used the genome assembly ASM198140v1 (https://www.ncbi.nlm.nih.gov/datasets/genome/GCA_001981405.1/).

### Validation of 5S rDNA loci sequences

Validation of *S. polyrhiza* 5S rDNA repeats and the clusters’ border sequences was performed by conventional sequencing of multiple DNA clones containing specific DNA fragments amplified by a previously described advanced PCR procedure^22,23^. The pairs of primers used to amplify rDNA units composed of 5S rRNA gene and NTS sequences, as well as the 5’ and 3’ border sequences of three 5S rDNA clusters are listed in Supplementary Table 5. After purification using an AxyPrepTM DNA Gel Extraction Kit (Axygen, United States), the generated fragments were cloned into the vector pMD19 (Takara, Dalian, China) and sequenced using the custom service provided by Sangon Biotech (Shanghai, China). The forward and reverse sequences obtained for each clone were assembled and analyzed using the CLC Main Workbench (Version 6.9.2, Qiagen) software.

### Assembly and characterization of 5S rDNA loci consensus sequences

The verified sequences of 5S rDNA units and the cluster borders were incorporated into the backbone ChrSp13 and ChrSp6 loci consensus ONT assemblies (Supplementary Figs. 5, 9), replacing corresponding imperfect original ONT sequence regions. In particular, the consensus sequences of DNA regions bordering 5S rDNA clusters were built using the available ONT reads and NGS assemblies (NCBI taxid: 29656) in CLC Main Workbench (Version 6.9.2, Qiagen). The consensus sequences of rDNA loci composed of two and 64 copies of 5S rDNA on ChrSp13, and 40 copies on ChrSp6, were assembled by head-to-tail stacking sequences of the corresponding manually sequenced 5S rDNA units following the general arrangement of the rDNA clusters in corresponding ONT reads. The G/C-plots of the loci sequences were built using CLC Main Workbench (Version 6.9.2, Qiagen), and G/C-content was estimated using Genomics G/C Content Calculator (https://www.sciencebuddies.org/science-fair-projects/references/genomics-g-c-content-calculator). Checking for possible regions of homology to the sequences of 5S rDNA loci within the *S. polyrhiza* genome was carried out by blasting the rDNA flanking regions against the whole genome sequence (NCBI taxid: 29656) using the megablast option for highly similar sequences (https://blast.ncbi.nlm.nih.gov/).

### Estimation of the 5S copy number by qPCR

The 5S rRNA gene copies were estimated by quantitative real-time PCR, relating the rates of sample DNA amplification to the standard curve based on the amplification reads of the dilution series of a reference plasmid containing sequences of 5S rDNA and a fragment of a single-copy actin gene^22^. The rDNA copy number was determined in qPCR reactions prepared with the UltraSybr Mixture (CWBio, Taizhou, China), run on the CFX Connect Real-Time detection system (Bio-Rad, Hercules, USA). Three biological replicates were included in the assays. For quantification of the 5S rDNA, we used the primers specific to the 5S rDNA sequence of *S. polyrhiza* (Supplementary Table 5). The samples and 10-fold dilution series of the reference plasmid were assayed in the same run. The quality of products was checked by the thermal denaturation cycle. Only the experiments providing a single peak were considered. Three technical replicates were performed for each biological replicate. The obtained data were analyzed using BIO-RAD CFX Manager 3.1 (Hercules, USA) and Microsoft Excel 2016 software.

### Cloning S. polyrhiza genes encoding TATA-binding proteins TBP1 and TBP2

DNA sequences of the *S. polyrhiza TBP1* and *TBP2* genes were obtained by cloning the PCR-amplified fragments using cDNA prepared from *S. polyrhiza* mRNA as a template. The PCR fragments were amplified with gene-specific primers (Supplementary Table 5), designed according to the *in silico* sequence information available at NCBI (taxid: 29656, GCA_900492545.1) for *S. polyrhiza*, ecotype 9509. The generated DNA fragments, cloned into the vector pMD19 (Takara, Dalian, China) following the manufacturer’s instructions, were custom sequenced (Sangon Biotech, Shanghai, China), and the obtained nucleotide sequences were analyzed using the CLC Main Workbench (Version 6.9.2, Qiagen) software.

### Construction of 3D models of the S. polyrhiza TBP1 pseudo-dimer in complex with TATA-like motifs of 5S rDNA repeat

The suitable structural template to construct the 3D protein model of *S. polyrhiza* TBP1 was identified using threading alignments in the Protein Data Bank (PDB). The crystal structure of the *Arabidopsis thaliana* TATA-box binding protein (TBP) (PDB accession 1QNA, chain A) in complex with the TATA-like motif 5’-GC<u>TTTAAAAG</u>GG-3’/ 5’-CC<u>CTTTTAAA</u>GC-3’^33^ was identified as a suitable template for structural modeling. The sequences of *S. polyrhiza* TBP1 and *Arabidopsis thaliana* TBP were aligned using MUSCLE^43^ and ProMals3D^32^, and the alignment qualities were checked by PSIPRED^44^ to confirm that secondary structural elements remained undisturbed. The sequence similarity was 92% and the sequence identity was 96% between the two sequences without gaps, as calculated by the pairwise EMBOSS Needle global algorithm^45^. The TATA-like AT-rich motif 5’-GC<u>TTTAAAAG</u>GG-3’/5’-CC<u>CTTTTAAA</u>GC-3’ of 1QNA, chains C and D, was transformed into the Sp5S-L motif 5’-GG<u>TATGAGGT</u>GG-3’/5’-CC<u>ACCTCAT</u>ACC-3’, while retaining the sharply bent sugar-phosphate backbone, using Coot^46^. The TATA-like AT-rich and the Sp5S-L motifs alongside the coordinates of *A. thaliana* TBP were used as the templates and inputted during 3D protein modeling of *S. polyrhiza* TBP1, using MODELLER v9.16^47^. Ramachandran plots of the model indicated that 100% of the residues were in the most favored and additionally allowed regions, when excluding glycine and proline residues. No residues were found in generously allowed or disallowed regions. The overall G-factor values evaluated by PROCHECK^48^ were 0.33, 0.05, -0.02 and -0.14 for *A. thaliana* TBP and *S. polyrhiza* TBP1, each in complex with TATA-like AT-rich and Sp5S-L motifs, respectively. The *Z*-score values, deduced from Prosa2003 reflecting combined statistical potential energy were -9.55, -9.82, -9.86 and -9.79 for *A. thaliana* TBP and *S. polyrhiza* TBP1, each in complex with TATA-like AT-rich and Sp5S-L motifs, respectively. The root-mean-square-deviation values of *A. thaliana* TBP and *S. polyrhiza* TBP1 with TATA-like AT-rich and Sp5S-L motifs determined with the PyMol ‘superposition’ algorithm were 0.117 Å and 0.556 Å for 182 residues, indicating their similar structural folds. Images of structures were generated in the PyMol Molecular Graphics System, Version 2.0.6 Schrödinger, LLC.

## Data availability

All raw ONT reads analysed in this study are listed in Supplementary Table 3. The sequences of all cloned PCR fragments have been submitted to the GenBank (accession numbers PQ963796 – PQ963840) and are available in Supplenentary Figures 1, 2, 3, 4, 6, 7, 8, 10. The assembled sequence of the 5S rDNA locus on ChrSp6 is represented in Supplenentary Figure 5, and the assembled sequence of the 5S rDNA locus on ChrSp13 is represented in Supplenentary Figure 9.

## Acknowledgements

We thank Dr. Andreas Houben (IPK, Gatersleben) for hosting N.B. in his lab during the summers of 2023, and 2024 and discussions. We thank Dr. Jörg Fuchs for his expert help with the preparation of *S. polyrhiza* nuclei for FISH analyses. We thank the German National Academy of Sciences Leopoldina for supporting A.S. with the Leopoldina Ukraine Distinguished Fellowship.

## Author contributions

A.S., V.S., G.C. and O.K. performed the experimental studies. A.S., I.S. and N.B. conceived experiments. A.S., V.S., T.P.M., E.L., M.H., I.S. and N.B. analyzed data. N.B. wrote the manuscript with substantial contributions from A.S., T.P.M., E.L., M.H. and I.S. I.S. and N.B. supervised the work.

## Competing interests

The authors declare no competing interest.

## Additional information

Supplementary information for this paper is available at:

## Notes

### Competing Interest Statement

The authors have declared no competing interest.

