## Supplementary Figures 1-10 and Supplementary Tables 1-5 for "Complete genome sequence assembly elucidates evolution and regulation of 5S rDNA loci in the greater duckweed *Spirodela polyrhiza*"

**Supplementary Figure 1.** **Nucleotide alignment of twenty sequences (S-1 to S-20) of cloned PCR fragments representing *S.polyrhiza* 5S rDNA repeats of shorter type (5S-Sp-S)**. Sequences encoding 5S rRNA are highlighted by green; SNPs in the non-transcribed spacer are marked by magenta.

```
S-1        CTTGGGCGAGAGTAGTACTAGGATGGGTGACCTCCTGGGAAGTCCTCGTGTTGCACCCCT
S-2        CTTGGGCGAGAGTAGTACTAGGATGGGTGACCTCCTGGGAAGTCCTCGTGTTGCACCCCT
S-3        CTTGGGCGAGAGTAGTACTAGGATGGGTGACCTCCTGGGAAGTCCTCGTGTTGCACCCCT
S-4        CTTGGGCGAGAGTAGTACTAGGATGGGTGACCTCCTGGGAAGTCCTCGTGTTGCACCCCT
S-5        CTTGGGCGAGAGTAGTACTAGGATGGGTGACCTCCTGGGAAGTCCTCGTGTTGCACCCCT
S-6        CTTGGGCGAGAGTAGTACTAGGATGGGTGACCTCCTGGGAAGTCCTCGTGTTGCACCCCT
S-7        CTTGGGCGAGAGTAGTACTAGGATGGGTGACCTCCTGGGAAGTCCTCGTGTTGCACCCCT
S-8        CTTGGGCGAGAGTAGTACTAGGATGGGTGACCTCCTGGGAAGTCCTCGTGTTGCACCCCT
S-9        CTTGGGCGAGAGTAGTACTAGGATGGGTGACCTCCTGGGAAGTCCTCGTGTTGCACCCCT
S-10       CTTGGGCGAGAGTAGTACTAGGATGGGTGACCTCCTGGGAAGTCCTCGTGTTGCACCCCT
S-11       CTTGGGCGAGAGTAGTACTAGGATGGGTGACCTCCTGGGAAGTCCTCGTGTTGCACCCCT
S-12       CTTGGGCGAGAGTAGTACTAGGATGGGTGACCTCCTGGGAAGTCCTCGTGTTGCACCCCT
S-13       CTTGGGCGAGAGTAGTACTAGGATGGGTGACCTCCTGGGAAGTCCTCGTGTTGCACCCCT
S-14       CTTGGGCGAGAGTAGTACTAGGATGGGTGACCTCCTGGGAAGTCCTCGTGTTGCACCCCT
S-15       CTTGGGCGAGAGTAGTACTAGGATGGGTGACCTCCTGGGAAGTCCTCGTGTTGCACCCCT
S-16       CTTGGGCGAGAGTAGTACTAGGATGGGTGACCTCCTGGGAAGTCCTCGTGTTGCACCCCT
S-17       CTTGGGCGAGAGTAGTACTAGGATGGGTGACCTCCTGGGAAGTCCTCGTGTTGCACCCCT
S-18       CTTGGGCGAGAGTAGTACTAGGATGGGTGACCTCCTGGGAAGTCCTCGTGTTGCACCCCT
S-19       CTTGGGCGAGAGTAGTACTAGGATGGGTGACCTCCTGGGAAGTCCTCGTGTTGCACCCCT
S-20       CTTGGGCGAGAGTAGTACTAGGATGGGTGACCTCCTGGGAAGTCCTCGTGTTGCACCCCT
           ************************************************************

S-1        TTTTCGCTGCCCGCGCGCACGTCTCTGCGGGTCCTTCCTCCCCCCTGCTCTGGTCTCTTG
S-2        TTTTCGCTGCCCGCGCGCACGTCTCTGCGGGTCCTTCCTCCCCCCTGCTCTGGTCTCTTG
S-3        TTTTCGCTGCCCGCGCGCACGTCTCTGCGGGTCCTTCCTCCCCCCTGCTCTGGTCTCTTG
S-4        TTTTCGCTGCCCGCGCGCACGTCTCTGCGGGTCCTTCCTCCCCCCTGCTCTGGTCTCTTG
S-5        TTTTCGCTGCCCGCGCGCACGTCTCTGCGGGTCCCTCCTCCCCCCTGCTCTGGTCTCTTG
S-6        TTTTCGCTGCCCGCGCGCACGTCTCTGCGGGTCCTTCCTCCCCCCTGCTCTGGTCTCTTG
S-7        TTTTCGCTGCCCGCGCGCACGTCTCTGCGGGTCCTTCCTCCCCCCTGCTCTGGTCTCTTG
S-8        TTTTCGCTGCCCGCGCGCACGTCTCTGCGGGTCCTTCCTCCCCCCTGCTCTGGTCTCTTG
S-9        TTTTCGCTGCCCGCGCGCACGTCTCTGCGGGTCCTTCCTCCCCCCTGCTCTGGTCTCTTG
S-10       TTTTCGCTGCCCGCGCGCACGTCTCTGCGGGTCCTTCCTCCCCCCTGCTCTGGTCTCTTG
S-11       TTTTCGCTGCCCGCGCGCACGTCTCTGCGGGTCCTTCCTCCCCCCTGCTCTGGTCTCTTG
S-12       TTTTCGCTGCCCGCGCGCACGTCTCTGCGGGTCCTTCCTCCCCCCTGCTCTGGTCTCTTG
S-13       TTTTCGCTGCCCGCGCGCACGTCTCTGCGGGTCCTTCCTCCCCCCTGCTCTGGTCTCTTG
S-14       TTTTCGCTGCCCGCGCGCACGTCTCTGCGGGTCCTTCCTCCCCCCTGCTCTGGTCTCTTG
S-15       TTTTCGCTGCCCGCGCGCACGTCTCTGCGGGTCCTCCCTCCCCCCTGCTCTGGTCTCTTG
S-16       TTTTCGCTGCCCGCGCGCACGTCTCTGCGGGTCCTTCCTCCCCCCTGCTCTGGTCTCTTG
S-17       TTTTCGCTGCCCGCGCGCACGTCTCTGCGGGTCCTTCCTCCCCCCTGCTCTGGTCTCTTG
S-18       TTTTCGCTGCCCGCGCGCACGTCTCTGCGGGTCCTTCCTCCCCCCTGCTCTGGTCTCTTG
S-19       TTTTCGCTGCCCGCGCGCACGTCTCTGCGGGTCCTTCCTCCCCCCTGCTCTGGTCTCTTG
S-20       TTTTCGCTGCCCGCGCGCACGTCTCTGCGGGTCCTTCCTCCCCCCTGCTCTGGTCTCTTG
           **********************************..************************

S-1        TTTTCCCCGCCGGCTTCGGCCATCCGGGCGAGCTGCCCCCACGCGTGCAGGTCTCGGGCG
S-2        TTTTCCCCGCCGGCTTCGGCCATCCGGGCGAGCTGCCCCCACGCGTGCAGGTCTCGGGCG
S-3        TTTTCCCCGCCGGCTTCGGCCATCCGGGCGAGCTGCCCCCACGCGTGCAGGTCTCGGGCG
S-4        TTTTCCCCGCCGGCTTCGACCATCCGGGCGAGCTGCCCCCACGCGTGCAGGTCTCGGGCG
S-5        TTTTCCCCGCCGGCTTCGACCATCCGGGCGAGCTGCCCCCACGCGTGCAGGTCTCGGGCG
S-6        TTTTCCCCGCCGGCTTCGGCCATCCGGGCGAGCTGCCCCCACGCGTGCAGGTCTCGGGCG
S-7        TTTTCCCCGCCGGCTTCGGCCATCCGGGCGAGCTGCCCCCACGCGTGCAGGTCTCGGGCG
S-8        TTTTCCCCGCCGGCTTCGGCCATCCGGGCGAGCTGCCCCCACGCGTGCAGGTCTCGGGCG
S-9        TTTTCCCCGCCGGCTTCGGCCATCCGGGCGAGCTGCCCCCGCGCGTGCAGGTCTCGGGCG
S-10       TTTTCCCCGCCGGCTTCGGCCATCCGGGCGAGCTGCCCCCACGCGTGCAGGTCTCGGGCG
S-11       TTTTCCCCGCCGGCTTCGGCCATCCGGGCGAGCTGCCCCCACGCGTGCAGGTCTCGGGCG
S-12       TTTTCCCCGCCGGCTTCGGCCATCCGGGCGAGCTGCCCCCACGCGTGCAGGTCTCGGGCG
S-13       TTTTCCCCGCCGGCTTCGGCCATCCGGGCGAGTTGCCCCCACGCGTGCAGGTCTCGGGCG
S-14       TTTTCCCCGCCGGCTTCGGCCATCCGGGCGAGCTGCCCCCACGCGTGCAGGTCTCGGGCG
S-15       TTTTCCCCGCCGGCTTCGGCCATCCGGGCGAGCTGCCCCCACGCGTGCAGGTCTCGGGCG
S-16       TTTTCCCCGCCGGCTTCGGCCATCCGGGCGAGCTGCCCCCACGCGTGCAGGTCTCGGGCG
S-17       TTTTCCCCGCCGGCTTCGACCATCCGGGCGAGCTGCCCCCACGCGTGCAGGTCTCGGGCG
S-18       TTTTCCCCGCCGGCTTCGGCCATCCGGGCGAGCTGCCCCCACGCGTGCAGGTCTCGGGCG
S-19       TTTTCCCCGCCGGCTTCGGCCATCCGGGCGAGCTGCCCCCACGCGTGCAGGTCTCGGGCG
S-20       TTTTCCCCGCCGGCTTCGGCCATCCGGGCGAGCTGCCCCCACGCGTGCAGGTCTCGGGCG
           ******************.*************.*******.*******************

S-1        CGGAGGGGCTGGCTAGCTATCTTTCGAGGGCGTTTCGGGCAGGTCTTGGGCGCCTGGGGC
S-2        CGGAGGGGCTGGCTAGCTATCTTTCGAGGGCGTTTCGGGCAGGTCTTGGGCGCCTGGGGC
S-3        CGGAGGGGCTGGCTAGCTATCTTTCGAGGGCGTTTCGGGCAGGTCTTGGGCGCCTGGGGC
S-4        CGGAGGGGCTGGCTAGCTATCTTTCGAGGGCGTTTCGGGCAGGTCTTGGGCGCCTGGGGC
S-5        CGGAGGGGCTGGCTAGCTATCTTTCGAGGGCGTTTCGGGCAGGTCTTGGGCGCCTGGGGC
S-6        CGGAGGGGCTGGCTAGCTATCTTTCGAGGGCGTTTCGGGCAGGTCTTGGGCGCCTGGGGC
S-7        CGGAGGGGCTGGCTAGCTATCTTTCGAGGGCGTTTCGGGCAGGTCTTGGGCGCCTGGGGC
S-8        CGGAGGGGCTGGCTAGCTATCTTTCGAGGGCGTTTCGGGCAGGTCTTGGGCGCCTGGGGC
S-9        CGGAGGGGCTGGCTAGCTATCTTTCGAGGGCGTTTCGGGCAGGTCTTGGGCGCCTGGGGC
S-10       CGGAGGGGCTGGCTAGCTATCTTTCGAGGGCGTTTCGGGCAGGTCTTGGGCGCCTGGGGC
S-11       CGGAGGGGCTGGCTAGCTATCTTTCGAGGGCGTTTCGGGCAGGTCTTGGGCGCCTGGGGC
S-12       CGGAGGGGCTGGCTAGCTATCTTTCGAGGGCGTTTCGGGCAGGTCTTGGGCGCCTGGGGC
S-13       CGGAGGGGCTGGCTAGCTATCTTTCGAGGACGTTTCGGGCAGGTCTTGGGCGCCTGGGGC
S-14       CGGAGGGGCTGGCTAGCTATCTTTCGAGGGCGTTTCGGGCAGGTCTTGGGCGCCTGGGGC
S-15       CGGAGGGGCTGGCTAGCTATCTTTCGAGGGCGTTTCGGGCAGGTCTTGGGCGCCTGGGGC
S-16       CGGAGGGGCTGGCTAGCTATCTTTCGAGGGCGTTTCGGGCAGGTCTTGGGCGCCTGGGGC
S-17       CGGAGGGGCTGGCTAGCTATCTTTCGAGGGCGTTTCGGGCAGGTCTTGGGCGCCTGGGGC
S-18       CGGAGGGGCTGGCTAGCTATCTTTCGAGGGCGTTTCGGGCAGGTCTTGGGCGCCTGGGGC
S-19       CGGAGGGGCTGGCTAGCTATCTTTCGAGGGCGTTTCGGGCAGGTCTTGGGCGCCTGGGGC
S-20       CGGAGGGGCTGGCTAGCTATCTTTCGAGGGCGTTTCGGGCAGGTCTTGGGCGCCTGGGGC
           *****************************.******************************

S-1        GGACTCCGGAGTCCACGGGGAGGGGGTTGTCCGCCCGCCTTGGGCACGAGCTCCCCCTGG
S-2        GGACTCCGGAGTCCACGGGGAGGGGGTTGTCCGCCCGCCTTGGGCACGAGCTCCCCCTGG
S-3        GGACTCCGGAGTCCACGGGGAGGGGGTTGTCCGCCCGCCTTGGGCACGAGCTCCCCCTGG
S-4        GGACTCCGGAGTTCACGGGGAGGGGGTTGTCCGCCCGCCTTGGGCACGAGCTCCCCCTGG
S-5        GGACTCCGGAGTCCACGGGGAGGGGGTTGTCCGCCCGCCTTGGGCACGAGCTCCCCCTGG
S-6        GGACTCCGGAGTCCACGGGGAGGGGGTTGTCCGCCCGCCTTGGGCACGAGCTCCCCCTGG
S-7        GGACTCCGGAGTCCACGGGGAGGGGGTTGTCCGCCCGCCTTGGGCACGAGCTCCCCCTGG
S-8        GGACTCCGGAGTCCACGGGGAGGGGGTTGTCCGCCCGCCTTGGGCACGAGCTCCCCCTGG
S-9        GGACTCCGGAGTCCACGGGGAGGGGGTTGTCCGCCCGCCTTGGGCACGAGCTCCCCCTGG
S-10       GGACTCCGGAGTCCACGGGGAGGGGGTTGTCCGCCCGCCTTGGGCACGAGCTCCCCCTGG
S-11       GGACTCCGGAGTCCACGGGGAGGGGGTTGTCCGCCCGCCTTGGGCACGAGCTCCCCCTGG
S-12       GGACTCCGGAGTCCACGGGGAGGGGGTTGCCCGCCCGCCTTGGGCACGAGCTCCCCCTGG
S-13       GGACTCCGGAGTCCACGGGGAGGGGGTTGTCCGCCCGCCTTGGGCACGAGCTCCCCCTGG
S-14       GGACTCCGGAGTCCACGGGGAGGGGGTTGTCCGACCGCCTTGGGCACGAGCTCCCCCTGG
S-15       GGACTCCGGAGTCCACGGGGAGGGGGTTGTCCGCCCGCCTTGGGCACGAGCTCCCCCTGG
S-16       GGACTCCGGAGTCCACGGGGAGGGGGTTGTCCGCCCGCCTTGGGCACGAGCTCCCCCTGG
S-17       GGACTCCGGAGTCCACGGGGAGGGGGTTGTCCGCCCGCCTTGGGCACGAGCTCCCCCTGG
S-18       GGACTCCGGAGTCCACGGGGAGGGGGTTGTCCGCCCGCCTTGGGCACGAGCTCCCCCTGG
S-19       GGACTCCGGAGTCCACGGGGAGGGGGTTGTCCGCCCGCCTTGGGCACGAGCTCCCCCTGG
S-20       GGACTCCGGAGTCCACGGGGAGGGGGTTGTCCGCCCGCCTTGGGCACGAGCTCCCCCTGG
           ************.****************.*** **************************

S-1        CGAGAGGCGTCCCGCATCCGCGCCGCTCGAGGGACACCACGCCGCCCCTTGGCGGTGGGT
S-2        CGAGAGGCGTCCCGCATCCGCGCCGCTCGAGGGACACCACGCCGCCCCTTGGCGGTGGGT
S-3        CGAGAGGCGTCCCGCATTCGCGCCGCTCGAGGGACACCACGCCGCCCCTTGGCGGTGGGT
S-4        CGAGAGGCGTCCCGCATCCGCGCCGCTCGAGGGACACCACGCCGCCCCTTGGCGGTGGGT
S-5        CGAGAGGCGTCCCGCATCCGCGCCGCTCGAGGGACACCACGCCGCCCCTTGGCGGTGGGT
S-6        CGAGAGGCGTCCCGCATCCGCGCCGCTCGAGGGACACTACGCCGCCCCTTGGCGGTGGGT
S-7        CGAGAGGCGTCCCGCATCCGCGCCGCTCAAGGGACACCACGCCGCCCCTTGGCGGTGGGT
S-8        CGAGAGGCGTCCCGCATCCGCGCCGCTCGAGGGACACCACGCCGCCCCTTGGCGGTGGGT
S-9        CGAGAGGCGTCCCGCATCCGCGCCGCTCGAGGGACACCACACCGCCCCTTGGCGGTGGGT
S-10       CGAGAGGCGTCCCGCATCCGCGCCGCTCGAGGGACACCACGCCGCCCCTTGGCGGTGGGT
S-11       CGAGAGGCGTCCCGCATCCGCGCCGCTCGAGGGACACCACGCCGCCCCTTGGCGGTGGGT
S-12       CGAGAGGCGTCCCGCATCCGCGCCGCTCGAGGGACACCACGCCGCCCCTTGGCGGTGGGT
S-13       CGAGAGGCGTCCCGCATCCGCGCCGCTCGAGGGACACCACGCCGCCCCTTGGCGGTGGGT
S-14       CGAGAGGCGTCCCGCATCCGCGCCGCTCGAGGGACACCACGCCGCCCCTTGGCGGTGGGT
S-15       CGAGAGGCGTCCCGCATCCGCGCCGCTCGAGGGACACCACGCCGCCCCTTGGCGGTGGGT
S-16       CGAGAGGCGTCCCGCATCCGCGCCGCTCGAGGGACACCACGCCGCCCCTTGGCGGTGGGT
S-17       CGAGAGGCGTCCCGCATCCGCACCGCTCGAGGGACACCACGCCGCCCCTTGGCGGTGGGT
S-18       CGAGAGGCGTCCCGCATCCGCGCCGCTCGAGGGACACCACGCCGCCCCTTGGCGGTGGGT
S-19       CGAGAGGCGTCCCGCATCCGCGCCGCTCGAGGGACACCACGCCGCCCCTTGGCGGTGGGT
S-20       CGAGAGGCGTCCCGCATCCGCGCCGCTCGAGGGACACCACGCCGCCCCTTGGCGGTGGGT
           *****************.***.******.********.**.*******************

S-1        CAATGATCTCGCCCGCCCCCTCTCCCTCTCCGTCTGTCAGGGAGGCGGCGCCGATGCCGG
S-2        CAATGATCTCGCCCGACCCCTCTCCCTCTCCGTCTGTCAGGGAGGCGGCGCCGATGCCGG
S-3        CAATGATCTCGCCCGCCCCCTCTCCCTCTCCGTCTGTCAGGGAGGCGGCGCCGATGCCGG
S-4        CAATGATCTCGCCCGCCCCCTCTCCCTCTCCGTCTGTCAGGGAGGCGGCGCCGATGCCGG
S-5        CAATGATCTCGCCCGCCCCCTCTCCCTCTCCGTCTGTCAGGGAGGCGGCGCCGATGCCGG
S-6        CAATGATCTCGCCCGCCCCCTCTCCCTCTCCGTCTGTCAGGGAGGCGGCGCCGATGCCGG
S-7        CAATGATCTCGCCCGCCCCCTCTCCCTCTCCGTCTGTCAGGGAGGCGGCGCCGATGCCGG
S-8        CAATGATCTCGCCCGCCCCCTCTCCCTCTCCGTCTGTCAGGGGGGCGGCGCCGATGCCGG
S-9        CAATGATCTCGCCCGCCCCCTCTCCCTCTCCGTCTGTCAGGGAGGCGGCGCCGATGCCGG
S-10       CAATGATCTCGCCCGCCCCCTCTCCCTCTCCGTCTGTCAGGGAGGCGGCGCCGATGCCGG
S-11       CAATGATCTCGCCCGCCCCCTCTCCCTCTCCGTCTGTCAGGGAGGCGGCGCCGATGCCGG
S-12       CAATGATCTCGCCCGCCCCCTCTCCCTCTCCGTCTGTCAGGGAGGCGGCGCCGATGCCGG
S-13       CAATGATCTCGCCCGCCCCCTCTCCCTCTCCGTCTGTCAGGGAGGCGGCGCCGATGCCGG
S-14       CAATGATCTCGCCCGCCCCCTCTCCCTCTCCGTCTGTCAGGGAGGCGGCGCCGATGCCGG
S-15       CAATGATCTCGCCCGCCCCCTCTCCCTCTCCGTCTGTCAGGGAGGCGGCGCCGATGCCGG
S-16       CAATGATCTCGCCCGCCCCCTCTCCCTCTCCGTCTGTCAGGGAGGCGGCGCCGATGCCGG
S-17       CAATGATCTCGCCCGCCCCCTCTCCCTCTCCGTCTGTCAGGGAGGCGGCGCCGATGCCGG
S-18       CAATGATCTCGCCCGCCCCCTCTCCCTCTCCGTCTGTCAGGGAGGCGGCGCCGATGCCGG
S-19       CAATGATCTCGCCCGCCCCCTCTCCCTCTCCGTCTGTCAGGGAGGCGGCGCCGATGCCGG
S-20       CAATGATCTCGCCCGCCCCCTCTCCCTCTCCGTCTGTCAGGGAGGCGGCGCCGATGCCGG
           *************** **************************.*****************

S-1        GGGGGAGAGGATTAGGGCTGACGCGGCGCTTGTCACATCGGGTGCGATCATACCAGCACT
S-2        GGGGGAGAGGATTAGGGCTGACGCGGCGCTTGTCACATTGGGTGCGATCACACCAGCACT
S-3        GGGGGAGAGGATTAGGGCTGACGCGGCGCTTGTCACATCGGGTGCGATCATACCAGCACT
S-4        GGGGGAGAGGATTAGGGCTGACGCGGCGCTTGTCACATCGGGTGCGATCATACCAGCACC
S-5        GGGGGAGAGGATTAGGGCTGACGCGGCGCTTGTCACTTCGGGTGCGATCATACCAGCACT
S-6        GGGGGAGAGGATTAGGGCTGACGCGGCGCTTGTCACATCGGGTGCGATCATACCAGCACC
S-7        GGGGGAGAGGATTAGGGCTGACGCGGCGCTTGTCACATCGGGTGCGATCATACCAGCACT
S-8        GGGGGAGAGGATTAGGGCTGACGCGGCGCTTGTCACATCGGGTGCGATCATACCAGCACT
S-9        GGGGGAGAGGATTAGGGCTGACGCGGCGCTTGTCACATCGGGTGCGATCATACCAGCACT
S-10       GGGGGAGAGGATTAGGGCTGACGCGGCGCTTGTCACATCGGGTGCGATCATACCAGCACT
S-11       GGGGGAGAGGATTAGGGCTGACGCGGCGCTTGTCACATCGGGTGCGATCATACCAGCACT
S-12       GGGGGAGAGGATTAGGGCTGACGCGGCGCTTGTCACATCGGGTGCGATCATACCAGCACT
S-13       GGGGGAGAGGATTAGGGCTGACGCGGTGCTTGTCACATCGGGTGCGATCATACCAGCACT
S-14       GGGGGAGAGGATTAGGGCTGACGCGGCGCTTGTCACATCGGGTGCGATCATACCAGCACT
S-15       GGGGGAGAGGATTAGGGCTGACGCGGCGCTTGTCACATCGGGTGCGATCATACCAGCACT
S-16       GGGGGAGAGGATTAGGGCTGACGCGGCGCTTGTCACATCGGGTGCGATCATACCAGCACT
S-17       GGGGGAGAGGATTAGGGCTGACGCGGCGCTTGTCACATCGGGTGCGATCATACCAGCACT
S-18       GGGGGAGAGGATTAGGGCTGACGCGGCGCTTGTCACATCGGGTGCGATCATACCAGCACT
S-19       GGGGGAGAGGATTAGGGCTGACGCGGCGCTTGTCACATCGGGTGCGATCATACCAGCACT
S-20       GGGGGAGAGGATTAGGGCTGACGCGGCGCTTGTCACATCGGGTGCGATCATACCAGCACT
           **************************.********* *.***********.********.

S-1        AATGCACCGGATCCCATCAGAACTCCGAAGTTAAGCGTG
S-2        AATGCACCGGATCCCATCAGAACTCCGAAGTTAAGCGTG
S-3        AATGCACCGGATCCCATCAGAACTCCGAAGTTAAGCGTG
S-4        AATGCACCGGATCCCATCAGAACTCCGAAGTTAAGCGTG
S-5        AATGCACCGGATCCCATCAGAACTCCGAAGTTAAGCGTG
S-6        AATGCACCGGATCCCATCAGAACTCCGAAGTTAAGCGTG
S-7        AATGCACCGGATCCCATCAGAACTCCGAAGTTAAGCGTG
S-8        AATGCACCGGATCCCATCAGAACTCCGAAGTTAAGCGTG
S-9        AATGCACCGGATCCCATCAGAACTCCGAAGTTAAGCGTG
S-10       AATGCACCGGATCCCATCAGAACTCCGAAGTTAAGCGTG
S-11       AATGCACCGGATCCCATCAGAACTCCGAAGTTAAGCGTG
S-12       AATGCACCGGATCCCATCAGAACTCCGAAGTTAAGCGTG
S-13       AATGCACCGGATCCCATCAGAACTCCGAAGTTAAGCGTG
S-14       AATGCACCGGGTCCCATCAGAACTCCGAAGTTAAGCGTG
S-15       AATGCACCGGATCCCATCAGAACTCCGAAGTTAAGCGTG
S-16       AATGCACCAGATCCCATCAGAACTCCGAAGTTAAGCGTG
S-17       AATGCACCGGATCCCATCAGAACTCCGAAGTTAAGCGTG
S-18       AATGCACCGGATCCCATCAGAACTCCGAAGTTAAGCGTG
S-19       AATGCACCGGATCCCATCAGAACTCCGAAGTTAAGCGTG
S-20       AATGCACCGGATCCCATCAGAACTCCGAAGTTAAGCGTG
           ********.*.****************************
```

**Supplementary Figure 2.** **Nucleotide alignment of twenty sequences (L-1 to L-20) of cloned PCR fragments representing *S.polyrhiza* 5S rDNA repeats of longer type (5S-Sp-L)**. Sequences encoding 5S rRNA are highlighted by green; SNPs in the non-transcribed spacer are marked by magenta, and the GAGA-motifs are highlighted by yellow.

```
L-1        CTTGGGCGAGAGTAGTACTAGGATGGGTGACCTCCTGGGAAGTCCTCGTGTTGCACCCCT
L-2        CTTGGGCGAGAGTAGTACTAGGATGGGTGACCTCCTGGGAAGTCCTCGTGTTGCACCCCT
L-3        CTTGGGCGAGAGTAGTACTAGGATGGGTGACCTCCTGGGAAGTCCTCGTGTTGCACCCCT
L-4        CTTGGGCGAGAGTAGTACTAGGATGGGTGACCTCCTGGGAAGTCCTCGTGTTGCACCCCT
L-5        CTTGGGCGAGAGTAGTACTAGGATGGGTGACCTCCTGGGAAGTCCTCGTGTTGCACCCCT
L-6        CTTGGGCGAGAGTAGTACTAGGATGGGTGACCTCCTGGGAAGTCCTCGTGTTGCACCCCT
L-7        CTTGGGCGAGAGTAGTACTAGGATGGGTGACCTCCTGGGAAGTCCTCGTGTTGCACCCCT
L-8        CTTGGGCGAGAGTAGTACTAGGATGGGTGACCTCCTGGGAAGTCCTCGTGTTGCACCCCT
L-9        CTTGGGCGAGAGTAGTACTAGGATGGGTGACCTCCTGGGAAGTCCTCGTGTTGCACCCCT
L-10       CTTGGGCGAGAGTAGTACTAGGATGGGTGACCTCCTGGGAAGTCCTCGTGTTGCACCCCT
L-11       CTTGGGCGAGAGTAGTACTAGGATGGGTGACCTCCTGGGAAGTCCTCGTGTTGCACCCCT
L-12       CTTGGGCGAGAGTAGTACTAGGATGGGTGACCTCCTGGGAAGTCCTCGTGTTGCACCCCT
L-13       CTTGGGCGAGAGTAGTACTAGGATGGGTGACCTCCTGGGAAGTCCTCGTGTTGCACCCCT
L-14       CTTGGGCGAGAGTAGTACTAGGATGGGTGACCTCCTGGGAAGTCCTCGTGTTGCACCCCT
L-15       CTTGGGCGAGAGTAGTACTAGGATGGGTGACCTCCTGGGAAGTCCTCGTGTTGCACCCCT
L-16       CTTGGGCGAGAGTAGTACTAGGATGGGTGACCTCCTGGGAAGTCCTCGTGTTGCACCCCT
L-17       CTTGGGCGAGAGTAGTACTAGGATGGGTGACCTCCTGGGAAGTCCTCGTGTTGCACCCCT
L-18       CTTGGGCGAGAGTAGTACTAGGATGGGTGACCTCCTGGGAAGTCCTCGTGTTGCACCCCT
L-19       CTTGGGCGAGAGTAGTACTAGGATGGGTGACCTCCTGGGAAGTCCTCGTGTTGCACCCCT
L-20       CTTGGGCGAGAGTAGTACTAGGATGGGTGACCTCCTGGGAAGTCCTCGTGTTGCACCCCT
           ************************************************************

L-1        TTTGCC-------------TGCCGCTCTCCCCGCAGCTTGCAATTGATGTCGCTGCCTCT
L-2        TTTGCC-------------TGCCGCTCTCCCCGCAGCTTGCAATTGATGTCGCTGCCTCT
L-3        TTTGCC-------------TGCCGCTCTCCCCGCAGCTTGCAATTGATGTCGCTGCCTCT
L-4        TTTGCC-------------TGCCGCTCTCCCCGCAGCTTGCAATTGATGTCGCTGCCTCT
L-5        TTTGCC-------------TGCCGCTCTCCCCGCAGCTTGCAATTGATGTCGCTGCCTCT
L-6        TTTGCC-------------TGCCGCTCTCCCCGCAGCTTGCAATTGATGTCGCTGCCTCT
L-7        TTTGCC-------------TGCCGCTCTCCCCGCAGCTTGCAATTGATGTCGCTGCCTCT
L-8        TTTGCC-------------TGCCGCTCTCCCCGCAGCTTGCAATTGATGTCGCTGCCTCT
L-9        TTTGCC-------------TGCCGCTCTCCCCGCAGCTTGCAATTGATGTCGCTGCCTCT
L-10       TTTGCC-------------TGCCGCTCTCCCCGCAGCTTGCAATTGATGTCGCTGCCTCT
L-11       TTTGCC-------------TGCCGCTCTCCCCGCAGCTTGCAATTGATGTCGCTGCCTCT
L-12       TTTGCC-------------TGCCGCTCTCCCCGCAGCTTGCAATTGATGTCGCTGCCTCT
L-13       TTTGCC-------------TGCCGCTCTCCCCGCAGCTTGCAATTGATGTCGCTGCCTCT
L-14       TTTGCC-------------TGCCGCTCTCCCCGCAGCCTGCAATTGATGTCGCTGCCTCT
L-15       TTTGCC-------------TGCCGCTCTCCCCGCAGCTTGCAATTGATGTCGCTGCCTCT
L-16       TTTGCC-------------TGCCGCTCTCCCCGCAGCTTGCAATTGATGTCGCTGCCTCT
L-17       TTTGCC-------------TGCCGCTCTCCCCGCAGCTTGCAATTGATGTCGCTGCCTCT
L-18       TTTGCCTGCCGCAGCTTTGTGCCGCTCTCCCCGCAGCTTGCAATTGATGTCGCTGCCTCT
L-19       TTTGCC-------------TGCCGCTCTCCCCGCAGCTTGCAATTGATGTCGCTGCCTCT
L-20       TTTGCC-------------TGCCGCTCTCCCCGCAGCTTGCAATTGATGTCGCTGCCTCT
           ******             ******************.**********************

L-1        GAGCTTCGTCGTGATCTCCTCACCTAGGGCGAGGGCCATCTTCCGGAGGCCCTTCCTCTG
L-2        GAGCTTCGTCGTGATCTCCTCACCTAGGGCGGGGGCCATCTTCCGGAGGCCCTTCCTCTG
L-3        GAGCTTCGTCGTGATCTCCTCACCTAGGGCGAGGGCCATCTTCCGGAGGCCCTTCCTCTG
L-4        GAGCTTCGTCGTGATCTCCTCACCTAGGGCGAGGGCCATCTTCCGGAGGCCCTTCCTCTG
L-5        GAGCTTCGTCGTGATCTCCTCACCTAGGGCGAGGGCCATCTTCCGGAGGCCCTTCCTCTG
L-6        GAGCTTCGTCGTGATCTCCTCACCTAGGGCGAGGGCCATCTTCCGGAGGCCCTTCCTCTG
L-7        GAGCTTCGTCGTGATCTCCTCACCTAGGGCGAGGGCCATCTTCCGGAGGCCCTTCCTCTG
L-8        GAGCTTCGTCGTGATCTCCTCACCTAGGGCGAGGGCCATCTTCCGGAGGCCCTTCCTCTG
L-9        GAGCTTCGTCGTGATCTCCTCACCTAGGGCGAGGGCCATCTTCCGGAGGCCCTTCCTCTG
L-10       GAGCTTCGTCGTGATCTCCTCACCTAGGGCGAGGGCCATCTTCCGGAGGCCCTTCCTCTG
L-11       GAGCTTCGTCGTGATCTCCTCACCTAGGGCGAGGGCCATCTTCCGGAGGCCCTTCCTCTG
L-12       GAGCTTCGTCGTGATCTCCTCACCTAGGGCGAGGGCCATCTTCCGGAGGCCCTTCCTCTG
L-13       GAGCTTCGTCGTGATCTCCTCACCTAGGGCGAGGGCCATCTTCCGGAGGCCCTTCCTCTG
L-14       GAGCTTCGTCGTGATCTCCTCACCTAGGGCGAGGGCCATCTTCCGGAGGCCCTTCCTCTG
L-15       GAGCTTCGTCGTGATCTCCTCACCTAGGGCGAGGGCCATCTTCCGGAGGCCCTTCCTCTG
L-16       GAGCTTCGTCGTGATCTCCTCACCTAGGGCGAGGGCCATCTTCCGGAGGCCCTTCCTCTG
L-17       GAGCTTCGTCGTGATCTCCTCACCTAGGGCGAGGGCCATCTTCCGGAGGCCCTTCCTCTG
L-18       GAGCTTCGTCGTGATCTCCTCACCTAGGGCGAGGGCCATCTTCCGGAGGCCCTTCCTCTG
L-19       GAGCTTCGTCGTGATCTCCTCACCTAGGGCGAGGGCCATCTTCCGGAGGCCCTTCCTCTG
L-20       GAGCTTCGTCGTGATCTCCTCACCTAGGGCGAGGGCCATCTTCCGGAGGCCCTTCCTCTG
           *******************************.****************************

L-1        GTCAGTTCATCCCCCTCTTTCCCTTCCTCCTCCGAACGCCCCGTCCGCCACACGCGAACT
L-2        GTCAGTTCATCCCCCTCTTTCCCTTCCTCCTCCGAACGCCCCGTCCGCCACACGCGAACT
L-3        GTCAGTTCATCCCCCTCTTTCCCTTCCTCCTCCGAACGCCCCGTCCGCCACGCGCGAACT
L-4        GTCAGTTCATCCCCCTCTTTCCCTTCCTCCTCCGAACGCCCCGTCCGCCACACGCGAACT
L-5        GTCAGTTCATCCCCCTCTTTCCCTTCCTCCTCCGAACGCCCCGTCCGCCACACGCGAACT
L-6        GTCAGTTCATCCCCCTCTTTCCCTTCCTCCTCCGAACGCCCCGTCCGCCACACGCGAACT
L-7        GTCAGTTCATCCCCCTCTTTCCCTTCCTCCTCCGAACGCCCCGTCCGCCACACGCGAACT
L-8        GTCAGTTCATCCCCCTCTTTCCCTTCCTCCTCCGAACGCCCCGTCCGCCACACGCGAACT
L-9        GTCAGTTCATCCCCCTCTTTCCCTTCCTCCTCCGAACGCCCCGTCCGCCACACGCGAACT
L-10       GTCAGTTCATCCCCCTCTTTCCCTTCCTCCTCCGAACGCCCCGTCCGCCACACGCGAACT
L-11       GTCAGTTCATCCCCCTCTTTCCCTTCCTCCTCCGAACGCCCCGTCCGCCACACGCGAACT
L-12       GTCAGTTCATCCCCCTCTTTCCCTTCCTCCTCCGAACGCCCCGTCCGCCACACGCGAACT
L-13       GTCAGTTCATCCCCCTCTTTCCCTTCCTCCTCCGAACGCCCCGTCCGCCACACGCGAACT
L-14       GTCAGTTCATCCCCCTCTTTCCCTTCCTCCTCCGAACGCCCCGTCCGCCACACGCGAACT
L-15       GTCAGTTCATCCCCCTCTTTCCCTTCCTCCTCCGAACGCCCCGTCCGCCACACGCGAACT
L-16       GTCAGTTCATCCCCCTCTTTCCCTTCCTCCTCCGAACGCCTCGTCCGCCACACGCGAACT
L-17       GTCAGTTCATCCCCCTCTTTCCCTTCCTCCTCCGAACGCCTCGTCCGCCACACGCGAACT
L-18       GTCAGTTCATCCCCCTCTTTCCCTTCCTCCTCCGAACGCCTCGTCCGCCACACGCGAACT
L-19       GTCAGTTCATCCCCCTCTTTCCCTTCCTCCTCCGAACGCCTCGTCCGCCACACGCGAACT
L-20       GTCAGTTCATCCCCCTCTTTCCCTTCCTCCTCCGAACGCCTCGTCCGCCACACGCGAACT
           ****************************************.**********.********

L-1        GGTTGCCGCACTACCATTTTGGGTATGCTGGGTCTTTTTTGGAGCTCAGTGGGTGCGGCT
L-2        GGTTGCCGCACTACCATTTTGGGTCTGCTGGGTCTTTTTTGGAGCTCAGTGGGTGCGGCT
L-3        GGTTGCCGCACTACCATTTTGGGTATGCTGGGTCTTTTTTGGAGCTCAGTGGGTGCGGCT
L-4        GGTTGCCGCACTACCATTTTGGGTATGCTGGGTCTTTTTTGGAGCTCAGTGGGTGCGGCT
L-5        GGTTGCCGCACTACCATTTTGGGTATGCTGGGTCTTTTTTGGAGCTCAGTGGGTGCGGCT
L-6        GGTTGCCGCACTACCATTTTGGGTATGCTGGGTCTTTTTTGGAGCTCAGTGGGTGCGGCT
L-7        GGTTGCCGCACTACCATTTTGGGTATGCTGGGTCTTTTTTGGAGCTCAGTGGGTGCGGCT
L-8        GGTTGCCGCACTACCATTTTGGGTATGCTGGGTCTTTTTTGGAGCTCAGTGGGTGCGGCT
L-9        GGTTGCCGCACTACCATTTTGGGTATGCTGGGTCTTTTTTGGAGCTCAGTGGGTGCGGCT
L-10       GGTTGCCGCACTACCATTTTGGGTATGCTGGGTCTTTTTTGGAGCTCAGTGGGTGCGGCT
L-11       GGTTGCCGCACTACCATTTTGGGTATGCTGGGTCTTTTTTGGAGCTCAGTGGGTGCGGCT
L-12       GGTTGCCGCACTACCATTTTGGGTATGCTGGGTCTTTTTTGGAGCTCAGTGGGTGCGGCT
L-13       GGTTGCCGCACTACCATTTTGGGTATGCTGGGTCTTTTTTGGAGCTCATTGGGTGCGGCT
L-14       GGTTGCCGCACTACCATTTTGGGTATGCTGGGTCTTTTTTGGAGCTCAGTGGGTGCGGCT
L-15       GGTTGCCGCACTACCATTTTGGGTATGCTGGGTCTTTTTTGGAGCTCAGTGGGTGCGGCT
L-16       GGTTGCCGCACTACCATTTTGGGTATGCTGGGTCTTTTTTGGAGCTCAGTGGGTGCGGCT
L-17       GGTTGCCGCACTACCATTTTGGGTATGCTGGGTCTTTTTTGGAGCTCAGTGGGTGCGGCT
L-18       GGTTGCCGCACTACCATTTTGGGTATGCTGGGTCTTTTTTGGAGCTCAGTGGGTGCGGCT
L-19       GGTTGCCGCACTACCATTTTGGGTATGCTGGGTCTTTTTTGGAGCTCAGTGGGTGCGGCT
L-20       GGTTGCCGCACTACCATTTTGGGTATGCTGGGTCTTTTTTGGAGCTCAGTGGGTGCGGCT
           ************************ *********************** ***********

L-1        TCCCTTGCAAAAGGTTTGCCTCTTTTATGTTGGCTAGCTAGCTAGTTGGGGTTTGGGCCT
L-2        TCCCTTGCAAAAGGTTTGCCTCTTTTATGTTGGCTAGCTAGCTAGTTGGGGTTTGGGCCT
L-3        TCCCTTGCAAAAGGTTTGCCTCTTTTATGTTGGCTAGCTAGCTAGTTGGGGTTTGGGCCT
L-4        TCCCTTGCAAAAGGTTTGCCTCTTTTATGTTGGCTAGCTAGCTAGTTGGGGTTTGGGCCT
L-5        TCCCTTGCAAAAGGTTTGCCTCTTTTATGTTGGCTAGCTAGCTAGTTGGGGTTTGGGCCT
L-6        TCCCTTGCAAAAGGTTTGCCTCTTTTATGTTGGCTAGCTAGCTAGTTGGGGTTTGGGCCT
L-7        TCCCTTGCAAAAAGTTTGCCTCTTTTATGTTGGCTAGCTAGCTAGTTGGGGTTTGGGCCT
L-8        TCCCTTGCAAAAGGTTTGCCTCTTTTATGTTGGCTAGCTAGCTAGTTGGGGTTTGGGCCT
L-9        TCCCTTGCAAAAGGTTTGCCTCTTTTATGTTGGCTAGCTAGCTAGTTGGGGTTTGGGCCT
L-10       TCCCTTGCAAAAGGTTTGCCTCTTTTATGTTGGCTAGCTAGCTAGTTGGGGTTTGGGCCT
L-11       TCCCTTGCAAAAGGTTTGCCTCTTTTATGTTGGCTAGCTAGCTAGTTGGGGTTTGGGCCT
L-12       TCCCTTGCAAAAGGTTTGCCTCTTTTATGTTGGCTAGCTAGCTAGTTGGGGTTTGGGCCT
L-13       TCCCTTGCAAAAGGTTTGCCTCTTTTATGTTGGCTAGCTAGCTAGTTGGGGTTTGGGCCT
L-14       TCCCTTGCAAAAGGTTTGCCTCTTTTATGTTGGCTAGCTAGCTAGTTGGGGTTTGGGCCT
L-15       TCCCTTGCAAAAGGTTTGCCTCTTTTATGTTGGCTAGCTAGCTAGTTGGGGTTTGGGCCT
L-16       TCCCTTGCAAAAGGTTTGCCTCTTTTATGTTGGCTAGCTAGCTAGTTGGGGTTTGGGCCT
L-17       TCCCTTGCAAAAGGTTTGCCTCTTTTATGTTGGCTAGCTAGCTAGTTGGGGTTTGGGCCT
L-18       TCCCTTGCAAAAGGTTTGCCTCTTTTATGTTGGCTAGCTAGCTAGTTGGGGTTTGGGCCT
L-19       TCCCTTGCAAAAGGTTTGCCTCTTTTATGTTGGCTAGCTAGCTAGTTGGGGTTTGGGCCT
L-20       TCCCTTGCAAAAGGTTTGCCTCTTTTATGTTGGCTAGCTAGCTAGTTGGGGTTTGGGCCT
           ************.***********************************************

L-1        ATTGGGCCCTTTTAGAGGGGGCACCGCCCTCATAAAAATTGCTTGCAATATTCAAATAAC
L-2        ATTGGGCCCTTTTAGAGGGGGCACCGCCCTCATAAAAATTGCTTGCAATATTCAAATAAC
L-3        ATTGGGCCCTTTTAGAGGGGGCACCGCCCTCATAAAAATTGCTTGCAATATTCAAATAAC
L-4        ATTGGGCCCTTTTAGAGGGGGCACCGCCCTCATAAAAATTGCTTGCAATATTCAAATAAC
L-5        ATTGGGCCCTTTTAGAGGGGGCACCGCCCTCATAAAAATTGCTTGCAATATTCAAATAAC
L-6        ATTGGGCCCTTTTAGAGGGGGCACCGCCCTCATAAAAATTGCTTGCAATATTCAAATAAC
L-7        ATTGGGCCCTTTTAGAGGGGGCACCGCCCTCATAAAAATTGCTTGCAATATTCAAATAAC
L-8        ATTGGGCCCTTTTAGAGGGGGCACCGCCCTCATAAAAATTGCTTGCAATATTCAAATAAC
L-9        ATTGGGCCCTTTTAGAGGGGGCACCGCCCTCATAAAAATTGCTTGCAATATTCAAATAAC
L-10       ATTGGGCCCTTTTAGAGGGGGCACCGCCCTCATAAAAATTGCTTGCAATATTCAAATAAC
L-11       ATTGGGCCCTTTTAGAGGGGGCACCGCCCTCATAAAAATTGCTTGCAATATTCAAATAAC
L-12       ATTGGGCCCTTTTAGAGGGGGCACCGCCCTCATAAAAATTGCTTGCAATATTCAAATAAC
L-13       ATTGGGCCCTTTTAGAGGGGGCACCGCCCTCATAAAAATTGCTTGCAATATTCAAATAAC
L-14       ATTGGGCCCTTTTAGAGGGGGCACCGCCCTCATAAAAATTGCTTGCAATATTCAAATAAC
L-15       ATTGGGCCCTTTTAGAGGGGGCACCGCCCTCATAAAAATTGCTTGCAATATTCAAATAAC
L-16       ATTGGGCCCTTTTAGAGGGGGCACCGCCCTCATAAAAATTGCTTGCAATATTCAAATAAC
L-17       ATTGGGCCCTTTTAGAGGGGGCACCGCCCTCATAAAAATTGCTTGCAATATTCAAATAAC
L-18       ATTGGGCCCTTTTAGAGGGGGCACCGCCCTCATAAAAATTGCTTGCAATATTCAAATAAC
L-19       ATTGGGCCCTTTTAGAGGGGGCACCGCCCTCATAAAAATTGCTTGCAATATTCAAATAAC
L-20       ATTGGGCCCTTTTAGAGGGGGCACCGCCCTCATAAAAATTGCTTGCAATATTCAAATAAC
           ************************************************************

L-1        TTGGGCATGAAACGCGACGGGAGGAAAAGAGAAAATAGGGCTATAAAGTGTGGAATATTA
L-2        TTGGGCATGAAACGCGACGGGAGGAAAAGAGAAAATAGGGCTATAAAGTGTGGAATATTA
L-3        TTGGGCATGAAACGCGACGGGAGGAAAGGAGAAAATAGGGCTATAAAGTGTGGAATATTA
L-4        TTGGGCATGAAACGCGACGGGAGGAAAAGAGAAAATAGGGCTATAAAGTGTGGAATATTA
L-5        TTGGGCATGAAACGCGACGGGAGGAAAAGAGAAAATAGGGCTATAAAGTGTGGAATATTA
L-6        TTGGGCATGAAACGCGACGGGAGGAAAAGAGAAAATAGGGCTATAAAGTGTGGAATATTA
L-7        TTGGGCATGAAACGCGACGGGAGGAAAAGAGAAAATAGGGCTATAAAGTGTGGAATATTA
L-8        TTGGGCATGAAACGCGACGGGAGGAAAAGAGAAAATAGGGCTATAAAGTGTGGAATATTA
L-9        TTGGGCATGAAACGCGACGGGAGGAAAAGAGAAAATAGGGCTATAAAGTGTGGAATATTA
L-10       TTGGGCATGAAACGCGACGGGAGGAAAAGAGAAAATAGGGCTATAAAGTGTGGAATATTA
L-11       TTGGGCATGAAACGCGACGGGAGGAAAAGAGAAAATAGGGCTATAAAGTGTGGAATATTA
L-12       TTGGGCATGAAACGCGACGGGAGGAAAAGAGAAAATAGGGCTATAAAGTGTGGAATATTA
L-13       TTGGGCATGAAACGCGACGGGAGGAAAAGAGAAAATAGGGCTATAAAGTGTGGAATATTA
L-14       TTGGGCATGAAACGCGACGGGAGGAAAAGAGAAAATAGGGCTATAAAGTGTGGAATATTA
L-15       TTGGGCATGAAACGCGACGGGAGGAAAAGAGAAAATAGGGCTATAAAGTGTGGAATATTA
L-16       TTGGGCATGAAACGCGACGGGAGGAAAAGAGAAAATAGGGCTATAAAGTGTGGAATATTA
L-17       TTGGGCATGAAACGCGACGGGAGGAAAAGAGAAAATAGGGCTATAAAGTGTGGAGTATTA
L-18       TTGGGCATGAAACGCGACGGGAGGAAAAGAGAAAATAGGGCTATAAAGTGTGGAATATTA
L-19       TTGGGCATGAAACGCGACGGGAGGAAAAGAGAAAATAGGGCTATAAAGTGTGGAATATTA
L-20       TTGGGCATGAAACGCGACGGGAGGAAAAGAGAAAATAGGGCTATAAAGTGTGGAATATTA
           ***************************.**************************.*****

L-1        AAGGAGGTTAGAGGAAGATATTAAATCGGAGGATGCGGATAGAAGATAGAGAAAGTCACG
L-2        AAGGAGGTTAGAGGAAGATATTAAATCGGAGGATGCGGATAGAAGATAGAGAAAGTCACG
L-3        AAGGAGGTTAGAGGAAGATATTAAATCGGAGGATGCGGATAGAAGATAGAGAAAGTCACG
L-4        AAGGAGGTTAGAGGAAGATATTAAATCGGAGGATGCGGATAGAAGATAGAGAAAGTCACG
L-5        AAGGAGGTTAGAGGAAGATATTAAATCGGAGGATGCGGATAGAAGATAGAGAAAGTCACG
L-6        AAGGAGGTTAGAGGAAGATATTAAATCGGAGGATGCGGATAGAAGATAGAGAAAGTCACG
L-7        AAGGAGGTTAGAGGAAGATATTAAATCGGAGGATGCGGATAGAAGATAGAGAAAGTCACG
L-8        AAGGAGGTTAGAGGAAGATATTAAATCGGAGGATGCGGATAGAAGATAGAGAAAGTCACG
L-9        AAGGAGGTTAGAGGAAGATATTAAATCGGAGGATGCGGATAGAAGATAGAGAAAGTCACG
L-10       AAGGAGGTTAGAGGAAGATATTAAATCGGAGGATGCGGATAGAAGATAGAGAAAGTCACG
L-11       AAGGAGGTTAGAGGAAGATATTAAATCGGAGGATGCGGATAGAAGATAGAGAAAGTCACG
L-12       AAGGAGGTTAGAGGAAGATATTAAATCGGAGGATGCGGATAGAAGATAGAGAAAGTCACG
L-13       AAGGAGGTTAGAGGAAGATATTAAATCGGAGGATGCGGATAGAAGATAGAGAAAGTCACG
L-14       AAGGAGGTTAGAGGAAGATATTAAATCGGAGGATGCGGATAGAAGATAGAGAAAGTCACG
L-15       AAGGAGGTTAGAGGAAGATATTAAATCGGAGGATGCGGATAGAAGATAGAGAAAGTCACG
L-16       AAGGAGGTTAGAGGAAGATATTAAATCGGAGGATGCGGATAGAAGATAGAGTAAGTCACG
L-17       AAGGAGGTTAGAGGAAGATATTAAATCGGAGGATGCGGATAGAAGATAGAGAAAGTCACG
L-18       AAGGAGGTTAGAGGAAGATATTAAATCGGAGGATGCGGATAGAAGATAGAGAAAGTCACG
L-19       AAGGAGGTTAGAGGAAGATATTAAATCGGAGGATGCGGATAGAAGATAGAGAAAGTCACG
L-20       AAGGAGGTTAGAGGAAGATATTAAATCGGAGGATGCGGATAGAAGATAGAGAAAGTCACG
           *************************************************** ********

L-1        CTAGAAATATTTCCATTTCGGAGTAGAAGCATGAATGGGAATGCGTGAGTGTTGATATCT
L-2        CTAGAAATATTTCCATTTCGGAGTAGAAGCATGAATGGGAATGCGTGAGTGTTGATATCT
L-3        CTAGAAATATTTCCATTTCGGAGTAGAAGCATGAGTGGGAATGCGTGAGTGTTGATATCT
L-4        CTAGAAATATTTCCATTTCGGAGTAGAAGCATGAATGGGAATGCGTGAGTGTTGATATCT
L-5        CTAGAAATATTTCCATTTCGGAGTAGAAGCATGAATGGGAATGCGTGAGTGTTGATATCT
L-6        CTAGAAATATTTCCATTTCGGAGTAGAAGCATGAATGGGAATGCGTGAGTGTTGATATCT
L-7        CTAGAAATATTTCCATTTCGGAGTAGAAGCATGAATGGGAATGCGTGAGTGTTGATATCT
L-8        CTAGAAATATTTCCATTTCGGAGTAGAAGCATGAATGGGAATGCGTGAGTGTTGATATCT
L-9        CTAGAAATATTTCCATTTCGGAGTAGAAGCATGAATGGGAATGCGTGAGTGTTGATATCT
L-10       CTAGAAATATTTCCATTTCGGAGTAGAAGCATGAATGGGAATGCGTGAGTGTTGATATCT
L-11       CTAGAAATATTTCCATTTCGGAGTAGAAGCATGAATGGGAATGCGTGAGTGTTGATATCT
L-12       CTAGAAATATTTCCATTTCGGAGTAGAAGCATGAATGGGAATGCGTGAGTGTTGATATCT
L-13       CTAGAAATATTTCCATTTCGGAGTAGAAGCATGAATGGGAATGCGTGAGTGTTGATATCT
L-14       CTAGAAATATTTCCATTTCGGAGTAGAAGCATGAATGGGAATGCGTGAGTGTTGATATCT
L-15       CTAGAAATATTTCCATTTCGGAGTAGAAGCATGAATGGGAATGCGTGAGTGTTGATATCT
L-16       CTAGAAATATTTCCATTTCGGAGTAGAAGCATGAATGGGAATGCGTGAGTGTTGATATCT
L-17       CTAGAAATATTTCCATTTCGGAGTAGAAGCATGAATGGGAATGCGTGAGTGTTGATATCT
L-18       CTAGAAATATTTCCATTTCGGAGTAGAAGCATGAATGGGAATGCGTGAGTGTTGATATCT
L-19       CTAGAAATATTTCCATTTCGGAGTAGAAGCATGAATGGGAATGCGTGAGTGTTGATATCT
L-20       CTAGAAATATTTCCATTTCGGAGTAGAAGCATGAATGGGAATGCGTGAGTGTTGATATCT
           **********************************.*************************

L-1        ATGCATAGATATCTAAGAGAGAGAGAGAGAGAATTTAGAGTGGAGCTATCCACTTTGAGG
L-2        ATGCATAGATATCTAAGAGAGAGAGAGAGAGAATTTAGAGTGGAGCTATCCACTTTGAGG
L-3        ATGCATAGATATCTAAGAGAGAGAGAGAGAGAATTTAGAGTGGAGCTATCCACTTTGAGG
L-4        ATGCATAGATATCTAAGAGAGAGAGAGAGAGAATTTAGAGTGGAGCTATCCACTTTGAGG
L-5        ATGCATAGATATCTAAGAGAGAGAGAGAGAGAATTTAGAGTGGAGCTATCCACTTTGAGG
L-6        ATGCATAGATATCTAAGAGAGAGAGAGAGAGAATTTAGAGTGGAGCTATCCACTTTGAGG
L-7        ATGCATAGATATCTAAGAGAGAGAGAGAGAGAATTTAGAGTGGAGCTATCCACTTTGAGG
L-8        ATGCATAGATATCTAAGAGAGAGAGAGAGAGAATTTAGAGTGGAGCTATCCACTTTGAGG
L-9        ATGCATAGATATCTAAGAGAGAGAGAGAGAGAATTTAGAGTGGAGCTATCCACTTTGAGG
L-10       ATGCATAGATATCTAAGAGAGAGAGAGAGAGAATTTAGAGTGGAGCTATCCACTTTGAGG
L-11       ATGCATAGATATCTAAGAGAGAGAGAGAGAGAATTTAGAGTGGAGCTATCCACTTTGAGG
L-12       ATGCATAGATATCTAAGAGAGAGAGAGAGAGAATTTAGAGTGGAGCTATCCACTTTGAGG
L-13       ATGCATAGATATCTAAGAGAGAGAGAGAGAGAATTTAGAGTGGAGCTATCCACTTTGAGG
L-14       ATGCATAGATATCTAAGAGAGAGAGAGAGAGAATTTAGAGTGGAGCTATCCACTTTGAGG
L-15       ATGCATAGATATCTAAGAGAGAGAGAGAGAGAATTTAGAGTGGAGCTATCCACTTTGAGG
L-16       ATGCATAGATATCTAAGAGAGAGAGAGAGAGAATTTAGAGTGGAGCTATCCACTTTGAGG
L-17       ATGCATAGATATCTAAGAGAGAGAGAGGGAGAATTTAGAGTGGAGCTATCCACTTTGAGG
L-18       ATGCATAGATATCTAAGAGAGAGAGAGAGAGAATTTAGAGTGGAGCTATCCACTTTGAGG
L-19       ATGCATAGATATCTAAGAGAGAGAGAGAGAGAATTTAGAGTGGAGCTATCCACTTTGAGG
L-20       ATGCATAGATATCTAAGAGAGAGAGAGAGAGAATTTAGAGTGGAGCTATCCACTTTGAGG
           ***************************.********************************

L-1        CTCTAGCTATTTCTAACCCTTAATTAGTTCACATATGGGTGACTTTATCATCAACTGATG
L-2        CTCTAGCTATTTCTAACCCTTAATTAGTTCACATATGGGTGACTTTATCATCAACTGATG
L-3        CTCTAGCTATTTCTAACCCTTAATTAGTTCACATATGGGTGACTTTATCATCAACTGATG
L-4        CTCTAGCTATTTCTAACCCTTAATTAGTTCACATATGGGTGACTTTATCATCAACTGATG
L-5        CTCTAGCTATTTCTAACCCTTAATTAGTTCACATATGGGTGACTTTATCATCAACTGATG
L-6        CTCTAGCTATTTCTAACCCTTAATTAGTTCACATATGGGTGACTTTATCATCAACTGATG
L-7        CTCTAGCTATTTCTAACCCTTAATTAGTTCACATATGGGTGACTTTATCATCAACTGATG
L-8        CTCTAGCTATTTCTAACCCTTAATTAGTTCACATATGGGTGACTTTATCATCAACTGATG
L-9        CTCTAGCTATTTCTAACCCTTAATTAGTTCACATATGGGTGACTTTATCATCAACTGATG
L-10       CTCTAGCTATTTCTAACCCTTAATTAGTTCACATATGGGTGACTTTATCATCAACTGATG
L-11       CTCTAGCTATTTCTAACCCTTAATTAGTTCACATATGGGTGACTTTATCATCAACTGATG
L-12       CTCTAGCTATTTCTAACCCTTAATTAGTTCACATATGGGTGACTTTATCATCAACTGATG
L-13       CTCTAGCTATTTCTAACCCTTAATTAGTTCACATATGGGTGACTTTATCATCAACTGATG
L-14       CTCTAGCTATTTCTAACCCTTAATTAGTTCACATATGGGTGACTTTATCATCAACTGATG
L-15       CTCTAGCTATTTCTAACCCTTAATTAGTTCACATATGGGTGACTTTATCATCAACTGATG
L-16       CTCTAGCTATTTCTAACCCTTAATTAGTTCACATATGGGTGACTTTATCATCAACTGATG
L-17       CTCTAGCTATTTCTAACCCTTAATTAGTTCACATATGGGTGACTTTATCATCAACTGATG
L-18       CTCTAGCTATTTCTAACCCTTAATTAGTTCACATATGGGTGACTTTATCATCAACTGATG
L-19       CTCTAGCTATTTCTAACCCTTAATTAGTTCACATATGGGTGACTTTATCATCAACTGATG
L-20       CTCTAGCTATTTCTAACCCTTAATTAGTTCACATATGGGTGACTTTATCATCAACTGATG
           ************************************************************

L-1        TGGCCCCATCACTCGCCGCAAACCATCTCTTGAGCCCCCTCCTCTTCAGGGCGAATTTCG
L-2        TGGCCCCATCACTCGCCGCAAACCATCTCTTGAGCCCCCTCCTCTTCAGGGCGAATTTCG
L-3        TGGCCCCATCACTCGCCGCAAACCATCTCTTGAGCCCCCTCCTCTTCAGGGCGAATTTCG
L-4        TGGCCCCATCACTCGCCGCAAACCATCTCTTGAGCCCCCTCCTCTTCAGGGCGAATTTCG
L-5        TGGCCCCATCACTCGCCGCAAACCATCTCTTGAGCCCCCTCCTCTTCAGGGCGAATTTCG
L-6        TGGCCCCATCACTCGCCGCAAACCATCTCTTGAGCCCCCTCCTCTTCAGGGCGAATTTCG
L-7        TGGCCCCATCACTCGCCGCAAACCATCTCTTGAGCCCCCTCCTCTTCAGGGCGAATTTCG
L-8        TGGCCCCATCACTCGCCGCAAACCATCTCTTGAGCCCCCTCCTCTTCAGGGCGAATTTCG
L-9        TGGCCCCATCACTCGCCGCAAACCATCTCTTGAGCCCCCTCCTCTTCAGGGCGAATTTCG
L-10       TGGCCCCATCACTCGCCGCAAACCATCTCTTGAGCCCCCTCCTCTTCAGGGCGAATTTCG
L-11       TGGCCCCATCACTCGCCGCAAACCATCTCTTGAGCCCCCTCCTCTTCAGGGCGAATTTCG
L-12       TGGCCCCATCACTCGCCGCAAACCATCTCTTGAGCCCCCTCCTCTTCAGGGCGAATTTCG
L-13       TGGCCCCATCACTCGCCGCAAACCATCTCTTGAGCCCCCTCCTCTTCAGGGCGAACTTCG
L-14       TGGCCCCATCACTCGCCGCAAACCATCTCTTGAGCCCCCTCCTCTTCAGGGCGAATTTCG
L-15       TGGCCCCATCACTCGCCGCAAACCATCTCTTGAGCCCCCTCCTCTTCAGGGCGAATTTCG
L-16       TGGCCCCATCACTCGCCGCAAACCATCTCTTGAGCCCCCTCCTCTTCAGGGCGAATTTCG
L-17       TGGCCCCATCACTCGCCGCAAACCATCTCTTGAGCCCCCTCCTCTTCAGGGCGAATTTCG
L-18       TGGCCCCATCACTCGCCGCAAACCATCTCTTGAGCCCCCTCCTCTTCAGGGCGAATTTCG
L-19       TGGCCCCATCACTCGCCGCAAACCATCTCTTGAGCCCCCTCCTCTTCAGGGCGAATTTCG
L-20       TGGCCCCATCACTCGCCGCAAACCATCTCTTGAGCCCCCTCCTCTTCAGGGCGAATTTCG
           *******************************************************.****

L-1        TCGCCGCCCCGAAGCGTTCTCGTGGGCTCCCTACGTCTGGCCTTTGACCGATCCGTGGAT
L-2        TCGCCGCCCCGAAGCGTTCTCGTGGGCTCCCTACGTCTGGCCTTTGACCGATCCGTGGAT
L-3        TCGCCGCCCCGAAGCGTTCTCGTGGGCTCCCTACGTCTGGCCTTTGACCGATCCGTGGAT
L-4        TCGCCGCCCCGAAGCGTTCTCGTGGGCTCCCTACGTCTGGCCTTTGACCGATCCGTGGAT
L-5        TCGCCGCCCCGAAGCGTTCTCGTGGGCTCCCTACGTCTGGCCTTTGACCGATCCGTGGAT
L-6        TCGCCGCCCCGAAGCGTTCTCGTGGGCTCCCTACGTCTGGCCTTTGACCGATCCGTGGAT
L-7        TCGCCGCCCCGAAGCGTTCTCGTGGGCTCCCTACGTCTGGCCTTTGACCGATCCGTGGAT
L-8        TCGCCGCCCCGAAGCGTTCTCGTGGGCTCCCTACGTCTGGCCTTTGACCGATCCGTGGAT
L-9        TCGCCGCCCCGAAGCGTTCTCGTGGGCTCCCTACGTCTGGCCTTTGACCGATCCGTGGAT
L-10       TCGCCGCCCCGAAGCGTTCTCGTGGGCTCCCTACGTCTGGCCTTTGACCGATCCGTGGAT
L-11       CCGCCGCCCCGAAGCGTTCTCGTGGGCTCCCTACGTCTGGCCTTTGACCGATCCGTGGAT
L-12       TCGCCGCCCCGAAGCGTTCTCGTGGGCTCCCTACGTCTGGCCTTTGACCGATCCGTGGAT
L-13       TCGCCGCCCCGAAGCGTTCTCGTGGGCTCCCTACGTCTGGCCTTTGACCGATCCGTGGAT
L-14       TCGCCGCCCCGAAGCGTTCTCGTGGGCTCCCTACGTCTGGCCTTTGACCGATCCGTGGAT
L-15       TCGCCGCCCCGAAGCGTTCTCGTGGGCTCCCTACGTCTGGCCTTTGACCGATCCGTGGAT
L-16       TCGCCGCCCCGAAGCGTTCTCGTGGGCTCCCTACGTCTGGCCTTTGACCGATCCGTGGAT
L-17       TCGCCGCCCCGAAGCGTTCTCGTGGGCTCCCTACGTCTGGCCTTTGACCGATCCGTGGAT
L-18       TCGCCGCCCCGAAGCGTTCTCGTGGGCTCCCTACGTCTGGCCTTTGACCGATCCGTGGAT
L-19       TCGCCGCCCCGAAGCGTTCTCGTGGGCTCCCTACGTCTGGCCTTTGACCGATCCGTGGAT
L-20       TCGCCGCCCCGAAGCGTTCTCGTGGGCTCCCTACGTCTGGCCTTTGACCGATCCGTGGAT
           .***********************************************************

L-1        GTCTTGCCACCCTCCCACGCGCAGAGTAGCTGCCTTTCGGGAGCGTTTTGGGCAGGTCTT
L-2        GTCTTGCCACCCTCCCACGCGCAGAGTAGCTGCCTTTCGGGAGCGTTTTGGGCAGGTCTT
L-3        GTCTTGCCACCCTCCCACGCGCAGAGTAGCTGCCTTTCGGGAGCGTTTTGGGCAGGTCTT
L-4        GTCTTGCCACCCTCCCACGCGCAGAGTAGCTGCCTTTCGGGAGCGTTTTGGGCAGGTCTT
L-5        GTCTTGCCACCCTCCCACGCGCAGAGTAGCTGCCTTTCGGGAGCGTTTTGGGCAGGTCTT
L-6        GTCTTGCCACCCTCCCACGCGCAGAGTAGCTGCCTTTCGGGAGCGTTTTGGGCAGGTCTT
L-7        GTCTTGCCACCCTCCCACGCGCAGAGTAGCTGCCTTTCGGGAGCGTTTTGGGCAGGTCTT
L-8        GTCTTGCCACCCTCCCACGCGCAGAGTAGCTGCCTTTCGGGAGCGTTTTGGGCAGGTCTT
L-9        GTCTTGCCACCCTCCCACGCGCAGAGTAGCTGCCTTTCGGGAGCGTTTTGGGCAGGTCTT
L-10       GTCTTGCCACCCTCCCACGCGCAGAGTAGCTGCCTTTCGGGAGCGTTTTGGGCAGGTCTT
L-11       GTCTTGCCACCCTCCCACGCGCAGAGTAGCTGCCTTTCGGGAGCGTTTTGGGCAGGTCTT
L-12       GTCTTGCCACCCTCCCACGCGCAGAGTAGCTGCCTTTCGGGAGCGTTTTGGGCAGGTCTT
L-13       GTCTTGCCACCCTCCCACGCGCAGAGTAGCTGCCTTTCGGGAGCGTTTTGGGCAGGTCTT
L-14       GTCTTGCCACCCTCCCACGCGCAGAGTAGCTGCCTTTCGGGAGCGTTTTGGGCAGGTCTT
L-15       GTCTTGCCACCCTCCCACGCGCAGAGTAGCTGCCTTTCGGGAGCGTTTTGGGCAGGTCTT
L-16       GTCTTGCCACCCTCCCACGCGCAGAGTAGCTGCCTTTCGGGAGCGTTTTGGGCGGGTCTT
L-17       GTCTTGCCACCCTCCTACGCGCAGAGTAGCTGCCTTTCGGGAGCGTTTTGGGCAGGTCTT
L-18       GTCTTGCCACCCTCCCACGCGCAGAGTAGCTGCCTTTCGGGAGCGTTTTGGGCAGGTCTT
L-19       GTCTTGCCACCCTCCCACGCGCAGAGTAGCTGCCTTTCGGGAGCGTTTTGGGCAGGTCTT
L-20       GTCTTGCCACCCTCCCACGCGCAGAGTAGCTGCCTTTCGGGAGCGTTTTGGGCAGGTCTT
           ***************.*************************************.******

L-1        GGGCACGTAGAGTGGACCTGAGAGTTGATGGGGACTTGAGGTGGGCTACCTGCTCGTCCT
L-2        GGGCACGTAGAGTGGACCTGAGAGTTGATGGGGACTTGAGGTGGGCTACCTGCTCGTCCT
L-3        GGGCACGTAGAGTGGACCTGAGAGTTGATGGGGACTTGAGGTGGGCTACCTGCTCGTCCT
L-4        GGGCACGTAGAGTGGACCTGAGAGTTGATGGGGACTTGAGGTGGGCTACCTGCTCGTCCT
L-5        GGGCACGTAGAGTGGACCTGAGAGTTGATGGGGACTTGAGGTGGGCTACCTGCTCGTCCT
L-6        GGGCACGTAGAGTGGACCTGAGAGTTGATGGGGACTTGAGGTGGGCTACCTGCTCGTCCT
L-7        GGGCACGTAGAGTGGACCTGAGAGTTGATGGGGACTTGAGGTGGGCTACCTGCTCGTCCT
L-8        GGGCACGTAGAGTGGACCTGAGAGTTGATGGGGACTTGAGGTGGGCTACCTGCTCGTCCT
L-9        GGGCACGTAGAGTGGACCTGAGAGTTGATGGGGACTTGAGGTGGGCTACCTGCTCGTCCT
L-10       GGGCACGTAGAGTGGACCTGAGAGTTGATGGGGACTTGAGGTGGGCTACCTGCTCGTCCT
L-11       GGGCACGTAGAGTGGACCTGAGAGTTGATGGGGACTTGAGGTGGGCTACCTGCTCGTCCT
L-12       GGGCACGTAGAGTGGACCTGAGAGTTGATGGGGACTTGAGGTGGGCTACCTGCTCGTCCT
L-13       GGGCACGTAGAGTGGACCTGAGAGTTGATGGGGACTTGAGGTGGGCTACCTGCTCGTCCT
L-14       GGGCACGTAGAGTGGACCTGAGAGTTGATGGGGACTTGAGGTGGGCTACCTGCTCGTCCT
L-15       GGGCACGTAGAGTGGACCTGAGAGTTGATGGGGACTTGAGGTGGGCTACCTGCTCGTCCT
L-16       GGGCACGTAGAGTGGACCTGAGAGTTGATGGGGACTTGAGGTGGGCTACCTGCTCGTCCT
L-17       GGGCACGTAGAGTGGACCTGAGAGTTGATGGGGACTTGAGGTGGGCTACCTGCTCGTCCT
L-18       GGGCACGTAGAGTGGACCTGAGAGTTGATGGGGACTTGAGGTGGGCTACCTGCTCGTCCT
L-19       GGGCACGTAGAGTGGACCTGAGAGTTGATGGGGACTTGAGGTGGGCTACCTGCTCGTCCT
L-20       GGGCACGTAGAGTGGACCTGAGAGTTGATGGGGACTTGAGGTGGGCTACCTGCTCGTCCT
           ************************************************************

L-1        ATGCACGATTCCCCCTGGCGAGGGACGCGCCGCCTTGGGGGTCGGTCATGTCACCGGCCC
L-2        ATGCACGATTCCCCCTGGCGAGGGACGCGCCGCCTTGGGGGTCGGTCATGTCACCGGCCC
L-3        ATGCACGATTCCCCCTGGCGAGGGACGCGCCGCCTTGGGGGTCGGTCATGTCACCGGCCC
L-4        ATGCACGATTCCCCCTGGCGAGGGACGCGCCGCCTTGGGGGTCGGTCATGTCACCGGCCC
L-5        ATGCACGATTCCCCCTGGCGAGGGACGCGCCGCCTTGGGGGTCGGTCATGTCACCGGCCC
L-6        ATGCACGATTCCCCCTGGCGAGGGACGCGCCGCCTTGGGGGTCGGTCATGTCACCGGCCC
L-7        ATGCACGATTCCCCCTGGCGAGGGACGCGCCGCCTTGGGGGTCGGTCATGTCACCGGCCC
L-8        ATGCACGATTCCCCCTGGCGAGGGACGCGCCGCCTTGGGGGTCGGTCATGTCACCGGCCC
L-9        ATGCACGATTCCCCCTGGCGAGGGACGCGCCGCCTTGGGGGTCGGTCATGTCACCGGCCC
L-10       ATGCACGATTCCCCCTGGCGAGGGACGCGCCGCCTTGGGGGTCGGTCATGTCACCGGCCC
L-11       ATGCACGATTCCCCCTGGCGAGGGACGCGCCGCCTTGGGGGTCGGTCATGTCACCGGCCC
L-12       ATGCACGATTCCCCCTGGCGAGGGACGCGCCGCCTTGGGGGTCGGTCATGTCACCGGCCC
L-13       ATGCACGATTCCCCCGGGCGAGGGACGCGCCGCCTTGGGGGTCGGTCATGTCACCGGCCC
L-14       ATGCACGATTCCCCCTGGCGAGGGACGCGCCGCCTTGGGGGTCGGTCATGTCACCGGCCC
L-15       ATGCACGATTCCCCCTGGCGAGGGACGCGCCGCCTTGGGGGTCGGTCATGTCACCGGCCC
L-16       ATGCACGATTCCCCCTGGCGAGGGACGCGCCGCCTTGGGGGTCGGTCATGTCACCGGCCC
L-17       ATGCACGATTCCCCCTGGCGAGGGACGCGCCGCCTTGGGGGTCGGTCATGTCACCGGCCC
L-18       ATGCACGATTCCCCCTGGCGAGGGACGCGCCGCCTTGGGGGTCGGTCATGTCACCGGCCC
L-19       ATGCACGATTCCCCCTGGCGAGGGACGCGCCGCCTTGGGGGTCGGTCATGTCACCGGCCC
L-20       ATGCACGATTCCCCCTGGCGAGGGACGCGCCGCCTTGGGGGTCGGTCATGTCACCGGCCC
           *************** ********************************************

L-1        TCTCCCTCTAGCGGGGCGGAAATGATCCGGTGCCACGTGGTGTTGGAGAGGGGGCAATCT
L-2        TCTCCCTCTAGCGGGGCGGAAATGATCCGGTGCCACGTGGTGTTGGAGAGGGGGCAATCT
L-3        TCTCCCTCTAGCGGGGCGGAAATGATCCGGTGCCACGTGGTGTTGGAGAGGGGGCAATCT
L-4        TCTCCCTCTAGCGGGGCGGAAATGATCCGGTGCCACGTGGTGTTGGAGAGGGGGCAATCT
L-5        TCTCCCTCTAGCGGGGCGGAAATGATCCGGTGCCACGTGGTGTTGGAGAGGGGGCAATCT
L-6        TCTCCCTCTAGCGGGGCGGAAATGATCCGGTGCCACGTGGTGTTGGAGAGGGGGCAATCT
L-7        TCTCCCTCTAGCGGGGCGGAAATGATCCGGTGCCACGTGGTGTTGGAGAGGGGGCAATCT
L-8        TCTCCCTCTAGCGGGGCGGAAATGATCCGGTGCCACGTGGTGTTGGAGAGGGGGCAATCT
L-9        TCTCCCTCTAGCGGGGCGGAAATGATCCGGTGCCACGTGGTGTTGGAGAGGGGGCAATCT
L-10       TCTCCCTCTAGCGGGGCGGAAATGATCCGGTGCCACGTGGTGTTGGAGAGGGGGCAATCT
L-11       TCTCCCTCTAGCGGGGCGGAAATGATCCGGTGCCACGTGGTGTTGGAGAGGGGGCAATCT
L-12       TCTCCCTCTAGCGGGGCGGAAATGATCCGGTGCCACGTGGTATTGGAGAGGGGGCAATCT
L-13       TCTCCCTCGAGCGGGGCGGAAATGATCCGGTGCCACGTGGTGTTGGAGAGGGGGCAATCT
L-14       TCTCCCTCTAGCGGGGCGGAAATGATCCGGTGCCACGTGGTGTTGGAGAGGGGGCAATCT
L-15       TCTCCCTCTAGCGGGGCGGAAATGATCCGGTGCCACGTGGTGTTGGAGAGGGGGCAATCT
L-16       TCTCCCTCTAGCGGGGCGGAAATGATCCGGTGCCACGTGGTGTTGGAGAGGGGGCAATCT
L-17       TCTCCCTCTAGCGGGGCGGAAATGATCCGGTGCCACGTGGTGTTGGAGAGGGGGCAATCT
L-18       TCTCCCTCTAGCGGGGCGGAAATGATCCGGTGCCACGTGGTGTTGGAGAGGGGGCAATCT
L-19       TCTCCCTCTAGCGGGGCGGAAATGATCCGGTGCCACGTGGTGTTGGAGAGGGGGCAATCT
L-20       TCTCCCTCTAGCGGGGCGGAAATGATCCGGTGCCACGTGGTGTTGGAGAGGGGGCAATCT
           ******** ********************************.******************

L-1        GGCGAGTGGGGACGAGGGTATGAGGTGGTGTGTGGCGCTAGTGATGTTGGGTGCGATCAT
L-2        GGCGAGTGGGGACGAGGGTATGAGGTGGTGTGTGGCGCTAGTGATGTTGGGTGCGATCAT
L-3        GGCGAGTGGGGACGAGGGTATGAGGTGGTGTGTGGCGCTAGTGATGTTGGGTGCGATCAT
L-4        GGCGAGTGGGGACGAGGGTATGAGGTGGTGTGTGGCGCTAGTGATGTTGGGTGCGATCAT
L-5        GGCGAGTGGGGACGAGGGTATGAGGTGGTGTGTGGCGCTAGTGATGTTGGGTGCGATCAT
L-6        GGCGAGTGGGGACGAGGGTATGAGGTGGTGTGTGGCGCTAGTGATGTTGGGTGCGATCAT
L-7        GGCGAGTGGGGACGAGGGTATGAGGTGGTGTGTGGCGCTAGTGATGTTGGGTGCGATCAT
L-8        GGCGAGTGGGGACGAGGGTATGAGGTGGTGTGTGGCGCTAGTGATGTTGGGTGCGATCAT
L-9        GGCGAGTGGGGACAAGGGTATGAGGTGGTGTGTGGCGCTAGTGATGTTGGGTGCGATCAT
L-10       GGCGAGTGGGGACGAGGGTATGAGGTGGTGTGTGGCGCTAGTGATGTTGGGTGCGATCAT
L-11       GGCGAGTGGGGACGAGGGTATGAGGTGGTGTGTGGCGCTAGTGATGTTGGGTGCGATCAT
L-12       GGCGAGTGGGGACGAGGGTATGAGGTGGTGTGTGGCGCTAGTGATGTTGGGTGCGATCAT
L-13       GGCGAGTGGGGACGAGGGTATGAGGTGGTGTGTGGCGCTAGTGATGTTGGGTGCGATCAT
L-14       GGCGAGTGGGGACGAGGGTATGAGGTGGTGTGTGGCGCTAGTGATGTTGGGTGCGATCAT
L-15       GGCGAGTGGGGACGAGGGTATGAGGTGGTGTGTGGCGCTAGTGATGTTGGGTGCGATCAT
L-16       GGCGAGTGGGGACGAGGGTATGAGGTGGTGTGTGGCGCTAGTGATGTTGGGTGCGATCAT
L-17       GGCGAGTGGGGACGAGGGTATGAGGTGGTGTGTGGCGCTAGTGATGTTGGGTGCGATCAT
L-18       GGCGAGTGGGGACGAGGGTATGAGGTGGTGTGTGGCGCTAGTGATGTTGGGTGCGATCAT
L-19       GGCGAGTGGGGACGAGGGTATGAGGTGGTGTGTGGCGCTAGTGATGTTGGGTGCGATCAT
L-20       GGCGAGTGGGGACGAGGGTATGAGGTGGTGTGTGGCGCTAGTGATGTTGGGTGCGATCAT
           *************.**********************************************

L-1        ACCAGCACTAATGCACCGGATCCCATCAGAACTCCGAAGTTAAGCGTG
L-2        ACCAGCACTAATGCACCGGATCCCATCAGAACTCCGAAGTTAAGCGTG
L-3        ACCAGCACTAATGCACCGGATCCCATCAGAACTCCGAAGTTAAGCGTG
L-4        ACCAGCACTAATGCACCGGATCCCATCAGAACTCCGAAGTTAAGCGTG
L-5        ACCAGCACTAATGCACCGGATCCCATCAGAACTCCGAAGTTAAGCGTG
L-6        ACCAGCACTAATGCACCGGATCCCATCAGAACTCCGAAGTTAAGCGTG
L-7        ACCAGCACTAATGCACCGGATCCCATCAGAACTCCGAAGTTAAGCGTG
L-8        ACCAGCACTAATGCACCGGATCCCATCAGAACTCCGAAGTTAAGCGTG
L-9        ACCAGCACTAATGCACCGGATCCCATCAGAACTCCGAAGTTAAGCGTG
L-10       ACCAGCACTAATGCACCGGATCCCATCAGAACTCCGAAGTTAAGCGTG
L-11       ACCAGCACTAATGCACCGGATCCCATCAGAACTCCGAAGTTAAGCGTG
L-12       ACCAGCACTAATGCACCGGATCCCATCAGAACTCCGAAGTTAAGCGTG
L-13       ACCAGCACTAATGCACCGGATCCCATCAGAACTCCGAAGTTAAGCGTG
L-14       ACCAGCACTAATGCACCGGATCCCATCAGAACTCCGAAGTTAAGCGTG
L-15       ACCAGCACTAATGCACCGGATCCCATCAGAACTCCGAAGTTAAGCGTG
L-16       ACCAGCACTAATGCACCGGATCCCATCAGAACTCCGAAGTTAAGCGTG
L-17       ACCAGCACTAATGCACCGGATCCCATCAGAACTCCGAAGTTAAGCGTG
L-18       ACCAGCACTAATGCACCGGATCCCATCAGAACTCCGAAGTTAAGCGTG
L-19       ACCAGCACTAATGCACCGGATCCCATCAGAACTCCGAAGTTAAGCGTG
L-20       ACCAGCACTAATGCACCGGATCCCATCAGAACTCCGAAGTTAAGCGTG
           ************************************************
```

**Supplementary Figure 3.** **Mapping the 5’-end border of the 40-copy array of 5S rDNA locus at *S.polyrhiza* ChrS6. a** Nucleotide alignment of PCR-amplified 5’-end border sequence of the 40-copy array (S-1-Up) with the sequence of representative 5S rDNA unit (S-1). Only an area of sequence homology is shown, partial sequence encoding 5S rRNA is highlighted by green. **b** Sequence of the PCR fragment representing array’s 5’- termini border. The partial IGS sequence is *underlined Italic*, and the part of sequence coding for 5S rRNA is highlighted by green.

**a**

```
S-1-UP     AAACCCAATAACTTTCTAACCAAAACTCTGAATTAATAATAAAACCACTCGTTTCGAAGC
S-1        --------------------------------------------------GTCTCGGGCG
                                                             **.***..

S-1-UP     TAAAGACCAGCGCTAGCTAACTATCTTTCGAGGGCGTTTCGGGCAGGTCTTGGGCGCCTG
S-1        CGGA----GGGGCTGGCTAGCTATCTTTCGAGGGCGTTTCGGGCAGGTCTTGGGCGCCTG
           ...*    .* ***.****.****************************************

S-1-UP     GGGCGGACTCCGGAGTCCACGGGGAGGGGGTTGTCCGCCCGCCTTGGGCACGAGCTCCCC
S-1        GGGCGGACTCCGGAGTCCACGGGGAGGGGGTTGTCCGCCCGCCTTGGGCACGAGCTCCCC
           ************************************************************

S-1-UP     CTGGCGAGAGGCGTCCCGCATCCGCGCCGCTCGAGGGACACCACGCCGCCCCTTGGCGGT
S-1        CTGGCGAGAGGCGTCCCGCATCCGCGCCGCTCGAGGGACACCACGCCGCCCCTTGGCGGT
           ************************************************************

S-1-UP     GGGTCAATGATCTCGCCCGCCCCCTCTCCCTCTCCGTCTGTCAGGGAGGCGGCGCCGATG
S-1        GGGTCAATGATCTCGCCCGCCCCCTCTCCCTCTCCGTCTGTCAGGGAGGCGGCGCCGATG
           ************************************************************

S-1-UP     CCGGGGGGGAGAGGATTAGGGCTGACGCGGCGCTTGTCACATCGGGTGCGATCATACCAG
S-1        CCGGGGGGGAGAGGATTAGGGCTGACGCGGCGCTTGTCACATCGGGTGCGATCATACCAG
           ************************************************************

S-1-UP     CACTAATGCACCGGATCCCATCAGAACTCCGAAGTTAAGCGTG
S-1        CACTAATGCACCGGATCCCATCAGAACTCCGAAGTTAAGCGTG
           *******************************************
```

**b**

**>S-1-UP**

**TCTTCTGCCTTGTACCTATTTCTAATTTTTATACAAAACACTAAGGAAAATTAAAGTCCAGGCCAAACAATAAAAAGGTAAAAGGTTTAATCAAAATCAAGTTCTGTATGATGTTTCTCAACAAACAAATAATCAATGCCTTAAACAACAAACAAGCTCTAGAATGTGAAAGTGACAATAAGAATAAGTTCTAATCATCCTCTTCATGCAATCTCTCAACATAACCATAGTCAACATCAAATAACACTTTAAAACATAAATTTGAGTAAATTCTGATTAACCTATTATTTTATCACCAAATGATGCTATCTTATAATCAATTGACTCACATGAGCCATGAACAAAATTTTCTTGAAATTTTTCATTAATGATCCATTATATAGTAAATGGTTAGTCCTCATTCAACCAACGAACAAGATTATTCTTGCAAAGTGGATATAGTATAACTATGTACACATTTCATCTAACAAAGAATCCCTTATTCTTCAACCAGAATAATTACTAACACTATGAAAAATGATAGAAACAATACTATCAATAGTGCACTAAAGTAAAATATTAGAGTCAACATAGTTCAACACAACCTAAATAAAAATGCTATCTAAAATCTAACCCAAAAAATTGACTGAACTAAAAGGTATTCCGCTAACCATAGATTTCTCACCACACAAGAAACATTTCCACAATCAACCACCTACAAACAAAAAATATCAAGATAGTAAAAACCCAATAACTTTCTAACCAAAACTCTGAATTAATAATAAAACCACTCGTTTCGAAGCTAAAGACCAGCG*CTAGCTAACTATCTTTCGAGGGCGTTTCGGGCAGGTCTTGGGCGCCTGGGGCGGACTCCGGAGTCCACGGGGAGGGGGTTGTCCGCCCGCCTTGGGCACGAGCTCCCCCTGGCGAGAGGCGTCCCGCATCCGCGCCGCTCGAGGGACACCACGCCGCCCCTTGGCGGTGGGTCAATGATCTCGCCCGCCCCCTCTCCCTCTCCGTCTGTCAGGGAGGCGGCGCCGATGCCGGGGGGGAGAGGATTAGGGCTGACGCGGCGCTTGTCACATC*GGGTGCGATCATACCAGCACTAATGCACCGGATCCCATCAGAACTCCGAAGTTAAGCGT**

**Supplementary Figure 4.** **Mapping the 3’-end border of the 40-copy array of 5S rDNA locus at *S.polyrhiza* ChrS6**. **a** Nucleotide alignment of PCR-amplified 3’-end border sequence of the 40-copy array (S-1-Down) with the sequence of representative 5S rDNA unit (S-20). Only an area of sequence homology is shown, partial sequence encoding 5S rRNA is highlighted by green. **b** The whole sequence of the PCR fragment representing array’s 3’- termini border. The 5S rDNA IGS sequence is *underlined Italic*, and the part of sequence coding for 5S rRNA is highlighted by green.

**a**

```
S-1-Down   CTTGGGCGAGAGTAGTACTAGGATGGGTGACCTCCTGGGAAGTCCTCGTGTTGCACCCCT
S-20       CTTGGGCGAGAGTAGTACTAGGATGGGTGACCTCCtggGAAGTCCTCGTGTTGCACCCCT
           ************************************************************

S-1-Down   TTTTCGCTGCCCGCGCGCACGTCTCTGCGGGTCCTTCCTCCCCCCTGCTCTGGTCTCTTG
S-20       TTTTCGCTGCCCGCGCGCACGTCTCTGCGGGTCCTTCCTCCCCCCTGCTCTGGTCTCTTG
           ************************************************************

S-1-Down   TTTTCCCCGCCGGCTTCGACCATCCGGGCGAGCTGCCCCCACGCGTGCAGGTCTCGGGCG
S-20       TTTTCCCCGCCGGCTTCGGCCATCCGGGCGAGCTGCCCCCACGCGTGCAGGTCTCGGGCG
           ******************.*****************************************

S-1-Down   CGGAGGGGCTGGCTAACTAAGTTCTATTAAAATAAATTGTTAATAATAAATTGAAATCAA
S-20       CGGAGGGGCTGGCTAGCTATCTTTCG----------------------------------
           ***************.***  **...
```

**b**

**>**S-1-Down

CTTGGGCGAGAGTAGTACTAGGATGGGTGACCTCCTGGGAAGTCCTCGTGTTGCACCCC*TTTTTCGCTGCCCGCGCGCACGTCTCTGCGGGTCCTTCCTCCCCCCTGCTCTGGTCTCTTGTTTTCCCCGCCGGCTTCGACCATCCGGGCGAGCTGCCCCCACGCGTGCAGGTCTCGGGCGCGGAGGGGCTGGCTAACTA*AGTTCTATTAAAATAAATTGTTAATAATAAATTGAAATCAAAAGTCAGCATATTAGTAGGCTATTTTCAAGTTGCCATTTTGAAGCTTTCTGAAAATTTTGTAAATGTAAAAATGTCTTTAAGGTACCTATAGAAATATCTTTGGCACCATAACTAATAGTCAAATTTTCAAATTATTATTTCACAGATAAACAATAATCATATTTGAGAAAAAAATAATACAATGTAGACTAGAAAGTGAAAATTCAGAGTTTCAGGATTTTCACATTTTATAAAGCTAATTTTTTTGCTTAAGATTTGTCCCTTGTTTTATCTCACTGCCTTAGTGGTGGATATTTTTCATTATTGTTAAATACTCTCTATCTAATAATATTTATAACAAAAATAATTTTAAATTATCCCATGTCATTGCAACCCAAAGGGACAATAAGCGTAGGCTTGGGTCATTTGGAAACCACATGAACTGGAGTTGGCATGACTCTGATGCACATCACCATGGACAAAAAATTGTGCCCTCTCATACAATCTTTTTCATTTATTTTAATTGTCACTAATTGGTTATGTTGGTTGATGGCCCCTTCATGTAACGTGATGATGTAGAACAAAATGGAAATACAAACTCATTATGGTGCCCAATTCTAAGTCTTATTTCTCCAACAGAGATTAATAATTTTCAAGAGTAAGGCTAGTTTCAGTCCAATTTTGATGTGCAATTTTGAATAACCAAAACAGTTAAGAAATCATCTGGTTCAAGGTGCTCAACAAACAAGGAATCAATTAATAGATTTCCATGGAAATCTCTCAATTAATCAAGCTGGGGAAAATGTAAATACTGTCTTTGAATGGTTTTCCAATAGCACCATCAAGAATATATCCTAAGATCAGGTTTTGGTGATTTGGTGGATATGATTGAGTGGAAAGTGATGTTATTAACATGTGGGT

**Supplementary Figure 5.** **Consensus nucleotide sequence of the *S. polyrhiza* 5S rDNA locus at ChrSp6.** The sequence is composed of 10kb long upstream chromosomal region followed by a cluster of 40 copies of 5S rRNA genes, followed by 10kb long locus’s downstream chromosome region. The clusters of 5S rDNA repeats are marked by *underlined Italic*; sequences coding for 5S rRNA are highlighted by green; GAGA stretches are highlighted by yellow; DNA stretches longer than 10bp with 90% or more of T/A are in **bold**; locations of primers used for amplification border sequences of the rDNA cluster are highlighted by grey.

**>ChrS6-5SrDNA** **(40,878 bp)**

TCTGTCTGTCTGTCTGTCTGTCTATCTATCTGTTTGTCTGTCTGTGTCGTCTGTCTGTCTGTCTGTCTGTCTGTCTGTCTGTCTGTCTGTCTGTTCTATCTATCTATCTATCTATCTATCTATCTATCTATCTATCTATCTATCTATTTATCTATCTCTCTAAGATTGATTCTGCAACGTACACTATAAAAGAGCATGCCTCATGCTACTTGTTGGATAGTGAGCTGATTAGGTGAGTTGGCTCCAATTAGTCAACCTAAAGTGTGTTGACCCATTTACTTCAACTTAGAGAAAATCAC**TAAAAAATTAATAGAAAT**GTG**AAATTTTTTAAAGAATTAA**CTTGTTCCTCTCTAGCTCCCTATTTAGTCAAAACCAGGATC**AATAATTCAAAT**CAGAGCTTACCTTAGAAGTATTATACTCACATACG**AATTTAATTGAATAAATTCATAAATAATTTAAATTAAAAATTAAAAATATTCAACTA**GGAATTTAGATGATGCATTTGACTTTCAGCATGCTGCAATATGTATG**TTATTGTATTTTA**CTACTTGAAGTCATTAC**TTTACATATTATT**GTGAAAACTATTCTTATTC**ATATAAACTATAATT**GTC**AAAAAAATAAGAATTT**GGACCTCTTGATC**AATGAATTTTTTT**G**AAGTAAAATAT**GGGTGTGATAACTTATCAAC**ATATTAGATAT**CTATG**ATCATAAAATATTTATTAGATTTTA**GAGATTAATGCTGTTGGACTTTTTTGAGGTCCAGCAACTTAATTCTAG**TTATTATTTTCTGAAATTATTTTT**GCATAACATAGG**TAAATTTAAT**GGAGTAAATGGATGATTTTGGAGCTAAAGGGGCTGGAGAGCTGGCCCAATGGAAGTCCAAGGACATGGGCACATGTCCAAGGGATAATTGGCCTGAAATGGCCAAATCAAGTGCACATCTTTCAAGACTCAAGGCCCAAGGGTGTGGAGCAATTAATGGTGGGAACTTTGACTTGGTGCCAACACGTGGAGCAAGAGCGTGTGTACATGCCCCCTGAAAAGTTGTTGTGTAGGAGAG**ATAAATGAAA**GTAAATGTGAATGTTTGGGGGGATAAAAAGGGAATCACCAAGG**ACTATTTAAT**GTAATGTTCTCCCTCTTGTTGGGGGTAATTCACAAGGTTTCAGACAAGGGCTAGCTACTGGAAAAGACCTAAGAAATCCCTCTTTTGATTAGATGG**TTTAAATGAA**G**TTTATTTCAA**CTACTCTAGTGTGATTAAGAGAGGCAACAAGGAATGGAAGAAGGAGATTAGCATAGAAGAGGAGGCACACCCATACGTAGGTGATTGGAGCCCTAGGTTGGTGAGGTAGACCCCAAGGTGAGTGTTCAATTCTTGTGCATATATTCCCTTCCCCTAATTCTGAAATCAGGTAGGTGTAGTAGTAGC**ATTTAATACTA**GATTAAAACCCACACTCACCTCTTAAAGTCACCCATTTCCACCCCAAATATCTCACTCTTGGGTGTGAGATATCTCTAGTCCTAGATCCACATCCTCTCCCTCTTTCACCTC**TTATTTATTACAT**GTCTCTTCTCCCCATTTAATCCTAAGATCAACTATTAACC**TAAATTTAAGA**CCATTCCTAGAGGAGACAATCCTAGCTC**ATTATTATGTA**CTTAATTCATCAAGAATTAGGC**TTATAAATTATTTTTGATTAAGTT**CTAGTTAGGATCATGACACCC**TAACTAAAAA**CTTATCAATGGGGTCCTCATGACCAACCAAACTAGGGTTAGGATGATCACAAAGATCAAGTAGACTTCACTCCAACAAATGCCAATTGTTCTTACAAG**ATGATAATATATCATTTTTAAAAT**GCAATGTGAGG**TAAAAGTAATATA**GTGTCG**TATGAATTTT**GTG**AAAATAATTTATAAATAGTGAAAAATAT**G**TTTGTTAATTTTTTTATTATTTTT**GG**TTAATTTTAATTATTTTCATAACTTATTT**CAAATGAAATGTGTC**ATTTATTATTTTATGATAT**C**AATAAATATAAATAAATAATAA**CAAAGGTGGTATTAGGCTCACCACATTTCGTGGGCCCAATGCAAGCAATCATGGGTTCACCATAGGCCAGTAATTGCATTAGACAATGTGCATTAAGAGGTCTACCGAGTTGTGATGCAAAAGACTCCACCATGCAAGCCCATGACATCATGGACCAAGGTCCACCAGGTCTACTGGAGGGCAACTATGAAGTCTCAATTTCCCTACCCATTATTAGTCCAATACAAGGCCTCTCTTAGGCCTAATAGGTGTTTGAGCCTAACCTCAAGTCAACTTTCAGGAGCTAATGATGAAATCTGGTCAGCCCATTTAGGAGAGGAACTTGATGAGAGAAGGTATTCC**AATTTATTAG**CATG**ATAATTCTAT**CAAAATAACCAATCATGGCCTCTAACAACTGGTGAACCTCTTCCTATCTCCCTTGGCTGCACTCAAAGCCTCCAACAAGTGGTGAAGTCATTCCTTCTTCCTC**ATAACAATAA**CATCAACCCTTCTAGCAAAATCTTTCTAGAGAGAAACATAAAGTCGCCATATGACTCTTATGTGCCTTTTGGTAGGGTTTAACATGACTAGATGATACAAGGTCACCACACCAGCATCTCTAACTACTCCAATGATGGTCTCATCAATGACAATGTTC**ATTTTCATTTAT**C**TTAGTTTTAT**GTTC**ATTTTTAGTT**GCATACAATGATTTC**TAGATTTTTT**CAG**AAATAATCAATA**GAACTCATGATTC**AATCAAAAAAT**GGTCTTCTTTTCTCTCTTAGATCCTTGATAACCCATTACAGATTGTCTTAAGAATC**ATTTTTAATCATTTTT**CATTAATGTCC**ATAAAAAAAT**GAGCTCAAAATCC**TTGTAATTTTT**CCATGAACACC**AATAAAATTTCT**AGGATATTTTCC**TAATTTTTAAGTTATT**CACCTAGTTTTCTAGTTCCATTTCAATG**TTTAATTCTTATTAATATATATAAAATA**G**ATTTAGTTATTTTAT**G**TATAATCTTA**CTCCTTTTTCGTAAATTCTCTCTCACGTATTCACTGGAAATCATGGTCAATATAGTGCGAGACATAGTGAGCAACAATTTTCTCTCTTTTATCATCTAAATTCTCTCTC**TAATTATCTT**CCTAAATTACCTC**TAAAAAATCATTT**GAAACGCAACATGTCTCCCTAATTTCTTCCCTCTAAATCCCTATAAACCTCACATCTATCTCTCTCCTACCTCTTTATCCTATTCTCTCCCCTCTTTTTTCCCTTGTATTCCCTTGAAATCTCATTTCTACCCTC**TTAATTTTCT**CCTTTGAATTTCTCCC**TTTTACTATT**CTTTCACTTG**AAAATAAGAAAAAAAT**CATAACTTCTAATG**ATTAATATTTTTAGTTTATTTAATTTATAAAAAATCATAAATTATTTTAAAA**C**TAAATCAAAATTTAT**CTTCATGATAG**ATTATTTTAA**CTCTATGTTCAACTAGAGACTCATTTC**TATACTAAATATTATTTAA**G**TTTTAATTAAAATTATTATTAGATA**G**TAGTTATTTAA**CACCAATGACCAATACCCATCACAAGATAGG**TATTTATAGA**CCACATGATATAAGGGAAGGAGTTCAAAGAGGCACATCTAATCAAATGTTGTGAACACCCCAAACAATGGTCATGGGGGTGAGTTGATTGACACAACTATTAGCTTGTAGGTAGTAGTTCCTTTAGTGGTTGTTGTCTTGTTTGGTGTTTGATAACTTGTGAAACTAACATACTAATC**TATTATCAAATTT**GGGAG**AATAATTCATATTT**GATCAAG**ATTTCAAAATTTAT**C**ATATTAACATAATT**CACG**TATTATTTAAATGAATTTAT**C**TAAATTATGAT**G**TAAAATAATATTTTATGATTAAAAATAAAAAAGTTAAT**GAGAGTC**ATTTATTATGAAATAAT**GTGATTGATGC**ATAAATCTTATTTATTT**CATGTTG**ATTTAATTTTTATTATGAT**GTGCTC**AATTTTTATAATTTTTAATGAATTATTTCTATT**G**AAACATATTTT**CTAC**ATTTTTTATTTTTTAATGAAATAA**G**AATAAAAAAATAGATTAAT**GGCTCACAAGTTAGTGGCCCTATGCTGGCCCAAGACCCATACACCTAATAGTTGATTCTCATGTCACATATCTAACAAATAGAAGAGTAACATGTCATTGAGTCCATAGACGTTACCCATCACTCAAATCTTGTGCTTGTAAAAGTAGAGCAATTTGACGTACCAAGAGGTGACACACTACTTGCCTCATCATGCCTACCAGTGGATGCATAGATCCTAGTAAAGGTAGGAGATAAGGTGATGGCTATCTGATGACAACTATGCAGCCACATCACTCTTCCTTTTTGTGGATTGGTGGTGTGTGGATTTGACACTCAAAGTTGAACCACATCACCAACCAACATCTCACGTGCTTTGATAAAGTGGACTCAAGCAAAAAGATTGTGACGGCCTCATATGACGACATATAAGTGGCCACATCACCAATGAATCATCATGGATTGGCAATGTAGGAATCATACGGTTCTCAACCTACCACATGACAAGGAAACACATGCAAATATCGATTATCATATGATAAACTAGGTATATCATGATACACATGCAGCTGGGCATATTTCCAAATATAGATTATCATC**TAGTAAAATA**GACAATATCCATTATTCTCACACACAATGG**TAATAAAAATTTAGAAAAAAAAT**CG**ATAAAAATATAAAAAAATTAAATAAAATT**G**AAAAAAATAG**CAAATCAAAACATAAAGCTTGATTTGCAGATCTAG**AGAATAAAAA**GGATGGAAGAGGAAGAGATGGAGGTCTTTTTTGC**AATTATTTTTTATGTT**CTAGGGTTAGAGATAGTTTTTCTTCTTG**TTTTTCTTTTATTTT**CAGCAACTTGATGTATGAGAGGATTTTTCTCTTTTCCTCAACATCAACTTAG**TAATTTTTTAGT**CAAACC**TAAGAAATTA**GATGTCATGAAGAGGC**TATAAAAATTATGTAAA**CCTTAGAGTTATCGG**TATTAAATCTAT**GAG**ATATAAAGAAATTTAT**GATCAAATGAATGATC**AATAAAATCTAAA**GAGTTAACCCAAAGTC**AAAATTAAGATAA**GATCCACC**ATAATTTGTA**GAGAACCAAATCACTACAAAATTCTTGATGTGCCCATTATCAAGAATTTTGTCATCTAGTGACATAGAGAATTAGAACAAC**TAAAAAAAACATAA**CATAGTCAATGAATAGAC**AAAATTAAAATCAAATA**GGTTGGTAGCAAC**TAAAAGAATA**G**ATAGAATTATAAAA**CC**TAATTTTACATA**GATGACTCTTTCTTGGGGAAGAGAATATGTTGAGTACATTAGTTTTAG**TTATAAACTTTATAA**GAATG**TTTATAGTTT**AGAGTGAGTAATGTTGTTTGATACATTAGAACAAGGCTAGATCTAATACCCAATAATCGATGTCATATGAAC**AATAAATGTAAAA**CATAAATGAGG**AAATAAGAATAAT**CAATGTTTGATATGTTGGAACATAAATG**ATGAAAAATAA**GCATAAGATGCATGAAAAGATC**AAAATATGTTT**GTTTATTCTG**AAATTAAGTTTTTATTTAAT**GTTC**TTCATATATAA**GGATGAAGTTGGG**TTTTAGATTA**GTGGGGATCAAGAAGAGGAGTTGAATTGAACTCCTTTTTCTCTCTTCTAAAGACTTCTTATTCGTGAGCTCCTTGCTAGTCTCTTAATGGGCTGATCTTTATTGGGCTGTGATCATGAACACATGGTAAACTCAATTTTAGC**ATTTGTTTATA**G**TCATTATTTT**GCTATGAG**TTTATACAAAATATTTATAATTTTTCTT**CCTGATTTGCTCTAGTTTTAGAATAGGGCAAGTAGGGGGTTAATCATCAAGGAATTAGGATTG**AAATTTTGATT**CTTTTAAGG**TTATAAAATAAGAAAAAAGATTA**GAGAGAAGAATGAGGGGAAGATGATGGAGGAATGAAAATCTTGATTTTCAAGATAGTTATTCCAAGAAAATGAAG**ATTCTAATTT**C**AAAATTTAAGAATTTTTTTATAA**CTACAATTAGGACTTAGGGAAATGAGTTGAGAGAAGAAAAGATGATGAAAGAGGGGAGGATCAAGGAGATCTTGATTTTCCTATGAAAATCAAGAAATCCAAGATCTCAAAGGAAAAGGGTGAATTAGGGTTTCGATTTTGAGAAGGAGAAAGGATGAGAGAAC**TAATTTTTCT**A**TTGAATATTA**GAGGATGAAATC**ATGAATTTTA**GGACCTAGAAGGGCATACCTTGGGTC**TGTATATTTT**GCCCAGACCAAATGGTTTTCTCCAACAAGAACACACTTCAAAGATCAAGAACAAATATTGCAATAATTCACCTCTATTCAG**ATTTATCAAT**CTGAATCCTAGGAAAGCTTTTTCTTGTGCCCAGCCTCTTCTAAGGTAATCTTTAGAATGAGG**AAAAATAAGAAAA**GTAAATAGG**TTTTATATATA**GGGAC**ATATTTCTAAA**GAGAACCGTCTATTTTCTCATTTATCTTCTCATGCCCTTTTATCC**TTCTTTATTTT**GATTGGGCCCCC**TTTAAAACTT**CAATTGTGTGGCCCATCTTAGAGAGGATGGTTGAAGATAGATTGGGCTGAAAATGGTTGTGCTCCCCTCTCCTTCTTTGGGCATTTCTTGGACCAAAGCTCAATCATCTCTTCTTCCAGCCCATTATAGAAGTCAAGCAAGGTTAGGC**ATAAAAAAATAAATCAA**GGAATGTTCGCATGTGCATGAAACTAGAGCAAGATATTTCAAGGTTAATCCTCACAAAATGGTAC**ACATTTTTTTAT**C**ATCAATAATTT**CATAGTAATCATGAATCATGTGTATATCATTG**ATTTATATAAGTA**GAAGTAAATATGAAGTAATTGCTC**ATATTCATTATTTTAATATT**GCTAGCATAATGTTGGAAAACTTCAAGGCTCAAAGCATTAGAATCATAGATATTGTCAC**TATATTTATCAT**GCTATCAATAAGTTATCAACAC**ACTATTTTATA**GTGGTTTGGCACTGACATACATTCATGCCCTTGGGTTCTACTAACTGATCATACATATCAC**AATCATTTATTT**CGTAGGCTCAACTCCCCCATGATTTCACTTATAAATC**TTCTTTTATTTT**CACGGCTTAAGGTGGACTTACACCAC**TTTTTTTTACAAAAT**GACTCCCCAAGTGATC**AAATAAATAATCAATA**CTGATAACTC**ACAAATATAAAACA**CTAACATTGTGAGTATATTCGGAAGATAAGCTTTGAATGTTCTAGATTGGTGTAGTGATAGAAGATGAATACTTTGGATATTAATTCTTTGTCTTATTGTTGTGATAGTTAGTTGGTATAGGGCTTAAAGGCCTATCGTTAGGCACAAATCGAGGGTGGAAAATATCTTGAATAGTTTCCTAGAGATTGATCATTCTAC**AATTAATGAATA**ACGAATTGATCAATCTTCTCTCTCTCCTCCTCTTGAATGCACTATGTGAAGACCAC**TAAAATAGAA**G**AAGAATAAATAT**AGTCATTGATGATTGATTCTTTTCATTGAAG**ATTAGTAATATTTAT**GGATG**AATTAAGTTAAA**GCTATTCATAGATTTTTCAAGATCTCTAATAACTAGATTCAC**TAAAAATAGAATTAAAGTTAAATTTTACATAAATTTAA**CACGACTTGTCATTTAACTACTTCAGATATCTTCTAAGAAATCAACACGTGAATC**TTTTTAAAACTTAA**CATTC**AAATCTAATA**GCAATCGATTC**AAATAAATCTAATA**GAAATCC**AATAAGATTTTTT**GTGAATCTAACATGACATTTCTTGCGAAAGTAATG**ATTTATCATA**CAAATC**ACATTTAAATT**CAG**TAAAAAAAGATATGA**CATAAGTATACTTTGTGAAATTCAATCTC**AATATTTTAAATCATAATATTTTT**GTCAAAATTGATCTGTATGATTTTC**AACATATATTT**GGTTAATTTGACACGACTTAGAATTTGGCTGATTCGATGTTTAAATC**ATCAAATTATTATAAAATAAATT**CATGGG**TATATAAATA**CCTAAATAAGCTTTCATGC**TCATATTTTA**C**TTTATCTAAA**CTCATTGATTTGTTTGATCAAATCATTACTC**TTTTAACAATAT**G**ATATATTTCT**CAAGATTTTATGACTCAACTACTTAG**ATTCATATAAT**GAGATTTTCATTC**ATAAATTAGTTTAT**CAATTGTATGTTTGAGATCATGACATTCATGCTTCATGTGTATGATTTCACATGTTACATCATAG**TAGATATTATATGTTTATATATAAT**GC**ATTTATCATTAAA**CCATGAATG**TATATTGTTAAA**GG**ATTATTCTTAATT**GAACTTTG**TGATTTTGTTAT**C**ATCAAAATATTAT**GTCTTCATAGCTTCAAC**AAAAGTTAATAA**CATC**TAAATTTTAGAAAAT**CTCAAATAC**TAATATCAAATAT**C**ATAATAGTATTTTAATGTATTTTAATAT**CCTAATAGTAAATCTTGATG**TATTATTGAATATT**GAG**ACTTAATATTA**GTG**AATTTAAATTATTTACTT**GG**TATATTATTTG**GATAGTATC**AATAATTACATAAATTA**CCACAATG**AATTTTACATTAT**GCATGTCAGG**ATAATAAAATTTATTTAAAAATTATGATTTA**CTTGAGTTACTTTTGGTAGTTTTGATTGTG**AAATCATTTTA**CTTCAAGATGCATCATCAACTTTTGTTCTG**AAAATAAAAAATTCTAATT**GAACTTGCTTTAGC**TAATAATCTTTAAAT**GCATGCACTAATTGTAC**TTACTTAAAT**GCATGCTTTGCATATCTG**AAATCTTAAA**GGTCATTTC**ATTTTTCTAATTTTTTTGAAATTTTTAAGTAATTTT**CCAATGGCTAATTAGTTCAC**AATAATCAAATAA**GACATTTCTTCTGCCTTGTACCTATTTC**TAATTTTTATACAAAA**CACTAAGGAAAATTAAAGTCCAGGCC**AAACAATAAAAA**GGTAAAAGG**TTTAATCAAAAT**CAAGTTCTGTATGATGTTTCTCAAC**AAACAAATAAT**CAATGCCTTAAACAACAAACAAGCTCTAGAATGTGAAAGTGAC**AATAAGAATAA**GTTCTAATCATCCTCTTCATGCAATCTCTCAACATAACCATAGTCAACATCAAATAACAC**TTTAAAACATAAATTT**GAGTAAATTCTGATTAACC**TATTATTTTAT**CACCAAATGATGCTATC**TTATAATCAATT**GACTCACATGAGCCATG**AACAAAATTTT**CTTG**AAATTTTTCATTAAT**GATCC**ATTATATAGTAAAT**GGTTAGTCCTCATTCAACCAACGAACAAGATTATTCTTGCAAAGTGG**ATATAGTATAA**CTATGTACACATTTCATC**TAACAAAGAAT**CCCTTATTCTTCAACC**AGAATAATTA**CTAACAC**TATGAAAAAT**GATAGAAACAATACTATCAATAGTGCAC**TAAAGTAAAATATTA**GAGTCAACATAGTTCAACACAACC**TAAATAAAAAT**GCTATCTAAAATCTAACCC**AAAAAATTGA**CTGAACTAAAAGGTATTCCGCTAACCATAGATTTCTCACCACACAAGAAACATTTCCACAATCAACCACCTAC**AAACAAAAAATAT**CAAG**ATAGTAAAAA**CCCAATAACTTTCTAACCAAAACTCTG**AATTAATAATAAAA**CCACTCGTTTCGAAGCTAAAGACCAGCG*CTAGCTAACTATCTTTCGAGGGCGTTTCGGGCAGGTCTTGGGCGCCTGGGGCGGACTCCGGAGTCCACGGGGAGGGGGTTGTCCGCCCGCCTTGGGCACGAGCTCCCCCTGGCGAGAGGCGTCCCGCATCCGCGCCGCTCGAGGGACACCACGCCGCCCCTTGGCGGTGGGTCAATGATCTCGCCCGCCCCCTCTCCCTCTCCGTCTGTCAGGGAGGCGGCGCCGATGCCGGGGGGGAGAGGATTAGGGCTGACGCGGCGCTTGTCACATCGGGTGCGATCATACCAGCACTAATGCACCGGATCCCATCAGAACTCCGAAGTTAAGCGTGCTTGGGCGAGAGTAGTACTAGGATGGGTGACCTCCTGGGAAGTCCTCGTGTTGCACCCCTTTTTCGCTGCCCGCGCGCACGTCTCTGCGGGTCCTTCCTCCCCCCTGCTCTGGTCTCTTGTTTTCCCCGCCGGCTTCGGCCATCCGGGCGAGCTGCCCCCACGCGTGCAGGTCTCGGGCGCGGAGGGGCTGGCTAGCTATCTTTCGAGGGCGTTTCGGGCAGGTCTTGGGCGCCTGGGGCGGACTCCGGAGTCCACGGGGAGGGGGTTGTCCGCCCGCCTTGGGCACGAGCTCCCCCTGGCGAGAGGCGTCCCGCATCCGCGCCGCTCGAGGGACACCACGCCGCCCCTTGGCGGTGGGTCAATGATCTCGCCCGCCCCCTCTCCCTCTCCGTCTGTCAGGGAGGCGGCGCCGATGCCGGGGGGGAGAGGATTAGGGCTGACGCGGCGCTTGTCACATCGGGTGCGATCATACCAGCACTAATGCACCGGATCCCATCAGAACTCCGAAGTTAAGCGTGCTTGGGCGAGAGTAGTACTAGGATGGGTGACCTCCTGGGAAGTCCTCGTGTTGCACCCCTTTTTCGCTGCCCGCGCGCACGTCTCTGCGGGTCCTTCCTCCCCCCTGCTCTGGTCTCTTGTTTTCCCCGCCGGCTTCGGCCATCCGGGCGAGCTGCCCCCACGCGTGCAGGTCTCGGGCGCGGAGGGGCTGGCTAGCTATCTTTCGAGGGCGTTTCGGGCAGGTCTTGGGCGCCTGGGGCGGACTCCGGAGTCCACGGGGAGGGGGTTGTCCGCCCGCCTTGGGCACGAGCTCCCCCTGGCGAGAGGCGTCCCGCATCCGCGCCGCTCGAGGGACACCACGCCGCCCCTTGGCGGTGGGTCAATGATCTCGCCCGACCCCTCTCCCTCTCCGTCTGTCAGGGAGGCGGCGCCGATGCCGGGGGGGAGAGGATTAGGGCTGACGCGGCGCTTGTCACATTGGGTGCGATCACACCAGCACTAATGCACCGGATCCCATCAGAACTCCGAAGTTAAGCGTGCTTGGGCGAGAGTAGTACTAGGATGGGTGACCTCCTGGGAAGTCCTCGTGTTGCACCCCTTTTTCGCTGCCCGCGCGCACGTCTCTGCGGGTCCTTCCTCCCCCCTGCTCTGGTCTCTTGTTTTCCCCGCCGGCTTCGGCCATCCGGGCGAGCTGCCCCCACGCGTGCAGGTCTCGGGCGCGGAGGGGCTGGCTAGCTATCTTTCGAGGGCGTTTCGGGCAGGTCTTGGGCGCCTGGGGCGGACTCCGGAGTCCACGGGGAGGGGGTTGTCCGCCCGCCTTGGGCACGAGCTCCCCCTGGCGAGAGGCGTCCCGCATTCGCGCCGCTCGAGGGACACCACGCCGCCCCTTGGCGGTGGGTCAATGATCTCGCCCGCCCCCTCTCCCTCTCCGTCTGTCAGGGAGGCGGCGCCGATGCCGGGGGGGAGAGGATTAGGGCTGACGCGGCGCTTGTCACATCGGGTGCGATCATACCAGCACTAATGCACCGGATCCCATCAGAACTCCGAAGTTAAGCGTGCTTGGGCGAGAGTAGTACTAGGATGGGTGACCTCCTGGGAAGTCCTCGTGTTGCACCCCTTTTTCGCTGCCCGCGCGCACGTCTCTGCGGGTCCTTCCTCCCCCCTGCTCTGGTCTCTTGTTTTCCCCGCCGGCTTCGACCATCCGGGCGAGCTGCCCCCACGCGTGCAGGTCTCGGGCGCGGAGGGGCTGGCTAGCTATCTTTCGAGGGCGTTTCGGGCAGGTCTTGGGCGCCTGGGGCGGACTCCGGAGTTCACGGGGAGGGGGTTGTCCGCCCGCCTTGGGCACGAGCTCCCCCTGGCGAGAGGCGTCCCGCATCCGCGCCGCTCGAGGGACACCACGCCGCCCCTTGGCGGTGGGTCAATGATCTCGCCCGCCCCCTCTCCCTCTCCGTCTGTCAGGGAGGCGGCGCCGATGCCGGGGGGGAGAGGATTAGGGCTGACGCGGCGCTTGTCACATCGGGTGCGATCATACCAGCACCAATGCACCGGATCCCATCAGAACTCCGAAGTTAAGCGTGCTTGGGCGAGAGTAGTACTAGGATGGGTGACCTCCTGGGAAGTCCTCGTGTTGCACCCCTTTTTCGCTGCCCGCGCGCACGTCTCTGCGGGTCCCTCCTCCCCCCTGCTCTGGTCTCTTGTTTTCCCCGCCGGCTTCGACCATCCGGGCGAGCTGCCCCCACGCGTGCAGGTCTCGGGCGCGGAGGGGCTGGCTAGCTATCTTTCGAGGGCGTTTCGGGCAGGTCTTGGGCGCCTGGGGCGGACTCCGGAGTCCACGGGGAGGGGGTTGTCCGCCCGCCTTGGGCACGAGCTCCCCCTGGCGAGAGGCGTCCCGCATCCGCGCCGCTCGAGGGACACCACGCCGCCCCTTGGCGGTGGGTCAATGATCTCGCCCGCCCCCTCTCCCTCTCCGTCTGTCAGGGAGGCGGCGCCGATGCCGGGGGGGAGAGGATTAGGGCTGACGCGGCGCTTGTCACTTCGGGTGCGATCATACCAGCACTAATGCACCGGATCCCATCAGAACTCCGAAGTTAAGCGTGCTTGGGCGAGAGTAGTACTAGGATGGGTGACCTCCTGGGAAGTCCTCGTGTTGCACCCCTTTTTCGCTGCCCGCGCGCACGTCTCTGCGGGTCCTTCCTCCCCCCTGCTCTGGTCTCTTGTTTTCCCCGCCGGCTTCGGCCATCCGGGCGAGCTGCCCCCACGCGTGCAGGTCTCGGGCGCGGAGGGGCTGGCTAGCTATCTTTCGAGGGCGTTTCGGGCAGGTCTTGGGCGCCTGGGGCGGACTCCGGAGTCCACGGGGAGGGGGTTGTCCGCCCGCCTTGGGCACGAGCTCCCCCTGGCGAGAGGCGTCCCGCATCCGCGCCGCTCGAGGGACACTACGCCGCCCCTTGGCGGTGGGTCAATGATCTCGCCCGCCCCCTCTCCCTCTCCGTCTGTCAGGGAGGCGGCGCCGATGCCGGGGGGGAGAGGATTAGGGCTGACGCGGCGCTTGTCACATCGGGTGCGATCATACCAGCACCAATGCACCGGATCCCATCAGAACTCCGAAGTTAAGCGTGCTTGGGCGAGAGTAGTACTAGGATGGGTGACCTCCTGGGAAGTCCTCGTGTTGCACCCCTTTTTCGCTGCCCGCGCGCACGTCTCTGCGGGTCCTTCCTCCCCCCTGCTCTGGTCTCTTGTTTTCCCCGCCGGCTTCGGCCATCCGGGCGAGCTGCCCCCACGCGTGCAGGTCTCGGGCGCGGAGGGGCTGGCTAGCTATCTTTCGAGGGCGTTTCGGGCAGGTCTTGGGCGCCTGGGGCGGACTCCGGAGTCCACGGGGAGGGGGTTGTCCGCCCGCCTTGGGCACGAGCTCCCCCTGGCGAGAGGCGTCCCGCATCCGCGCCGCTCAAGGGACACCACGCCGCCCCTTGGCGGTGGGTCAATGATCTCGCCCGCCCCCTCTCCCTCTCCGTCTGTCAGGGAGGCGGCGCCGATGCCGGGGGGGAGAGGATTAGGGCTGACGCGGCGCTTGTCACATCGGGTGCGATCATACCAGCACTAATGCACCGGATCCCATCAGAACTCCGAAGTTAAGCGTGCTTGGGCGAGAGTAGTACTAGGATGGGTGACCTCCTGGGAAGTCCTCGTGTTGCACCCCTTTTTCGCTGCCCGCGCGCACGTCTCTGCGGGTCCTTCCTCCCCCCTGCTCTGGTCTCTTGTTTTCCCCGCCGGCTTCGGCCATCCGGGCGAGCTGCCCCCACGCGTGCAGGTCTCGGGCGCGGAGGGGCTGGCTAGCTATCTTTCGAGGGCGTTTCGGGCAGGTCTTGGGCGCCTGGGGCGGACTCCGGAGTCCACGGGGAGGGGGTTGTCCGCCCGCCTTGGGCACGAGCTCCCCCTGGCGAGAGGCGTCCCGCATCCGCGCCGCTCGAGGGACACCACGCCGCCCCTTGGCGGTGGGTCAATGATCTCGCCCGCCCCCTCTCCCTCTCCGTCTGTCAGGGGGGCGGCGCCGATGCCGGGGGGGAGAGGATTAGGGCTGACGCGGCGCTTGTCACATCGGGTGCGATCATACCAGCACTAATGCACCGGATCCCATCAGAACTCCGAAGTTAAGCGTGCTTGGGCGAGAGTAGTACTAGGATGGGTGACCTCCTGGGAAGTCCTCGTGTTGCACCCCTTTTTCGCTGCCCGCGCGCACGTCTCTGCGGGTCCTTCCTCCCCCCTGCTCTGGTCTCTTGTTTTCCCCGCCGGCTTCGGCCATCCGGGCGAGCTGCCCCCGCGCGTGCAGGTCTCGGGCGCGGAGGGGCTGGCTAGCTATCTTTCGAGGGCGTTTCGGGCAGGTCTTGGGCGCCTGGGGCGGACTCCGGAGTCCACGGGGAGGGGGTTGTCCGCCCGCCTTGGGCACGAGCTCCCCCTGGCGAGAGGCGTCCCGCATCCGCGCCGCTCGAGGGACACCACACCGCCCCTTGGCGGTGGGTCAATGATCTCGCCCGCCCCCTCTCCCTCTCCGTCTGTCAGGGAGGCGGCGCCGATGCCGGGGGGGAGAGGATTAGGGCTGACGCGGCGCTTGTCACATCGGGTGCGATCATACCAGCACTAATGCACCGGATCCCATCAGAACTCCGAAGTTAAGCGTGCTTGGGCGAGAGTAGTACTAGGATGGGTGACCTCCTGGGAAGTCCTCGTGTTGCACCCCTTTTTCGCTGCCCGCGCGCACGTCTCTGCGGGTCCTTCCTCCCCCCTGCTCTGGTCTCTTGTTTTCCCCGCCGGCTTCGGCCATCCGGGCGAGCTGCCCCCACGCGTGCAGGTCTCGGGCGCGGAGGGGCTGGCTAGCTATCTTTCGAGGGCGTTTCGGGCAGGTCTTGGGCGCCTGGGGCGGACTCCGGAGTCCACGGGGAGGGGGTTGTCCGCCCGCCTTGGGCACGAGCTCCCCCTGGCGAGAGGCGTCCCGCATCCGCGCCGCTCGAGGGACACCACGCCGCCCCTTGGCGGTGGGTCAATGATCTCGCCCGCCCCCTCTCCCTCTCCGTCTGTCAGGGAGGCGGCGCCGATGCCGGGGGGGAGAGGATTAGGGCTGACGCGGCGCTTGTCACATCGGGTGCGATCATACCAGCACTAATGCACCGGATCCCATCAGAACTCCGAAGTTAAGCGTGCTTGGGCGAGAGTAGTACTAGGATGGGTGACCTCCTGGGAAGTCCTCGTGTTGCACCCCTTTTTCGCTGCCCGCGCGCACGTCTCTGCGGGTCCTTCCTCCCCCCTGCTCTGGTCTCTTGTTTTCCCCGCCGGCTTCGGCCATCCGGGCGAGCTGCCCCCACGCGTGCAGGTCTCGGGCGCGGAGGGGCTGGCTAGCTATCTTTCGAGGGCGTTTCGGGCAGGTCTTGGGCGCCTGGGGCGGACTCCGGAGTCCACGGGGAGGGGGTTGTCCGCCCGCCTTGGGCACGAGCTCCCCCTGGCGAGAGGCGTCCCGCATCCGCGCCGCTCGAGGGACACCACGCCGCCCCTTGGCGGTGGGTCAATGATCTCGCCCGCCCCCTCTCCCTCTCCGTCTGTCAGGGAGGCGGCGCCGATGCCGGGGGGGAGAGGATTAGGGCTGACGCGGCGCTTGTCACATCGGGTGCGATCATACCAGCACTAATGCACCGGATCCCATCAGAACTCCGAAGTTAAGCGTGCTTGGGCGAGAGTAGTACTAGGATGGGTGACCTCCTGGGAAGTCCTCGTGTTGCACCCCTTTTTCGCTGCCCGCGCGCACGTCTCTGCGGGTCCTTCCTCCCCCCTGCTCTGGTCTCTTGTTTTCCCCGCCGGCTTCGGCCATCCGGGCGAGCTGCCCCCACGCGTGCAGGTCTCGGGCGCGGAGGGGCTGGCTAGCTATCTTTCGAGGGCGTTTCGGGCAGGTCTTGGGCGCCTGGGGCGGACTCCGGAGTCCACGGGGAGGGGGTTGCCCGCCCGCCTTGGGCACGAGCTCCCCCTGGCGAGAGGCGTCCCGCATCCGCGCCGCTCGAGGGACACCACGCCGCCCCTTGGCGGTGGGTCAATGATCTCGCCCGCCCCCTCTCCCTCTCCGTCTGTCAGGGAGGCGGCGCCGATGCCGGGGGGGAGAGGATTAGGGCTGACGCGGCGCTTGTCACATCGGGTGCGATCATACCAGCACTAATGCACCGGATCCCATCAGAACTCCGAAGTTAAGCGTGCTTGGGCGAGAGTAGTACTAGGATGGGTGACCTCCTGGGAAGTCCTCGTGTTGCACCCCTTTTTCGCTGCCCGCGCGCACGTCTCTGCGGGTCCTTCCTCCCCCCTGCTCTGGTCTCTTGTTTTCCCCGCCGGCTTCGGCCATCCGGGCGAGTTGCCCCCACGCGTGCAGGTCTCGGGCGCGGAGGGGCTGGCTAGCTATCTTTCGAGGACGTTTCGGGCAGGTCTTGGGCGCCTGGGGCGGACTCCGGAGTCCACGGGGAGGGGGTTGTCCGCCCGCCTTGGGCACGAGCTCCCCCTGGCGAGAGGCGTCCCGCATCCGCGCCGCTCGAGGGACACCACGCCGCCCCTTGGCGGTGGGTCAATGATCTCGCCCGCCCCCTCTCCCTCTCCGTCTGTCAGGGAGGCGGCGCCGATGCCGGGGGGGAGAGGATTAGGGCTGACGCGGTGCTTGTCACATCGGGTGCGATCATACCAGCACTAATGCACCGGATCCCATCAGAACTCCGAAGTTAAGCGTGCTTGGGCGAGAGTAGTACTAGGATGGGTGACCTCCTGGGAAGTCCTCGTGTTGCACCCCTTTTTCGCTGCCCGCGCGCACGTCTCTGCGGGTCCTTCCTCCCCCCTGCTCTGGTCTCTTGTTTTCCCCGCCGGCTTCGGCCATCCGGGCGAGCTGCCCCCACGCGTGCAGGTCTCGGGCGCGGAGGGGCTGGCTAGCTATCTTTCGAGGGCGTTTCGGGCAGGTCTTGGGCGCCTGGGGCGGACTCCGGAGTCCACGGGGAGGGGGTTGTCCGACCGCCTTGGGCACGAGCTCCCCCTGGCGAGAGGCGTCCCGCATCCGCGCCGCTCGAGGGACACCACGCCGCCCCTTGGCGGTGGGTCAATGATCTCGCCCGCCCCCTCTCCCTCTCCGTCTGTCAGGGAGGCGGCGCCGATGCCGGGGGGGAGAGGATTAGGGCTGACGCGGCGCTTGTCACATCGGGTGCGATCATACCAGCACTAATGCACCGGGTCCCATCAGAACTCCGAAGTTAAGCGTGCTTGGGCGAGAGTAGTACTAGGATGGGTGACCTCCTGGGAAGTCCTCGTGTTGCACCCCTTTTTCGCTGCCCGCGCGCACGTCTCTGCGGGTCCTCCCTCCCCCCTGCTCTGGTCTCTTGTTTTCCCCGCCGGCTTCGGCCATCCGGGCGAGCTGCCCCCACGCGTGCAGGTCTCGGGCGCGGAGGGGCTGGCTAGCTATCTTTCGAGGGCGTTTCGGGCAGGTCTTGGGCGCCTGGGGCGGACTCCGGAGTCCACGGGGAGGGGGTTGTCCGCCCGCCTTGGGCACGAGCTCCCCCTGGCGAGAGGCGTCCCGCATCCGCGCCGCTCGAGGGACACCACGCCGCCCCTTGGCGGTGGGTCAATGATCTCGCCCGCCCCCTCTCCCTCTCCGTCTGTCAGGGAGGCGGCGCCGATGCCGGGGGGGAGAGGATTAGGGCTGACGCGGCGCTTGTCACATCGGGTGCGATCATACCAGCACTAATGCACCGGATCCCATCAGAACTCCGAAGTTAAGCGTGCTTGGGCGAGAGTAGTACTAGGATGGGTGACCTCCTGGGAAGTCCTCGTGTTGCACCCCTTTTTCGCTGCCCGCGCGCACGTCTCTGCGGGTCCTTCCTCCCCCCTGCTCTGGTCTCTTGTTTTCCCCGCCGGCTTCGGCCATCCGGGCGAGCTGCCCCCACGCGTGCAGGTCTCGGGCGCGGAGGGGCTGGCTAGCTATCTTTCGAGGGCGTTTCGGGCAGGTCTTGGGCGCCTGGGGCGGACTCCGGAGTCCACGGGGAGGGGGTTGTCCGCCCGCCTTGGGCACGAGCTCCCCCTGGCGAGAGGCGTCCCGCATCCGCGCCGCTCGAGGGACACCACGCCGCCCCTTGGCGGTGGGTCAATGATCTCGCCCGCCCCCTCTCCCTCTCCGTCTGTCAGGGAGGCGGCGCCGATGCCGGGGGGGAGAGGATTAGGGCTGACGCGGCGCTTGTCACATCGGGTGCGATCATACCAGCACTAATGCACCAGATCCCATCAGAACTCCGAAGTTAAGCGTGCTTGGGCGAGAGTAGTACTAGGATGGGTGACCTCCTGGGAAGTCCTCGTGTTGCACCCCTTTTTCGCTGCCCGCGCGCACGTCTCTGCGGGTCCTTCCTCCCCCCTGCTCTGGTCTCTTGTTTTCCCCGCCGGCTTCGACCATCCGGGCGAGCTGCCCCCACGCGTGCAGGTCTCGGGCGCGGAGGGGCTGGCTAGCTATCTTTCGAGGGCGTTTCGGGCAGGTCTTGGGCGCCTGGGGCGGACTCCGGAGTCCACGGGGAGGGGGTTGTCCGCCCGCCTTGGGCACGAGCTCCCCCTGGCGAGAGGCGTCCCGCATCCGCACCGCTCGAGGGACACCACGCCGCCCCTTGGCGGTGGGTCAATGATCTCGCCCGCCCCCTCTCCCTCTCCGTCTGTCAGGGAGGCGGCGCCGATGCCGGGGGGGAGAGGATTAGGGCTGACGCGGCGCTTGTCACATCGGGTGCGATCATACCAGCACTAATGCACCGGATCCCATCAGAACTCCGAAGTTAAGCGTGCTTGGGCGAGAGTAGTACTAGGATGGGTGACCTCCTGGGAAGTCCTCGTGTTGCACCCCTTTTTCGCTGCCCGCGCGCACGTCTCTGCGGGTCCTTCCTCCCCCCTGCTCTGGTCTCTTGTTTTCCCCGCCGGCTTCGGCCATCCGGGCGAGCTGCCCCCACGCGTGCAGGTCTCGGGCGCGGAGGGGCTGGCTAGCTATCTTTCGAGGGCGTTTCGGGCAGGTCTTGGGCGCCTGGGGCGGACTCCGGAGTCCACGGGGAGGGGGTTGTCCGCCCGCCTTGGGCACGAGCTCCCCCTGGCGAGAGGCGTCCCGCATCCGCGCCGCTCGAGGGACACCACGCCGCCCCTTGGCGGTGGGTCAATGATCTCGCCCGCCCCCTCTCCCTCTCCGTCTGTCAGGGAGGCGGCGCCGATGCCGGGGGGGAGAGGATTAGGGCTGACGCGGCGCTTGTCACATCGGGTGCGATCATACCAGCACTAATGCACCGGATCCCATCAGAACTCCGAAGTTAAGCGTGCTTGGGCGAGAGTAGTACTAGGATGGGTGACCTCCTGGGAAGTCCTCGTGTTGCACCCCTTTTTCGCTGCCCGCGCGCACGTCTCTGCGGGTCCTTCCTCCCCCCTGCTCTGGTCTCTTGTTTTCCCCGCCGGCTTCGGCCATCCGGGCGAGCTGCCCCCACGCGTGCAGGTCTCGGGCGCGGAGGGGCTGGCTAGCTATCTTTCGAGGGCGTTTCGGGCAGGTCTTGGGCGCCTGGGGCGGACTCCGGAGTCCACGGGGAGGGGGTTGTCCGCCCGCCTTGGGCACGAGCTCCCCCTGGCGAGAGGCGTCCCGCATCCGCGCCGCTCGAGGGACACCACGCCGCCCCTTGGCGGTGGGTCAATGATCTCGCCCGCCCCCTCTCCCTCTCCGTCTGTCAGGGAGGCGGCGCCGATGCCGGGGGGGAGAGGATTAGGGCTGACGCGGCGCTTGTCACATCGGGTGCGATCATACCAGCACTAATGCACCGGATCCCATCAGAACTCCGAAGTTAAGCGTGCTTGGGCGAGAGTAGTACTAGGATGGGTGACCTCCTGGGAAGTCCTCGTGTTGCACCCCTTTTTCGCTGCCCGCGCGCACGTCTCTGCGGGTCCTTCCTCCCCCCTGCTCTGGTCTCTTGTTTTCCCCGCCGGCTTCGGCCATCCGGGCGAGCTGCCCCCACGCGTGCAGGTCTCGGGCGCGGAGGGGCTGGCTAGCTATCTTTCGAGGGCGTTTCGGGCAGGTCTTGGGCGCCTGGGGCGGACTCCGGAGTCCACGGGGAGGGGGTTGTCCGCCCGCCTTGGGCACGAGCTCCCCCTGGCGAGAGGCGTCCCGCATCCGCGCCGCTCGAGGGACACCACGCCGCCCCTTGGCGGTGGGTCAATGATCTCGCCCGCCCCCTCTCCCTCTCCGTCTGTCAGGGAGGCGGCGCCGATGCCGGGGGGGAGAGGATTAGGGCTGACGCGGCGCTTGTCACATCGGGTGCGATCATACCAGCACTAATGCACCGGATCCCATCAGAACTCCGAAGTTAAGCGTGCTTGGGCGAGAGTAGTACTAGGATGGGTGACCTCCTGGGAAGTCCTCGTGTTGCACCCCTTTTTCGCTGCCCGCGCGCACGTCTCTGCGGGTCCTTCCTCCCCCCTGCTCTGGTCTCTTGTTTTCCCCGCCGGCTTCGGCCATCCGGGCGAGCTGCCCCCACGCGTGCAGGTCTCGGGCGCGGAGGGGCTGGCTAGCTATCTTTCGAGGGCGTTTCGGGCAGGTCTTGGGCGCCTGGGGCGGACTCCGGAGTCCACGGGGAGGGGGTTGTCCGCCCGCCTTGGGCACGAGCTCCCCCTGGCGAGAGGCGTCCCGCATCCGCGCCGCTCGAGGGACACCACGCCGCCCCTTGGCGGTGGGTCAATGATCTCGCCCGCCCCCTCTCCCTCTCCGTCTGTCAGGGAGGCGGCGCCGATGCCGGGGGGGAGAGGATTAGGGCTGACGCGGCGCTTGTCACATCGGGTGCGATCATACCAGCACTAATGCACCGGATCCCATCAGAACTCCGAAGTTAAGCGTGCTTGGGCGAGAGTAGTACTAGGATGGGTGACCTCCTGGGAAGTCCTCGTGTTGCACCCCTTTTTCGCTGCCCGCGCGCACGTCTCTGCGGGTCCTTCCTCCCCCCTGCTCTGGTCTCTTGTTTTCCCCGCCGGCTTCGGCCATCCGGGCGAGCTGCCCCCACGCGTGCAGGTCTCGGGCGCGGAGGGGCTGGCTAGCTATCTTTCGAGGGCGTTTCGGGCAGGTCTTGGGCGCCTGGGGCGGACTCCGGAGTCCACGGGGAGGGGGTTGTCCGCCCGCCTTGGGCACGAGCTCCCCCTGGCGAGAGGCGTCCCGCATCCGCGCCGCTCGAGGGACACCACGCCGCCCCTTGGCGGTGGGTCAATGATCTCGCCCGACCCCTCTCCCTCTCCGTCTGTCAGGGAGGCGGCGCCGATGCCGGGGGGGAGAGGATTAGGGCTGACGCGGCGCTTGTCACATTGGGTGCGATCACACCAGCACTAATGCACCGGATCCCATCAGAACTCCGAAGTTAAGCGTGCTTGGGCGAGAGTAGTACTAGGATGGGTGACCTCCTGGGAAGTCCTCGTGTTGCACCCCTTTTTCGCTGCCCGCGCGCACGTCTCTGCGGGTCCTTCCTCCCCCCTGCTCTGGTCTCTTGTTTTCCCCGCCGGCTTCGGCCATCCGGGCGAGCTGCCCCCACGCGTGCAGGTCTCGGGCGCGGAGGGGCTGGCTAGCTATCTTTCGAGGGCGTTTCGGGCAGGTCTTGGGCGCCTGGGGCGGACTCCGGAGTCCACGGGGAGGGGGTTGTCCGCCCGCCTTGGGCACGAGCTCCCCCTGGCGAGAGGCGTCCCGCATTCGCGCCGCTCGAGGGACACCACGCCGCCCCTTGGCGGTGGGTCAATGATCTCGCCCGCCCCCTCTCCCTCTCCGTCTGTCAGGGAGGCGGCGCCGATGCCGGGGGGGAGAGGATTAGGGCTGACGCGGCGCTTGTCACATCGGGTGCGATCATACCAGCACTAATGCACCGGATCCCATCAGAACTCCGAAGTTAAGCGTGCTTGGGCGAGAGTAGTACTAGGATGGGTGACCTCCTGGGAAGTCCTCGTGTTGCACCCCTTTTTCGCTGCCCGCGCGCACGTCTCTGCGGGTCCTTCCTCCCCCCTGCTCTGGTCTCTTGTTTTCCCCGCCGGCTTCGACCATCCGGGCGAGCTGCCCCCACGCGTGCAGGTCTCGGGCGCGGAGGGGCTGGCTAGCTATCTTTCGAGGGCGTTTCGGGCAGGTCTTGGGCGCCTGGGGCGGACTCCGGAGTTCACGGGGAGGGGGTTGTCCGCCCGCCTTGGGCACGAGCTCCCCCTGGCGAGAGGCGTCCCGCATCCGCGCCGCTCGAGGGACACCACGCCGCCCCTTGGCGGTGGGTCAATGATCTCGCCCGCCCCCTCTCCCTCTCCGTCTGTCAGGGAGGCGGCGCCGATGCCGGGGGGGAGAGGATTAGGGCTGACGCGGCGCTTGTCACATCGGGTGCGATCATACCAGCACCAATGCACCGGATCCCATCAGAACTCCGAAGTTAAGCGTGCTTGGGCGAGAGTAGTACTAGGATGGGTGACCTCCTGGGAAGTCCTCGTGTTGCACCCCTTTTTCGCTGCCCGCGCGCACGTCTCTGCGGGTCCCTCCTCCCCCCTGCTCTGGTCTCTTGTTTTCCCCGCCGGCTTCGACCATCCGGGCGAGCTGCCCCCACGCGTGCAGGTCTCGGGCGCGGAGGGGCTGGCTAGCTATCTTTCGAGGGCGTTTCGGGCAGGTCTTGGGCGCCTGGGGCGGACTCCGGAGTCCACGGGGAGGGGGTTGTCCGCCCGCCTTGGGCACGAGCTCCCCCTGGCGAGAGGCGTCCCGCATCCGCGCCGCTCGAGGGACACCACGCCGCCCCTTGGCGGTGGGTCAATGATCTCGCCCGCCCCCTCTCCCTCTCCGTCTGTCAGGGAGGCGGCGCCGATGCCGGGGGGGAGAGGATTAGGGCTGACGCGGCGCTTGTCACTTCGGGTGCGATCATACCAGCACTAATGCACCGGATCCCATCAGAACTCCGAAGTTAAGCGTGCTTGGGCGAGAGTAGTACTAGGATGGGTGACCTCCTGGGAAGTCCTCGTGTTGCACCCCTTTTTCGCTGCCCGCGCGCACGTCTCTGCGGGTCCTTCCTCCCCCCTGCTCTGGTCTCTTGTTTTCCCCGCCGGCTTCGGCCATCCGGGCGAGCTGCCCCCACGCGTGCAGGTCTCGGGCGCGGAGGGGCTGGCTAGCTATCTTTCGAGGGCGTTTCGGGCAGGTCTTGGGCGCCTGGGGCGGACTCCGGAGTCCACGGGGAGGGGGTTGTCCGCCCGCCTTGGGCACGAGCTCCCCCTGGCGAGAGGCGTCCCGCATCCGCGCCGCTCGAGGGACACTACGCCGCCCCTTGGCGGTGGGTCAATGATCTCGCCCGCCCCCTCTCCCTCTCCGTCTGTCAGGGAGGCGGCGCCGATGCCGGGGGGGAGAGGATTAGGGCTGACGCGGCGCTTGTCACATCGGGTGCGATCATACCAGCACCAATGCACCGGATCCCATCAGAACTCCGAAGTTAAGCGTGCTTGGGCGAGAGTAGTACTAGGATGGGTGACCTCCTGGGAAGTCCTCGTGTTGCACCCCTTTTTCGCTGCCCGCGCGCACGTCTCTGCGGGTCCTTCCTCCCCCCTGCTCTGGTCTCTTGTTTTCCCCGCCGGCTTCGGCCATCCGGGCGAGCTGCCCCCACGCGTGCAGGTCTCGGGCGCGGAGGGGCTGGCTAGCTATCTTTCGAGGGCGTTTCGGGCAGGTCTTGGGCGCCTGGGGCGGACTCCGGAGTCCACGGGGAGGGGGTTGTCCGCCCGCCTTGGGCACGAGCTCCCCCTGGCGAGAGGCGTCCCGCATCCGCGCCGCTCAAGGGACACCACGCCGCCCCTTGGCGGTGGGTCAATGATCTCGCCCGCCCCCTCTCCCTCTCCGTCTGTCAGGGAGGCGGCGCCGATGCCGGGGGGGAGAGGATTAGGGCTGACGCGGCGCTTGTCACATCGGGTGCGATCATACCAGCACTAATGCACCGGATCCCATCAGAACTCCGAAGTTAAGCGTGCTTGGGCGAGAGTAGTACTAGGATGGGTGACCTCCTGGGAAGTCCTCGTGTTGCACCCCTTTTTCGCTGCCCGCGCGCACGTCTCTGCGGGTCCTTCCTCCCCCCTGCTCTGGTCTCTTGTTTTCCCCGCCGGCTTCGGCCATCCGGGCGAGCTGCCCCCACGCGTGCAGGTCTCGGGCGCGGAGGGGCTGGCTAGCTATCTTTCGAGGGCGTTTCGGGCAGGTCTTGGGCGCCTGGGGCGGACTCCGGAGTCCACGGGGAGGGGGTTGTCCGCCCGCCTTGGGCACGAGCTCCCCCTGGCGAGAGGCGTCCCGCATCCGCGCCGCTCGAGGGACACCACGCCGCCCCTTGGCGGTGGGTCAATGATCTCGCCCGCCCCCTCTCCCTCTCCGTCTGTCAGGGGGGCGGCGCCGATGCCGGGGGGGAGAGGATTAGGGCTGACGCGGCGCTTGTCACATCGGGTGCGATCATACCAGCACTAATGCACCGGATCCCATCAGAACTCCGAAGTTAAGCGTGCTTGGGCGAGAGTAGTACTAGGATGGGTGACCTCCTGGGAAGTCCTCGTGTTGCACCCCTTTTTCGCTGCCCGCGCGCACGTCTCTGCGGGTCCTTCCTCCCCCCTGCTCTGGTCTCTTGTTTTCCCCGCCGGCTTCGGCCATCCGGGCGAGCTGCCCCCGCGCGTGCAGGTCTCGGGCGCGGAGGGGCTGGCTAGCTATCTTTCGAGGGCGTTTCGGGCAGGTCTTGGGCGCCTGGGGCGGACTCCGGAGTCCACGGGGAGGGGGTTGTCCGCCCGCCTTGGGCACGAGCTCCCCCTGGCGAGAGGCGTCCCGCATCCGCGCCGCTCGAGGGACACCACACCGCCCCTTGGCGGTGGGTCAATGATCTCGCCCGCCCCCTCTCCCTCTCCGTCTGTCAGGGAGGCGGCGCCGATGCCGGGGGGGAGAGGATTAGGGCTGACGCGGCGCTTGTCACATCGGGTGCGATCATACCAGCACTAATGCACCGGATCCCATCAGAACTCCGAAGTTAAGCGTGCTTGGGCGAGAGTAGTACTAGGATGGGTGACCTCCTGGGAAGTCCTCGTGTTGCACCCCTTTTTCGCTGCCCGCGCGCACGTCTCTGCGGGTCCTTCCTCCCCCCTGCTCTGGTCTCTTGTTTTCCCCGCCGGCTTCGGCCATCCGGGCGAGCTGCCCCCACGCGTGCAGGTCTCGGGCGCGGAGGGGCTGGCTAGCTATCTTTCGAGGGCGTTTCGGGCAGGTCTTGGGCGCCTGGGGCGGACTCCGGAGTCCACGGGGAGGGGGTTGTCCGCCCGCCTTGGGCACGAGCTCCCCCTGGCGAGAGGCGTCCCGCATCCGCGCCGCTCGAGGGACACCACGCCGCCCCTTGGCGGTGGGTCAATGATCTCGCCCGCCCCCTCTCCCTCTCCGTCTGTCAGGGAGGCGGCGCCGATGCCGGGGGGGAGAGGATTAGGGCTGACGCGGCGCTTGTCACATCGGGTGCGATCATACCAGCACTAATGCACCGGATCCCATCAGAACTCCGAAGTTAAGCGTGCTTGGGCGAGAGTAGTACTAGGATGGGTGACCTCCTGGGAAGTCCTCGTGTTGCACCCCTTTTTCGCTGCCCGCGCGCACGTCTCTGCGGGTCCTTCCTCCCCCCTGCTCTGGTCTCTTGTTTTCCCCGCCGGCTTCGGCCATCCGGGCGAGCTGCCCCCACGCGTGCAGGTCTCGGGCGCGGAGGGGCTGGCTAGCTATCTTTCGAGGGCGTTTCGGGCAGGTCTTGGGCGCCTGGGGCGGACTCCGGAGTCCACGGGGAGGGGGTTGTCCGCCCGCCTTGGGCACGAGCTCCCCCTGGCGAGAGGCGTCCCGCATCCGCGCCGCTCGAGGGACACCACGCCGCCCCTTGGCGGTGGGTCAATGATCTCGCCCGCCCCCTCTCCCTCTCCGTCTGTCAGGGAGGCGGCGCCGATGCCGGGGGGGAGAGGATTAGGGCTGACGCGGCGCTTGTCACATCGGGTGCGATCATACCAGCACTAATGCACCGGATCCCATCAGAACTCCGAAGTTAAGCGTGCTTGGGCGAGAGTAGTACTAGGATGGGTGACCTCCTGGGAAGTCCTCGTGTTGCACCCCTTTTTCGCTGCCCGCGCGCACGTCTCTGCGGGTCCTTCCTCCCCCCTGCTCTGGTCTCTTGTTTTCCCCGCCGGCTTCGGCCATCCGGGCGAGCTGCCCCCACGCGTGCAGGTCTCGGGCGCGGAGGGGCTGGCTAGCTATCTTTCGAGGGCGTTTCGGGCAGGTCTTGGGCGCCTGGGGCGGACTCCGGAGTCCACGGGGAGGGGGTTGCCCGCCCGCCTTGGGCACGAGCTCCCCCTGGCGAGAGGCGTCCCGCATCCGCGCCGCTCGAGGGACACCACGCCGCCCCTTGGCGGTGGGTCAATGATCTCGCCCGCCCCCTCTCCCTCTCCGTCTGTCAGGGAGGCGGCGCCGATGCCGGGGGGGAGAGGATTAGGGCTGACGCGGCGCTTGTCACATCGGGTGCGATCATACCAGCACTAATGCACCGGATCCCATCAGAACTCCGAAGTTAAGCGTGCTTGGGCGAGAGTAGTACTAGGATGGGTGACCTCCTGGGAAGTCCTCGTGTTGCACCCCTTTTTCGCTGCCCGCGCGCACGTCTCTGCGGGTCCTTCCTCCCCCCTGCTCTGGTCTCTTGTTTTCCCCGCCGGCTTCGGCCATCCGGGCGAGTTGCCCCCACGCGTGCAGGTCTCGGGCGCGGAGGGGCTGGCTAGCTATCTTTCGAGGACGTTTCGGGCAGGTCTTGGGCGCCTGGGGCGGACTCCGGAGTCCACGGGGAGGGGGTTGTCCGCCCGCCTTGGGCACGAGCTCCCCCTGGCGAGAGGCGTCCCGCATCCGCGCCGCTCGAGGGACACCACGCCGCCCCTTGGCGGTGGGTCAATGATCTCGCCCGCCCCCTCTCCCTCTCCGTCTGTCAGGGAGGCGGCGCCGATGCCGGGGGGGAGAGGATTAGGGCTGACGCGGTGCTTGTCACATCGGGTGCGATCATACCAGCACTAATGCACCGGATCCCATCAGAACTCCGAAGTTAAGCGTGCTTGGGCGAGAGTAGTACTAGGATGGGTGACCTCCTGGGAAGTCCTCGTGTTGCACCCCTTTTTCGCTGCCCGCGCGCACGTCTCTGCGGGTCCTTCCTCCCCCCTGCTCTGGTCTCTTGTTTTCCCCGCCGGCTTCGGCCATCCGGGCGAGCTGCCCCCACGCGTGCAGGTCTCGGGCGCGGAGGGGCTGGCTAGCTATCTTTCGAGGGCGTTTCGGGCAGGTCTTGGGCGCCTGGGGCGGACTCCGGAGTCCACGGGGAGGGGGTTGTCCGACCGCCTTGGGCACGAGCTCCCCCTGGCGAGAGGCGTCCCGCATCCGCGCCGCTCGAGGGACACCACGCCGCCCCTTGGCGGTGGGTCAATGATCTCGCCCGCCCCCTCTCCCTCTCCGTCTGTCAGGGAGGCGGCGCCGATGCCGGGGGGGAGAGGATTAGGGCTGACGCGGCGCTTGTCACATCGGGTGCGATCATACCAGCACTAATGCACCGGGTCCCATCAGAACTCCGAAGTTAAGCGTGCTTGGGCGAGAGTAGTACTAGGATGGGTGACCTCCTGGGAAGTCCTCGTGTTGCACCCCTTTTTCGCTGCCCGCGCGCACGTCTCTGCGGGTCCTCCCTCCCCCCTGCTCTGGTCTCTTGTTTTCCCCGCCGGCTTCGGCCATCCGGGCGAGCTGCCCCCACGCGTGCAGGTCTCGGGCGCGGAGGGGCTGGCTAGCTATCTTTCGAGGGCGTTTCGGGCAGGTCTTGGGCGCCTGGGGCGGACTCCGGAGTCCACGGGGAGGGGGTTGTCCGCCCGCCTTGGGCACGAGCTCCCCCTGGCGAGAGGCGTCCCGCATCCGCGCCGCTCGAGGGACACCACGCCGCCCCTTGGCGGTGGGTCAATGATCTCGCCCGCCCCCTCTCCCTCTCCGTCTGTCAGGGAGGCGGCGCCGATGCCGGGGGGGAGAGGATTAGGGCTGACGCGGCGCTTGTCACATCGGGTGCGATCATACCAGCACTAATGCACCGGATCCCATCAGAACTCCGAAGTTAAGCGTGCTTGGGCGAGAGTAGTACTAGGATGGGTGACCTCCTGGGAAGTCCTCGTGTTGCACCCCTTTTTCGCTGCCCGCGCGCACGTCTCTGCGGGTCCTTCCTCCCCCCTGCTCTGGTCTCTTGTTTTCCCCGCCGGCTTCGGCCATCCGGGCGAGCTGCCCCCACGCGTGCAGGTCTCGGGCGCGGAGGGGCTGGCTAGCTATCTTTCGAGGGCGTTTCGGGCAGGTCTTGGGCGCCTGGGGCGGACTCCGGAGTCCACGGGGAGGGGGTTGTCCGCCCGCCTTGGGCACGAGCTCCCCCTGGCGAGAGGCGTCCCGCATCCGCGCCGCTCGAGGGACACCACGCCGCCCCTTGGCGGTGGGTCAATGATCTCGCCCGCCCCCTCTCCCTCTCCGTCTGTCAGGGAGGCGGCGCCGATGCCGGGGGGGAGAGGATTAGGGCTGACGCGGCGCTTGTCACATCGGGTGCGATCATACCAGCACTAATGCACCAGATCCCATCAGAACTCCGAAGTTAAGCGTGCTTGGGCGAGAGTAGTACTAGGATGGGTGACCTCCTGGGAAGTCCTCGTGTTGCACCCCTTTTTCGCTGCCCGCGCGCACGTCTCTGCGGGTCCTTCCTCCCCCCTGCTCTGGTCTCTTGTTTTCCCCGCCGGCTTCGACCATCCGGGCGAGCTGCCCCCACGCGTGCAGGTCTCGGGCGCGGAGGGGCTGGCTAGCTATCTTTCGAGGGCGTTTCGGGCAGGTCTTGGGCGCCTGGGGCGGACTCCGGAGTCCACGGGGAGGGGGTTGTCCGCCCGCCTTGGGCACGAGCTCCCCCTGGCGAGAGGCGTCCCGCATCCGCACCGCTCGAGGGACACCACGCCGCCCCTTGGCGGTGGGTCAATGATCTCGCCCGCCCCCTCTCCCTCTCCGTCTGTCAGGGAGGCGGCGCCGATGCCGGGGGGGAGAGGATTAGGGCTGACGCGGCGCTTGTCACATCGGGTGCGATCATACCAGCACTAATGCACCGGATCCCATCAGAACTCCGAAGTTAAGCGTGCTTGGGCGAGAGTAGTACTAGGATGGGTGACCTCCTGGGAAGTCCTCGTGTTGCACCCCTTTTTCGCTGCCCGCGCGCACGTCTCTGCGGGTCCTTCCTCCCCCCTGCTCTGGTCTCTTGTTTTCCCCGCCGGCTTCGGCCATCCGGGCGAGCTGCCCCCACGCGTGCAGGTCTCGGGCGCGGAGGGGCTGGCTAGCTATCTTTCGAGGGCGTTTCGGGCAGGTCTTGGGCGCCTGGGGCGGACTCCGGAGTCCACGGGGAGGGGGTTGTCCGCCCGCCTTGGGCACGAGCTCCCCCTGGCGAGAGGCGTCCCGCATCCGCGCCGCTCGAGGGACACCACGCCGCCCCTTGGCGGTGGGTCAATGATCTCGCCCGCCCCCTCTCCCTCTCCGTCTGTCAGGGAGGCGGCGCCGATGCCGGGGGGGAGAGGATTAGGGCTGACGCGGCGCTTGTCACATCGGGTGCGATCATACCAGCACTAATGCACCGGATCCCATCAGAACTCCGAAGTTAAGCGTGCTTGGGCGAGAGTAGTACTAGGATGGGTGACCTCCTGGGAAGTCCTCGTGTTGCACCCCTTTTTCGCTGCCCGCGCGCACGTCTCTGCGGGTCCTTCCTCCCCCCTGCTCTGGTCTCTTGTTTTCCCCGCCGGCTTCGGCCATCCGGGCGAGCTGCCCCCACGCGTGCAGGTCTCGGGCGCGGAGGGGCTGGCTAGCTATCTTTCGAGGGCGTTTCGGGCAGGTCTTGGGCGCCTGGGGCGGACTCCGGAGTCCACGGGGAGGGGGTTGTCCGCCCGCCTTGGGCACGAGCTCCCCCTGGCGAGAGGCGTCCCGCATCCGCGCCGCTCGAGGGACACCACGCCGCCCCTTGGCGGTGGGTCAATGATCTCGCCCGCCCCCTCTCCCTCTCCGTCTGTCAGGGAGGCGGCGCCGATGCCGGGGGGGAGAGGATTAGGGCTGACGCGGCGCTTGTCACATCGGGTGCGATCATACCAGCACTAATGCACCGGATCCCATCAGAACTCCGAAGTTAAGCGTGCTTGGGCGAGAGTAGTACTAGGATGGGTGACCTCCTGGGAAGTCCTCGTGTTGCACCCCTTTTTCGCTGCCCGCGCGCACGTCTCTGCGGGTCCTTCCTCCCCCCTGCTCTGGTCTCTTGTTTTCCCCGCCGGCTTCGGCCATCCGGGCGAGCTGCCCCCACGCGTGCAGGTCTCGGGCGCGGAGGGGCTGGCTAGCTATCTTTCGAGGGCGTTTCGGGCAGGTCTTGGGCGCCTGGGGCGGACTCCGGAGTCCACGGGGAGGGGGTTGTCCGCCCGCCTTGGGCACGAGCTCCCCCTGGCGAGAGGCGTCCCGCATCCGCGCCGCTCGAGGGACACCACGCCGCCCCTTGGCGGTGGGTCAATGATCTCGCCCGCCCCCTCTCCCTCTCCGTCTGTCAGGGAGGCGGCGCCGATGCCGGGGGGGAGAGGATTAGGGCTGACGCGGCGCTTGTCACATCGGGTGCGATCATACCAGCACTAATGCACCGGATCCCATCAGAACTCCGAAGTTAAGCGTGCTTGGGCGAGAGTAGTACTAGGATGGGTGACCTCCTGGGAAGTCCTCGTGTTGCACCCCTTTTTCGCTGCCCGCGCGCACGTCTCTGCGGGTCCTTCCTCCCCCCTGCTCTGGTCTCTTGTTTTCCCCGCCGGCTTCGACCATCCGGGCGAGCTGCCCCCACGCGTGCAGGTCTCGGGCGCGGAGGGGCTGGCTAACTA*AG**TTCTATTAAAATAAATTGTTAATAATAAATTGAAAT**CAAAAGTCAGCATATTAGTAGGCTATTTTCAAGTTGCCATTTTGAAGCTTTCTG**AAAATTTTGTAAATGTAAAAAT**GTCTTTAAGGTACC**TATAGAAATAT**CTTTGGCACC**ATAACTAATA**GTC**AAATTTTCAAATTATTATTT**CACAG**ATAAACAATAATCATATTTGAGAAAAAAATAATA**CAATGTAGACTAGAAAGTGAAAATTCAGAGTTTCAGGATTTTCACATTTTATAAAGCTAATTTTTTTGCTTAAGATTTGTCCCTTGTTTTATCTCACTGCCTTAGTGGTGGATATTTTTCATTATTGTTAAATACTCTCTATCTAATAATATTTATAACAAAAATAATTTTAAATTATCCCATGTCATTGCAACCCAAAGGGACAATAAGCGTAGGCTTGGGTCATTTGGAAACCACATGAACTGGAGTTGGCATGACTCTGATGCACATCACCATGGACAAAAAATTGTGCCCTCTCATACAATCTTTTTCATTTATTTTAATTGTCACTAATTGGTTATGTTGGTTGATGGCCCCTTCATGTAACGTGATGATGTAGAACAAAATGGAAATACAAACTCATTATGGTGCCCAATTCTAAGTCTTATTTCTCCAACAGAGATTAATAATTTTCAAGAGTAAGGCTAGTTTCAGTCCAATTTTGATGTGCAATTTTGAATAACCAAAACAGTTAAGAAATCATCTGGTTCAAGGTGCTCAACAAACAAGGAATCAATTAATAGATTTCCATGGAAATCTCTCAATTAATCAAGCTGGGGAAAATGTAAATACTGTCTTTGAATGGTTTTCCAATAGCACCATCAAGAATATATCCTAAGATCAGGTTTTGGTGATTTGGTGGATATGATTGAGTGGAAAGTGATGTTATTAACATGTGGGTAGTGAAAATGGTAGCAGTCCTCACTCCGCTCTAGCCTCTCCTTCATGGAGCAATTAACATGATAAAATCCAATTTCAATACCTCCATATCATGATTACATGGATCTCTATAAAGTTTTCAAGAATAGAGAAAGTATCCAGACATATCAGCGATTAACAGTGGACTAATAATAGATGAAATCGTGTTGTTGACATGCAATGTGTGTATGTTTCAACGGCGAGCCTTTGTTTCATCTTTCATAGCTCTAGTTCTCCCGCATCCCCATCACCTGGTTTTTATTTAGGTCATCCCATTAGATTGAACAACCTCTGCCTATGATCATGGCTTAACCCAGTGGCGATCGAAGTCTGTGAGCCATGTGCCTCAAGCTAAATCTCAATTAGGAAATTGAGGCACTTCAGACCCTTGTTAGATGATTAGACCCACATTGCAAGGGGACTTGTGTTTTGTAAAATCTTAGTCGGCAATAAAATGGTGTTTATAATTCTGAATTTTCTAATGCTTGAAAAGCGACTGAGCCCGCATCTTCAATATCAGCCCTTCATGCTATCAAGGAACAGACGAGTACGTACCCCTTTTGGAAATGTTTTCATCCAAGGGTACGAATCCAAAGGGAGAACATCGTAATTTACGGGATCCGAATTGCGCCGGCAGTGATTGACCTTCTCCAAGGGTATCCTGGGCAACCCCCTTTGTCAAGAAATCAGACTGCAGTGATTCCTGACCGAACTCCTCGAGGGCAGCAGCTGGAGAGAGAGAGAGAGAGAGAGAGAGAGAGAGAGAGAGAGGAGAGGGTTGAGGGAGTAGGGGGACAAGCGGTTTCAAGATGATATTTTTCTTCTCCTCTGAGGTGAAGTTGGGGTAAAATACCTTCCCCGCAAAAGTCCTCTTCCTCCCTCTCCGGCACGGCTTCGTTTGTCTCTCATTCTTGGGAGTTGGGGCTTCCTTCCCTCGTCCCCGAGGAGGCAATTGACGGAGTTCCGCCCCTTCCTCCCAACCCTAGCGGGAGGGGATTTCTTGATCCGATCCGTGGAGGGCGAGAGGCCGTATTGGCGCTCTCCCAGTCTCTGATAAGCTGTCATCTGTTTTCCCCCAGGAGGAAATCTCCCGTTGTCCCTCTCTATCAGGGGAAGGAAAATTGTGGGCTCTGGTAACCGAGGAATTGGCGTCCCCCTGGATTCGAACATGTCGTCGCGGGTGCATTCTATGCATTCGCAGCGGCGCCGGCAGGGGGGTGTCGGTGGTGGCGGGAGTTCGTGGGGCTTCAGATCCAATTGGTCCAAGAGTTGGATCAACGACGGCAGTATTCGCCCGGGGGATGGCAGCTACTCCACCTATTCTTGTGCGCAGGACCTATCCGCCGTCAGTCATCGGCAGGGCTATTTTGCCGGCGGCGAAAACGCCTCCCTAGCACCTGCTTTGCTGAGGAGGAAGCCCATGTGGGAGAAAGACTTCGATGGTGGTAGTCAGCCCTGGAACCCTGGCCCTGCCGTCCGCTACTGCAGCGATGGCCCTTCTACCAGCAATAGTTCCTTTCCGCATGTGACCACAGCAGGTTGGGTAGCTGCCAAACGCGACCCAGATTGCCGATTCTTCAGCACCGACCTCGATGCGCCTTCCACAAGAAGTCGGGCGAGATTGGAGGACACCGTGGTATTTATGTCGAGGGACGAGCTTGAACGTTGCTCTCCCTCTAGAAAGGACGGCATCAATCTTCTGCGAGAGACGCATCTCCGATACTCATACTGCGCGTTCCTCCAGAACTTAGGCATCCAGCTGAATCTGTAAGTTGCCCGTCTGCTTACTTTCTCTATCTCATTCTGTACAATGTCCGTCTGTTTATTTTCTCTATCTCTATCTTTAAGTTGGCCGGATATTTATTTTCTTTATCTCAATCTGAAAGTTGTCCGTTCGTTTATTGTATTCAGGTGCCTAACTGGGTAGTTGTGCCATATGTTTTTCTTGATTGTACAGGCCTCAAACGACAATTGGGACTGCAATGGTTCTCTGCCACCGTTTTTTTGTTCGCAGATCTCATGCTTGTCATGACAGATTTGTGAGTTCTAAGTCCTTATCTTAATATTTTCCTTTATTTTTATTCGTATTCTCTTCGTACTTTGAATGTCTTAGTCATCAAATTGCCGTATTAACTCTTGAATGCTCGTCTCTATTAATGGCTTCCGTTCTGACTTTCCCAACCTCGGTTCTGTGCAGCTGGAGTGCTATGGTGTAAACACCAAAAAATTAATAATCTAGCCTCAACACAACATTCATTATTTTCATTGAAATGTGACGATGAAAAGACTTGGGGGACAAAAATTCCTCTCATTAAACAGTGTTTTTGGCAAAATATTGTGGACGCTGCTACTGTTTTTGGCTGTCATTTGAAGGTGAAGATAATGGAGGATTTCATAGATGAGGGGATGAATGGTGGCTAGATTGCTTTTCATTTAGGTATTCATTTTCCTTAGAGATGTAGCTTATGATTCTAAGTATACTAGGCATGCTACTCTTTGGATGATTTAGTATGAAGAATGGTCAGGCTTTTATTTACCACAGTTACCCAGTGCTGACGGTGGCGAGTACTTTATGCTCTAAATTGCGAGTCTGGCTTTGGGCGTTGTAATAATGGACGTTTGATTCCTTTATTTTATGGAGGCCGTCTTATCTCCTTGGTCTTGACTGTGAAAATATGACAAAATTTTCATTAGTCATAACCTGAATATTCATCTCTATTTTTAATATTTAACGTAAATGTGAAATTATCTATTCTCAGAAGGAAACTGAATGTGATTGTGATTGTGGCCGCGAGGCATTACATTGAAAGGTTCATTGAGTCAACAAACTCATCACTGTACTCTGCTTTCCCCATGCCCTAGTGATGATACTTAGTCTCCATTTGTGACTGGTTTTCTAAAAGTGAAGTGGGCTTAAGAAACTCCATCTCTCATGGAGTGGGTGAAACCAGAATGAACGGAAAACTTGGATCATTAGTTGATTCAGTGTGTCTATTTTATTGCCTTCAGATCTGCCATCGCTTGTCATTCAGTGAAGTTATCTAGTGTCAGAAAAAGTATTAAAACATTCTTTTATAACATATTTTTCAGGTTCTTGACACGTCTGTGAAGATATATTCATTTTCATTTTCTTCTGGATTTCAAACCTACAAAATTGTCTTATGTTAAATATGATCTCTTTTAATTCTTTTAATCTCTTTGGTCTTTGTAAATGTTAACGTCTCTTATTTCTGTAAAGGCATGTTTTGTTTTTAATGTCAAGTCCCTGTTTGTGGACATTTTTTGACATGTTATCTGGATAGCATGCCAGTGTAAAACTAAATTACTTCTCGACTACTAATTTAGTTCGGTCTGCAGTTGATTGCCACAGCAGCTCTGTTCCTTGCAGCAAAGTCGGAGGAGACACCATGTCCTTTAAACTCTGTACTCAGGGTATCTTGGGAAATTTGTGAAAAGCAGAATCCCTCTGTTATTCAATGTTTACTCCCTATTGTGAGTATTCTTTTGGAATCACTTTTGTTCATTTTCTTTTGAAGCTTGTTTAACTGAAATGGACTAATTCTACTAGGCATCATGGGGCTTCATTTTTGCACAATTTTCTTCACTACTTTCCATATGCAGCATATAATGGCTGTGCGAAATGCTTTTCATAAAAATAGTTTGTGGAGTAAACAGTGTAAACTTAACAGATTAAAATACAAACAAGTACCCTTGTGAATACACTGTCCATTCGTCTCTCGATAAATGTTTGTAGATAGAATGTTAGTTGTGATGCGAAATAAATGGTCGAATGTAAATGTTGATGTGCTCATCTTTTTCCACAAATATCATTCATTGTGCAAAATGTGCCATTGAATTTTAGAGTGTCCAAAAACATTGTTTTTCTAATTATCCCTTTTCCTTAAGGACTTGGTCTTGGCTTAGTTGGTATGCCTGTTAGATAGATGTGAGGAGAGTAATTCCGAGTAAATAAGTTGATGTGCCAGACTTCTTTCACTAATTACCTCACGTCATCAATATATTTTCTAAAATTTCTATTACATTTAAATACATATGAGCAGAAATTTTATGGATTTTGATATGCTTTTCTTTTGTTTGTTTTGGCAGGACTGGTTTGAACAATATCGAGAGTCGGTCATTGGCGCAGAGCACATGATACTAACAACCTTAAATTTTGAACTTGATGTCCAACATCCTTATACGCCTCTTAAATCTGTCCTTCACAAATTAGGGCTTTCCCAATCAGTCCTGCTGAATTTGGCTTTGAACTTGATAAATGAAGGGTGAGTATCTTGGTTTCAGTGATGCTTAGATAGCTGGTATTTATAGAGATGCTCGTTCTTTTATTTTTCGTGATTTGTTTGGGAAAAATTTTGTTCGGTATAGTGTCTTATTTTGAGATCTTCATTCTATTAAAGGAAAAACCATGGACAGAAAGACATTCAATAAGAACTTTCTCCAAATTATGAATTCTCGTTAATATTTAAATGGTCCCTGTATGATTAGTTGGAGCTTGCCCAAGTGTCTATGGTCGACAACAACTTTGTTGTGATTCATGGTCGTTCATTTGTACGAACTCAAGTTTTTAGATTTTAAATTTACAGCCTTCCTCATAAGTTTCCGATGGTTCATCAATTCCCATTCATCTCTCCATATCCGCCACTGATAAATGGGTTTGAGTATGATCTTCGTGTCAAATTTTGTGTTATTTCTTCTCTAGAGAATTGTTTCTTCACAGGATATTGATATTTTTGGTTGGAAATAATTGCAGGGTCTTCACGAGAATCTGTGCTCTCCTATGTGATATTCTTAGCTGGAGGATAGATAATGCTGTTCATGGAAAGCATTGTTTTATCACTGAAGTTGCGAGTGGTTTATTTTGATATATTTTAATAAAATTTGGGAAACCAACATCGCTAACTAGTTTTTGCCGATACCATCTAGTACTGCCTTCTGGGTAGTCTATTTGTTTATGTATGCATAATTTAGAATTTTTATTTTTTTATCTTATTTTAAGTTTTTTTTCTAGGTCTACCTATTTCTCCCTTTTTTCCTTCAATGTGAAGTATTTTTATCAAAACCTTTTGTATTTTATTTTTAAGTTTCACAGATTTGTGTCCCACATTTCCTATCACTCCCCCGAAAATAGTTTTACCCGTTTCTGCTGTGATTATTTGATCAAGATCAAATCTTCAAAGTTTTCTGTAGTTTCATTTATTATTGGCAGAATCGAGCAAATGTCCTTTCACCATGGGTAGCTTATATTACCAATCAGTTAAAATATAAATTTGTGTTTGGCAGTCTTGAACCAGCTACTAACTATCCATGACACCTCTGGTGGTGTCTCATCCACATCTGCCTTTACTCCAAGAAAATAGGTTAACATTCCCCCGGTCCCTGCCCCCAAAAAGAGAAAAAGAAAAGCAAGGGACATTGCTATTCGTTCATACCAAATTGACTTCATAGACTGATTCCTCTCCATATATTGTTGACTTTTTTTAGTAATACCTTTTTACTAATTTATCACCTAAAATTTAAAAAAAAACCTTTTCATCGTTCGTTTCACTTCATATTTTATTTCGTGCATCTATGTTCTTTATCTTTCCATTTTTTTATTTTTAACTCCGTCCACCATATACTTGAATGTGGAGTGAAGATTGGGAAAGTTCAGACAAGTGGGTGATAGAGATGTCTCAATGTACTTGAATTATTGTAAATGTGTGGTCACTGTGCCACACCTATATTCCCTCGAGATATTTGTATTTGTTTGCAGCGTGAGGATGGTAAAAAGAGACAACCTCCTGTGGAGTTGTTGCTGATTCCCCTTTTTTCAATGCCCATGAATTGCTGAAGGGGAAGGGCAGTAATGGGCATCCAGTCATCTCTTGCAAACGCTTTTTTTTGTGGGGGTAGGGGTCTTTACGTTGCAACTTCACTGGAGATTCCTCATCTTCAAAAATATCGTTGATAGCAAAGGTTTTCTTGGCCTTTCCCCAATTTTTCTTTTTTCTCTGACTTGGCACTGGATTACCCCACACGTGTTCATTATGCATTTTAATGGACTTATTGGCCTTTGGCTGATAAACAACCTCTATGATGAAGTATGGGGTCACAGCATTGCTGATCTAAATGTGGACGCTGAGAGCTTCCCCTATTCAGAAGAGGACTCTTTGAGTGAGTCTTTGGATCCTGATTGCCAGTTTGCTTGGCATCTTTATGATGCTTGACCTGTGTTGATATTTCCTTAAAAAGTCGATAGCTGGAATGAAGAGCAGTCATAGGTATGAATTTGGAGGGTTCCAATCGTGTCTACTTTGCATATGACGGTCTGCATCTCTACTGTTTCGCTGTGTACTTTGTTTCTGGATTTCAATGTGCACCTAGCATGATGGGTCCTGAAGATGACTAAATAATCAGATGCAATCCAAAGAGCTCAGCCTACGACTAGTTTTCTCTCCTTGTCGGCACTTCAGACAGAAAGATTTATATTCCACTGGGCATCTTATTTGCTAGTTGGTAAGGAAAGAAACACTCAGTGGCTTGTTTGCTTTCTGTATCTCTGTGTAATTTTTCCCTCCCAGCCTGTAAGTTCCCCATGCTGATGTTTTGACACTATGGTATTCTTGCCTCTGATTGATGGTTTATTTACGATTTTTCATGAATATTTATTGGTTAATGAATTAAGTATAACGGAGTCTATCCTTCAGCATTCAAATTCTATATTATTTTTCAGATTAAGAAGCTCACTTTGGCTCCAATTCAAGCCTCATCATATAGCTGCAGGGGCTGCATTCCTAGCCGCTCAGTTCCTCAATTTGGACCTGGCCTACTACCAAGGTATCTGGCATGAGTTTCAGACCACACCCTCCATTCTTCAAGGTACGGTGACACCTTCTCTGTACTGCGTCAGCCTCCTTACCAAGGTGTGCTTATTTATGATTTTGCTTGAATTGCAGATGTTGTCCAGCAGCTGTCTGAGCTTCTCTAATATGTGAAGTTGTTGTCAGTGGAGATGATGCCATTTACAAAGGTCCTGAGAGCGTTATTGTAATGTATTGGGTCAACCGTGCATTATTAATCACAGTCCTCTTCTCTCACGAGTGACGTGAGCAGAAGATGTCATGGCGCGACCTCTTCTTAATGCAAATTTTTCATTAGTATCGGATTCTATACTTCGCTGAATGGTGTTTGTACACAACTGAAGATCATCTCATTTTTCCTCACGATCTGTCTTTGGGTGTGTGTGTGTGTGTGTGAGAGAGAGAGAGAGAGAGACTGGAGCTGTCCGGCTGATTTCCTTGCGCCTCTAAACCCTCCAGCACTCTGGCGTCATGCTTCTAGAGTAAGGTACTCCCACCTATTTGGTGCCAGGCAGCCCGAAGTGTGGAGTTGTGTTTCAATGCTCTCAACACCATTAGATTATAGTACAAAAGGAGACAGATGTACCAGATGGTTCCTTCCTCTCCTTGATCATTTTCATCATCTTATCATTGTGTCTGAAGCCTATTATCGCACAATGCTCATGTACATGAACCTGATATAAGGCATGGATGGTGAGTCTGGTGCTCCAATTAATGTGTCCATGCCTAATTAAAGTTTAAGAAACCCACATTTTTAGAGCATGCAAAGCATGGGAATCATTGACTTTCCAATATTGAGTCTAAAGTCCTATGTTCTGATGCCTAAGTTGCATGACACCTTCCTTCTACATTTATTTAAGGAAGTCTAGCTTGCACTTGGGCTTATCAAGAAATCGTCTTAATTATACTTTTATAAGGCATATTATTAATCTATTTTGATTATGATATCTGGTTTACTGTTTCTTACTTCAAGACATACTTTATATAATTACATTTTTTTTAGTTTCTACGACAGTCCAAAGAGACCATTGCACCACGTATCTAGGAAGTTATTTTGGACTCAACAACATTGACTCTTGAAGCAGAAATGTTGGCCTTTTTTGTGCATCCTCTTGAAAACTTATTTGATAAATATTTACGTCTTTCGAAGCAAGAAATGGTATAATATTATATTTTTATCTAATTTATTTAATTCTATTTTCTATTTTATATTGCCATAAAATCATTTCTCTATACGAGGTAGGTGAATTCCTCTTGCGGGACATGAGCTATGAAAATCATGCCTCCTCTAATTAGTAGTTAAGTTTTTTTTTGTTCCCAAGCTGGGGTAGGTAGATTATTAAGAGGAGGATGGATTGTTTTTTCATGTCAATTTCATATTTAAAATTTTTACATTACTTATCCTAGGTAGACTTGGGTTGTCACAAAGATGCACAGGCTGCCGACACATAAGTTTGGGCTGAAACAAATAGGTTTGAGTTAGTACAAGCTAGATTGGGTTGACACGTGTTAATTTCGGTTGACACATTAACTTGACACGTTATAGTGATTTGGTTGAGTTGACACACTAAGATTTATAATTCCTCAACCTTTCCTAGTCTATATCCACTTATAAAAGGTTCATACCACATGTCTTTACTATTTTCTCTCAAATAAACTAAAACTGAAGTTTATGGGATCAGAAAATCACTATACATTATGGATTTAATGTGTTTATTTCACTATGGTAGATACAAAAGGTATTAGTTTACCTCTCTTTCCTTTTTTTAGTGACACTCATCAAACGCCATCACATAAAAAAGGGTAATTTAAGAACTTAAAAATCAATCTTGAGATCAAATAGACGAGTTTACCCATGTTATACATTTCAAGACTTAGCTCTTATTTCTACCCTCATTAAAAAAAACAGCCCATTGAGTATATTCAAGTTTCAAGTTCTAAGAACTTAGAGAGAGAAATCTGGATGCGGAAAGATTAGCTTCCCTACTAATTTTTTCTAGTGGATTGGGAGAAACCATGATAAGATGATTTTTCATTAAC

**Supplementary Figure 6.** Revealing the 5’-end border of the 60-copy array of 5S rDNA locus at *S.polyrhiza* ChrS13. **a** Nucleotide alignment of PCR-amplified 5’-end border sequence of the 60-copy array (L-1_Up) with the sequence of representative 5S rDNA unit (L-1). **b** The array border with the 5’- termini of 5S rDNA specific sequence underlined, and the sequence coding for 5S rRNA highlighted by green.

a

```
L-1_Up     TTTTGATGTTTAAGCTATGTAACTAGATACTAGACACTATTTATTTCAAATTCTTCAAGT
L-1        TTTTG-----------------------------------------------CCTGCCGC
           *****                                               *.*   *.

L-1_Up     TCTTCATACATAATAATGATTCAAATTGTATCTAACAAATCATCATCAATATTTTTTAGT
L-1        TCTCCCCGCA--------------------------------------------------
           ***.* ..**

L-1_Up     TCAATTTGAGAAGTAAATGTCTTGTTGATAATATCTTCATTTTTTAGATGTAATGTGATC
L-1        -------------------GCTTGCAATTGATGTCGCTGCCTCTGAGCTTCGTCGTGATC
                               ****. . *.**.** .....*.* ** * .. .******

L-1_Up     GAAAATAATAATACGTGGCATGAATTTCATCAAATACTTATTTTTTGTGAATTTTAGTTA
L-1        TCCTCACCTAGGGCGAGGGCCA---TCTTCCGGAGGCCCTTCCTCTGGTCAGTTCATCCC
                   **. .** **  ..   *.. .*..* .*.. *..*.**   * **.* ..

L-1_Up     ACTTTTATCATTTATCTACGATTAAAATAGATTACCTACTATTTTATTAAATAAATAAAT
L-1        CCTCTTTCCCTTCCTCCTC---------CGAACGCCCCGTCCGCCACACGCGAACTGGTT
            **.** .* **. **. *          ** ..**.  * . ..*.  .  ** *.. *

L-1_Up     AAATAAATAATATATCTATTATCGAGCATAGCCATTTGATTTCTGTTTGATCTGATCCCT
L-1        GCCGCACTACCATTTTGGGTATGCTGGGTCTTTTTTGGAGCTCAGTGGGTGCGGCTTCCC
           .    * ** .** *. . ***   * .*  .. ** ** .** **  *  * * *.**.

L-1_Up     CTCAACAAGCCTCATGACTCATTTTCTAG-TAGCATATCAGCGGTGGTTAAATACCA---
L-1        TTGCAAAAGGTTTGCCTCTTTTATGTTGGCTAGCTAGCTAGTTGGGGTTTGGGCCTATTG
           .*  * *** .*...  **. * * .*.* ****  ...**. * **** ..  *.*

L-1_Up     -----------------------------ATAACAAGCTTCTTATTTCTAACTTTTCAGA
L-1        GGCCCTTTTAGAGGGGGCACCGCCCTCATAAAAATTGCTTGCAATATTCAAATAACTTGG
                                        * **   **** . ** *..** *  .. *.

L-1_Up     ATGTACCATGTACCATTAAAATTTAATTAAATATAACAGATCATGGTGCAGTAATCTCAC
L-1        GCATGAAACGCGACGGGAGGAAAAGAGAAAATAGGGCTATAAAGTGTGGAATATTAAAGG
           ...*.  *.*.. *.  *..*   .*  ***** ..* .   *  *** *.** *   .

L-1_Up     AACATCTAGACTGCCATGCGCCCCCA----ATGCATAAGTGATTGATTGTATTACATGAG
L-1        AGGTTAGAGGAAGATATTAAATCGGAGGATGCGGATAGAAGATAGAGAAAGTCACGCTAG
           *.  *  **.  * .**  . .*  *    ..* ***.. *** **  . .*.**.. **

L-1_Up     ATGTA-----------------ATGATCAGATAGTAAAGAATGTGTCGAACAATCAATGC
L-1        AAATATTTCCATTTCGGAGTAGAAGCATGAATGGGAATGCGTGAGTGTTGATATCTATGC
           * .**                 * *  ...**.* ** * .** **   .  *** ****

L-1_Up     ACTTATGTTCCAAAGAAGCATATAAACAAATATAGATTACAGACAACCA-----AAACCA
L-1        ATAGATATCTAAGAGAGAGAGAGAGA-GAATTTAGAGTGGAGCTATCCACTTTGAGGCTC
           *.  **.*.. *.***.. * * *.* .*** **** *. ** .* ***     *..*.

L-1_Up     TAATTATCAATTATTTTCCATTCAAT-AAATCAACAAAGCTTCAACGATAATCAGCAATC
L-1        TAGCTATTTCTAACCCTTAATTAGTTCACATATGGGTGACTTTATCATCAACTGATGTGG
           **..***.  * *...*. *** . * * **  . . ..***.* *. .**......

L-1_Up     CCTTATCTCCCTCTATCCTGCGATCAAGGCAGTTTGCAACCACCCCACACTCCTAAATCC
L-1        CCCCATCACTCGCCGC--------------------AAACCATCTCTTGAGCCCCCTCCT
           **..*** *.* *...                     *****.*.* ..  **.   .*.

L-1_Up     CTTTGATACAAAAATAAAAATTGTTCAAAGTAAAAACTCCATAACTTTGCAACCACAACT
L-1        CTTCAGGGCGAATTTCGTCGCCGCCCCGA--AGCGTTCTCGTGGGCTCCCTACGTCTGGC
           ***... .*.**  * .  ...*..* .*  *. . ...*.*.. .*. * **  * . .

L-1_Up     CATATCAACCAATAAAGCCATTCGTTACGAAGCTAACAAAAAACGCAAGCTAACTATCTT
L-1        C--TTTGACCGATCCGTGGATGTCTTGCCACCCTCCC---ACGCGCAGAGTAGCTGCCTT
           *   *..***.**  .   ** . **.* *  **  *   * .****.. **.**..***

L-1_Up     TCGGGAGCGTTTTGGGCAGGTCTTGGGCACGTAGAGTGGACCTGAGAGTTGATGGGGACT
L-1        TCGGGAGCGTTTTGGGCAGGTCTTGGGCACGTAGAGTGGACCTGAGAGTTGATGGGGACT
           ************************************************************

L-1_Up     TGAGGTGGGCTACCTGCTCGTCCTATGCACGATTCCCCCTGGCGAGGGACGCGCCGCCTT
L-1        TGAGGTGGGCTACCTGCTCGTCCTATGCACGATTCCCCCTGGCGAGGGACGCGCCGCCTT
           ************************************************************

L-1_Up     GGGGGTCGGTCATGTCACCGGCCCTCTCCCTCTAGCGGGGCGGAAATGATCCGGTGCCAC
L-1        GGGGGTCGGTCATGTCACCGGCCCTCTCCCTCTAGCGGGGCGGAAATGATCCGGTGCCAC
           ************************************************************

L-1_Up     GTGGTGTTGGAGAGGGGGCAATCTGGCGAGTGGGGACGAGGGTATGAGGTGGTGTGTGGC
L-1        GTGGTGTTGGAGAGGGGGCAATCTGGCGAGTGGGGACGAGGGTATGAGGTGGTGTGTGGC
           ************************************************************

L-1_Up     GCTAGTGATGTTGGGTGCGATCATACCAGCACTAATGCACCGGATCCCATCAGAACTCCG
L-1        GCTAGTGATGTTGGGTGCGATCATACCAGCACTAATGCACCGGATCCCATCAGAACTCCG
           ************************************************************

L-1_Up     AAGTTAAGCGTG
L-1        AAGTTAAGCGTG
           ************
```

**b**

**>L-1_Up**

TTTTGATGTTTAAGCTATGTAACTAGATACTAGACACTATTTATTTCAAATTCTTCAAGTTCTTCATACATAATAATGATTCAAATTGTATCTAACAAATCATCATCAATATTTTTTAGTTCAATTTGAGAAGTAAATGTCTTGTTGATAATATCTTCATTTTTTAGATGTAATGTGATCGAAAATAATAATACGTGGCATGAATTTCATCAAATACTTATTTTTTGTGAATTTTAGTTAACTTTTATCATTTATCTACGATTAAAATAGATTACCTACTATTTTATTAAATAAATAAATAAATAAATAATATATCTATTATCGAGCATAGCCATTTGATTTCTGTTTGATCTGATCCCTCTCAACAAGCCTCATGACTCATTTTCTAGTAGCATATCAGCGGTGGTTAAATACCAATAACAAGCTTCTTATTTCTAACTTTTCAGAATGTACCATGTACCATTAAAATTTAATTAAATATAACAGATCATGGTGCAGTAATCTCACAACATCTAGACTGCCATGCGCCCCCAATGCATAAGTGATTGATTGTATTACATGAGATGTAATGATCAGATAGTAAAGAATGTGTCGAACAATCAATGCACTTATGTTCCAAAGAAGCATATAAACAAATATAGATTACAGACAACCAAAACCATAATTATCAATTATTTTCCATTCAATAAATCAACAAAGCTTCAACGATAATCAGCAATCCCTTATCTCCCTCTATCCTGCGATCAAGGCAGTTTGCAACCACCCCACACTCCTAAATCCCTTTGATACAAAAATAAAAATTGTTCAAAGTAAAAACTCCATAACTTTGCAACCACAACTCATATCAACCAATAAAGCCATTCGTTACGAAGCTAACAAAAAACGCAAGCTAACTATCTTTCGGGAGCGTTTTGGGCAGGTCTTGGGCACGTAGAGTGGACCTGAGAGTTGATGGGGACTTGAGGTGGGCTACCTGCTCGTCCTATGCACGATTCCCCCTGGCGAGGGACGCGCCGCCTTGGGGGTCGGTCATGTCACCGGCCCTCTCCCTCTAGCGGGGCGGAAATGATCCGGTGCCACGTGGTGTTGGAGAGGGGGCAATCTGGCGAGTGGGGACGAGGGTATGAGGTGGTGTGTGGCGCTAGTGATGTTGGGTGCGATCATACCAGCACTAATGCACCGGATCCCATCAGAACTCCGAAGTTAAGCGTG

**Supplementary Figure 7.** **Revealing the 3’-end border of the 60-copy array of 5S rDNA locus at *S.polyrhiza* Chr13.** **a** Nucleotide alignment of PCR-amplified 3’-end border sequence of the 60-copy array (L-1_Up) with the sequence of representative 5S rDNA unit (L-18). Sequence encoding 5S rRNA is highlighted by green; a 13bp insert of the sequence L-18 is marked by red. **b** The array border with the 3’- termini of 5S rDNA specific sequence underlined, and the sequence coding for 5S rRNA highlighted by green.

**a**

```
5S-Down    CTTGGGCGAGAGTAGTACTAGGATGGGTGACCTCCTGGGAAGTCCTCGTGTTGCACCCCT
L-18       CTTGGGCGAGAGTAGTACTAGGATGGGTGACCTCCTGGGAAGTCCTCGTGTTGCACCCCT
           ************************************************************

5S-Down    TTTGCC-------------TGCCGCTCTCCCCGCAGCTTGCAATTGATGTCGCTGCCTCT
L-18       TTTGCCTGCCGCAGCTTTGTGCCGCTCTCCCCGCAGCTTGCAATTGATGTCGCTGCCTCT
           ******             *****************************************

5S-Down    GAGCTTCGTCGTGATCTCCTCACCTAGGGCGAGGGCCATCTTCCGGAGGCCCTTCCTCTG
L-18       GAGCTTCGTCGTGATCTCCTCACCTAGGGCGAGGGCCATCTTCCGGAGGCCCTTCCTCTG
           ************************************************************

5S-Down    GTCAGTTCATCCCCCTCTTTCCCTTCCTCCTCCGAACGCCTCGTCCGCCACACGCGAACT
L-18       GTCAGTTCATCCCCCTCTTTCCCTTCCTCCTCCGAACGCCTCGTCCGCCACACGCGAACT
           ************************************************************

5S-Down    GGTTGCCGCACTACCATTTTGGGTATGCTGGGTCTTTTTTGGAGCTCAGTGGGTGCGGCT
L-18       GGTTGCCGCACTACCATTTTGGGTATGCTGGGTCTTTTTTGGAGCTCAGTGGGTGCGGCT
           ************************************************************

5S-Down    TCCCTTGCAAAAGGTTTGCCTCTTTTATGTTGGCTAGCTAGCTAGTTGGGGTTTGGGCCT
L-18       TCCCTTGCAAAAGGTTTGCCTCTTTTATGTTGGCTAGCTAGCTAGTTGGGGTTTGGGCCT
           ************************************************************

5S-Down    ATTGGGCCCTTTTAGAGGGGGCACCGCCCTCATAAAAATTGCTTGCAATATTCAAATAAC
L-18       ATTGGGCCCTTTTAGAGGGGGCACCGCCCTCATAAAAATTGCTTGCAATATTCAAATAAC
           ************************************************************

5S-Down    TTGGGCATGAAACGCGACGGGAGGAAAAGAGAAAATAGGGCTATAAAGTGTGGAATATTA
L-18       TTGGGCATGAAACGCGACGGGAGGAAAAGAGAAAATAGGGCTATAAAGTGTGGAATATTA
           ************************************************************

5S-Down    AAGGAGGTTAGAGGAAGATATTAAATCGGAGGATGCGGATAGAAGATAGAGAAAGTCACG
L-18       AAGGAGGTTAGAGGAAGATATTAAATCGGAGGATGCGGATAGAAGATAGAGAAAGTCACG
           ************************************************************

5S-Down    CTAGAAATATTTCCATTTCGGAGTAGAAGCATGAATGGGAATGCGTGAGTGTTGATATCT
L-18       CTAGAAATATTTCCATTTCGGAGTAGAAGCATGAATGGGAATGCGTGAGTGTTGATATCT
           ************************************************************

5S-Down    ATGCATAGATATCTAAGAGAGAGAGAGAGAGAATTTAGAGTGGAGCTATCCACTTTGAGG
L-18       ATGCATAGATATCTAAGAGAGAGAGAGAGAGAATTTAGAGTGGAGCTATCCACTTTGAGG
           ************************************************************

5S-Down    CTCTAGCTATTTCTAACCCTTAATTAGTTCACATATGGGTGACTTTATCATCAACTGATG
L-18       CTCTAGCTATTTCTAACCCTTAATTAGTTCACATATGGGTGACTTTATCATCAACTGATG
           ************************************************************

5S-Down    TGGCCCCATCACTCGCCGCAAACCATCTCTTGAGCCCCCTCCTCTTCAGGGCGAATTTCG
L-18       TGGCCCCATCACTCGCCGCAAACCATCTCTTGAGCCCCCTCCTCTTCAGGGCGAATTTCG
           ************************************************************

5S-Down    TCGCCGCCCCGAAGCGTTCTCGTGGGCTCCCTACGTCTAGCCTTTGACCGATCCGTGGAT
L-18       TCGCCGCCCCGAAGCGTTCTCGTGGGCTCCCTACGTCTGGCCTTTGACCGATCCGTGGAT
           **************************************.*********************

5S-Down    GTCTTGCCATCCTCCCACGCGCAGAGTAGCTGCCTTTCGGGAGCGTTTTGGGCAGGTCTT
L-18       GTCTTGCCACCCTCCCACGCGCAGAGTAGCTGCCTTTCGGGAGCGTTTTGGGCAGGTCTT
           *********.**************************************************

5S-Down    TG----------------------------------------------------TGTTTT
L-18       GGGCACGTAGAGTGGACCTGAGAGTTGATGGGGACTTGAGGTGGGCTACCTGCTCGTCCT
            *                                                    .**..*

5S-Down    GT---TTTTTCTTCGTTGTAAGGAGTGCATT-----------------TGTCTCTAGCCC
L-18       ATGCACGATTCCCCCTGGCGAGGGACGCGCCGCCTTGGGGGTCGGTCATGTCACCGGCCC
           .*   .  ***..* * *..***...**...                 **** *..****

5S-Down    TTTCC-------------------------------------------------------
L-18       TCTCCCTCTAGCGGGGCGGAAATGATCCGGTGCCACGTGGTGTTGGAGAGGGGGCAATCT
           *.***

5S-Down    ----------------------------------------------TA
L-18       GGCGAGTGGGGACGAGGGTATGAGGTGGTGTGTGGCGCTAGTGATGTT
                                                         *
```

**b**

**>5S-Down**

CTTGGGCGAGAGTAGTACTAGGATGGGTGACCTCCTGGGAAGTCCTCGTGTTGCACCCCTTTTGCCTGCCGCTCTCCCCGCAGCTTGCAATTGATGTCGCTGCCTCTGAGCTTCGTCGTGATCTCCTCACCTAGGGCGAGGGCCATCTTCCGGAGGCCCTTCCTCTGGTCAGTTCATCCCCCTCTTTCCCTTCCTCCTCCGAACGCCTCGTCCGCCACACGCGAACTGGTTGCCGCACTACCATTTTGGGTATGCTGGGTCTTTTTTGGAGCTCAGTGGGTGCGGCTTCCCTTGCAAAAGGTTTGCCTCTTTTATGTTGGCTAGCTAGCTAGTTGGGGTTTGGGCCTATTGGGCCCTTTTAGAGGGGGCACCGCCCTCATAAAAATTGCTTGCAATATTCAAATAACTTGGGCATGAAACGCGACGGGAGGAAAAGAGAAAATAGGGCTATAAAGTGTGGAATATTAAAGGAGGTTAGAGGAAGATATTAAATCGGAGGATGCGGATAGAAGATAGAGAAAGTCACGCTAGAAATATTTCCATTTCGGAGTAGAAGCATGAATGGGAATGCGTGAGTGTTGATATCTATGCATAGATATCTAAGAGAGAGAGAGAGAGAATTTAGAGTGGAGCTATCCACTTTGAGGCTCTAGCTATTTCTAACCCTTAATTAGTTCACATATGGGTGACTTTATCATCAACTGATGTGGCCCCATCACTCGCCGCAAACCATCTCTTGAGCCCCCTCCTCTTCAGGGCGAATTTCGTCGCCGCCCCGAAGCGTTCTCGTGGGCTCCCTACGTCTAGCCTTTGACCGATCCGTGGATGTCTTGCCATCCTCCCACGCGCAGAGTAGCTGCCTTTCGGGAGCGTTTTGGGCAGGTCTTTGTGTTTTGTTTTTTCTTCGTTGTAAGGAGTGCATTTGTCTCTAGCCCTTTCCTA

Supplementary Figure 8. Verification of the small 5S rDNA array at the *S.polyrhiza* ChrS13 locus.

**a** Schematic representation of the array structure with two genes encoding 5S rRNA (green box), IGS sequences (thick grey line) and non-rDNA chromosomal sequences (black line). Arrows indicate locations of primers used for amplification of PCR fragments; vertical red box shows position of the 13bp insert.

**b** Nucleotide sequence alignment of assembled small cluster 5S rDNA array (Chr13-cl-S) with 3’-end sequences of the major 64-copy array (positions 97,741 to 100,947, according to the sequence in Supplementary Fig. 9 ). The Chr13-cl-S sequence is reversed-complemented; the positions of primers used to clone the cluster borders are highlighted by grey; sequences coding for 5S rRNA are highlighted by green; GAGA stretches are highlighted by yellow; positions of 13bp inserts in the corresponding NTS sequence is marked by red. c rDNA array (Chr13-cl-S) with sequences representing 5S rDNA repeats marked by *underlined Italic*.

**a**

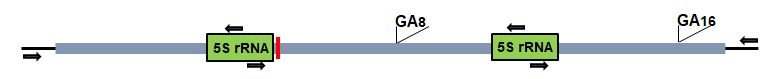

**b**

```
Chr13-cl-S TATATCTCCTCACCCAAGGTGAGGGCCATCTTCTGGAGACCTTTCCTCTAGTCAGTTCAT
Chr13-cl-L GTGATCTCCTCACCTAGGGCGAGGGCCATCTTCCGGAGGCCCTTCCTCTGGTCAGTTCAT
              ***********.*.**.*************.****.**.*******.**********

Chr13-cl-S CCCCCTCTTTCCCTTTCTCCTCCGAACGCTTTGTCTGCCGCATGCGAACTAGTTCCCGTG
Chr13-cl-L CCCCCTCTTTCCCTTCCTCCTCCGAACGCCTCGTCCGCCACACGCGAACTGGTTGCCGCA
           ***************.*************.*.***.***.**.*******.*** ***..

Chr13-cl-S CTACCATTTTGAGTATGCTGGGTCTTCTTTGGAGCTCAATGGATGTGGCTTCCCTTACAA
Chr13-cl-L CTACCATTTTGGGTATGCTGGGTCTTTTTTGGAGCTCAGTGGGTGCGGCTTCCCTTGCAA
           ***********.**************.***********.***.**.**********.***

Chr13-cl-S AAGGTTTGCCTCTTTTATGTTGGCTAGCTAGCTAGTTGGGGTTTGGGCCTATTGGGCCCT
Chr13-cl-L AAGGTTTGCCTCTTTTATGTTGGCTAGCTAGCTAGTTGGGGTTTGGGCCTATTGGGCCCT
           ************************************************************

Chr13-cl-S TTTAGAGGGGGCACCGCC--TATAAAAATTGCTTGCAATATTCAAATAACTTGGGCATGA
Chr13-cl-L TTTAGAGGGGGCACCGCCCTCATAAAAATTGCTTGCAATATTCAAATAACTTGGGCATGA
           ******************  .***************************************

Chr13-cl-S AATGCGACGGGAGGAAGAGAGAAAATAGGGCTATAAAGTGTGGAATATTAAAGGAGGTTA
Chr13-cl-L AACGCGACGGGAGGAAAAGAGAAAATAGGGCTATAAAGTGTGGAATATTAAAGGAGGTTA
           **.*************.*******************************************

Chr13-cl-S GACAAAGGTATTAAATCGGAGGTTGTGGATAGAAGATAGAGAAAGTCATGTTAGAAATAT
Chr13-cl-L GAGGAAGATATTAAATCGGAGGATGCGGATAGAAGATAGAGAAAGTCACGCTAGAAATAT
           ** .***.************** **.**********************.*.*********

Chr13-cl-S TTTAATTT-AGAGTAGATGCATGAATGGGAATGCTTAAGTGTTGATATCTATGCATAGAT
Chr13-cl-L TTCCATTTCGGAGTAGAAGCATGAATGGGAATGCGTGAGTGTTGATATCTATGCATAGAT
           **. **** .******* **************** *.***********************

Chr13-cl-S ATCTA-----------------------AGTGGAGCTATCCACTTTGAGGCTCTAGCTAT
Chr13-cl-L ATCTAAGAGAGAGAGAGAGAGAATTTAGAGTGGAGCTATCCACTTTGAGGCTCTAGCTAT
           *****                       ********************************

Chr13-cl-S TTCTAACCCGTAATTAGTTCACATATGGGTGACTTTATCATCAACTGATGTGGCCCCATC
Chr13-cl-L TTCTAACCCTTAATTAGTTCACATATGGGTGACTTTATCATCAACTGATGTGGCCCCATC
           ********* **************************************************

Chr13-cl-S ACTCGCTGCAAATTATCTCTTGAGCCCCCTCCTCTTCTAGGCAAATTTCGTCACTGCCCC
Chr13-cl-L ACTCGCCGCAAACCATCTCTTGAGCCCCCTCCTCTTCAGGGCGAATTTCGTCGCCGCCCC
           ******.*****..*********************** .***.*********.*.*****

Chr13-cl-S GAAGCGTTCTCGTGGGCTCCCTACGTCTGGCCCTTGACTGATCCGTGGATGTCTTGCCAC
Chr13-cl-L GAAGCGTTCTCGTGGGCTCCCTACGTCTGGCCTTTGACCGATCCGTGGATGTCTTGCCAC
           ********************************.*****.*********************

Chr13-cl-S CCCCCCACGCACAGAGTAGCTGCCTTTCGGGAGCGTTTTGGGCAGGTCTTGGGCACGTAG
Chr13-cl-L CCTCCCACGCGCAGAGTAGCTGCCTTTCGGGAGCGTTTTGGGCAGGTCTTGGGCACGTAG
           **.*******.*************************************************

Chr13-cl-S AGTGGACCTGAGAGTTGATGGGGACTTGAGGTGGGCTACCTGCTCGTCCTATGCACGATT
Chr13-cl-L AGTGGACCTGAGAGTTGATGGGGACTTGAGGTGGGCTACCTGCTCGTCCTATGCACGATT
           ************************************************************

Chr13-cl-S CCCCCTGGCGAGGGACGCGCCGCCTTGGGGGTCGGTCATGTCACCGGCCCTCTCCCTCTA
Chr13-cl-L CCCCCTGGCGAGGGACGCGCCGCCTTGGGGGTCGGTCATGTCACCGGCCCTCTCCCTCTA
           ************************************************************

Chr13-cl-S GCGGGGCGGAAATGATCCGGTGCCACGTGgTGTTGGAGAGGGGGCAATCTGGCGAGTGGG
Chr13-cl-L GCGGGGCGGAAATGATCCGGTGCCACGTGGTGTTGGAGAGGGGGCAATCTGGCGAGTGGG
           ************************************************************

Chr13-cl-S GACGAGGGTATGAGTTGTCGTGTGGCGCTAGTGATGTTGGGTGCGATCATACTAGCACTA
Chr13-cl-L GACGAGGGTATGAGGTGGTGTGTGGCGCTAGTGATGTTGGGTGCGATCATACCAGCACTA
           ************** ** .*********************************.*******

Chr13-cl-S ATGCACCGGATCCCATCAGAACTCCGAAGTTAAGCGTGCTTGGGCGAGAGTAGTACTAGG
Chr13-cl-L ATGCACCGGATCCCATCAGAACTCCGAAGTTAAGCGTGCTTGGGCGAGAGTAGTACTAGG
           ************************************************************

Chr13-cl-S ATGGGTGACCTCTTGGGAAGTCCTCGTGTTGCACCCCTTTTGCCTGCCGCAGCTTTGTGC
Chr13-cl-L ATGGGTGACCTCCTGGGAAGTCCTCGTGTTGCACCCCTTTTGCCTGCCGCAGCTTTGTGC
           ************.***********************************************

Chr13-cl-S CGCTCTCCCCGCAGCTTGCAATTGATGTCGCTGCCTCTGAGCTTCGTCGTGATCTCCTCA
Chr13-cl-L CGCTCTCCCCGCAGCTTGCAATTGATGTCGCTGCCTCTGAGCTTCGTCGTGATCTCCTCA
           ************************************************************

Chr13-cl-S CCTAGGGCGAGGGCCATCTTCCGGAGGCCCTTCCTCTGGTCAGTTCATCCCCCTCTTTCC
Chr13-cl-L CCTAGGGCGAGGGCCATCTTCCGGAGGCCCTTCCTCTGGTCAGTTCATCCCCCTCTTTCC
           ************************************************************

Chr13-cl-S CTTCCTCCTCCGAACGCCTCGTCCGCCACACGCGAACTGGTTGCCGCACTATCATTTTGG
Chr13-cl-L CTTCCTCCTCCGAACGCCTCGTCCGCCACACGCGAACTGGTTGCCGCACTACCATTTTGG
           ***************************************************.********

Chr13-cl-S GTATGCTGGGTCTTTTTTGGAGCTCAGTGGGTGCGGCTTCCCTTGCAAAAGGTTTGCCTC
Chr13-cl-L GTATGCTGGGTCTTTTTTGGAGCTCAGTGGGTGCGGCTTCCCTTGCAAAAGGTTTGCCTC
           ************************************************************

Chr13-cl-S TTTTATGTTGGCTAGCTAGCTAGTTGGGGTTTGGGCCTATTGGGCCCTTTTAGAGGGGGC
Chr13-cl-L TTTTATGTTGGCTAGCTAGCTAGTTGGGGTTTGGGCCTATTGGGCCCTTTTAGAGGGGGC
           ************************************************************

Chr13-cl-S ACCGCCCTCATAAAAATTGCTTGCAATATTCAAATAACTTGGGCATGAAACGCGACGGGA
Chr13-cl-L ACCGCCCTCATAAAAATTGCTTGCAATATTCAAATAACTTGGGCATGAAACGCGACGGGA
           ************************************************************

Chr13-cl-S GGAAAAGAGAAAATAGGGCTATAAAGTGTGGAATATTAAAGGAGGTTAGAGGAAGATATT
Chr13-cl-L GGAAAAGAGAAAATAGGGCTATAAAGTGTGGAATATTAAAGGAGGTTAGAGGAAGATATT
           ************************************************************

Chr13-cl-S AAATCGGAGGATGCGGGTAGAAGATAGAGAAAGTCACGCTAGAAATATTTCCATTTCGGA
Chr13-cl-L AAATCGGAGGATGCGGATAGAAGATAGAGAAAGTCACGCTAGAAATATTTCCATTTCGGA
           ****************.*******************************************

Chr13-cl-S GTAGAAGCATGAATGGGAATGCGTGAGTGTTGATATCTATGCATAGATATCTAAGAGAGA
Chr13-cl-L GTAGAAGCATGAATGGGAATGCGTGAGTGTTGATATCTATGCATAGATATCTAAGAGAGA
           ************************************************************

Chr13-cl-S GAGAGAGAGAATTTAGAGTGGAGCTATCCACTTTGAGGCTCTAGCTATTTCTAACCCTTA
Chr13-cl-L GAGAGAGAGAATTTAGAGTGGAGCTATCCACTTTGAGGCTCTAGCTATTTCTAACCCTTA
           ************************************************************

Chr13-cl-S ATTAGTTCACATATGGGTGACTTTATCATCAACTGATGTGGCCCCATCACTCGCCGCAAA
Chr13-cl-L ATTAGTTCACATATGGGTGACTTTATCATCAACTGATGTGGCCCCATCACTCGCCGCAAA
           ************************************************************

Chr13-cl-S CCATCTCTTGAGCCCCCTCCTCTTCAGGGCGAATTTCGTCGCCGCCCCGAAGCGTTCTCG
Chr13-cl-L CCATCTCTTGAGCCCCCTCCTCTTCAGGGCGAATTTCGTCGCCGCCCCGAAGCGTTCTCG
           ************************************************************

Chr13-cl-S TGGGCTCCCTACGTCTGGCCTTTGACCGATCCGTGGATGTCTTGCCACCCTCCCACGCGC
Chr13-cl-L TGGGCTCCCTACGTCTGGCCTTTGACCGATCCGTGGATGTCTTGCCACCCTCCCACGCGC
           ************************************************************

Chr13-cl-S AGAGTAGCTGCCTTTCGGGAGCGTTTTGGGCAGGTCTTGGGCACGTAGAGTGGACCTGAG
Chr13-cl-L AGAGTAGCTGCCTTTCGGGAGCGTTTTGGGCAGGTCTTGGGCACGTAGAGTGGACCTGAG
           ************************************************************

Chr13-cl-S AGTTGATGGGGACTTGAGGTGGGCTACCTGCTCGTCCTATGCACGATTCCCCCTGGCGAG
Chr13-cl-L AGTTGATGGGGACTTGAGGTGGGCTACCTGCTCGTCCTATGCACGATTCCCCCTGGCGAG
           ************************************************************

Chr13-cl-S GGACGCGCCGCCTTGGGGGTCGGTCATGTCACCGGCCCTCTCCCTCTAGCGGGGCGGAAA
Chr13-cl-L GGACGCGCCGCCTTGGGGGTCGGTCATGTCACCGGCCCTCTCCCTCTAGCGGGGCGGAAA
           ************************************************************

Chr13-cl-S TGATCCGGTGCCACGTGGTGTTGGAGAGGGGGCAATCTGGCGAGTGGGGACGAGGGTATG
Chr13-cl-L TGATCCGGTGCCACGTGGTGTTGGAGAGGGGGCAATCTGGCGAGTGGGGACGAGGGTATG
           ************************************************************

Chr13-cl-S AGGTGGTGTGTGGCGCTAGTGATGTTGGGTGCGATCATACTAGCACTAATGCACCGGATC
Chr13-cl-L AGGTGGTGTGTGGCGCTAGTGATGTTGGGTGCGATCATACCAGCACTAATGCACCGGATC
           ****************************************.*******************

Chr13-cl-S CCATCAGAACTCCGAAGTTAAGCGTGCTTGGGCGAGAGTAGTACTAGGATGGGTGACCTC
Chr13-cl-L CCATCAGAACTCCGAAGTTAAGCGTGCTTGGGCGAGAGTAGTACTAGGATGGGTGACCTC
           ************************************************************

Chr13-cl-S CTGGGAAGTCCTCGTGTTGCACCCCTTTTGCCTGCCGCTCTCCCCGCAGCTTGCAATTGA
Chr13-cl-L CTGGGAAGTCCTCGTGTTGCACCCCTTTTGCCTGCCGCTCTCCCCGCAGCTTGCAATTGA
           ************************************************************

Chr13-cl-S TGTCGCTGCCTCTGAGCTTCGTCGTGATCTCCTCACCTAGGGCGAGGGCCATCTTCCGGA
Chr13-cl-L TGTCGCTGCCTCTGAGCTTCGTCGTGATCTCCTCACCTAGGGCGAGGGCCATCTTCCGGA
           ************************************************************

Chr13-cl-S GGCCCTTCCTCTGGTCAGTTCATCCCCCTCTTTCCCTTCCTCCTCCGAACGCcTCGTCCG
Chr13-cl-L GGCCCTTCCTCTGGTCAGTTCATCCCCCTCTTTCCCTTCCTCCTCCGAACGCCTCGTCCG
           ************************************************************

Chr13-cl-S CCACACGCGAACTGGTTGCCGCACTACCATTTTGGGTATGCTGGGTCTTTTTTGGAGCTC
Chr13-cl-L CCACACGCGAACTGGTTGCCGCACTACCATTTTGGGTATGCTGGGTCTTTTTTGGAGCTC
           ************************************************************

Chr13-cl-S AGTGGGTGCGGCTTCCCTTGCAAAAGGTTTGCCTCTTTTATGTTGGCTAGCTAGCTAGTT
Chr13-cl-L AGTGGGTGCGGCTTCCCTTGCAAAAGGTTTGCCTCTTTTATGTTGGCTAGCTAGCTAGTT
           ************************************************************

Chr13-cl-S GGGGTTTGGGcCTATTGGGCCCTTTTAGAGGGGGCACCGCCCTCATAAAAATTGCTTGCA
Chr13-cl-L GGGGTTTGGGCCTATTGGGCCCTTTTAGAGGGGGCACCGCCCTCATAAAAATTGCTTGCA
           ************************************************************

Chr13-cl-S ATATTCAAATAACTTGGGCATGAAACGCGACGGGAGAAAAAGAGAAAATAGGGCTATAAA
Chr13-cl-L ATATTCAAATAACTTGGGCATGAAACGCGACGGGAGGAAAAGAGAAAATAGGGCTATAAA
           ************************************.***********************

Chr13-cl-S GTGTGGAATATTAAAGGAGGTTAGAGGAAGATATTAAATCGGAGGATGCGGATAGAAGAT
Chr13-cl-L GTGTGGAATATTAAAGGAGGTTAGAGGAAGATATTAAATCGGAGGATGCGGATAGAAGAT
           ************************************************************

Chr13-cl-S AGAGAAAGTCACGCTAGAAATATTTCCATTTCGGAGTAGAAGCATGAATGGGAATGCATG
Chr13-cl-L AGAGAAAGTCACGCTAGAAATATTTCCATTTCGGAGTAGAAGCATGAATGGGAATGCGTG
           *********************************************************.**

Chr13-cl-S AGTGTTGATATCTATGCATAGATATCTAAGAGAGAGAGAGAGAGAGAGAGAGAGAGAGAG
Chr13-cl-L AGTGTTGATATCTATGCATAGATATCTA----------------AGAGAGAGAGAGAGAG
           ****************************                ****************

Chr13-cl-S AATTTAGAGTGGAGCTATCCACTTTGAGGCTCTAGCTATTTCTAACCCTTAATTAGTTCA
Chr13-cl-L AATTTAGAGTGGAGCTATCCACTTTGAGGCTCTAGCTATTTCTAACCCTTAATTAGTTCA
           ************************************************************

Chr13-cl-S CATATGGGTGACTTTATCATCAACTGATGTGGCCCCATCACTCGCCGCAAACCATCTCTT
Chr13-cl-L CATATGGGTGACTTTATCATCAACTGATGTGGCCCCATCACTCGCCGCAAACCATCTCTT
           ************************************************************

Chr13-cl-S GAGCCCCCTCCTCTtCAGGGCGAATTTCGTCGCCGCCCCGAAGCGTTCTCGTGGGCTCCC
Chr13-cl-L GAGCCCCCTCCTCTTCAGGGCGAATTTCGTCGCCGCCCCGAAGCGTTCTCGTGGGCTCCC
           ************************************************************

Chr13-cl-S TACGTCTGGCCTTTGACCGATCCGTGGATGTCTTGCCACCCTCCCACGCGCAGAGTAGCT
Chr13-cl-L TACGTCTAGCCTTTGACCGATCCGTGGATGTCTTGCCATCCTCCCACGCGCAGAGTAGCT
           *******.******************************.*********************

Chr13-cl-S GCCTTTCGGGAGCGTTTTGGGCAGGTCTTGGGCACGTAGAGTGGACCCGAGAGTTGATGG
Chr13-cl-L GCCTTTCGGGAGCGTTTTGGGCAGGTCTTTG-----------------------------
           ***************************** *

Chr13-cl-S GGACTTGAGGTGGGCTAACTGCTTGTCCTATGCACGATTCCCCTTGGCGAAGGATGCGTC
Chr13-cl-L -----------------------TGTTTTGT---TTTTTCTTCGTTGTAAGGAGTGCATT
                                  ***..*.*   .  ***..* * *..*.*..***.*.


Chr13-cl-S GCCTTGGGGGTCGGTCATGTCACTGGCCCCATCCCTCTAGTGGAGCTATGTGGATGCGCT
Chr13-cl-L -----------------TGTCTCTAGCCCTTTCCTAAGAGTTAGGGTTTGG---------
                            **** **.****. ***.   *** ..* * **

Chr13-cl-S TCCCTT
Chr13-cl-L ------
```

**c**

Chr13-cl-S

TAT*ATCTCCTCACCCAAGGTGAGGGCCATCTTCTGGAGACCTTTCCTCTAGTCAGTTCATCCCCCTCTTTCCCTTTCTCCTCCGAACGCTTTGTCTGCCGCATGCGAACTAGTTCCCGTGCTACCATTTTGAGTATGCTGGGTCTTCTTTGGAGCTCAATGGATGTGGCTTCCCTTACAAAAGGTTTGCCTCTTTTATGTTGGCTAGCTAGCTAGTTGGGGTTTGGGCCTATTGGGCCCTTTTAGAGGGGGCACCGCCTATAAAAATTGCTTGCAATATTCAAATAACTTGGGCATGAAATGCGACGGGAGGAAGAGAGAAAATAGGGCTATAAAGTGTGGAATATTAAAGGAGGTTAGACAAAGGTATTAAATCGGAGGTTGTGGATAGAAGATAGAGAAAGTCATGTTAGAAATATTTTAATTTAGAGTAGATGCATGAATGGGAATGCTTAAGTGTTGATATCTATGCATAGATATCTAAGTGGAGCTATCCACTTTGAGGCTCTAGCTATTTCTAACCCGTAATTAGTTCACATATGGGTGACTTTATCATCAACTGATGTGGCCCCATCACTCGCTGCAAATTATCTCTTGAGCCCCCTCCTCTTCTAGGCAAATTTCGTCACTGCCCCGAAGCGTTCTCGTGGGCTCCCTACGTCTGGCCCTTGACTGATCCGTGGATGTCTTGCCACCCCCCCACGCACAGAGTAGCTGCCTTTCGGGAGCGTTTTGGGCAGGTCTTGGGCACGTAGAGTGGACCTGAGAGTTGATGGGGACTTGAGGTGGGCTACCTGCTCGTCCTATGCACGATTCCCCCTGGCGAGGGACGCGCCGCCTTGGGGGTCGGTCATGTCACCGGCCCTCTCCCTCTAGCGGGGCGGAAATGATCCGGTGCCACGTGgTGTTGGAGAGGGGGCAATCTGGCGAGTGGGGACGAGGGTATGAGTTGTCGTGTGGCGCTAGTGATGTTGGGTGCGATCATACTAGCACTAATGCACCGGATCCCATCAGAACTCCGAAGTTAAGCGTGCTTGGGCGAGAGTAGTACTAGGATGGGTGACCTCTTGGGAAGTCCTCGTGTTGCACCCCTTTTGCCTGCCGCAGCTTTGTGCCGCTCTCCCCGCAGCTTGCAATTGATGTCGCTGCCTCTGAGCTTCGTCGTGATCTCCTCACCTAGGGCGAGGGCCATCTTCCGGAGGCCCTTCCTCTGGTCAGTTCATCCCCCTCTTTCCCTTCCTCCTCCGAACGCCTCGTCCGCCACACGCGAACTGGTTGCCGCACTATCATTTTGGGTATGCTGGGTCTTTTTTGGAGCTCAGTGGGTGCGGCTTCCCTTGCAAAAGGTTTGCCTCTTTTATGTTGGCTAGCTAGCTAGTTGGGGTTTGGGCCTATTGGGCCCTTTTAGAGGGGGCACCGCCCTCATAAAAATTGCTTGCAATATTCAAATAACTTGGGCATGAAACGCGACGGGAGGAAAAGAGAAAATAGGGCTATAAAGTGTGGAATATTAAAGGAGGTTAGAGGAAGATATTAAATCGGAGGATGCGGGTAGAAGATAGAGAAAGTCACGCTAGAAATATTTCCATTTCGGAGTAGAAGCATGAATGGGAATGCGTGAGTGTTGATATCTATGCATAGATATCTAAGAGAGAGAGAGAGAGAATTTAGAGTGGAGCTATCCACTTTGAGGCTCTAGCTATTTCTAACCCTTAATTAGTTCACATATGGGTGACTTTATCATCAACTGATGTGGCCCCATCACTCGCCGCAAACCATCTCTTGAGCCCCCTCCTCTTCAGGGCGAATTTCGTCGCCGCCCCGAAGCGTTCTCGTGGGCTCCCTACGTCTGGCCTTTGACCGATCCGTGGATGTCTTGCCACCCTCCCACGCGCAGAGTAGCTGCCTTTCGGGAGCGTTTTGGGCAGGTCTTGGGCACGTAGAGTGGACCTGAGAGTTGATGGGGACTTGAGGTGGGCTACCTGCTCGTCCTATGCACGATTCCCCCTGGCGAGGGACGCGCCGCCTTGGGGGTCGGTCATGTCACCGGCCCTCTCCCTCTAGCGGGGCGGAAATGATCCGGTGCCACGTGGTGTTGGAGAGGGGGCAATCTGGCGAGTGGGGACGAGGGTATGAGGTGGTGTGTGGCGCTAGTGATGTTGGGTGCGATCATACTAGCACTAATGCACCGGATCCCATCAGAACTCCGAAGTTAAGCGTGCTTGGGCGAGAGTAGTACTAGGATGGGTGACCTCCTGGGAAGTCCTCGTGTTGCACCCCTTTTGCCTGCCGCTCTCCCCGCAGCTTGCAATTGATGTCGCTGCCTCTGAGCTTCGTCGTGATCTCCTCACCTAGGGCGAGGGCCATCTTCCGGAGGCCCTTCCTCTGGTCAGTTCATCCCCCTCTTTCCCTTCCTCCTCCGAACGCcTCGTCCGCCACACGCGAACTGGTTGCCGCACTACCATTTTGGGTATGCTGGGTCTTTTTTGGAGCTCAGTGGGTGCGGCTTCCCTTGCAAAAGGTTTGCCTCTTTTATGTTGGCTAGCTAGCTAGTTGGGGTTTGGGcCTATTGGGCCCTTTTAGAGGGGGCACCGCCCTCATAAAAATTGCTTGCAATATTCAAATAACTTGGGCATGAAACGCGACGGGAGAAAAAGAGAAAATAGGGCTATAAAGTGTGGAATATTAAAGGAGGTTAGAGGAAGATATTAAATCGGAGGATGCGGATAGAAGATAGAGAAAGTCACGCTAGAAATATTTCCATTTCGGAGTAGAAGCATGAATGGGAATGCATGAGTGTTGATATCTATGCATAGATATCTAAGAGAGAGAGAGAGAGAGAGAGAGAGAGAGAGAATTTAGAGTGGAGCTATCCACTTTGAGGCTCTAGCTATTTCTAACCCTTAATTAGTTCACATATGGGTGACTTTATCATCAACTGATGTGGCCCCATCACTCGCCGCAAACCATCTCTTGAGCCCCCTCCTCTtCAGGGCGAATTTCGTCGCCGCCCCGAAGCGTTCTCGTGGGCTCCCTACGTCTGGCCTTTGACCGATCCGTGGATGTCTTGCCACCCTCCCACGCGCAGAGTAGCTGCCTTTCGGGAGCGTTTTGGGCAGGTCTT*GGGCACGTAGAGTGGACCCGAGAGTTGATGGGGACTTGAGGTGGGCTAACTGCTTGTCCTATGCACGATTCCCCTTGGCGAAGGATGCGTCGCCTTGGGGGTCGGTCATGTCACTGGCCCCATCCCTCTAGTGGAGCTATGTGGATGCGCTTCCCTT

**Supplementary Figure 9.** **Consensus nucleotide sequence of the *Spirodela polyrhiza* 5S rDNA locus at ChrS13_B.** The sequence is composed of 10kb long upstream chromosomal region followed by a cluster of two copies of 5S rRNA genes (reverse orientation), followed by 12,359 bp of intermediate chromosomal region, followed by a cluster of sixty copies of 5S rRNA genes (direct orientation), followed by 10kb long locus’s downstream chromosome region. The clusters of 5S rDNA repeats are marked by *underlined Italic*; sequences coding for 5S rRNA are highlighted by green; GAGA stretches are highlighted by yellow; DNA stretches with more than 90% of T/A are in **bold**; positions of 13bp inserts in the corresponding NTS sequences are marked by red; locations of primers used for amplification borders sequences of the rDNA clusters are highlighted by grey.

**>ChrS13_B-5SrDNA (110,911 bp)**

C**TAAAAATTTACT**CCATGTCATGAAACTGGATGTCTATACCAATAAGCCCTCAGTACACGTGGGACACTGTCTCTATTCATTGATCTCGTAGTTCTGTCCACAGTTTTTTGCTCTTCTTGTCCACCATTTCCTACATGCAGATTTCAAGGCAGTTCAAGTCATTTACCACAGCTCAAATC**TATTTCATTAAT**GCTGCC**TTCATTTTTTATT**CTCATCC**TTTTGTTATTAT**CGGACAGTAGGCACAGATTAAATGGTCTATTGGTAATTCATTCACTGCCCTAAACCTTCTAG**AAAACATTTTTT**CAG**ATTTAATTATTTAATTTT**CTGAATGTC**AATAAAAAACAAAT**GAC**TTTTTTCAATA**GC**TTAATGAATA**GAAGCCCCTCTTGTTTATACTTGAATTCAAGTGGTATGTTCTCTCATATCATTCCC**TAAAATGATTAA**GGTGATTGTTCTCAGATTGTCCATTACCCATTTCCATGTCTGGTTTTCTGCATGAATGAACTACAGGTTGGTGCGCATGAGGTTAGCCCTGTTGAACTCTCGCAACTGCTTGAACCTCCTAC**ATATTTCTATT**GCCCGTTCGACACTCGAAAACACCACCGGCTGGTCCTCGTTCGGGCTGAAGATGATGATGCATGTCTGCACATTCGGAGAGCATATGCAGCTCCTCGGCCTTCTTAATGATTCTGCTCCTTCTCTTCTTCAAGGTTTCCCGTCTTCGAGCATCATCCTCTAGCCACCGGATCCTGATCTTCTTCCTGGAAATGAAAGCTCTCCCGGTGCGTCTATGTCGTGCCTGAGGAGAGGAGGGAATGGTGGGTACATGATGGGATCTAATAACACCCTCTCCCCCAGGTCCCATTATGCGCCGACCATTTCCCCCTCTGCCTCAGGCGCAAGATAGACGCACCAGGGAATTTTCGTTTCCAGGGAAAAG**ATCAAAATAT**GGTGGTTAGAGGATGGCGCTCGAAGGAGGGCAGCCTTGAAGAAGAGAAGGAGCGCAATCATTAAGAGAGCCGAGGAGCTGCATATCCTCTTCGATGTGCAGATATGCATCATGATTTTCAGCTCTAACAAAGACTGGTCGGAGGTGTTTTGGAATTTTGAACGGTCAGTGGAAATGTGTAAGAGGTTCAAGCAGCTGCCAGAGTTCAACATAGCTAACCTCATGCGCACCAACCTGTAGTTCATTCATGCAGAAAACCAGACATGGAAATTGGTAATGGACAATCTGACAACAATCTCC**TTAATCATTTTA**GGGAATGATATGAGAGAACATACCGCTTGAATTCAAG**TATAAACAAT**GGGGGCTTC**TATTCATTAA**GCTATTGAAGAAAGTCATCTGATTTGTATTGACATTCAG**AAAATTAAATAATTAAA**GCTG**AAAAATGTTTT**CTAGATGGTCGAAGGTAGTGAATAAATTACCAATGGACCATTTAATCTGTCCCTATTGTCAGATAAGAACAAAAGGGGAG**AATAAAACAAA**GGTAGTGTTACCGGCTCTTATAAGTGACTTGGGCTGCCTTGAAATCTGCATGCAGGAAATGGTGGACAAGAAGAGCTAAAAGCTGTCGACTGAGAACAACGAGATAAATGATGTAGAGACATGTGTCCCATGTGTACTGAGAGCTCATTGGTACAGAAATCAAAACTCATGACATGGAG**TAAAGTTTTA**GGGTGCAAGAGATGGTCCAAAGATTGTTGAATCAACACTACAAGCAGTACAGCACACTTCGCTCGGTGGCTCCATTTTCTCAATTACCATAGGCATTGCAACAGCAGCCAGGGATGAGCCTGGTTGGATTGGATGCTGAAGAACCCTCTGTAGGCTCTCATTTGCCGATTCCTTTATGGGGAACACAAACCCCCACGTTAGACAGAGCAGATGTCATCTCAACAACTAAGATTAGGGAATCTAATCCTTGGTTTCCTTGTTGGGTAGGACCAGTTATTTGTTGATTATTGATGATATTTCTC**AAACATTTTTT**CATGTGTTTCGTTTCCTTGCACCATCAACTTATATTCACAATTACAGGGAAAACAATGCCTGTGACTTTGCACCATTATGTTTCTGAATGGTTATTTGCTTCACTAATAGTAAGTGCCTTCAGATTACATTGAACGAATGAGTCTTC**TACATTTTTTT**CAAGTCGTAGGTAAAC**ATTTATTAATCATTAAA**CAAAACTTGACCTTTCATGCAAAATCGGAATCACATCTATAGGTCAGTTTGAAGTTAGTACG**AATGAATTTTA**CATTTTCC**ATATATATTGTTTA**CCCTGTAGCCCATCTAATCAAAGGCATGCTGAAC**ATTTCATTTT**GTGTTGATGAACATTTCATTGTATGTTATGGTTATCATGAAACCATTTAACTTATCAGAGTGCATAATCAATCCTATTTTC**TTACATATTT**CAAGAATCATTGAAG**TTTTTATTTTATTGAATTAAAT**C**ATTATTAATTCAATTATATTT**C**TAAAATTTCTATAAT**GTGAAATGCC**AAAAACTTAAT**CATCATGCTCTATCAAAGAATTGCATGTCCCTTTGAAAGAAAGGC**ATATATTAAGAT**CTCACC**ATATAATATTT**CC**ACAAAATATA**CTG**ATATTAAAATAATTTATCTTAAA**G**TATTATTACAAT**GATCCAAACCACTAAATAGCATAACATCCATCATGGAATTTTCACTCCCAAGAAATGTATACAACTTGTGACATG**TAAACATTTAATTAAT**C**AAAAAACTTT**GCATGCACAATTTGGCCAGCTGGCTAACTTGAGGGATAAAC**TTTAGTATTATT**CGTCCCCCTTCGAACTTCTTCATGCCCTCTAGAAACTTCAATTTG**ATCAAAAAATA**GGACAGCTATGTCGTGTTACTAAC**ATAATGATTTTTTT**CCTCTAATTAGGAAAACATAC**TTATATGAAAA**CCTATTGACTCC**TAATTCATTTTTT**GATTCATTG**AAAAAAAATTCTT**GGCTTAATTGTAACTGG**AATTATAATTAAAATTTATTA**GCTTAGTTGTGATATTGC**ATATTTTATCTTATT**GGACAC**ATTAGTATTT**CTCATCAACC**AAATATATTGT**GGTAGCTAAGTTCAGTCTTTATCAAACC**TTATAAAATAGTTTA**GTTATGACTAATGTGTGCACAAAATTTAGGGGAC**AATTTTTTGTTT**CCACC**TTAAATTTGTT**CACAAGAATATTATTTGGAGGCTTCAAATATTTTCTTATCATTCATGAAGGTCAATTATGAGAAATTAGAGCAGTTTAAGCCTACTACAGCCTTAGTGGCAATGTCTAAGAATGCTAACACACTACCATTAGTATGTTGAATCACTTAGCAACAATGGCCAATTTAGAGGTATAATCTATGGTTGATTAGTTAGTGCCAATTTGTTGACACAAAAGATCTGTGAATATGAAAGGAATAGATATCCTCCATTGTTTGACCCCCTATGCTCCTAAATGTAGAATATAACTAATCATTGACATTTTGACTTTAGAG**TAATTAAATATATT**GAAGGTAGTGTGGACACCCTTGGACATGTTTTTCG**ATATTAAATTTAT**GAG**AAAATATATTATTAA**GG**TTATATATTATTAA**GGTAGTGTCGATGCCCTTAGAC**AATAAATTGA**GTGGTTTACAACAAATCATGATTC**AACAATAATAAA**CCCAACAACAAGCTATTCAACAATTGCTGTAGTAAC**AATAAGAAAAT**CAACCTATGCATCAACTCACATAAAAACCATATGCAGGTCAACAATTTAGACTAAATGGAGGCCAGCTACCAAGTCAACCCATAGTCAATCCACAAAATCGTCCTCTTGGTTTTCAGC**ATCAAATTAATAT**GGTAAATACAAGTTTTGAAGAACCTTTTGTACCATCTCAAATGGACCATGAAGGTCATTTTTCAGC**TATAAAATAGAAA**GGATTTGGGTGATCCATATCGAAAAAAGGAAGATGGAAAAGACCTCTTTTCTATG**AAAAAAAATTTAGAA**GATC**AAAAATAGAAAA**CTATAGAAGAAGATGATGAATTTAGTGATGATGAAGAGGTTG**AAAAATCAAA**GGTTC**AAGTAAATATT**GATCTAGAGGAATATGTTCCTAGTACACCTTTTCCTCATGCATTAAAACC**AAAAGTTAAAAA**GGGATTGATACCTAATGATTAGCTG**ATAAAACTTTTT**GAGCAAGTGCAGATCACGATTCCTCTCTTAGATGCCATCAAACATCTACTATG**TTATGTAAAATTTTT**GAAGGAGATGTGCACATCTAAAAGAGAGAAGAGAATCACTCTTTAATCACAAGTGTAATTACTCTTGATCAATTACCCATGAAATGTAAAGATCTAAGTGCACTTCTAATC**TCTTATAAAATT**GATGAAC**TTACTTTTAATAAT**GCATTACTTGATTTAGGTGCTAGTG**TTAATCTTTTA**CTCACTGCCATTGTTGAAC**ATTTTAGAATT**GGTGAGCTCAAACCAGCCTC**TATAATTCTT**C**AATTTGTAAATA**GATCTACAAAGAAACTTAGAG**AAATATTATAAGAT**GTAATTGACAAGGTAGGCG**AATGTTATTTT**CCAGTTG**ATTTTCTATTTTTA**GACATGATATAGTCACATG**AATTAAAACAAAT**GGC**TATCATTTTAA**GATGCTCATTTCTTGCTACAGC**AAAAGTAAATATTAATT**GTGAGTCAATAAGGTTAG**ATATTTTCTTTA**GGG**AACATAAAAT**GCCTC**TTAATATTTTTA**GACCAGCTAATTATCCAAAGGGAGAAGAAGATTATTCTGACATTGATAAGACAG**ATATTTTTCTT**G**ATAATATATTTTTA**CATGATAAATCATCTG**ATTTATATTCTA**CAACACTATAGGAAGAAG**AATTTTTTCTT**CCTTCCCTTGCCTATCCAACATTAGATCTCAAACCTTTGCTCAAAAGTCTC**AAATATAATCAAT**CCATACATGCCAC**TTTTTTAGAA**CCCTTGCTTG**AATTTCTTTTT**CTTCTTCATCCTTGATGATTGTGAAACCTCTTTTCTTTGGTACGACCATAATTGAGCTCAGCCATTTGCTGTTAGAAATAGG**ATATATAAAATTA**GCATTTAACC**ATTTAATTATCT**CATCATTAACAACATCTTTCATACTGAGATTTAATCTCTTTTGCATCTTACTCCTTAAGTCTTGCATCTTCCTCTAAATAGATGCTATGCATGTAAGTTTCTAGATCAATATCTTTGAAATC**AGAAATATTT**CACCCAATTGCATTATTAC**ATGATAATAAAA**CATCAATCAAACATGATTCTTGGGTAGAAG**TTAAGTTTTTT**G**TAATAATGATA**GGG**TTAAATGTAT**GTACTAAGAAAGATCATTTTCCCTTACC**ATTTATGAATTAAATTTTT**GAGATGATAGTAGTTC**AAAATATTTTTTCTTT**CTTGATGGCTACTCTGG**TTATAATCAAATA**GCCATCAATCTAAATGATCAAGAAAAG**ATTATATTTACAT**GACCGTTTGGTACTTTTGCTTTC**ATTAGAATAT**C**TTTTGATTTAT**GTAATACTCTTACTACATTTTAGTGATACATG**TTATCTATTTTTTAAAAAT**GATTCGAG**AATTTTTAGAA**GC**ATTATAAACAATTTAT**CAATCTTTAGATATTCCTTTGATGATTGCC**TTATTAATCTT**GAGAAAGTATTAACCAAATACATTG**AATCAAATTT**GGTTTTGAGTTGGGAGAAGAGTC**ATTTCTAAAA**GATCTAAGGGCTAGACAAAATACCAAGAGGTGGGGTGAATTGGTGTTTCAAGAATCTTGGATTTTCTTTCTAGATAATAGATGTGTCAACACGTGCTAACATGGCTCTATG**TAATTATCTA**CATATAGGCTTACTTTCTCTCTCCTCTAGTGGGTTCTCTCATGAGTTTCTCTCTCTCACTAATGAGATAAAGAC**TTTTTATACTA**GTTCGTCTCTCATCTAGCTACTTTCAGTACTGAAGGACCTTTCCTGCTAG**TTTTTAACTATTT**CTCTTAGAAATACAC**ACTAAAAATAAT**GTGATTAACAACCTAATACCTCAGGTTTGGCCTCTGTAGAGATCCAACTCCTAGGTCAAATAGGGCTTAAAGCACCTTGTGCCTTTGCTCAAAACCACTCTCCCTCATAACC**TAATTATCTTA**GGGCTTGATAGAGAGCTTCTTTGAGAGATTCAGGC**TTTTTTTTATCAT**C**AATTATTGAT**GAAGAATGATGATGTAGTCTATGCACAGTGATGTTTATTGATC**ATAAGAAATATTTTT**CGTTATAGATTGATGTAGAATCATTGTTGTTAATCTACATAGCTTTCTTTGCTTATAAGGCATCATCCTCTAATAG**ATTTATGTATTTTT**CTCAAGAG**ATAAAATTGAAAA**GTTAGTTCAAATACTTATC**ATTTTCTAAAA**CGTGTGGCAATGACCTTTGATCATTGCGAATTTG**ATAACTTATTATT**CCTTAGTC**TATTTAATTTAAAATTT**GAAGGAAGATTTATCTCTCATATATGAACAACTTTTTTGTTGACACTATTTTGAG**ATTTAAACATTAA**GCTAGTC**ATTTTTTTCAT**GTAAGTAGC**TAATATGTTAT**CGTCC**AAATTTCTTATTT**GAGACTCATGAGAGC**TCTTTAATATTT**CGAGTC**ATTTTTATTTTCTTTTAAT**GATGCATGTGAC**TAATAAATAAA**GGG**TTATACTTAAAT**GGTCAAATAATC**ATAGATAAAT**GAAC**TTAAAACATAA**CCATCATTTTTCCATCTCATTTACC**TATTAAATATTTATTTT**C**TAATAAATATTTATTGATATAAAT**GTGACGCTTCACTTATTCATCAATTCACTTAATCTGAAAG**TTTAATTTCTT**GGATCAATGGG**TTAAACATAATAT**CACAAGATAAATGTAGAGTTTATTG**TATAAAGTATATTA**GAC**ATTTATGATAAT**CTTGTGAACAATATCACATATC**AATTATATATTTT**CTC**ATTTAATTTTAATATCAAAATA**CATGTCCAAGGGCATCAACAATCTCTCATATTTTGATACTTACAAAACAATCTTAAG**AAAATATATATATATTA**GCACAATGATTTGCACAAAC**AATATAACAATA**GATTCACGCAC**ATAGTAAATAT**GAAGGCTTT**GACAATAATAAAAATAA**GTATCATGTGTCAACATGTGTCAAAGTAATGCACAAATAGAACATTCAAAGTTTAGTATTCACATCTATTCACACAATTCATCCACAGAGGTTCAAGC**AATAAAATTTTAATT**GC**ATTTAATACAT**CATTACTTTCCTTGAGCAAGTGCTCCTTATAAGTACGTGCAGTCATATAGATTCTCAATCTTTCTCCC**ATTTTTTAAATTTT**CTCAATCTTTCTCTCTCTTTTTG**TAAGATAAAAAAAAT**C**AAATCATAAATA**GTCGAGAAAATAG**TTATAGTTTTT**GCTCGAGTAGAAAATCATGAAAAATGCAAATGAATCTG**TAAAATATTGT**CTTCAAGGTCTTGAAGTCATTCTTTCATCATATCAAGATATTACTTCACTTGTTCCACGCTTTTGTTG**AATTTATTAACAAT**GGGATGAATGTTAAAGAAGGTGAAAGTATGAGATGATGTATGAGAGGATATGGATGATGAAGCCACAGACCATCATTATGAGAGTAATTCATTCTATGAAGAGTGGATTTGTCAATAATGTCATGTGAAATACTACTGATGAATTCTTCGATAAAGAC**ATTTATATTGTAATA**CTTGAGGACATGTGTAATGACATAGATAAGTGAAAGAATGAGATCATATTTCAAGATTGTTTCCATCTGTG**TTAATAATAGAT**CTTCGAAATCAAGTGGAATGTCATGAATG**ATACAATATAA**GATAGAC**ATTTATTTTTTT**GACAACCTATCATGTCCACCACTTCTATGCTATATAAGTGATGACCAATAAGAAAATGAATGACTCTATATGATAGTGGAAG**ATTAAATACTTTTATATTATT**GAGATATACATGTTATGTTCCATTCAATGTGGCAAAAAAAGCATTGATGGTTTCTTTGGTGATGGCCAAGGGAGACCTAGAGG**AAATATTACTAAT**GAC**ATTTAAGATTTTA**GGAGTAAGATCAATAAATCCTCCTATTACATATGTGGTGAGGCCAACAAGGGTGATAGAGGCAC**TATTATAAAACAT**CATAGCAAGATTTGG**ATAATTTTTAATAT**GCTC**ATAAACAAAAT**GAG**TAATTATAAAGA**C**ATCAAAATAAATATT**C**ATATTAATCT**GCTACGATTCAAAGAATTTCAAATCAAGAAATTTGTAGCAAAAGACTTTACAGTCTTTGAAAG**ATTTCTAAAATTTT**GCTTCATGGGCTGTAG**TAAGAAATAATA**GAG**AATAATAATAA**GGAAACCTATCTAAGATATGCTAGGGTACAACACCTTTTCTAGTTCTTTTGGAACTCTGTTTCTTTGCTTTGAGTTCCTACAAGG**TTATTTTTTTTGTA**GC**TTTTGATTTT**CGTATAGTGGAAGGCTTTCTAATGATTGATTTGGTCTTTTTCATG**ATTTGTAATA**GTTC**AAATAGTTTT**GGGCTGTGGGCAGAGTTAGGG**TTTTCTATTT**CTGTTTCATTAATGATGGGCGAGACTTAGGGTTATATATGCTTG**AATTTAAAAAAAATTGAAAT**GGAGGCGCTAACATTTGTAG**ATAATTTTTGAA**CAACCACTACGATTTGGTCATCAATCATCATAGGTGATGGAGGAATGGGTGATTTAGGTCTTGTAGGAGAACCAAATGGAGGGGTTAAAAAAGGCATATG**TAATTTTTCAA**CCATGGGAGAAGGTGGGAAAGTCTATTTAGGAGATGAATCAATGACAATGTTTTTCTCTTTATCCTTCTTGTAAATTCTTAGATGCCGATGGGTTCTTGAAGGAGGAATAGAAGGGTAGGGTGTCGGGTATCAACCATGGTGAATCTAGGG**TTTTGAAAAA**GAGAGTTGATTGAGATGAGATATGAATACAATGCATTGCATAGAATAAGATACTTTTTGTTGACTCAAGAGTTGACCCGCTATGATGTGAGCTGTGGCTTGAAGTCTTGTGGGCATGGTGGAATGAAGGTCTTCAAATGGTGAAAACGAATGAGAAGGTAAAGTTATCGAACAATGAAGTGTGGCCTTTTAATGGAAGAAGAGGGATATAGAC**AAAATATATAATTTTTTTATATTTT**CC**TTAAAAAAAAT**C**TCTTAATTAA**CACTTCAAAATCATGTGATGAAGACCGTGTCAACATAGAATCTACACATATGGGTTTC**ATATCAATTATA**G**AATGATAATTA**G**AATATATTTTCAATAAAT**CACAACGTATCATC**AAAAATATTAGAAATTT**GA**TAAAAATATCAAT**CAATTTGC**ATATTATAAGTTAT**C**AAATAATTGAATTAAAT**CTGTCTTCGGAAAGTCTTTTTCCTC**ATTAAAAAATTCA**CCTTTAGGGAAGAATTCTCTTTGGCTTGTAAGATGAAACAATGGTTGATTGTCTCCATCACCCCCATTCATTGTTATGAAATACTTTGATATGCTTTGGTATGTTATAGATCAATACTAGTATGAAAATGACCTTTGTTCATATTGTACATAGAGGTGTGCACTAGTATGACATGGATTAACGTTGACAATAAGAAACTGTGCAAAACTGTATTTGGAGTGTTTGTTGGCACTAGGCAATCAACATCAACATTAGAAATCACC**TAAAATAGTA**CCAGTGC**AAAAAATGAT**GTGAAATGGTGTATGCAAGTG**ATGTTTATAAT**GCTTAAGTAGTATAGAGGAGTATGGGGCACAAGAAACCCAGTGCAAAGTGTACATTGGCACATACCAACATCTCTAATTCAGTGAGAGTGAGCTGGCATTACTCCTTGAGTGGGAGTCACAATATCATGAACTAGAAGGTTCATGAAACTAACCAAACAGTGCAAAACAGTGGAATGATAATTGTACTCATTATGGTGCAGCCCAATAGGCCCAAACCCCAGCTAGCTAGCCAACATAAAAGGGGCAAACCTTTTGCAAGGGAAGCGCATCCACATAGCTCCA*CTAGAGGGaTGGGGCCAGTGACATGACCGACCCCCAAGGCGACGCATCCTTCGCCAAGGGGAATCGTGCATAGGACAAGCAGTTAGCCCACCTCAAGTCCCCATCAACTCTCGGGTCCACTCTACGTGCCCAAGACCTGCCCAAAACGCTCCCGAAAGGCAGCTACTCTGCGCGTGGGAGGGTGGCAAGACATCCACGGATCGGTCAAAGGCCAGACGTAGGGAGCCCACGAGAACGCTTCGGGGCGGCGACGAAATTCGCCCTGaAGAGGAGGGGGCTCAAGAGATGGTTTGCGGCGAGTGATGGGGCCACATCAGTTGATGATAAAGTCACCCATATGTGAACTAATTAAGGGTTAGAAATAGCTAGAGCCTCAAAGTGGATAGCTCCACTCTAAATTCTCTCTCTCTCTCTCTCTCTCTCTCTCTCTCTTAGATATCTATGCATAGATATCAACACTCATGCATTCCCATTCATGCTTCTACTCCGAAATGGAAATATTTCTAGCGTGACTTTCTCTATCTTCTATCCGCATCCTCCGATTTAATATCTTCCTCTAACCTCCTTTAATATTCCACACTTTATAGCCCTATTTTCTCTTTTTCTCCCGTCGCGTTTCATGCCCAAGTTATTTGAATATTGCAAGCAATTTTTATGAGGGCGGTGCCCCCTCTAAAAGGGCCCAATAGgCCCAAACCCCAACTAGCTAGCTAGCCAACATAAAAGAGGCAAACCTTTTGCAAGGGAAGCCGCACCCACTGAGCTCCAAAAAAGACCCAGCATACCCAAAATGGTAGTGCGGCAACCAGTTCGCGTGTGGCGGACGAgGCGTTCGGAGGAGGAAGGGAAAGAGGGGGATGAACTGACCAGAGGAAGGGCCTCCGGAAGATGGCCCTCGCCCTAGGTGAGGAGATCACGACGAAGCTCAGAGGCAGCGACATCAATTGCAAGCTGCGGGGAGAGCGGCAGGCAAAAGGGGTGCAACACGAGGACTTCCCAGGAGGTCACCCATCCTAGTACTACTCTCGCCCAAGCACGCTTAACTTCGGAGTTCTGATGGGATCCGGTGCATTAGTGCTAGTATGATCGCACCCAACATCACTAGCGCCACACACCACCTCATACCCTCGTCCCCACTCGCCAGATTGCCCCCTCTCCAACACCACGTGGCACCGGATCATTTCCGCCCCGCTAGAGGGAGAGGGCCGGTGACATGACCGACCCCCAAGGCGGCGCGTCCCTCGCCAGGGGGAATCGTGCATAGGACGAGCAGGTAGCCCACCTCAAGTCCCCATCAACTCTCAGGTCCACTCTACGTGCCCAAGACCTGCCCAAAACGCTCCCGAAAGGCAGCTACTCTGCGCGTGGGAGGGTGGCAAGACATCCACGGATCGGTCAAAGGCCAGACGTAGGGAGCCCACGAGAACGCTTCGGGGCGGCGACGAAATTCGCCCTGAAGAGGAGGGGGCTCAAGAGATGGTTTGCGGCGAGTGATGGGGCCACATCAGTTGATGATAAAGTCACCCATATGTGAACTAATTAAGGGTTAGAAATAGCTAGAGCCTCAAAGTGGATAGCTCCACTCTAAATTCTCTCTCTCTCTCTCTTAGATATCTATGCATAGATATCAACACTCACGCATTCCCATTCATGCTTCTACTCCGAAATGGAAATATTTCTAGCGTGACTTTCTCTATCTTCTACCCGCATCCTCCGATTTAATATCTTCCTCTAACCTCCTTTAATATTCCACACTTTATAGCCCTATTTTCTCTTTTCCTCCCGTCGCGTTTCATGCCCAAGTTATTTGAATATTGCAAGCAATTTTTATGAGGGCGGTGCCCCCTCTAAAAGGGCCCAATAGGCCCAAACCCCAACTAGCTAGCTAGCCAACATAAAAGAGGCAAACCTTTTGCAAGGGAAGCCGCACCCACTGAGCTCCAAAAAAGACCCAGCATACCCAAAATGATAGTGCGGCAACCAGTTCGCGTGTGGCGGACGAGGCGTTCGGAGGAGGAAGGGAAAGAGGGGGATGAACTGACCAGAGGAAGGGCCTCCGGAAGATGGCCCTCGCCCTAGGTGAGGAGATCACGACGAAGCTCAGAGGCAGCGACATCAATTGCAAGCTGCGGGGAGAGCGGCACAAAGCTGCGGCAGGCAAAAGGGGTGCAACACGAGGACTTCCCAAGAGGTCACCCATCCTAGTACTACTCTCGCCCAAGCACGCTTAACTTCGGAGTTCTGATGGGATCCGGTGCATTAGTGCTAGTATGATCGCACCCAACATCACTAGCGCCACACGACAACTCATACCCTCGTCCCCACTCGCCAGATTGCCCCCTCTCCAACAcCACGTGGCACCGGATCATTTCCGCCCCGCTAGAGGGAGAGGGCCGGTGACATGACCGACCCCCAAGGCGGCGCGTCCCTCGCCAGGGGGAATCGTGCATAGGACGAGCAGGTAGCCCACCTCAAGTCCCCATCAACTCTCAGGTCCACTCTACGTGCCCAAGACCTGCCCAAAACGCTCCCGAAAGGCAGCTACTCTGTGCGTGGGGGGGTGGCAAGACATCCACGGATCAGTCAAGGGCCAGACGTAGGGAGCCCACGAGAACGCTTCGGGGCAGTGACGAAATTTGCCTAGAAGAGGAGGGGGCTCAAGAGATAATTTGCAGCGAGTGATGGGGCCACATCAGTTGATGATAAAGTCACCCATATGTGAACTAATTACGGGTTAGAAATAGCTAGAGCCTCAAAGTGGATAGCTCCACTTAGATATCTATGCATAGATATCAACACTTAAGCATTCCCATTCATGCATCTACTCTAAATTAAAATATTTCTAACATGACTTTCTCTATCTTCTATCCACAACCTCCGATTTAATACCTTTGTCTAACCTCCTTTAATATTCCACACTTTATAGCCCTATTTTCTCTCTTCCTCCCGTCGCATTTCATGCCCAAGTTATTTGAATATTGCAAGCAATTTTTATAGGCGGTGCCCCCTCTAAAAGGGCCCAATAGGCCCAAACCCCAACTAGCTAGCTAGCCAACATAAAAGAGGCAAACCTTTTGTAAGGGAAGCCACATCCATTGAGCTCCAAAGAAGACCCAGCATACTCAAAATGGTAGCACGGGAACTAGTTCGCATGCGGCAGACAAAGCGTTCGGAGGAGAAAGGGAAAGAGGGGGATGAACTGACTAGAGGAAAGGTCTCCAGAAGATGGCCCTC*ACCTTGGGTGAGGAG**ATATAAATAAAAAA**CTTTGAAAG**TAATAAGAAAAATT**C**AAATTTTAAAATAAAT**GAACTAATACATTG**AAAAAAAATAAAATTTTA**CAACCTTTAGTATACTTGAGTTCTCCTTTGTAAGATCTAAAAGGCTAAGTC**AAATACAAAAA**GGGGAGGTGAATTGGTGTTTCTACGATCTC**AAATTTTCTT**CTTAATCACAGATATGTCAACACGTGTACTAACATTGTTCTTCATTCTTGTTCACATGTGGGCTTTCTTTCTTTTTCCTC**TAAATTATTTT**CTCATGATTTTCTCTCACACATAAATGGGACTC**AAATATTTTTATATTA**GCTTTTCCCTCACATAACTACTTCCAATTTTAGAGGATCTTTCCTACCATATTTCGTTATTTTCCTAAGAAATACACATTGAACCTTAAGAGATCAGTGACCTAATAACTCAAG**TTTTATCTTT**GCAAAGATTTAGTCCCTTCTAAGCCAACTTGGGTTTAGAGCAACTCATGCCTTGGCCCAAAACCTATTTTCCTTCACAACTTAGTTACCTTAAAGCTTGATGAAGCTATTGATTAAGAGATTTTGATCATTTTCATTGATCATC**AATAAATAATGAA**GATGGATGAAGTAGTTTGC**ACATTTTTAAA**GCTCTACAGTGAATTTGCTTGATTGTAAGGAAGATTCTTCATAGTAGATCTTTGTGAGATCGATTTTCTTTCAATG**ATTTTCTTTATT**TGTAAGGCTTCATCCTCTAATG**ATTTTCTATTTTTTA**CTCAAG**ATAAATATAAAAGTTA**GTTCAAATTCTTACC**ATTTTTCAAA**CTGTGTCATAACGAC**TTTTGATTATTA**CG**AATTCAATAAATTTTTTATTCTTTT**CTTC**ATTACATTT**G**AAATTTGAATT**GGTGATTCAGGTTGATGTTGACTTCAAGTAAGAATG**TTAAAATTGTTTT**CTCC**AAATAAATA**GCCAATATGTCATTGTCC**AAATTTTTTGTT**CGAG**ACTTATAAAA**GC**TTATTTAATATTTTAAA**CTCTTTTC**ATTTTATTTTCATAATTTAT**GTGACTAATAC**TTAACTAATT**ACACTTAAATGG**TTAAATAATCATAGATAAATTAATCAAAATAATAA**CCCCTCATTTACCTTATATCATTTACCC**ATGTAAATATTTATTTTCTAAAAAATATTTATTCTTATAAATTT**GACTCTTCACT**TATTTATCAAATTATTTAA**TCTCAATGTCTAATCCC**TTAGATTAATA**GGTCATGCATAATATCAC**AAGATAAATAATTA**GGTATAATACATCAAAGATACCAAGCATTTGTAATGATCTCGTGGATGATGTAAAATCTTGAATC**ATATATTTTCT**CG**TTAAATATAATATCAAAA**CACATGCCCACAGAC**ATCAACAATTT**CTCC**TTTTTTGATA**C**TTACAAAATTAT**GTTAAGA**AACTATAT**C**TATTTGTATTAATAT**GTGCAC**AAAGAAAATAATAAT**GGATTCTCACACATAAG**AAATCAAATTT**TGGCAATAACAATCGTAAGCATCATGTGTCAAGACGTGTCAAAGCAATAACACTCAAATAG**AATATTTAAAT**CCATTCACATAAGTCAACCATGAAGGTTCAAG**TAATAAAGAT**GTCAATGCTCTGAATAC**ATAATTTCTT**CCTCTAATTTTGTAC**AATCATATAAATTTT**CAACATTTATCAATCTTTGTCCCCC**TTTTTCTAAGATAAAAAAATAAAATATTAATCAAT**GGGGAGAAGCAC**ACAAAAAATA**GCCTCAACTCTAGTAAGAACAAAATCAAGAAAAACACGAATGAATTTCTGAAATATTGTCTTGAAGGTCATGAAGTTGTTCTTCCATCATGTCGAGGAGTTGCTTCATGTGTTTCATGC**TTTTATTAAATTTCTCAATAATATGAAT**GATGTCGAGGAAGGTGAGAGCACAAGATGATGTATGAGAGGTGGAAGATATATGATGAAGCTATTGACCATCAATGTGAG**AATAATTCATT**CTATGAAGAATGG**ATTTTTCAAT**GATGATATATGG**AATACTATTAATA**GATTCCTCTCTATAGATATCTACTTGATAATACTTGAGGGCACACATGATGACAAACCCAAGAGGAAGAATCAAGTCATATTTCAAGGC**TATTTTCATAT**G**TTATAAGAATA**GATCACCAAAATTAAGTGAAATGCCATGAATTATGCAATATAAGATAGACATAGATTTTCTAACAATGTATCATAAGAGATGACCAACAAGAAAATGTATGTCATGATGAGATAGTAG**AAGATTAAATA**CTTTCATGTTATACACATGCATTTTCCAATCTAGTCAAATTGATCAAAAAGGTATTGATGGTTTCTTTGGTGATGGTTAAGGGAGGTTTAGATTGAATATC**AAAAATGTAATTTAAGATTTTA**GGAGTAAGATCAATGAACACTCC**ATTGATATATA**TGGTGAGGCTGATAAGGGTGAC**TTTGAAATTATTATA**GAACATCT**CAATAAGATTTAAAT**GGTTATCAACATGTTC**AAAAACTAAA**TGTGTAAGTCCAAGGGCATCAAAGTATTTC**TTTATATTGATTT**GTGGTGAC**TTAAAGAATTTTA**GGTTAATACAC**TTGAAATAAA**GGACTTTATAGTC**TTTAAATGATTTT**GG**AATTTGATTT**CATGAATTGGCTTGAGGTGTAGC**ATTTTTTCTA**GGGTGTTTGGGATTCTTTTCTTAGCCTTGATGTCCTACAAGGTAATTTCTTCTATAGGTTTCATTTCACACATAGGAGACAATCTTTTGA**TAATAGATTTAA**CCCTTCTCTTGAGCTGCAAAGGTTTTGACAATTTTGGATTATTGGTAGGTGGATCGGGGATAGGG**TTTTTTAATTCAA**GTTTTGTAGGAATAGATGAGTTAATGGGTTTG**ATATTTTCAT**GTTCAATGAATAGTAAGAGAGGTGCTGGTGTCC**ATAAATAGTTT**TGAATGATTGTACCAACTTGTTTGTCTATTTTCAATGGTAATAGACATAGGTGATCCATTTTGGGCAGAAGATCTACATAGGGGGGGGGGGGGGGGATAGGAAATAGTGTAGGTAGTTTTCTTTTCATGAGAGATGATGGGGGGAGTCCATTCAAGAGATGTGTCAATGATCATATCCTTTCCTTTATCTTTCTTCG**AGATTTTTTA**GTGTCTATAGGCTCTTGAAGGAGGGACATAAAGGGTAGAACATCAAGTGTCAACCATCATAAAACTAGGATTTAGAAAGGGAGAAATTGACTAGAAGGAGATGTGACTAGAATGCAATGAAAGAAACACAATACCTCATGTTAACTCAAGAGTC**TGATAATTATAAATTA**GTAGTTACATC**TATGTATTAATATCTTATATAA**GG**ATTTGAATTAAT**GGATAGATATAATTATTGATAATTATAAATTAATAGTCATGTATGTGC**ATTAATATGTTATATATA**G**ATACTAATTA**GTGGGTAG**ATAATGTAATTATA**CGTGTTACGAACTCATTATGAC**ATTTATAACATT**CTACCATTTAAGTCCTAATACAC**ATTAATTTACTTATTATATTAATATTAT**GAGACTAATCCCACTTGACTTTTATAAAGTCAAGTGGCGGTTCTCTCACATTTTCTTG**ATAATGATTTAATTATAA**GGACTTAGGTGAAGAATGAGAGAAGCCCGAGTTCTTAGCCCAAGGAAAAGCCCATCTTACAAAGAGGATCGGCCTAAGACATTTTTGGGCCAAAGAGAGGGTGTCATCTTGGTCAAGCCCATTTGGACCTTAAGTAGCACATGACCTTGGCCCAGCCC**ATTTAAATTTAAAA**GGGCTATC**AAGAAATAAAAT**CTAGGCTGTCTACCAAGAGAGTCGGTTAC**TAATCATTATT**GTCCATTAGTGGGGGTTAGTTTTGTAGAGCTTTAATGCCTCTCATCTCCCTCTTGAAGGATGCTTTTAGCTCTCTCTCTCTCTAGCAGTAGCTAGCCAAAAGAAAAGGAGGGAAATCAGGGCCTAGTCAATGTTATTCTTGTGCTCAAGATCGATATGGAGAAGAGATTACTTGGAACACATC**AAAATTATCA**CGAATCAAGCACACTCTGCCTAGGACTAGCACACAATAAGGTTTGTGCAAAATCTAGATTTAAGGTGTCCTTATAATCCTTTCTATTGATAGATTAACTCTTAGTTTGAACTAGTTTC**ATTTTTTATTCTAT**C**ATTATATTGT**GTTCTGCCTATTCTTTCATCATCATCACCTTCCCATTCCTCTTATCTTCACATTATCAC**ATTTAATTTCA**CCCATCACCATCTC**TATTATATATTTGTATATATAAGAAATAA**C**ATAAATTCATA**CTCTAATGTTTACGTTCCCTGAGGAGTGACCCTTACTC**ATTTTAAATAGA**GGAATGAGG**TTTTATAAATATATTT**GCATGAACTTGATTGCACTC**ATAATGAAAAATTAA**CTTC**TTAATCAAATTAAT**CAATTATACATGTATTAACGACCCATTATGACATTC**ATAATATTTTA**CCGTTTAAGTC**ATAATGTATATTATTTTATTTATTTTATTAATATTT**GAG**AATTATAAATTAATAGTTTTATTTATGTATTAATATTTTATATATAT**G**TTAATTAGTA**GATAAAGAATG**TAATTATACAT**GTTCTATCGACTC**ATTATAATATTTATAATGTTTTATCATTTAA**GTCGTAATG**TATATTATTTTACTTATTTTATTGATATTATTAA**GTCC**TAATATATTATGA**GACTAACCCCCCGTG**ACTTTTATGAA**GTCAAGTGGTGATTCTCTCAC**ATTTTTTTGATAATTATTT**GATTGCAAGGATTTAGGTGAGCTCTTATTATGAGTTGAAAGTAATGGAAGAAGCCTAAGTTCTCAACCCATGGAAAAGCCCAAATCTTATGGAGAGAGTTGACC**TCATATATTTAT**GGGCCAAGGCATCATCTTGGTCAGGCCTATTTAAACCTTAAGTGGCATATAACCTTGGTCTAACCAATTTGGACTTAAAAGGGCTATCAAGAG**ATGAAAAATA**GGACACCCATCTTAAGGTACCTTAAGATGAGATTAAG**AAGAAGATTA**GAGCACTAGCCTCATAGAAGAATCTAGGTCGTCCACCAAAAGAGTTGG**TTATTAATATAACTATTAATTA**GAGGAGCCTAATTCTCTAGGGGTTTAAGCCTCTTGCCCCCCCTTTTAAAG**AATACTTTTT**GGCTCTCTCTAGTAGCAGCTAGCTAAAGGAAAAAGAGGGAAATCAAGGCCCAGTCAATGACATTCTCGTGATCAAGGTTATTGTGGATAAGAGACTACTTGAAGTGCATCAAGATCATCATAGATCAAACATCACTTGGCTATGACTGGCACATAACAAGGTTTGTGCAAGATCTAGATTTGGGATGTCCTTGTAATCCTCTCTATAGATAGATTAACTC**TTATTTTGAA**CTAGTTTG**ATTATTTATTCTAT**CATTCTATTGTG**TTCTATTTTTTTTT**CCATCATCATCACCTTCTCCATCCCTCTTATCTTTATC**ATTATCATATTT**GATTTTGCCCATCACCATCTTAAG**TATATCTATATATATATAT**C**TAAGAAATAAATTAAATAAAATTCTTA**CTATCATTATTCACATTCCTTGATGATCGACCTTTACTCATC**TTATATTGAA**GAGTGAGG**TTTATAAGTATTATTT**GCGTCAACTTGGTTACATCACG**ATGATATATTTAA**CCTGTAAATCAAG**TTCATTAATATTAT**GAGACTAACCCCACTTGACTTTTATGAAGTCAAGTAGTAATCCTCTCACATTTTCTTG**ATAATGATTT**GATTGCAAGGATTTATGTGAAC**TTTTATTATGA**GTTGGAAGTAAATGGAGAAGCTCAAGTTCTTAGC**TAATAGATTTTA**GTC**ATTTTCAAATTTTATATTTATAAA**CCTTATCCAACTATATGTATATCTCAAATATATAGTGGATTCAAGGTCAAACATAGGGACAACTATGTGAGTAGATGATCACTATTG**ATTGATAAAT**CTATTCAAGTTTGGTGTTGG**TTCTAAATTATGATAATAAAAGAATAAA**CATAGATCTCTTGTTGGCGTTGGAAGAAATTGAGTCCATATTGAAAGGAGAAATCCTCTCAAAGTCTCATCAGCCACCACATAAAAGAGTTATGGATGTCTAATGAGATGGATCCTCAAATATGTTGGCACATTCAAATGATCAAATCACTGGTATGAGGAAAGTCAACAC**ATATCAAATT**GC**AAAATTAATTAAA**CAGAAATCACAAATTCTAGTTTGACAATTGAAAGTGAATACAATAAGTTGATTATTCTCCCTCTTCTTTTGATGAACATAGGGGCTTGAGTGGTTGATCCTTGTCTTATTGCACCAAGAAAGGAGATCCACGATCTTCACCGATGTTGTCACCACATGAGGAAGAGGAGAAGGAAGAGGAGGCCCGCCTATGGATTGCAGACAGCACCAGGGAAAAAGAGGTGATGAGATGGCCAACTCTGTAGTGGAGTAGAGTCACACACCCAAGGGAGAGAGACGGATGTCTGATTGTGAGTGCCCTCTTGAGTGGCAACTACAATCCTTTTGGGATTGTGTACCCTATCAGGCACCTTCCAAGAATGCCCTTGCTCCCTGCAAGAATTACAAG**TAAATAATGTTTATAATTTTATGTTTTATAT**CCA**AGTATTATTAAAAA**GCCCATGATAAC**ATTAAATCAA**GATATATGTGCATGATCATGAATTGTTCTGCTTCCACTCAAGATCTAGATTTGAGCTTCTTTGTAATTCTCTCC**ATATTTCAAATT**GTGTTTATCATTTATTCTATCATTCTATCGTACTCCATCCATTCTCTCCTCATTGCCACTTTCTCCATCCCTCTTATCTTCATCATGATC**ACATATAATTTT**GTCCATCACCATC**TTAAGTATATT**G**ATGTATATATA**G**AAAATAATGT**GAGC**AAAATTTATACA**CTCATGGACCCTTATATCCTAAATTGAAAGAGTGAGG**TCTTATAAATATTATTT**GCATGAACTTGGTTGTACTCACCATGATCATTTAACCACTAAGTCGAGTACATAGGAGTCAACCTAATATGATGCGAGCTTCAACTTGAAGGCTTACAAGTAGAGGTGAAATAAAGGTTTTTGGATGGTGAAAGAGAGAGAAAGGTAAAGTAATTG**AAAGTATTT**GAACTTGTGAAGTTTAACCTTTTGATAGAAGAAGAGAGACATAGACAC**AATATAGTATTT**CTTAAGATATGCGACAACATATCATGGGTTGCTCATGTCC**TTAATAAAAATCAA**CTAAATAGAAG**AATTTCAAAT**CACAATATCATCATCAATTTGTACATTCCAAG**TTTAGAAATAATT**G**AATTAAATCT**GTCTTTAGAAAGGG**ATTTTGTAAAAATATCAAATAATTATTTTGATATAT**C**AATAATGTAT**GATTTTATGTCTCC**TTTTTATATGT**G**ATTATGAATAAAAT**GATGTATGATTTCAATGTGCTTAGTTCTAGAATGTATG**ATAAGATTTTTA**GTCATAATG**ATATCATTAT**GTTGTCACAATACATTGGTATCTTCTCTAATTTAAGTCCATAGTC**TTTTAATTGATATTTTATCTAAATTATTT**GTGC**ATAATAAGTT**CTAGTAGACACATACTTGGCCTCTATAGTGGATAATGC**AATAGAATTTTATTT**CTTGAAACTCCAAAAAATAAGCAACC**ATTTAAAAATAAACAT**GACCTTGATAATCTCAAATTAGGACTTGTATAG**TCATTATTTT**GCTTTAAGCTTATACC**AAAATATTTATACTTTTT**CC**TTTTGATTTA**CTC**TATTTTTAGAATAA**GGTAAGTAGGGGTCACTCC**TTAAGTAATTAT**G**ATTGAAATTAT**G**ATTCTTTTAA**GG**TTATAAATGAA**GAAGAAAGATTAGAGAGAAAAGATTGAGAAGAAGGAGGAATGAGAATCTTGATTTTCAAGATAGCTAGTTATCTCAAGAAAATCAAGATTCAGATTTCAG**AATTTAATATTTTGTAT**G**ACTATATTTA**GAACTTAGGGGAATGATATGAGAGAAGAAAAGATGATGAAAGGTAAGAGGATCAAGGAGATCTTGATTTTTCCTATG**AAAATCAATAAAT**CCAAGGTCTCAATGGGAAAGG**TTTGAATTAT**GG**TTTTGATTTT**GAGAAGTAAGAAGAGAGGATGAG**AAAACTAATTTTT**CTATCGAATATTAGAGAACGAAATCACTAATTTCAAGACCTAGGAGGGCATACCTTGAGCC**TATATACTTT**GCCCAAATTAGGTGGTTTTCTCCAACAAGAACACACTTCAAAGATC**AAAAACAAA**CTTTGCAATAATTCATAATGCAGGACAAATGAGACCTTTAAGTATGAATCCTTCAAGAGTAGTGATTGTACTTAAGAGTTGAATCAAGTAAGG**TTTAATAAAATA**CGAGATCCCACAATTTGAAAGATGGATCATC**TTAATTATTGTAATA**GGGAATTCTACTTCAAAGAGTTAG**AAATAATATAACAACTATAT**GTATGGATCACCAAATCAATGAGAGAGGGCTAATCTATGATCACTCACACTAAGAGCATGTGG**ATTATTTTAGAATT**GACCAAATATC**ATGAAATTTAA**GAAGTCAATGTACCTTAATGGAGATCC**TTTTTATAAAGA**GTTTTTGTGATTTCTGAAGGTGGCTCTCTTTGATAAAACTCCCACATTTGGGAAGAGGGTGAAAGTCTAGAAAGGAAAGAGAGATAGATAAGGAGGTACCATAGGAGAAACAAAGTCAGAGAGAGAGTGAAACCGATGGAGAGAGGAGAGAGAGGGAGAGAGAAATAGAGAATAAATGGGAAGATTGAAAGGAAAGGACGTGAAGAAAATCGTTAGAGAAATAGAGGACAACTAGGAGAGCGTGAGCAGATGATAGAGGGCGTGAGAGAGAGAGGATGACAAGGGAAAGGAGAAAGAGAGGGAAATGACACACAGAGAAGAGAGAGAAAGAGGAAACCGTCCACAAGAGAGAGAGAAAGAGAGGGAACGACGTGCAGAAGAGAGAGAGAGAGGGCAACATGCAAGGGAGAGAGAGAGAGAAGACGATGCACGGGATGGAGAGGCCTACCTAGAGGGGGAAAGAGAAAGAGGGGGTAATGCAGGAAAGAGAGCAAGAGTGATGGGGATCAGGAACTGACTGTTCATGGGACAGTACGACAAACGGACTCTGGTTCCATTTATCCAGGCAGTGCAACGGCTGAGCGTGCAGAGGGAGATGCCCTGTCGCTTGACACCATTGCCCCCATAGACAGACAACCACTACCCCTAGGGGACCCAGACTGGTAAAGTCGACGATGCCTTTGTCTATTATTAGCTGTTCGGCGACGTCGTCGTCGACCACTGGTGCTTTCTATTTCATTTCTGTATGTAAGTCGTGCTTGTTTAAAAACCCCCCCTCAGGACGAGGTGAGGGGGTAATCTGAGTCTTGAAGTTTTGGTTCCCTCTAGAGTTCGGCTCTGCCGTGATGGCATCCTTGTGAGCATTCATTGCCATTTCCTCGCTACTTACCTCAACACCCTCACTCCCCCAGCTTGCCCAACAAAGAGAGAAAGAGGGGAAAGGTGAGGAAGGCCGTGGGAAAGAGATAGAAAGAGAGAGAGGGGATGCATGAAGAAGGGATAGAAAGAAAGAGCGCGTGGGAGTGGAACACATGAGGGAGAGAAAGGAAGAGAGAGAATGCACAAGTGAGAAACAGGCGAGAAAGAGGAAAGAGAGAGAACACACAAGTGAGAAACACGTGAGGAAGGAAGAGGAAAGAGAGAGAACACGCAAGTGAGAAACACACTGAGGAAGGAAGAGAAAGAGAGAGAAGATAGAAGAGAtAATC**TATAAACAAA**GGATGCCTTTGGTGCCAAGTGGCACCCCTTCAAAGGATGCCATGAGATGGTTCCCATTGGTGCAATGTGGTGCCAATTCATTGGTCCCTAGTGGCATTGAGTGTCCTCTGGCACAGCCTTCGTAGCTTATTTATTGGTGCCCAGTG**AATTAATAGA**GAAAATATTGACTGATTGTCCCCATCAATTCATCTCTATTC**AAATTTATCAATTT**GGATCCTGGGAGAAC**TTTTTCTTATT**GTCAACATTTGTTGGGCAATCTCTAGAATGAGAAACAAGG**AAATAAAAGA**GGTTTTATACGTAGTGACATATTTCTCAAAAGAACCCTCTCTTTTCTCCATTATCTTTCCATGCCCTCATCTCTAAATCATCTCTTCTTCTAGGCCAATATCCTTCAAACTC**TTTAATTAATTACTT**GATTAAATTCACCTAAATCTTGCACACTATACACTAGCAATATTCCATAGTAACC**ATAAATCATAT**GTATATCAC**TAATTTTTCATA**GGC**AAAAATAAATATAAAGTAATTT**CTCAC**ATTCATTTATTATTTTAAT**GTTGCTAGCATGATATTAGAGATCTTCAAAGCTCATAAGC**ATATAAAATCAT**AGATATTACC**ATTATATTTATCAT**GTTATC**AATAAGTTAT**CAAACCTCACTAGTGCTTTTTC**TATCAATTTTATAA**CCTACAGAATTAGAATCATAATATC**TTAAAAGATTTA**CATCTTGATGC**TTTAAATATTATATACATAA**GGCTGTTATATCAATCAG**ACATATAAATATTTTTTT**GACATCGTGAGGTGTGACTCTTTAGGGTTGATTTGAAACCTAAAGAGTCATTCC**TAAATATTATATTAAATAT**CAGATCTACTTATTGTC**AAATATGATAAA**GACCTAATCATTCCTCTATCTAACTTCTTACC**AATTTTTTTTCCTTTTTTATTTAT**CAAGTTTGACATTTGTTCCCATAGGAG**TTATTTTTGAAAT**CTAAATTATTTATTTTAAATTTCTTAACTAACTCTTTTGTATATTTTTCTTGAGATATGAAATTATTTCATACTCATTTCAAATTCAAAATTAAATATG**ATAAATCATT**GCATAGAGATTCATCAC**TTAAATTGAAAATAATAT**CATG**AAACATATATTT**CAACAATTAACACATGTATTTCTACTCTTTTAAG**AAATAAAGTT**GTGTCCACATTTCATTTGTC**AAAATTATTTT**CTG**AAAGAAATAAA**CTCAATTCCTCATACCAAGCTCATAGAGCTTGTTTC**AAATCATATAAA**CTC**TTTTTTAATCTAAAA**GTATG**ATTAAGAAATTTAGAAATTTTAAAAT**CAAGAGTCTATTCAACATAAAC**TTCTTTAATAAAA**CC**ATTTAGTAAATTA**CCTTTAATATCTATC**TAGTATACTTTAAATTATTA**GCACATGC**AAAAAATAACATAATTATAATA**GCATCAAGTCTAGAATATAAGCATAAGTTTTCTCATAATCTATCATCTTATTAGCCTTGAACAACTAATCATATTTCACTCCTAATTACTTCTCATTGTTCATTTAAGCTATTTCTAAATACCCACTTTATTCCTCTAATATTTTGATGTTTAAGCTATGTAACTAGATACTAGACACTATTTATTTCAAATTCTTCAAGTTCTTCATACATAATAATGATTCAAATTGTATCTAACAAATCATCATC**AATATTTTTTAGTT**CAATTTGAGAAGTAAATGTCTTGTTGATAATATCTTCATTTTTTAGATGTAATGTGATCG**AAAATAATAATA**CGTGGCATGAATTTCATC**AAATACTTATTTTTT**GTGAATTTTAGTTAACTTTTATCATTTATCTACGATTAAAATAGATTACCTAC**TATTTTATTAAATAAATAAATAAATAAATAATATAT**CTATTATCGAGCATAGCCATTTGATTTCTGTTTGATCTGATCCCTCTCAACAAGCCTCATGACTCATTTTCTAGTAGCATATCAGCGGTGGTTAAATACCAATAACAAGCTTCTTATTTCTAACTTTTCAGAATGTACCATGTACCATTAAAATTTAATTAAATATAACAGATCATGGTGCAGTAATCTCACAACATCTAGACTGCCATGCGCCCCCAATGCATAAGTGATTGATTGTATTACATGAGATGTAATGATCAGATAGTAAAGAATGTGTCGAACAATCAATGCACTTATGTTCCAAAGAAGC**ATATAAACAAATATA**GATTACAGACAACCAAAACC**ATAATTATCAATTATTTT**CCATTCAATAAATCAACAAAGCTTCAACGATAATCAGCAATCCCTTATCTCCCTCTATCCTGCGATCAAGGCAGTTTGCAACCACCCCACACTCCTAAATCCCTTTGATAC**AAAAATAAAAATT**GTTCAAAGTAAAAACTCCATAACTTTGCAACCACAACTCATATCAACCAATAAAGCCATTCGTTACGAAGCTAACAAAAAACGCAAGCTAACTAT*CTTTCGGGAGCGTTTTGGGCAGGTCTTGGGCACGTAGAGTGGACCTGAGAGTTGATGGGGACTTGAGGTGGGCTACCTGCTCGTCCTATGCACGATTCCCCCTGGCGAGGGACGCGCCGCCTTGGGGGTCGGTCATGTCACCGGCCCTCTCCCTCTAGCGGGGCGGAAATGATCCGGTGCCACGTGGTGTTGGAGAGGGGGCAATCTGGCGAGTGGGGACGAGGGTATGAGGTGGTGTGTGGCGCTAGTGATGTTGGGTGCGATCATACCAGCACTAATGCACCGGATCCCATCAGAACTCCGAAGTTAAGCGTGCTTGGGCGAGAGTAGTACTAGGATGGGTGACCTCCTGGGAAGTCCTCGTGTTGCACCCCTTTTGCCTGCCGCTCTCCCCGCAGCTTGCAATTGATGTCGCTGCCTCTGAGCTTCGTCGTGATCTCCTCACCTAGGGCGAGGGCCATCTTCCGGAGGCCCTTCCTCTGGTCAGTTCATCCCCCTCTTTCCCTTCCTCCTCCGAACGCCCCGTCCGCCACACGCGAACTGGTTGCCGCACTACCATTTTGGGTATGCTGGGTCTTTTTTGGAGCTCAGTGGGTGCGGCTTCCCTTGCAAAAGGTTTGCCTCTTTTATGTTGGCTAGCTAGCTAGTTGGGGTTTGGGCCTATTGGGCCCTTTTAGAGGGGGCACCGCCCTCATAAAAATTGCTTGCAATATTCAAATAACTTGGGCATGAAACGCGACGGGAGGAAAAGAGAAAATAGGGCTATAAAGTGTGGAATATTAAAGGAGGTTAGAGGAAGATATTAAATCGGAGGATGCGGATAGAAGATAGAGAAAGTCACGCTAGAAATATTTCCATTTCGGAGTAGAAGCATGAATGGGAATGCGTGAGTGTTGATATCTATGCATAGATATCTAAGAGAGAGAGAGAGAGAATTTAGAGTGGAGCTATCCACTTTGAGGCTCTAGCTATTTCTAACCCTTAATTAGTTCACATATGGGTGACTTTATCATCAACTGATGTGGCCCCATCACTCGCCGCAAACCATCTCTTGAGCCCCCTCCTCTTCAGGGCGAATTTCGTCGCCGCCCCGAAGCGTTCTCGTGGGCTCCCTACGTCTGGCCTTTGACCGATCCGTGGATGTCTTGCCACCCTCCCACGCGCAGAGTAGCTGCCTTTCGGGAGCGTTTTGGGCAGGTCTTGGGCACGTAGAGTGGACCTGAGAGTTGATGGGGACTTGAGGTGGGCTACCTGCTCGTCCTATGCACGATTCCCCCTGGCGAGGGACGCGCCGCCTTGGGGGTCGGTCATGTCACCGGCCCTCTCCCTCTAGCGGGGCGGAAATGATCCGGTGCCACGTGGTGTTGGAGAGGGGGCAATCTGGCGAGTGGGGACGAGGGTATGAGGTGGTGTGTGGCGCTAGTGATGTTGGGTGCGATCATACCAGCACTAATGCACCGGATCCCATCAGAACTCCGAAGTTAAGCGTGCTTGGGCGAGAGTAGTACTAGGATGGGTGACCTCCTGGGAAGTCCTCGTGTTGCACCCCTTTTGCCTGCCGCTCTCCCCGCAGCTTGCAATTGATGTCGCTGCCTCTGAGCTTCGTCGTGATCTCCTCACCTAGGGCGGGGGCCATCTTCCGGAGGCCCTTCCTCTGGTCAGTTCATCCCCCTCTTTCCCTTCCTCCTCCGAACGCCCCGTCCGCCACACGCGAACTGGTTGCCGCACTACCATTTTGGGTCTGCTGGGTCTTTTTTGGAGCTCAGTGGGTGCGGCTTCCCTTGCAAAAGGTTTGCCTCTTTTATGTTGGCTAGCTAGCTAGTTGGGGTTTGGGCCTATTGGGCCCTTTTAGAGGGGGCACCGCCCTCATAAAAATTGCTTGCAATATTCAAATAACTTGGGCATGAAACGCGACGGGAGGAAAAGAGAAAATAGGGCTATAAAGTGTGGAATATTAAAGGAGGTTAGAGGAAGATATTAAATCGGAGGATGCGGATAGAAGATAGAGAAAGTCACGCTAGAAATATTTCCATTTCGGAGTAGAAGCATGAATGGGAATGCGTGAGTGTTGATATCTATGCATAGATATCTAAGAGAGAGAGAGAGAGAATTTAGAGTGGAGCTATCCACTTTGAGGCTCTAGCTATTTCTAACCCTTAATTAGTTCACATATGGGTGACTTTATCATCAACTGATGTGGCCCCATCACTCGCCGCAAACCATCTCTTGAGCCCCCTCCTCTTCAGGGCGAATTTCGTCGCCGCCCCGAAGCGTTCTCGTGGGCTCCCTACGTCTGGCCTTTGACCGATCCGTGGATGTCTTGCCACCCTCCCACGCGCAGAGTAGCTGCCTTTCGGGAGCGTTTTGGGCAGGTCTTGGGCACGTAGAGTGGACCTGAGAGTTGATGGGGACTTGAGGTGGGCTACCTGCTCGTCCTATGCACGATTCCCCCTGGCGAGGGACGCGCCGCCTTGGGGGTCGGTCATGTCACCGGCCCTCTCCCTCTAGCGGGGCGGAAATGATCCGGTGCCACGTGGTGTTGGAGAGGGGGCAATCTGGCGAGTGGGGACGAGGGTATGAGGTGGTGTGTGGCGCTAGTGATGTTGGGTGCGATCATACCAGCACTAATGCACCGGATCCCATCAGAACTCCGAAGTTAAGCGTGCTTGGGCGAGAGTAGTACTAGGATGGGTGACCTCCTGGGAAGTCCTCGTGTTGCACCCCTTTTGCCTGCCGCTCTCCCCGCAGCTTGCAATTGATGTCGCTGCCTCTGAGCTTCGTCGTGATCTCCTCACCTAGGGCGAGGGCCATCTTCCGGAGGCCCTTCCTCTGGTCAGTTCATCCCCCTCTTTCCCTTCCTCCTCCGAACGCCCCGTCCGCCACGCGCGAACTGGTTGCCGCACTACCATTTTGGGTATGCTGGGTCTTTTTTGGAGCTCAGTGGGTGCGGCTTCCCTTGCAAAAGGTTTGCCTCTTTTATGTTGGCTAGCTAGCTAGTTGGGGTTTGGGCCTATTGGGCCCTTTTAGAGGGGGCACCGCCCTCATAAAAATTGCTTGCAATATTCAAATAACTTGGGCATGAAACGCGACGGGAGGAAAGGAGAAAATAGGGCTATAAAGTGTGGAATATTAAAGGAGGTTAGAGGAAGATATTAAATCGGAGGATGCGGATAGAAGATAGAGAAAGTCACGCTAGAAATATTTCCATTTCGGAGTAGAAGCATGAGTGGGAATGCGTGAGTGTTGATATCTATGCATAGATATCTAAGAGAGAGAGAGAGAGAATTTAGAGTGGAGCTATCCACTTTGAGGCTCTAGCTATTTCTAACCCTTAATTAGTTCACATATGGGTGACTTTATCATCAACTGATGTGGCCCCATCACTCGCCGCAAACCATCTCTTGAGCCCCCTCCTCTTCAGGGCGAATTTCGTCGCCGCCCCGAAGCGTTCTCGTGGGCTCCCTACGTCTGGCCTTTGACCGATCCGTGGATGTCTTGCCACCCTCCCACGCGCAGAGTAGCTGCCTTTCGGGAGCGTTTTGGGCAGGTCTTGGGCACGTAGAGTGGACCTGAGAGTTGATGGGGACTTGAGGTGGGCTACCTGCTCGTCCTATGCACGATTCCCCCTGGCGAGGGACGCGCCGCCTTGGGGGTCGGTCATGTCACCGGCCCTCTCCCTCTAGCGGGGCGGAAATGATCCGGTGCCACGTGGTGTTGGAGAGGGGGCAATCTGGCGAGTGGGGACGAGGGTATGAGGTGGTGTGTGGCGCTAGTGATGTTGGGTGCGATCATACCAGCACTAATGCACCGGATCCCATCAGAACTCCGAAGTTAAGCGTGCTTGGGCGAGAGTAGTACTAGGATGGGTGACCTCCTGGGAAGTCCTCGTGTTGCACCCCTTTTGCCTGCCGCTCTCCCCGCAGCTTGCAATTGATGTCGCTGCCTCTGAGCTTCGTCGTGATCTCCTCACCTAGGGCGAGGGCCATCTTCCGGAGGCCCTTCCTCTGGTCAGTTCATCCCCCTCTTTCCCTTCCTCCTCCGAACGCCCCGTCCGCCACACGCGAACTGGTTGCCGCACTACCATTTTGGGTATGCTGGGTCTTTTTTGGAGCTCAGTGGGTGCGGCTTCCCTTGCAAAAGGTTTGCCTCTTTTATGTTGGCTAGCTAGCTAGTTGGGGTTTGGGCCTATTGGGCCCTTTTAGAGGGGGCACCGCCCTCATAAAAATTGCTTGCAATATTCAAATAACTTGGGCATGAAACGCGACGGGAGGAAAAGAGAAAATAGGGCTATAAAGTGTGGAATATTAAAGGAGGTTAGAGGAAGATATTAAATCGGAGGATGCGGATAGAAGATAGAGAAAGTCACGCTAGAAATATTTCCATTTCGGAGTAGAAGCATGAATGGGAATGCGTGAGTGTTGATATCTATGCATAGATATCTAAGAGAGAGAGAGAGAGAATTTAGAGTGGAGCTATCCACTTTGAGGCTCTAGCTATTTCTAACCCTTAATTAGTTCACATATGGGTGACTTTATCATCAACTGATGTGGCCCCATCACTCGCCGCAAACCATCTCTTGAGCCCCCTCCTCTTCAGGGCGAATTTCGTCGCCGCCCCGAAGCGTTCTCGTGGGCTCCCTACGTCTGGCCTTTGACCGATCCGTGGATGTCTTGCCACCCTCCCACGCGCAGAGTAGCTGCCTTTCGGGAGCGTTTTGGGCAGGTCTTGGGCACGTAGAGTGGACCTGAGAGTTGATGGGGACTTGAGGTGGGCTACCTGCTCGTCCTATGCACGATTCCCCCTGGCGAGGGACGCGCCGCCTTGGGGGTCGGTCATGTCACCGGCCCTCTCCCTCTAGCGGGGCGGAAATGATCCGGTGCCACGTGGTGTTGGAGAGGGGGCAATCTGGCGAGTGGGGACGAGGGTATGAGGTGGTGTGTGGCGCTAGTGATGTTGGGTGCGATCATACCAGCACTAATGCACCGGATCCCATCAGAACTCCGAAGTTAAGCGTGCTTGGGCGAGAGTAGTACTAGGATGGGTGACCTCCTGGGAAGTCCTCGTGTTGCACCCCTTTTGCCTGCCGCTCTCCCCGCAGCTTGCAATTGATGTCGCTGCCTCTGAGCTTCGTCGTGATCTCCTCACCTAGGGCGAGGGCCATCTTCCGGAGGCCCTTCCTCTGGTCAGTTCATCCCCCTCTTTCCCTTCCTCCTCCGAACGCCCCGTCCGCCACACGCGAACTGGTTGCCGCACTACCATTTTGGGTATGCTGGGTCTTTTTTGGAGCTCAGTGGGTGCGGCTTCCCTTGCAAAAGGTTTGCCTCTTTTATGTTGGCTAGCTAGCTAGTTGGGGTTTGGGCCTATTGGGCCCTTTTAGAGGGGGCACCGCCCTCATAAAAATTGCTTGCAATATTCAAATAACTTGGGCATGAAACGCGACGGGAGGAAAAGAGAAAATAGGGCTATAAAGTGTGGAATATTAAAGGAGGTTAGAGGAAGATATTAAATCGGAGGATGCGGATAGAAGATAGAGAAAGTCACGCTAGAAATATTTCCATTTCGGAGTAGAAGCATGAATGGGAATGCGTGAGTGTTGATATCTATGCATAGATATCTAAGAGAGAGAGAGAGAGAATTTAGAGTGGAGCTATCCACTTTGAGGCTCTAGCTATTTCTAACCCTTAATTAGTTCACATATGGGTGACTTTATCATCAACTGATGTGGCCCCATCACTCGCCGCAAACCATCTCTTGAGCCCCCTCCTCTTCAGGGCGAATTTCGTCGCCGCCCCGAAGCGTTCTCGTGGGCTCCCTACGTCTGGCCTTTGACCGATCCGTGGATGTCTTGCCACCCTCCCACGCGCAGAGTAGCTGCCTTTCGGGAGCGTTTTGGGCAGGTCTTGGGCACGTAGAGTGGACCTGAGAGTTGATGGGGACTTGAGGTGGGCTACCTGCTCGTCCTATGCACGATTCCCCCTGGCGAGGGACGCGCCGCCTTGGGGGTCGGTCATGTCACCGGCCCTCTCCCTCTAGCGGGGCGGAAATGATCCGGTGCCACGTGGTGTTGGAGAGGGGGCAATCTGGCGAGTGGGGACGAGGGTATGAGGTGGTGTGTGGCGCTAGTGATGTTGGGTGCGATCATACCAGCACTAATGCACCGGATCCCATCAGAACTCCGAAGTTAAGCGTGCTTGGGCGAGAGTAGTACTAGGATGGGTGACCTCCTGGGAAGTCCTCGTGTTGCACCCCTTTTGCCTGCCGCTCTCCCCGCAGCTTGCAATTGATGTCGCTGCCTCTGAGCTTCGTCGTGATCTCCTCACCTAGGGCGAGGGCCATCTTCCGGAGGCCCTTCCTCTGGTCAGTTCATCCCCCTCTTTCCCTTCCTCCTCCGAACGCCCCGTCCGCCACACGCGAACTGGTTGCCGCACTACCATTTTGGGTATGCTGGGTCTTTTTTGGAGCTCAGTGGGTGCGGCTTCCCTTGCAAAAGGTTTGCCTCTTTTATGTTGGCTAGCTAGCTAGTTGGGGTTTGGGCCTATTGGGCCCTTTTAGAGGGGGCACCGCCCTCATAAAAATTGCTTGCAATATTCAAATAACTTGGGCATGAAACGCGACGGGAGGAAAAGAGAAAATAGGGCTATAAAGTGTGGAATATTAAAGGAGGTTAGAGGAAGATATTAAATCGGAGGATGCGGATAGAAGATAGAGAAAGTCACGCTAGAAATATTTCCATTTCGGAGTAGAAGCATGAATGGGAATGCGTGAGTGTTGATATCTATGCATAGATATCTAAGAGAGAGAGAGAGAGAATTTAGAGTGGAGCTATCCACTTTGAGGCTCTAGCTATTTCTAACCCTTAATTAGTTCACATATGGGTGACTTTATCATCAACTGATGTGGCCCCATCACTCGCCGCAAACCATCTCTTGAGCCCCCTCCTCTTCAGGGCGAATTTCGTCGCCGCCCCGAAGCGTTCTCGTGGGCTCCCTACGTCTGGCCTTTGACCGATCCGTGGATGTCTTGCCACCCTCCCACGCGCAGAGTAGCTGCCTTTCGGGAGCGTTTTGGGCAGGTCTTGGGCACGTAGAGTGGACCTGAGAGTTGATGGGGACTTGAGGTGGGCTACCTGCTCGTCCTATGCACGATTCCCCCTGGCGAGGGACGCGCCGCCTTGGGGGTCGGTCATGTCACCGGCCCTCTCCCTCTAGCGGGGCGGAAATGATCCGGTGCCACGTGGTGTTGGAGAGGGGGCAATCTGGCGAGTGGGGACGAGGGTATGAGGTGGTGTGTGGCGCTAGTGATGTTGGGTGCGATCATACCAGCACTAATGCACCGGATCCCATCAGAACTCCGAAGTTAAGCGTGCTTGGGCGAGAGTAGTACTAGGATGGGTGACCTCCTGGGAAGTCCTCGTGTTGCACCCCTTTTGCCTGCCGCTCTCCCCGCAGCTTGCAATTGATGTCGCTGCCTCTGAGCTTCGTCGTGATCTCCTCACCTAGGGCGAGGGCCATCTTCCGGAGGCCCTTCCTCTGGTCAGTTCATCCCCCTCTTTCCCTTCCTCCTCCGAACGCCCCGTCCGCCACACGCGAACTGGTTGCCGCACTACCATTTTGGGTATGCTGGGTCTTTTTTGGAGCTCAGTGGGTGCGGCTTCCCTTGCAAAAAGTTTGCCTCTTTTATGTTGGCTAGCTAGCTAGTTGGGGTTTGGGCCTATTGGGCCCTTTTAGAGGGGGCACCGCCCTCATAAAAATTGCTTGCAATATTCAAATAACTTGGGCATGAAACGCGACGGGAGGAAAAGAGAAAATAGGGCTATAAAGTGTGGAATATTAAAGGAGGTTAGAGGAAGATATTAAATCGGAGGATGCGGATAGAAGATAGAGAAAGTCACGCTAGAAATATTTCCATTTCGGAGTAGAAGCATGAATGGGAATGCGTGAGTGTTGATATCTATGCATAGATATCTAAGAGAGAGAGAGAGAGAATTTAGAGTGGAGCTATCCACTTTGAGGCTCTAGCTATTTCTAACCCTTAATTAGTTCACATATGGGTGACTTTATCATCAACTGATGTGGCCCCATCACTCGCCGCAAACCATCTCTTGAGCCCCCTCCTCTTCAGGGCGAATTTCGTCGCCGCCCCGAAGCGTTCTCGTGGGCTCCCTACGTCTGGCCTTTGACCGATCCGTGGATGTCTTGCCACCCTCCCACGCGCAGAGTAGCTGCCTTTCGGGAGCGTTTTGGGCAGGTCTTGGGCACGTAGAGTGGACCTGAGAGTTGATGGGGACTTGAGGTGGGCTACCTGCTCGTCCTATGCACGATTCCCCCTGGCGAGGGACGCGCCGCCTTGGGGGTCGGTCATGTCACCGGCCCTCTCCCTCTAGCGGGGCGGAAATGATCCGGTGCCACGTGGTGTTGGAGAGGGGGCAATCTGGCGAGTGGGGACGAGGGTATGAGGTGGTGTGTGGCGCTAGTGATGTTGGGTGCGATCATACCAGCACTAATGCACCGGATCCCATCAGAACTCCGAAGTTAAGCGTGCTTGGGCGAGAGTAGTACTAGGATGGGTGACCTCCTGGGAAGTCCTCGTGTTGCACCCCTTTTGCCTGCCGCTCTCCCCGCAGCTTGCAATTGATGTCGCTGCCTCTGAGCTTCGTCGTGATCTCCTCACCTAGGGCGAGGGCCATCTTCCGGAGGCCCTTCCTCTGGTCAGTTCATCCCCCTCTTTCCCTTCCTCCTCCGAACGCCCCGTCCGCCACACGCGAACTGGTTGCCGCACTACCATTTTGGGTATGCTGGGTCTTTTTTGGAGCTCAGTGGGTGCGGCTTCCCTTGCAAAAGGTTTGCCTCTTTTATGTTGGCTAGCTAGCTAGTTGGGGTTTGGGCCTATTGGGCCCTTTTAGAGGGGGCACCGCCCTCATAAAAATTGCTTGCAATATTCAAATAACTTGGGCATGAAACGCGACGGGAGGAAAAGAGAAAATAGGGCTATAAAGTGTGGAATATTAAAGGAGGTTAGAGGAAGATATTAAATCGGAGGATGCGGATAGAAGATAGAGAAAGTCACGCTAGAAATATTTCCATTTCGGAGTAGAAGCATGAATGGGAATGCGTGAGTGTTGATATCTATGCATAGATATCTAAGAGAGAGAGAGAGAGAATTTAGAGTGGAGCTATCCACTTTGAGGCTCTAGCTATTTCTAACCCTTAATTAGTTCACATATGGGTGACTTTATCATCAACTGATGTGGCCCCATCACTCGCCGCAAACCATCTCTTGAGCCCCCTCCTCTTCAGGGCGAATTTCGTCGCCGCCCCGAAGCGTTCTCGTGGGCTCCCTACGTCTGGCCTTTGACCGATCCGTGGATGTCTTGCCACCCTCCCACGCGCAGAGTAGCTGCCTTTCGGGAGCGTTTTGGGCAGGTCTTGGGCACGTAGAGTGGACCTGAGAGTTGATGGGGACTTGAGGTGGGCTACCTGCTCGTCCTATGCACGATTCCCCCTGGCGAGGGACGCGCCGCCTTGGGGGTCGGTCATGTCACCGGCCCTCTCCCTCTAGCGGGGCGGAAATGATCCGGTGCCACGTGGTGTTGGAGAGGGGGCAATCTGGCGAGTGGGGACGAGGGTATGAGGTGGTGTGTGGCGCTAGTGATGTTGGGTGCGATCATACCAGCACTAATGCACCGGATCCCATCAGAACTCCGAAGTTAAGCGTGCTTGGGCGAGAGTAGTACTAGGATGGGTGACCTCCTGGGAAGTCCTCGTGTTGCACCCCTTTTGCCTGCCGCTCTCCCCGCAGCTTGCAATTGATGTCGCTGCCTCTGAGCTTCGTCGTGATCTCCTCACCTAGGGCGAGGGCCATCTTCCGGAGGCCCTTCCTCTGGTCAGTTCATCCCCCTCTTTCCCTTCCTCCTCCGAACGCCCCGTCCGCCACACGCGAACTGGTTGCCGCACTACCATTTTGGGTATGCTGGGTCTTTTTTGGAGCTCAGTGGGTGCGGCTTCCCTTGCAAAAGGTTTGCCTCTTTTATGTTGGCTAGCTAGCTAGTTGGGGTTTGGGCCTATTGGGCCCTTTTAGAGGGGGCACCGCCCTCATAAAAATTGCTTGCAATATTCAAATAACTTGGGCATGAAACGCGACGGGAGGAAAAGAGAAAATAGGGCTATAAAGTGTGGAATATTAAAGGAGGTTAGAGGAAGATATTAAATCGGAGGATGCGGATAGAAGATAGAGAAAGTCACGCTAGAAATATTTCCATTTCGGAGTAGAAGCATGAATGGGAATGCGTGAGTGTTGATATCTATGCATAGATATCTAAGAGAGAGAGAGAGAGAATTTAGAGTGGAGCTATCCACTTTGAGGCTCTAGCTATTTCTAACCCTTAATTAGTTCACATATGGGTGACTTTATCATCAACTGATGTGGCCCCATCACTCGCCGCAAACCATCTCTTGAGCCCCCTCCTCTTCAGGGCGAATTTCGTCGCCGCCCCGAAGCGTTCTCGTGGGCTCCCTACGTCTGGCCTTTGACCGATCCGTGGATGTCTTGCCACCCTCCCACGCGCAGAGTAGCTGCCTTTCGGGAGCGTTTTGGGCAGGTCTTGGGCACGTAGAGTGGACCTGAGAGTTGATGGGGACTTGAGGTGGGCTACCTGCTCGTCCTATGCACGATTCCCCCTGGCGAGGGACGCGCCGCCTTGGGGGTCGGTCATGTCACCGGCCCTCTCCCTCTAGCGGGGCGGAAATGATCCGGTGCCACGTGGTGTTGGAGAGGGGGCAATCTGGCGAGTGGGGACAAGGGTATGAGGTGGTGTGTGGCGCTAGTGATGTTGGGTGCGATCATACCAGCACTAATGCACCGGATCCCATCAGAACTCCGAAGTTAAGCGTGCTTGGGCGAGAGTAGTACTAGGATGGGTGACCTCCTGGGAAGTCCTCGTGTTGCACCCCTTTTGCCTGCCGCTCTCCCCGCAGCTTGCAATTGATGTCGCTGCCTCTGAGCTTCGTCGTGATCTCCTCACCTAGGGCGAGGGCCATCTTCCGGAGGCCCTTCCTCTGGTCAGTTCATCCCCCTCTTTCCCTTCCTCCTCCGAACGCCCCGTCCGCCACACGCGAACTGGTTGCCGCACTACCATTTTGGGTATGCTGGGTCTTTTTTGGAGCTCAGTGGGTGCGGCTTCCCTTGCAAAAGGTTTGCCTCTTTTATGTTGGCTAGCTAGCTAGTTGGGGTTTGGGCCTATTGGGCCCTTTTAGAGGGGGCACCGCCCTCATAAAAATTGCTTGCAATATTCAAATAACTTGGGCATGAAACGCGACGGGAGGAAAAGAGAAAATAGGGCTATAAAGTGTGGAATATTAAAGGAGGTTAGAGGAAGATATTAAATCGGAGGATGCGGATAGAAGATAGAGAAAGTCACGCTAGAAATATTTCCATTTCGGAGTAGAAGCATGAATGGGAATGCGTGAGTGTTGATATCTATGCATAGATATCTAAGAGAGAGAGAGAGAGAATTTAGAGTGGAGCTATCCACTTTGAGGCTCTAGCTATTTCTAACCCTTAATTAGTTCACATATGGGTGACTTTATCATCAACTGATGTGGCCCCATCACTCGCCGCAAACCATCTCTTGAGCCCCCTCCTCTTCAGGGCGAATTTCGTCGCCGCCCCGAAGCGTTCTCGTGGGCTCCCTACGTCTGGCCTTTGACCGATCCGTGGATGTCTTGCCACCCTCCCACGCGCAGAGTAGCTGCCTTTCGGGAGCGTTTTGGGCAGGTCTTGGGCACGTAGAGTGGACCTGAGAGTTGATGGGGACTTGAGGTGGGCTACCTGCTCGTCCTATGCACGATTCCCCCTGGCGAGGGACGCGCCGCCTTGGGGGTCGGTCATGTCACCGGCCCTCTCCCTCTAGCGGGGCGGAAATGATCCGGTGCCACGTGGTGTTGGAGAGGGGGCAATCTGGCGAGTGGGGACGAGGGTATGAGGTGGTGTGTGGCGCTAGTGATGTTGGGTGCGATCATACCAGCACTAATGCACCGGATCCCATCAGAACTCCGAAGTTAAGCGTGCTTGGGCGAGAGTAGTACTAGGATGGGTGACCTCCTGGGAAGTCCTCGTGTTGCACCCCTTTTGCCTGCCGCTCTCCCCGCAGCTTGCAATTGATGTCGCTGCCTCTGAGCTTCGTCGTGATCTCCTCACCTAGGGCGAGGGCCATCTTCCGGAGGCCCTTCCTCTGGTCAGTTCATCCCCCTCTTTCCCTTCCTCCTCCGAACGCCCCGTCCGCCACACGCGAACTGGTTGCCGCACTACCATTTTGGGTATGCTGGGTCTTTTTTGGAGCTCAGTGGGTGCGGCTTCCCTTGCAAAAGGTTTGCCTCTTTTATGTTGGCTAGCTAGCTAGTTGGGGTTTGGGCCTATTGGGCCCTTTTAGAGGGGGCACCGCCCTCATAAAAATTGCTTGCAATATTCAAATAACTTGGGCATGAAACGCGACGGGAGGAAAAGAGAAAATAGGGCTATAAAGTGTGGAATATTAAAGGAGGTTAGAGGAAGATATTAAATCGGAGGATGCGGATAGAAGATAGAGAAAGTCACGCTAGAAATATTTCCATTTCGGAGTAGAAGCATGAATGGGAATGCGTGAGTGTTGATATCTATGCATAGATATCTAAGAGAGAGAGAGAGAGAATTTAGAGTGGAGCTATCCACTTTGAGGCTCTAGCTATTTCTAACCCTTAATTAGTTCACATATGGGTGACTTTATCATCAACTGATGTGGCCCCATCACTCGCCGCAAACCATCTCTTGAGCCCCCTCCTCTTCAGGGCGAATTTCGCCGCCGCCCCGAAGCGTTCTCGTGGGCTCCCTACGTCTGGCCTTTGACCGATCCGTGGATGTCTTGCCACCCTCCCACGCGCAGAGTAGCTGCCTTTCGGGAGCGTTTTGGGCAGGTCTTGGGCACGTAGAGTGGACCTGAGAGTTGATGGGGACTTGAGGTGGGCTACCTGCTCGTCCTATGCACGATTCCCCCTGGCGAGGGACGCGCCGCCTTGGGGGTCGGTCATGTCACCGGCCCTCTCCCTCTAGCGGGGCGGAAATGATCCGGTGCCACGTGGTGTTGGAGAGGGGGCAATCTGGCGAGTGGGGACGAGGGTATGAGGTGGTGTGTGGCGCTAGTGATGTTGGGTGCGATCATACCAGCACTAATGCACCGGATCCCATCAGAACTCCGAAGTTAAGCGTGCTTGGGCGAGAGTAGTACTAGGATGGGTGACCTCCTGGGAAGTCCTCGTGTTGCACCCCTTTTGCCTGCCGCTCTCCCCGCAGCTTGCAATTGATGTCGCTGCCTCTGAGCTTCGTCGTGATCTCCTCACCTAGGGCGAGGGCCATCTTCCGGAGGCCCTTCCTCTGGTCAGTTCATCCCCCTCTTTCCCTTCCTCCTCCGAACGCCCCGTCCGCCACACGCGAACTGGTTGCCGCACTACCATTTTGGGTATGCTGGGTCTTTTTTGGAGCTCAGTGGGTGCGGCTTCCCTTGCAAAAGGTTTGCCTCTTTTATGTTGGCTAGCTAGCTAGTTGGGGTTTGGGCCTATTGGGCCCTTTTAGAGGGGGCACCGCCCTCATAAAAATTGCTTGCAATATTCAAATAACTTGGGCATGAAACGCGACGGGAGGAAAAGAGAAAATAGGGCTATAAAGTGTGGAATATTAAAGGAGGTTAGAGGAAGATATTAAATCGGAGGATGCGGATAGAAGATAGAGAAAGTCACGCTAGAAATATTTCCATTTCGGAGTAGAAGCATGAATGGGAATGCGTGAGTGTTGATATCTATGCATAGATATCTAAGAGAGAGAGAGAGAGAATTTAGAGTGGAGCTATCCACTTTGAGGCTCTAGCTATTTCTAACCCTTAATTAGTTCACATATGGGTGACTTTATCATCAACTGATGTGGCCCCATCACTCGCCGCAAACCATCTCTTGAGCCCCCTCCTCTTCAGGGCGAATTTCGTCGCCGCCCCGAAGCGTTCTCGTGGGCTCCCTACGTCTGGCCTTTGACCGATCCGTGGATGTCTTGCCACCCTCCCACGCGCAGAGTAGCTGCCTTTCGGGAGCGTTTTGGGCAGGTCTTGGGCACGTAGAGTGGACCTGAGAGTTGATGGGGACTTGAGGTGGGCTACCTGCTCGTCCTATGCACGATTCCCCCTGGCGAGGGACGCGCCGCCTTGGGGGTCGGTCATGTCACCGGCCCTCTCCCTCTAGCGGGGCGGAAATGATCCGGTGCCACGTGGTATTGGAGAGGGGGCAATCTGGCGAGTGGGGACGAGGGTATGAGGTGGTGTGTGGCGCTAGTGATGtTGGGTGCGATCATACCAGCACTAATGCACCGGATCCCATCAGAACTCCGAAGTTAAGCGTGCTTGGGCGAGAGTAGTACTAGGATGGGTGACCTCCTGGGAAGTCCTCGTGTTGCACCCCTTTTGCCTGCCGCTCTCCCCGCAGCTTGCAATTGATGTCGCTGCCTCTGAGCTTCGTCGTGATCTCCTCACCTAGGGCGAGGGCCATCTTCCGGAGGCCCTTCCTCTGGTCAGTTCATCCCCCTCTTTCCCTTCCTCCTCCGAACGCCCCGTCCGCCACACGCGAACTGGTTGCCGCACTACCATTTTGGGTATGCTGGGTCTTTTTTGGAGCTCATTGGGTGCGGCTTCCCTTGCAAAAGGTTTGCCTCTTTTATGTTGGCTAGCTAGCTAGTTGGGGTTTGGGCCTATTGGGCCCTTTTAGAGGGGGCACCGCCCTCATAAAAATTGCTTGCAATATTCAAATAACTTGGGCATGAAACGCGACGGGAGGAAAAGAGAAAATAGGGCTATAAAGTGTGGAATATTAAAGGAGGTTAGAGGAAGATATTAAATCGGAGGATGCGGATAGAAGATAGAGAAAGTCACGCTAGAAATATTTCCATTTCGGAGTAGAAGCATGAATGGGAATGCGTGAGTGTTGATATCTATGCATAGATATCTAAGAGAGAGAGAGAGAGAATTTAGAGTGGAGCTATCCACTTTGAGGCTCTAGCTATTTCTAACCCTTAATTAGTTCACATATGGGTGACTTTATCATCAACTGATGTGGCCCCATCACTCGCCGCAAACCATCTCTTGAGCCCCCTCCTCTTCAGGGCGAACTTCGTCGCCGCCCCGAAGCGTTCTCGTGGGCTCCCTACGTCTGGCCTTTGACCGATCCGTGGATGTCTTGCCACCCTCCCACGCGCAGAGTAGCTGCCTTTCGGGAGCGTTTTGGGCAGGTCTTGGGCACGTAGAGTGGACCTGAGAGTTGATGGGGACTTGAGGTGGGCTACCTGCTCGTCCTATGCACGATTCCCCCGGGCGAGGGACGCGCCGCCTTGGGGGTCGGTCATGTCACCGGCCCTCTCCCTCGAGCGGGGCGGAAATGATCCGGTGCCACGTGGTGTTGGAGAGGGGGCAATCTGGCGAGTGGGGACGAGGGTATGAGGTGGTGTGTGGCGCTAGTGATGTTGGGTGCGATCATACCAGCACTAATGCACCGGATCCCATCAGAACTCCGAAGTTAAGCGTGCTTGGGCGAGAGTAGTACTAGGATGGGTGACCTCCTGGGAAGTCCTCGTGTTGCACCCCTTTTGCCTGCCGCTCTCCCCGCAGCCTGCAATTGATGTCGCTGCCTCTGAGCTTCGTCGTGATCTCCTCACCTAGGGCGAGGGCCATCTTCCGGAGGCCCTTCCTCTGGTCAGTTCATCCCCCTCTTTCCCTTCCTCCTCCGAACGCCCCGTCCGCCACACGCGAACTGGTTGCCGCACTACCATTTTGGGTATGCTGGGTCTTTTTTGGAGCTCAGTGGGTGCGGCTTCCCTTGCAAAAGGTTTGCCTCTTTTATGTTGGCTAGCTAGCTAGTTGGGGTTTGGGCCTATTGGGCCCTTTTAGAGGGGGCACCGCCCTCATAAAAATTGCTTGCAATATTCAAATAACTTGGGCATGAAACGCGACGGGAGGAAAAGAGAAAATAGGGCTATAAAGTGTGGAATATTAAAGGAGGTTAGAGGAAGATATTAAATCGGAGGATGCGGATAGAAGATAGAGAAAGTCACGCTAGAAATATTTCCATTTCGGAGTAGAAGCATGAATGGGAATGCGTGAGTGTTGATATCTATGCATAGATATCTAAGAGAGAGAGAGAGAGAATTTAGAGTGGAGCTATCCACTTTGAGGCTCTAGCTATTTCTAACCCTTAATTAGTTCACATATGGGTGACTTTATCATCAACTGATGTGGCCCCATCACTCGCCGCAAACCATCTCTTGAGCCCCCTCCTCTTCAGGGCGAATTTCGTCGCCGCCCCGAAGCGTTCTCGTGGGCTCCCTACGTCTGGCCTTTGACCGATCCGTGGATGTCTTGCCACCCTCCCACGCGCAGAGTAGCTGCCTTTCGGGAGCGTTTTGGGCAGGTCTTGGGCACGTAGAGTGGACCTGAGAGTTGATGGGGACTTGAGGTGGGCTACCTGCTCGTCCTATGCACGATTCCCCCTGGCGAGGGACGCGCCGCCTTGGGGGTCGGTCATGTCACCGGCCCTCTCCCTCTAGCGGGGCGGAAATGATCCGGTGCCACGTGGTGTTGGAGAGGGGGCAATCTGGCGAGTGGGGACGAGGGTATGAGGTGGTGTGTGGCGCTAGTGATGTTGGGTGCGATCATACCAGCACTAATGCACCGGATCCCATCAGAACTCCGAAGTTAAGCGTG**CTTGGGCGAGAGTAGTACTAGGATGGGTGACCTCCTGGGAAGTCCTCGTGTTGCACCCCTTTTGCCTGCCGCTCTCCCCGCAGCTTGCAATTGATGTCGCTGCCTCTGAGCTTCGTCGTGATCTCCTCACCTAGGGCGAGGGCCATCTTCCGGAGGCCCTTCCTCTGGTCAGTTCATCCCCCTCTTTCCCTTCCTCCTCCGAACGCCCCGTCCGCCACACGCGAACTGGTTGCCGCACTACCATTTTGGGTATGCTGGGTCTTTTTTGGAGCTCAGTGGGTGCGGCTTCCCTTGCAAAAGGTTTGCCTCTTTTATGTTGGCTAGCTAGCTAGTTGGGGTTTGGGCCTATTGGGCCCTTTTAGAGGGGGCACCGCCCTCATAAAAATTGCTTGCAATATTCAAATAACTTGGGCATGAAACGCGACGGGAGGAAAAGAGAAAATAGGGCTATAAAGTGTGGAATATTAAAGGAGGTTAGAGGAAGATATTAAATCGGAGGATGCGGATAGAAGATAGAGAAAGTCACGCTAGAAATATTTCCATTTCGGAGTAGAAGCATGAATGGGAATGCGTGAGTGTTGATATCTATGCATAGATATCTAAGAGAGAGAGAGAGAGAATTTAGAGTGGAGCTATCCACTTTGAGGCTCTAGCTATTTCTAACCCTTAATTAGTTCACATATGGGTGACTTTATCATCAACTGATGTGGCCCCATCACTCGCCGCAAACCATCTCTTGAGCCCCCTCCTCTTCAGGGCGAATTTCGTCGCCGCCCCGAAGCGTTCTCGTGGGCTCCCTACGTCTGGCCTTTGACCGATCCGTGGATGTCTTGCCACCCTCCCACGCGCAGAGTAGCTGCCTTTCGGGAGCGTTTTGGGCAGGTCTTGGGCACGTAGAGTGGACCTGAGAGTTGATGGGGACTTGAGGTGGGCTACCTGCTCGTCCTATGCACGATTCCCCCTGGCGAGGGACGCGCCGCCTTGGGGGTCGGTCATGTCACCGGCCCTCTCCCTCTAGCGGGGCGGAAATGATCCGGTGCCACGTGGTGTTGGAGAGGGGGCAATCTGGCGAGTGGGGACGAGGGTATGAGGTGGTGTGTGGCGCTAGTGATGTTGGGTGCGATCATACCAGCACTAATGCACCGGATCCCATCAGAACTCCGAAGTTAAGCGTGCTTGGGCGAGAGTAGTACTAGGATGGGTGACCTCCTGGGAAGTCCTCGTGTTGCACCCCTTTTGCCTGCCGCTCTCCCCGCAGCTTGCAATTGATGTCGCTGCCTCTGAGCTTCGTCGTGATCTCCTCACCTAGGGCGAGGGCCATCTTCCGGAGGCCCTTCCTCTGGTCAGTTCATCCCCCTCTTTCCCTTCCTCCTCCGAACGCCTCGTCCGCCACACGCGAACTGGTTGCCGCACTACCATTTTGGGTATGCTGGGTCTTTTTTGGAGCTCAGTGGGTGCGGCTTCCCTTGCAAAAGGTTTGCCTCTTTTATGTTGGCTAGCTAGCTAGTTGGGGTTTGGGCCTATTGGGCCCTTTTAGAGGGGGCACCGCCCTCATAAAAATTGCTTGCAATATTCAAATAACTTGGGCATGAAACGCGACGGGAGGAAAAGAGAAAATAGGGCTATAAAGTGTGGAATATTAAAGGAGGTTAGAGGAAGATATTAAATCGGAGGATGCGGATAGAAGATAGAGTAAGTCACGCTAGAAATATTTCCATTTCGGAGTAGAAGCATGAATGGGAATGCGTGAGTGTTGATATCTATGCATAGATATCTAAGAGAGAGAGAGAGAGAATTTAGAGTGGAGCTATCCACTTTGAGGCTCTAGCTATTTCTAACCCTTAATTAGTTCACATATGGGTGACTTTATCATCAACTGATGTGGCCCCATCACTCGCCGCAAACCATCTCTTGAGCCCCCTCCTCTTCAGGGCGAATTTCGTCGCCGCCCCGAAGCGTTCTCGTGGGCTCCCTACGTCTGGCCTTTGACCGATCCGTGGATGTCTTGCCACCCTCCCACGCGCAGAGTAGCTGCCTTTCGGGAGCGTTTTGGGCGGGTCTTGGGCACGTAGAGTGGACCTGAGAGTTGATGGGGACTTGAGGTGGGCTACCTGCTCGTCCTATGCACGATTCCCCCTGGCGAGGGACGCGCCGCCTTGGGGGTCGGTCATGTCACCGGCCCTCTCCCTCTAGCGGGGCGGAAATGATCCGGTGCCACGTGGTGTTGGAGAGGGGGCAATCTGGCGAGTGGGGACGAGGGTATGAGGTGGTGTGTGGCGCTAGTGATGTTGGGTGCGATCATACCAGCACTAATGCACCGGATCCCATCAGAACTCCGAAGTTAAGCGTGCTTGGGCGAGAGTAGTACTAGGATGGGTGACCTCCTGGGAAGTCCTCGTGTTGCACCCCTTTTGCCTGCCGCTCTCCCCGCAGCTTGCAATTGATGTCGCTGCCTCTGAGCTTCGTCGTGATCTCCTCACCTAGGGCGAGGGCCATCTTCCGGAGGCCCTTCCTCTGGTCAGTTCATCCCCCTCTTTCCCTTCCTCCTCCGAACGCCTCGTCCGCCACACGCGAACTGGTTGCCGCACTACCATTTTGGGTATGCTGGGTCTTTTTTGGAGCTCAGTGGGTGCGGCTTCCCTTGCAAAAGGTTTGCCTCTTTTATGTTGGCTAGCTAGCTAGTTGGGGTTTGGGCCTATTGGGCCCTTTTAGAGGGGGCACCGCCCTCATAAAAATTGCTTGCAATATTCAAATAACTTGGGCATGAAACGCGACGGGAGGAAAAGAGAAAATAGGGCTATAAAGTGTGGAGTATTAAAGGAGGTTAGAGGAAGATATTAAATCGGAGGATGCGGATAGAAGATAGAGAAAGTCACGCTAGAAATATTTCCATTTCGGAGTAGAAGCATGAATGGGAATGCGTGAGTGTTGATATCTATGCATAGATATCTAAGAGAGAGAGAGGGAGAATTTAGAGTGGAGCTATCCACTTTGAGGCTCTAGCTATTTCTAACCCTTAATTAGTTCACATATGGGTGACTTTATCATCAACTGATGTGGCCCCATCACTCGCCGCAAACCATCTCTTGAGCCCCCTCCTCTTCAGGGCGAATTTCGTCGCCGCCCCGAAGCGTTCTCGTGGGCTCCCTACGTCTGGCCTTTGACCGATCCGTGGATGTCTTGCCACCCTCCTACGCGCAGAGTAGCTGCCTTTCGGGAGCGTTTTGGGCAGGTCTTGGGCACGTAGAGTGGACCTGAGAGTTGATGGGGACTTGAGGTGGGCTACCTGCTCGTCCTATGCACGATTCCCCCTGGCGAGGGACGCGCCGCCTTGGGGGTCGGTCATGTCACCGGCCCTCTCCCTCTAGCGGGGCGGAAATGATCCGGTGCCACGTGGTGTTGGAGAGGGGGCAATCTGGCGAGTGGGGACGAGGGTATGAGGTGGTGTGTGGCGCTAGTGATGTTGGGTGCGATCATACCAGCACTAATGCACCGGATCCCATCAGAACTCCGAAGTTAAGCGTGCTTGGGCGAGAGTAGTACTAGGATGGGTGACCTCCTGGGAAGTCCTCGTGTTGCACCCCTTTTGCCTGCCGCTCTCCCCGCAGCTTGCAATTGATGTCGCTGCCTCTGAGCTTCGTCGTGATCTCCTCACCTAGGGCGAGGGCCATCTTCCGGAGGCCCTTCCTCTGGTCAGTTCATCCCCCTCTTTCCCTTCCTCCTCCGAACGCCTCGTCCGCCACACGCGAACTGGTTGCCGCACTACCATTTTGGGTATGCTGGGTCTTTTTTGGAGCTCAGTGGGTGCGGCTTCCCTTGCAAAAGGTTTGCCTCTTTTATGTTGGCTAGCTAGCTAGTTGGGGTTTGGGCCTATTGGGCCCTTTTAGAGGGGGCACCGCCCTCATAAAAATTGCTTGCAATATTCAAATAACTTGGGCATGAAACGCGACGGGAGGAAAAGAGAAAATAGGGCTATAAAGTGTGGAATATTAAAGGAGGTTAGAGGAAGATATTAAATCGGAGGATGCGGATAGAAGATAGAGAAAGTCACGCTAGAAATATTTCCATTTCGGAGTAGAAGCATGAATGGGAATGCGTGAGTGTTGATATCTATGCATAGATATCTAAGAGAGAGAGAGAGAGAATTTAGAGTGGAGCTATCCACTTTGAGGCTCTAGCTATTTCTAACCCTTAATTAGTTCACATATGGGTGACTTTATCATCAACTGATGTGGCCCCATCACTCGCCGCAAACCATCTCTTGAGCCCCCTCCTCTTCAGGGCGAATTTCGTCGCCGCCCCGAAGCGTTCTCGTGGGCTCCCTACGTCTGGCCTTTGACCGATCCGTGGATGTCTTGCCACCCTCCCACGCGCAGAGTAGCTGCCTTTCGGGAGCGTTTTGGGCAGGTCTTGGGCACGTAGAGTGGACCTGAGAGTTGATGGGGACTTGAGGTGGGCTACCTGCTCGTCCTATGCACGATTCCCCCTGGCGAGGGACGCGCCGCCTTGGGGGTCGGTCATGTCACCGGCCCTCTCCCTCTAGCGGGGCGGAAATGATCCGGTGCCACGTGGTGTTGGAGAGGGGGCAATCTGGCGAGTGGGGACGAGGGTATGAGGTGGTGTGTGGCGCTAGTGATGTTGGGTGCGATCATACCAGCACTAATGCACCGGATCCCATCAGAACTCCGAAGTTAAGCGTGCTTGGGCGAGAGTAGTACTAGGATGGGTGACCTCCTGGGAAGTCCTCGTGTTGCACCCCTTTTGCCTGCCGCTCTCCCCGCAGCTTGCAATTGATGTCGCTGCCTCTGAGCTTCGTCGTGATCTCCTCACCTAGGGCGAGGGCCATCTTCCGGAGGCCCTTCCTCTGGTCAGTTCATCCCCCTCTTTCCCTTCCTCCTCCGAACGCCTCGTCCGCCACACGCGAACTGGTTGCCGCACTATCATTTTGGGTATGCTGGGTCTTTTTTGGAGCTCAGTGGGTGCGGCTTCCCTTGCAAAAGGTTTGCCTCTTTTATGTTGGCTAGCTAGCTAGTTGGGGTTTGGGCCTATTGGGCCCTTTTAGAGGGGGCACCGCCCTCATAAAAATTGCTTGCAATATTCAAATAACTTGGGCATGAAACGCGACGGGAGGAAAAGAGAAAATAGGGCTATAAAGTGTGGAATATTAAAGGAGGTTAGAGGAAGATATTAAATCGGAGGATGCGGATAGAAGATAGAGAAAGTCACGCTAGAAATATTTCCATTTCGGAGTAGAAGCATGAATGGGAATGCGTGAGTGTTGATATCTATGCATAGATATCTAAGAGAGAGAGAGAGAGAATTTAGAGTGGAGCTATCCACTTTGAGGCTCTAGCTATTTCTAACCCTTAATTAGTTCACATATGGGTGACTTTATCATCAACTGATGTGGCCCCATCACTCGCCGCAAACCATCTCTTGAGCCCCCTCCTCTTCAGGGCGAATTTCGTCGCCGCCCCGAAGCGTTCTCGTGGGCTCCCTACGTCTGGCCTTTGACCGATCCGTGGATGTCTTGCCACCCTCCCACGCGCAGAGTAGCTGCCTTTCGGGAGCGTTTTGGGCAGGTCTTGAGCACGTAGAGTGGACCTGAGAGTTGATGGGGACTTGAGGTGGGCTACCTGCTCGTCCTATCCACGATTCCCCCTGGCGAGGGACGCGCCGCCTTGGGGGTCGGTCATGTCACCGGCCCTCTCCCTCTAGCGGGGCGGAAATGATCCGGTGCCACGTGGTGTTGGAGAGGGGGCAATCTGGCGAGTGGGGACGAGGGTATGAGGTGGTGTGTGGCGCTAGTGATGTTGGGTGCGATCATACCAGCACTAATGCACCGGATCCCATCAGAACTCCGAAGTTAAGCGTGCTTGGGCGAGAGTAGTACTAGGATGGGTGACCTCCTGGGAAGTCCTCGTGTTGCACCCCTTTTGCCTGCCGCTCTCCCCGCAGCTTGCAATTGATGTCGCTGCCTCTGAGCTTCGTCGTGATCTCCTCACCTAGGGCGAGGGCCATCTTCCGGAGGCCCTTCCTCTGGTCAGTTCATCCCCCTCTTTCCCTTCCTCCTCCGAACGCCCCGTCCGCCACACGCGAACTGGTTGCCGCACTACCATTTTGGGTATGCTGGGTCTTTTTTGGAGCTCAGTGGGTGCGGCTTCCCTTGCAAAAGGTTTGCCTCTTTTATGTTGGCTAGCTAGCTAGTTGGGGTTTGGGCCTATTGGGCCCTTTTAGAGGGGGCACCGCCCTCATAAAAATTGCTTGCAATATTCAAATAACTTGGGCATGAAACGCGACGGGAGGAAAAGAGAAAATAGGGCTATAAAGTGTGGAATATTAAAGGAGGTTAGAGGAAGATATTAAATCGGAGGATGCGGATAGAAGATAGAGAAAGTCACGCTAGAAATATTTCCATTTCGGAGTAGAAGCATGAATGGGAATGCGTGAGTGTTGATATCTATGCATAGATATCTAAGAGAGAGAGAGAGAGAATTTAGAGTGGAGCTATCCACTTTGAGGCTCTAGCTATTTCTAACCCTTAATTAGTTCACATATGGGTGACTTTATCATCAACTGATGTGGCCCCATCACTCGCCGCAAACCATCTCTTGAGCCCCCTCCTCTTCAGGGCGAATTTCGTCGCCGCCCCGAAGCGTTCTCGTGGGCTCCCTACGTCTGGCCTTTGACCGATCCGTGGATGTCTTGCCACCCTCCCACGCGCAGAGTAGCTGCCTTTCGGGAGCGTTTTGGGCAGGTCTTGGGCACGTAGAGTGGACCTGAGAGTTGATGGGGACTTGAGGTGGGCTACCTGCTCGTCCTATGCACGATTCCCCCTGGCGAGGGACGCGCCGCCTTGGGGGTCGGTCATGTCACCGGCCCTCTCCCTCTAGCGGGGCGGAAATGATCCGGTGCCACGTGGTGTTGGAGAGGGGGCAATCTGGCGAGTGGGGACGAGGGTATGAGGTGGTGTGTGGCGCTAGTGATGTTGGGTGCGATCATACCAGCACTAATGCACCGGATCCCATCAGAACTCCGAAGTTAAGCGTGCTTGGGCGAGAGTAGTACTAGGATGGGTGACCTCCTGGGAAGTCCTCGTGTTGCACCCCTTTTGCCTGCCGCTCTCCCCGCAGCTTGCAATTGATGTCGCTGCCTCTGAGCTTCGTCGTGATCTCCTCACCTAGGGCGGGGGCCATCTTCCGGAGGCCCTTCCTCTGGTCAGTTCATCCCCCTCTTTCCCTTCCTCCTCCGAACGCCCCGTCCGCCACACGCGAACTGGTTGCCGCACTACCATTTTGGGTCTGCTGGGTCTTTTTTGGAGCTCAGTGGGTGCGGCTTCCCTTGCAAAAGGTTTGCCTCTTTTATGTTGGCTAGCTAGCTAGTTGGGGTTTGGGCCTATTGGGCCCTTTTAGAGGGGGCACCGCCCTCATAAAAATTGCTTGCAATATTCAAATAACTTGGGCATGAAACGCGACGGGAGGAAAAGAGAAAATAGGGCTATAAAGTGTGGAATATTAAAGGAGGTTAGAGGAAGATATTAAATCGGAGGATGCGGATAGAAGATAGAGAAAGTCACGCTAGAAATATTTCCATTTCGGAGTAGAAGCATGAATGGGAATGCGTGAGTGTTGATATCTATGCATAGATATCTAAGAGAGAGAGAGAGAGAATTTAGAGTGGAGCTATCCACTTTGAGGCTCTAGCTATTTCTAACCCTTAATTAGTTCACATATGGGTGACTTTATCATCAACTGATGTGGCCCCATCACTCGCCGCAAACCATCTCTTGAGCCCCCTCCTCTTCAGGGCGAATTTCGTCGCCGCCCCGAAGCGTTCTCGTGGGCTCCCTACGTCTGGCCTTTGACCGATCCGTGGATGTCTTGCCACCCTCCCACGCGCAGAGTAGCTGCCTTTCGGGAGCGTTTTGGGCAGGTCTTGGGCACGTAGAGTGGACCTGAGAGTTGATGGGGACTTGAGGTGGGCTACCTGCTCGTCCTATGCACGATTCCCCCTGGCGAGGGACGCGCCGCCTTGGGGGTCGGTCATGTCACCGGCCCTCTCCCTCTAGCGGGGCGGAAATGATCCGGTGCCACGTGGTGTTGGAGAGGGGGCAATCTGGCGAGTGGGGACGAGGGTATGAGGTGGTGTGTGGCGCTAGTGATGTTGGGTGCGATCATACCAGCACTAATGCACCGGATCCCATCAGAACTCCGAAGTTAAGCGTGCTTGGGCGAGAGTAGTACTAGGATGGGTGACCTCCTGGGAAGTCCTCGTGTTGCACCCCTTTTGCCTGCCGCTCTCCCCGCAGCTTGCAATTGATGTCGCTGCCTCTGAGCTTCGTCGTGATCTCCTCACCTAGGGCGAGGGCCATCTTCCGGAGGCCCTTCCTCTGGTCAGTTCATCCCCCTCTTTCCCTTCCTCCTCCGAACGCCCCGTCCGCCACGCGCGAACTGGTTGCCGCACTACCATTTTGGGTATGCTGGGTCTTTTTTGGAGCTCAGTGGGTGCGGCTTCCCTTGCAAAAGGTTTGCCTCTTTTATGTTGGCTAGCTAGCTAGTTGGGGTTTGGGCCTATTGGGCCCTTTTAGAGGGGGCACCGCCCTCATAAAAATTGCTTGCAATATTCAAATAACTTGGGCATGAAACGCGACGGGAGGAAAGGAGAAAATAGGGCTATAAAGTGTGGAATATTAAAGGAGGTTAGAGGAAGATATTAAATCGGAGGATGCGGATAGAAGATAGAGAAAGTCACGCTAGAAATATTTCCATTTCGGAGTAGAAGCATGAGTGGGAATGCGTGAGTGTTGATATCTATGCATAGATATCTAAGAGAGAGAGAGAGAGAATTTAGAGTGGAGCTATCCACTTTGAGGCTCTAGCTATTTCTAACCCTTAATTAGTTCACATATGGGTGACTTTATCATCAACTGATGTGGCCCCATCACTCGCCGCAAACCATCTCTTGAGCCCCCTCCTCTTCAGGGCGAATTTCGTCGCCGCCCCGAAGCGTTCTCGTGGGCTCCCTACGTCTGGCCTTTGACCGATCCGTGGATGTCTTGCCACCCTCCCACGCGCAGAGTAGCTGCCTTTCGGGAGCGTTTTGGGCAGGTCTTGGGCACGTAGAGTGGACCTGAGAGTTGATGGGGACTTGAGGTGGGCTACCTGCTCGTCCTATGCACGATTCCCCCTGGCGAGGGACGCGCCGCCTTGGGGGTCGGTCATGTCACCGGCCCTCTCCCTCTAGCGGGGCGGAAATGATCCGGTGCCACGTGGTGTTGGAGAGGGGGCAATCTGGCGAGTGGGGACGAGGGTATGAGGTGGTGTGTGGCGCTAGTGATGTTGGGTGCGATCATACCAGCACTAATGCACCGGATCCCATCAGAACTCCGAAGTTAAGCGTGCTTGGGCGAGAGTAGTACTAGGATGGGTGACCTCCTGGGAAGTCCTCGTGTTGCACCCCTTTTGCCTGCCGCTCTCCCCGCAGCTTGCAATTGATGTCGCTGCCTCTGAGCTTCGTCGTGATCTCCTCACCTAGGGCGAGGGCCATCTTCCGGAGGCCCTTCCTCTGGTCAGTTCATCCCCCTCTTTCCCTTCCTCCTCCGAACGCCCCGTCCGCCACACGCGAACTGGTTGCCGCACTACCATTTTGGGTATGCTGGGTCTTTTTTGGAGCTCAGTGGGTGCGGCTTCCCTTGCAAAAGGTTTGCCTCTTTTATGTTGGCTAGCTAGCTAGTTGGGGTTTGGGCCTATTGGGCCCTTTTAGAGGGGGCACCGCCCTCATAAAAATTGCTTGCAATATTCAAATAACTTGGGCATGAAACGCGACGGGAGGAAAAGAGAAAATAGGGCTATAAAGTGTGGAATATTAAAGGAGGTTAGAGGAAGATATTAAATCGGAGGATGCGGATAGAAGATAGAGAAAGTCACGCTAGAAATATTTCCATTTCGGAGTAGAAGCATGAATGGGAATGCGTGAGTGTTGATATCTATGCATAGATATCTAAGAGAGAGAGAGAGAGAATTTAGAGTGGAGCTATCCACTTTGAGGCTCTAGCTATTTCTAACCCTTAATTAGTTCACATATGGGTGACTTTATCATCAACTGATGTGGCCCCATCACTCGCCGCAAACCATCTCTTGAGCCCCCTCCTCTTCAGGGCGAATTTCGTCGCCGCCCCGAAGCGTTCTCGTGGGCTCCCTACGTCTGGCCTTTGACCGATCCGTGGATGTCTTGCCACCCTCCCACGCGCAGAGTAGCTGCCTTTCGGGAGCGTTTTGGGCAGGTCTTGGGCACGTAGAGTGGACCTGAGAGTTGATGGGGACTTGAGGTGGGCTACCTGCTCGTCCTATGCACGATTCCCCCTGGCGAGGGACGCGCCGCCTTGGGGGTCGGTCATGTCACCGGCCCTCTCCCTCTAGCGGGGCGGAAATGATCCGGTGCCACGTGGTGTTGGAGAGGGGGCAATCTGGCGAGTGGGGACGAGGGTATGAGGTGGTGTGTGGCGCTAGTGATGTTGGGTGCGATCATACCAGCACTAATGCACCGGATCCCATCAGAACTCCGAAGTTAAGCGTGCTTGGGCGAGAGTAGTACTAGGATGGGTGACCTCCTGGGAAGTCCTCGTGTTGCACCCCTTTTGCCTGCCGCTCTCCCCGCAGCTTGCAATTGATGTCGCTGCCTCTGAGCTTCGTCGTGATCTCCTCACCTAGGGCGAGGGCCATCTTCCGGAGGCCCTTCCTCTGGTCAGTTCATCCCCCTCTTTCCCTTCCTCCTCCGAACGCCCCGTCCGCCACACGCGAACTGGTTGCCGCACTACCATTTTGGGTATGCTGGGTCTTTTTTGGAGCTCAGTGGGTGCGGCTTCCCTTGCAAAAGGTTTGCCTCTTTTATGTTGGCTAGCTAGCTAGTTGGGGTTTGGGCCTATTGGGCCCTTTTAGAGGGGGCACCGCCCTCATAAAAATTGCTTGCAATATTCAAATAACTTGGGCATGAAACGCGACGGGAGGAAAAGAGAAAATAGGGCTATAAAGTGTGGAATATTAAAGGAGGTTAGAGGAAGATATTAAATCGGAGGATGCGGATAGAAGATAGAGAAAGTCACGCTAGAAATATTTCCATTTCGGAGTAGAAGCATGAATGGGAATGCGTGAGTGTTGATATCTATGCATAGATATCTAAGAGAGAGAGAGAGAGAATTTAGAGTGGAGCTATCCACTTTGAGGCTCTAGCTATTTCTAACCCTTAATTAGTTCACATATGGGTGACTTTATCATCAACTGATGTGGCCCCATCACTCGCCGCAAACCATCTCTTGAGCCCCCTCCTCTTCAGGGCGAATTTCGTCGCCGCCCCGAAGCGTTCTCGTGGGCTCCCTACGTCTGGCCTTTGACCGATCCGTGGATGTCTTGCCACCCTCCCACGCGCAGAGTAGCTGCCTTTCGGGAGCGTTTTGGGCAGGTCTTGGGCACGTAGAGTGGACCTGAGAGTTGATGGGGACTTGAGGTGGGCTACCTGCTCGTCCTATGCACGATTCCCCCTGGCGAGGGACGCGCCGCCTTGGGGGTCGGTCATGTCACCGGCCCTCTCCCTCTAGCGGGGCGGAAATGATCCGGTGCCACGTGGTGTTGGAGAGGGGGCAATCTGGCGAGTGGGGACGAGGGTATGAGGTGGTGTGTGGCGCTAGTGATGTTGGGTGCGATCATACCAGCACTAATGCACCGGATCCCATCAGAACTCCGAAGTTAAGCGTGCTTGGGCGAGAGTAGTACTAGGATGGGTGACCTCCTGGGAAGTCCTCGTGTTGCACCCCTTTTGCCTGCCGCTCTCCCCGCAGCTTGCAATTGATGTCGCTGCCTCTGAGCTTCGTCGTGATCTCCTCACCTAGGGCGAGGGCCATCTTCCGGAGGCCCTTCCTCTGGTCAGTTCATCCCCCTCTTTCCCTTCCTCCTCCGAACGCCCCGTCCGCCACACGCGAACTGGTTGCCGCACTACCATTTTGGGTATGCTGGGTCTTTTTTGGAGCTCAGTGGGTGCGGCTTCCCTTGCAAAAGGTTTGCCTCTTTTATGTTGGCTAGCTAGCTAGTTGGGGTTTGGGCCTATTGGGCCCTTTTAGAGGGGGCACCGCCCTCATAAAAATTGCTTGCAATATTCAAATAACTTGGGCATGAAACGCGACGGGAGGAAAAGAGAAAATAGGGCTATAAAGTGTGGAATATTAAAGGAGGTTAGAGGAAGATATTAAATCGGAGGATGCGGATAGAAGATAGAGAAAGTCACGCTAGAAATATTTCCATTTCGGAGTAGAAGCATGAATGGGAATGCGTGAGTGTTGATATCTATGCATAGATATCTAAGAGAGAGAGAGAGAGAATTTAGAGTGGAGCTATCCACTTTGAGGCTCTAGCTATTTCTAACCCTTAATTAGTTCACATATGGGTGACTTTATCATCAACTGATGTGGCCCCATCACTCGCCGCAAACCATCTCTTGAGCCCCCTCCTCTTCAGGGCGAATTTCGTCGCCGCCCCGAAGCGTTCTCGTGGGCTCCCTACGTCTGGCCTTTGACCGATCCGTGGATGTCTTGCCACCCTCCCACGCGCAGAGTAGCTGCCTTTCGGGAGCGTTTTGGGCAGGTCTTGGGCACGTAGAGTGGACCTGAGAGTTGATGGGGACTTGAGGTGGGCTACCTGCTCGTCCTATGCACGATTCCCCCTGGCGAGGGACGCGCCGCCTTGGGGGTCGGTCATGTCACCGGCCCTCTCCCTCTAGCGGGGCGGAAATGATCCGGTGCCACGTGGTGTTGGAGAGGGGGCAATCTGGCGAGTGGGGACGAGGGTATGAGGTGGTGTGTGGCGCTAGTGATGTTGGGTGCGATCATACCAGCACTAATGCACCGGATCCCATCAGAACTCCGAAGTTAAGCGTGCTTGGGCGAGAGTAGTACTAGGATGGGTGACCTCCTGGGAAGTCCTCGTGTTGCACCCCTTTTGCCTGCCGCTCTCCCCGCAGCTTGCAATTGATGTCGCTGCCTCTGAGCTTCGTCGTGATCTCCTCACCTAGGGCGAGGGCCATCTTCCGGAGGCCCTTCCTCTGGTCAGTTCATCCCCCTCTTTCCCTTCCTCCTCCGAACGCCCCGTCCGCCACACGCGAACTGGTTGCCGCACTACCATTTTGGGTATGCTGGGTCTTTTTTGGAGCTCAGTGGGTGCGGCTTCCCTTGCAAAAAGTTTGCCTCTTTTATGTTGGCTAGCTAGCTAGTTGGGGTTTGGGCCTATTGGGCCCTTTTAGAGGGGGCACCGCCCTCATAAAAATTGCTTGCAATATTCAAATAACTTGGGCATGAAACGCGACGGGAGGAAAAGAGAAAATAGGGCTATAAAGTGTGGAATATTAAAGGAGGTTAGAGGAAGATATTAAATCGGAGGATGCGGATAGAAGATAGAGAAAGTCACGCTAGAAATATTTCCATTTCGGAGTAGAAGCATGAATGGGAATGCGTGAGTGTTGATATCTATGCATAGATATCTAAGAGAGAGAGAGAGAGAATTTAGAGTGGAGCTATCCACTTTGAGGCTCTAGCTATTTCTAACCCTTAATTAGTTCACATATGGGTGACTTTATCATCAACTGATGTGGCCCCATCACTCGCCGCAAACCATCTCTTGAGCCCCCTCCTCTTCAGGGCGAATTTCGTCGCCGCCCCGAAGCGTTCTCGTGGGCTCCCTACGTCTGGCCTTTGACCGATCCGTGGATGTCTTGCCACCCTCCCACGCGCAGAGTAGCTGCCTTTCGGGAGCGTTTTGGGCAGGTCTTGGGCACGTAGAGTGGACCTGAGAGTTGATGGGGACTTGAGGTGGGCTACCTGCTCGTCCTATGCACGATTCCCCCTGGCGAGGGACGCGCCGCCTTGGGGGTCGGTCATGTCACCGGCCCTCTCCCTCTAGCGGGGCGGAAATGATCCGGTGCCACGTGGTGTTGGAGAGGGGGCAATCTGGCGAGTGGGGACGAGGGTATGAGGTGGTGTGTGGCGCTAGTGATGTTGGGTGCGATCATACCAGCACTAATGCACCGGATCCCATCAGAACTCCGAAGTTAAGCGTGCTTGGGCGAGAGTAGTACTAGGATGGGTGACCTCCTGGGAAGTCCTCGTGTTGCACCCCTTTTGCCTGCCGCTCTCCCCGCAGCTTGCAATTGATGTCGCTGCCTCTGAGCTTCGTCGTGATCTCCTCACCTAGGGCGAGGGCCATCTTCCGGAGGCCCTTCCTCTGGTCAGTTCATCCCCCTCTTTCCCTTCCTCCTCCGAACGCCCCGTCCGCCACACGCGAACTGGTTGCCGCACTACCATTTTGGGTATGCTGGGTCTTTTTTGGAGCTCAGTGGGTGCGGCTTCCCTTGCAAAAGGTTTGCCTCTTTTATGTTGGCTAGCTAGCTAGTTGGGGTTTGGGCCTATTGGGCCCTTTTAGAGGGGGCACCGCCCTCATAAAAATTGCTTGCAATATTCAAATAACTTGGGCATGAAACGCGACGGGAGGAAAAGAGAAAATAGGGCTATAAAGTGTGGAATATTAAAGGAGGTTAGAGGAAGATATTAAATCGGAGGATGCGGATAGAAGATAGAGAAAGTCACGCTAGAAATATTTCCATTTCGGAGTAGAAGCATGAATGGGAATGCGTGAGTGTTGATATCTATGCATAGATATCTAAGAGAGAGAGAGAGAGAATTTAGAGTGGAGCTATCCACTTTGAGGCTCTAGCTATTTCTAACCCTTAATTAGTTCACATATGGGTGACTTTATCATCAACTGATGTGGCCCCATCACTCGCCGCAAACCATCTCTTGAGCCCCCTCCTCTTCAGGGCGAATTTCGTCGCCGCCCCGAAGCGTTCTCGTGGGCTCCCTACGTCTGGCCTTTGACCGATCCGTGGATGTCTTGCCACCCTCCCACGCGCAGAGTAGCTGCCTTTCGGGAGCGTTTTGGGCAGGTCTTGGGCACGTAGAGTGGACCTGAGAGTTGATGGGGACTTGAGGTGGGCTACCTGCTCGTCCTATGCACGATTCCCCCTGGCGAGGGACGCGCCGCCTTGGGGGTCGGTCATGTCACCGGCCCTCTCCCTCTAGCGGGGCGGAAATGATCCGGTGCCACGTGGTGTTGGAGAGGGGGCAATCTGGCGAGTGGGGACGAGGGTATGAGGTGGTGTGTGGCGCTAGTGATGTTGGGTGCGATCATACCAGCACTAATGCACCGGATCCCATCAGAACTCCGAAGTTAAGCGTGCTTGGGCGAGAGTAGTACTAGGATGGGTGACCTCCTGGGAAGTCCTCGTGTTGCACCCCTTTTGCCTGCCGCTCTCCCCGCAGCTTGCAATTGATGTCGCTGCCTCTGAGCTTCGTCGTGATCTCCTCACCTAGGGCGAGGGCCATCTTCCGGAGGCCCTTCCTCTGGTCAGTTCATCCCCCTCTTTCCCTTCCTCCTCCGAACGCCCCGTCCGCCACACGCGAACTGGTTGCCGCACTACCATTTTGGGTATGCTGGGTCTTTTTTGGAGCTCAGTGGGTGCGGCTTCCCTTGCAAAAGGTTTGCCTCTTTTATGTTGGCTAGCTAGCTAGTTGGGGTTTGGGCCTATTGGGCCCTTTTAGAGGGGGCACCGCCCTCATAAAAATTGCTTGCAATATTCAAATAACTTGGGCATGAAACGCGACGGGAGGAAAAGAGAAAATAGGGCTATAAAGTGTGGAATATTAAAGGAGGTTAGAGGAAGATATTAAATCGGAGGATGCGGATAGAAGATAGAGAAAGTCACGCTAGAAATATTTCCATTTCGGAGTAGAAGCATGAATGGGAATGCGTGAGTGTTGATATCTATGCATAGATATCTAAGAGAGAGAGAGAGAGAATTTAGAGTGGAGCTATCCACTTTGAGGCTCTAGCTATTTCTAACCCTTAATTAGTTCACATATGGGTGACTTTATCATCAACTGATGTGGCCCCATCACTCGCCGCAAACCATCTCTTGAGCCCCCTCCTCTTCAGGGCGAATTTCGTCGCCGCCCCGAAGCGTTCTCGTGGGCTCCCTACGTCTGGCCTTTGACCGATCCGTGGATGTCTTGCCACCCTCCCACGCGCAGAGTAGCTGCCTTTCGGGAGCGTTTTGGGCAGGTCTTGGGCACGTAGAGTGGACCTGAGAGTTGATGGGGACTTGAGGTGGGCTACCTGCTCGTCCTATGCACGATTCCCCCTGGCGAGGGACGCGCCGCCTTGGGGGTCGGTCATGTCACCGGCCCTCTCCCTCTAGCGGGGCGGAAATGATCCGGTGCCACGTGGTGTTGGAGAGGGGGCAATCTGGCGAGTGGGGACAAGGGTATGAGGTGGTGTGTGGCGCTAGTGATGTTGGGTGCGATCATACCAGCACTAATGCACCGGATCCCATCAGAACTCCGAAGTTAAGCGTGCTTGGGCGAGAGTAGTACTAGGATGGGTGACCTCCTGGGAAGTCCTCGTGTTGCACCCCTTTTGCCTGCCGCTCTCCCCGCAGCTTGCAATTGATGTCGCTGCCTCTGAGCTTCGTCGTGATCTCCTCACCTAGGGCGAGGGCCATCTTCCGGAGGCCCTTCCTCTGGTCAGTTCATCCCCCTCTTTCCCTTCCTCCTCCGAACGCCCCGTCCGCCACACGCGAACTGGTTGCCGCACTACCATTTTGGGTATGCTGGGTCTTTTTTGGAGCTCAGTGGGTGCGGCTTCCCTTGCAAAAGGTTTGCCTCTTTTATGTTGGCTAGCTAGCTAGTTGGGGTTTGGGCCTATTGGGCCCTTTTAGAGGGGGCACCGCCCTCATAAAAATTGCTTGCAATATTCAAATAACTTGGGCATGAAACGCGACGGGAGGAAAAGAGAAAATAGGGCTATAAAGTGTGGAATATTAAAGGAGGTTAGAGGAAGATATTAAATCGGAGGATGCGGATAGAAGATAGAGAAAGTCACGCTAGAAATATTTCCATTTCGGAGTAGAAGCATGAATGGGAATGCGTGAGTGTTGATATCTATGCATAGATATCTAAGAGAGAGAGAGAGAGAATTTAGAGTGGAGCTATCCACTTTGAGGCTCTAGCTATTTCTAACCCTTAATTAGTTCACATATGGGTGACTTTATCATCAACTGATGTGGCCCCATCACTCGCCGCAAACCATCTCTTGAGCCCCCTCCTCTTCAGGGCGAATTTCGTCGCCGCCCCGAAGCGTTCTCGTGGGCTCCCTACGTCTGGCCTTTGACCGATCCGTGGATGTCTTGCCACCCTCCCACGCGCAGAGTAGCTGCCTTTCGGGAGCGTTTTGGGCAGGTCTTGGGCACGTAGAGTGGACCTGAGAGTTGATGGGGACTTGAGGTGGGCTACCTGCTCGTCCTATGCACGATTCCCCCTGGCGAGGGACGCGCCGCCTTGGGGGTCGGTCATGTCACCGGCCCTCTCCCTCTAGCGGGGCGGAAATGATCCGGTGCCACGTGGTGTTGGAGAGGGGGCAATCTGGCGAGTGGGGACGAGGGTATGAGGTGGTGTGTGGCGCTAGTGATGTTGGGTGCGATCATACCAGCACTAATGCACCGGATCCCATCAGAACTCCGAAGTTAAGCGTGCTTGGGCGAGAGTAGTACTAGGATGGGTGACCTCCTGGGAAGTCCTCGTGTTGCACCCCTTTTGCCTGCCGCTCTCCCCGCAGCTTGCAATTGATGTCGCTGCCTCTGAGCTTCGTCGTGATCTCCTCACCTAGGGCGAGGGCCATCTTCCGGAGGCCCTTCCTCTGGTCAGTTCATCCCCCTCTTTCCCTTCCTCCTCCGAACGCCCCGTCCGCCACACGCGAACTGGTTGCCGCACTACCATTTTGGGTATGCTGGGTCTTTTTTGGAGCTCAGTGGGTGCGGCTTCCCTTGCAAAAGGTTTGCCTCTTTTATGTTGGCTAGCTAGCTAGTTGGGGTTTGGGCCTATTGGGCCCTTTTAGAGGGGGCACCGCCCTCATAAAAATTGCTTGCAATATTCAAATAACTTGGGCATGAAACGCGACGGGAGGAAAAGAGAAAATAGGGCTATAAAGTGTGGAATATTAAAGGAGGTTAGAGGAAGATATTAAATCGGAGGATGCGGATAGAAGATAGAGAAAGTCACGCTAGAAATATTTCCATTTCGGAGTAGAAGCATGAATGGGAATGCGTGAGTGTTGATATCTATGCATAGATATCTAAGAGAGAGAGAGAGAGAATTTAGAGTGGAGCTATCCACTTTGAGGCTCTAGCTATTTCTAACCCTTAATTAGTTCACATATGGGTGACTTTATCATCAACTGATGTGGCCCCATCACTCGCCGCAAACCATCTCTTGAGCCCCCTCCTCTTCAGGGCGAATTTCGCCGCCGCCCCGAAGCGTTCTCGTGGGCTCCCTACGTCTGGCCTTTGACCGATCCGTGGATGTCTTGCCACCCTCCCACGCGCAGAGTAGCTGCCTTTCGGGAGCGTTTTGGGCAGGTCTTGGGCACGTAGAGTGGACCTGAGAGTTGATGGGGACTTGAGGTGGGCTACCTGCTCGTCCTATGCACGATTCCCCCTGGCGAGGGACGCGCCGCCTTGGGGGTCGGTCATGTCACCGGCCCTCTCCCTCTAGCGGGGCGGAAATGATCCGGTGCCACGTGGTGTTGGAGAGGGGGCAATCTGGCGAGTGGGGACGAGGGTATGAGGTGGTGTGTGGCGCTAGTGATGTTGGGTGCGATCATACCAGCACTAATGCACCGGATCCCATCAGAACTCCGAAGTTAAGCGTGCTTGGGCGAGAGTAGTACTAGGATGGGTGACCTCCTGGGAAGTCCTCGTGTTGCACCCCTTTTGCCTGCCGCTCTCCCCGCAGCTTGCAATTGATGTCGCTGCCTCTGAGCTTCGTCGTGATCTCCTCACCTAGGGCGAGGGCCATCTTCCGGAGGCCCTTCCTCTGGTCAGTTCATCCCCCTCTTTCCCTTCCTCCTCCGAACGCCCCGTCCGCCACACGCGAACTGGTTGCCGCACTACCATTTTGGGTATGCTGGGTCTTTTTTGGAGCTCAGTGGGTGCGGCTTCCCTTGCAAAAGGTTTGCCTCTTTTATGTTGGCTAGCTAGCTAGTTGGGGTTTGGGCCTATTGGGCCCTTTTAGAGGGGGCACCGCCCTCATAAAAATTGCTTGCAATATTCAAATAACTTGGGCATGAAACGCGACGGGAGGAAAAGAGAAAATAGGGCTATAAAGTGTGGAATATTAAAGGAGGTTAGAGGAAGATATTAAATCGGAGGATGCGGATAGAAGATAGAGAAAGTCACGCTAGAAATATTTCCATTTCGGAGTAGAAGCATGAATGGGAATGCGTGAGTGTTGATATCTATGCATAGATATCTAAGAGAGAGAGAGAGAGAATTTAGAGTGGAGCTATCCACTTTGAGGCTCTAGCTATTTCTAACCCTTAATTAGTTCACATATGGGTGACTTTATCATCAACTGATGTGGCCCCATCACTCGCCGCAAACCATCTCTTGAGCCCCCTCCTCTTCAGGGCGAATTTCGTCGCCGCCCCGAAGCGTTCTCGTGGGCTCCCTACGTCTGGCCTTTGACCGATCCGTGGATGTCTTGCCACCCTCCCACGCGCAGAGTAGCTGCCTTTCGGGAGCGTTTTGGGCAGGTCTTGGGCACGTAGAGTGGACCTGAGAGTTGATGGGGACTTGAGGTGGGCTACCTGCTCGTCCTATGCACGATTCCCCCTGGCGAGGGACGCGCCGCCTTGGGGGTCGGTCATGTCACCGGCCCTCTCCCTCTAGCGGGGCGGAAATGATCCGGTGCCACGTGGTATTGGAGAGGGGGCAATCTGGCGAGTGGGGACGAGGGTATGAGGTGGTGTGTGGCGCTAGTGATGtTGGGTGCGATCATACCAGCACTAATGCACCGGATCCCATCAGAACTCCGAAGTTAAGCGTGCTTGGGCGAGAGTAGTACTAGGATGGGTGACCTCCTGGGAAGTCCTCGTGTTGCACCCCTTTTGCCTGCCGCTCTCCCCGCAGCTTGCAATTGATGTCGCTGCCTCTGAGCTTCGTCGTGATCTCCTCACCTAGGGCGAGGGCCATCTTCCGGAGGCCCTTCCTCTGGTCAGTTCATCCCCCTCTTTCCCTTCCTCCTCCGAACGCCCCGTCCGCCACACGCGAACTGGTTGCCGCACTACCATTTTGGGTATGCTGGGTCTTTTTTGGAGCTCATTGGGTGCGGCTTCCCTTGCAAAAGGTTTGCCTCTTTTATGTTGGCTAGCTAGCTAGTTGGGGTTTGGGCCTATTGGGCCCTTTTAGAGGGGGCACCGCCCTCATAAAAATTGCTTGCAATATTCAAATAACTTGGGCATGAAACGCGACGGGAGGAAAAGAGAAAATAGGGCTATAAAGTGTGGAATATTAAAGGAGGTTAGAGGAAGATATTAAATCGGAGGATGCGGATAGAAGATAGAGAAAGTCACGCTAGAAATATTTCCATTTCGGAGTAGAAGCATGAATGGGAATGCGTGAGTGTTGATATCTATGCATAGATATCTAAGAGAGAGAGAGAGAGAATTTAGAGTGGAGCTATCCACTTTGAGGCTCTAGCTATTTCTAACCCTTAATTAGTTCACATATGGGTGACTTTATCATCAACTGATGTGGCCCCATCACTCGCCGCAAACCATCTCTTGAGCCCCCTCCTCTTCAGGGCGAACTTCGTCGCCGCCCCGAAGCGTTCTCGTGGGCTCCCTACGTCTGGCCTTTGACCGATCCGTGGATGTCTTGCCACCCTCCCACGCGCAGAGTAGCTGCCTTTCGGGAGCGTTTTGGGCAGGTCTTGGGCACGTAGAGTGGACCTGAGAGTTGATGGGGACTTGAGGTGGGCTACCTGCTCGTCCTATGCACGATTCCCCCGGGCGAGGGACGCGCCGCCTTGGGGGTCGGTCATGTCACCGGCCCTCTCCCTCGAGCGGGGCGGAAATGATCCGGTGCCACGTGGTGTTGGAGAGGGGGCAATCTGGCGAGTGGGGACGAGGGTATGAGGTGGTGTGTGGCGCTAGTGATGTTGGGTGCGATCATACCAGCACTAATGCACCGGATCCCATCAGAACTCCGAAGTTAAGCGTGCTTGGGCGAGAGTAGTACTAGGATGGGTGACCTCCTGGGAAGTCCTCGTGTTGCACCCCTTTTGCCTGCCGCTCTCCCCGCAGCCTGCAATTGATGTCGCTGCCTCTGAGCTTCGTCGTGATCTCCTCACCTAGGGCGAGGGCCATCTTCCGGAGGCCCTTCCTCTGGTCAGTTCATCCCCCTCTTTCCCTTCCTCCTCCGAACGCCCCGTCCGCCACACGCGAACTGGTTGCCGCACTACCATTTTGGGTATGCTGGGTCTTTTTTGGAGCTCAGTGGGTGCGGCTTCCCTTGCAAAAGGTTTGCCTCTTTTATGTTGGCTAGCTAGCTAGTTGGGGTTTGGGCCTATTGGGCCCTTTTAGAGGGGGCACCGCCCTCATAAAAATTGCTTGCAATATTCAAATAACTTGGGCATGAAACGCGACGGGAGGAAAAGAGAAAATAGGGCTATAAAGTGTGGAATATTAAAGGAGGTTAGAGGAAGATATTAAATCGGAGGATGCGGATAGAAGATAGAGAAAGTCACGCTAGAAATATTTCCATTTCGGAGTAGAAGCATGAATGGGAATGCGTGAGTGTTGATATCTATGCATAGATATCTAAGAGAGAGAGAGAGAGAATTTAGAGTGGAGCTATCCACTTTGAGGCTCTAGCTATTTCTAACCCTTAATTAGTTCACATATGGGTGACTTTATCATCAACTGATGTGGCCCCATCACTCGCCGCAAACCATCTCTTGAGCCCCCTCCTCTTCAGGGCGAATTTCGTCGCCGCCCCGAAGCGTTCTCGTGGGCTCCCTACGTCTGGCCTTTGACCGATCCGTGGATGTCTTGCCACCCTCCCACGCGCAGAGTAGCTGCCTTTCGGGAGCGTTTTGGGCAGGTCTTGGGCACGTAGAGTGGACCTGAGAGTTGATGGGGACTTGAGGTGGGCTACCTGCTCGTCCTATGCACGATTCCCCCTGGCGAGGGACGCGCCGCCTTGGGGGTCGGTCATGTCACCGGCCCTCTCCCTCTAGCGGGGCGGAAATGATCCGGTGCCACGTGGTGTTGGAGAGGGGGCAATCTGGCGAGTGGGGACGAGGGTATGAGGTGGTGTGTGGCGCTAGTGATGTTGGGTGCGATCATACCAGCACTAATGCACCGGATCCCATCAGAACTCCGAAGTTAAGCGTGCTTGGGCGAGAGTAGTACTAGGATGGGTGACCTCCTGGGAAGTCCTCGTGTTGCACCCCTTTTGCCTGCCGCTCTCCCCGCAGCTTGCAATTGATGTCGCTGCCTCTGAGCTTCGTCGTGATCTCCTCACCTAGGGCGAGGGCCATCTTCCGGAGGCCCTTCCTCTGGTCAGTTCATCCCCCTCTTTCCCTTCCTCCTCCGAACGCCCCGTCCGCCACACGCGAACTGGTTGCCGCACTACCATTTTGGGTATGCTGGGTCTTTTTTGGAGCTCAGTGGGTGCGGCTTCCCTTGCAAAAGGTTTGCCTCTTTTATGTTGGCTAGCTAGCTAGTTGGGGTTTGGGCCTATTGGGCCCTTTTAGAGGGGGCACCGCCCTCATAAAAATTGCTTGCAATATTCAAATAACTTGGGCATGAAACGCGACGGGAGGAAAAGAGAAAATAGGGCTATAAAGTGTGGAATATTAAAGGAGGTTAGAGGAAGATATTAAATCGGAGGATGCGGATAGAAGATAGAGAAAGTCACGCTAGAAATATTTCCATTTCGGAGTAGAAGCATGAATGGGAATGCGTGAGTGTTGATATCTATGCATAGATATCTAAGAGAGAGAGAGAGAGAATTTAGAGTGGAGCTATCCACTTTGAGGCTCTAGCTATTTCTAACCCTTAATTAGTTCACATATGGGTGACTTTATCATCAACTGATGTGGCCCCATCACTCGCCGCAAACCATCTCTTGAGCCCCCTCCTCTTCAGGGCGAATTTCGTCGCCGCCCCGAAGCGTTCTCGTGGGCTCCCTACGTCTGGCCTTTGACCGATCCGTGGATGTCTTGCCACCCTCCCACGCGCAGAGTAGCTGCCTTTCGGGAGCGTTTTGGGCAGGTCTTGGGCACGTAGAGTGGACCTGAGAGTTGATGGGGACTTGAGGTGGGCTACCTGCTCGTCCTATGCACGATTCCCCCTGGCGAGGGACGCGCCGCCTTGGGGGTCGGTCATGTCACCGGCCCTCTCCCTCTAGCGGGGCGGAAATGATCCGGTGCCACGTGGTGTTGGAGAGGGGGCAATCTGGCGAGTGGGGACGAGGGTATGAGGTGGTGTGTGGCGCTAGTGATGTTGGGTGCGATCATACCAGCACTAATGCACCGGATCCCATCAGAACTCCGAAGTTAAGCGTGCTTGGGCGAGAGTAGTACTAGGATGGGTGACCTCCTGGGAAGTCCTCGTGTTGCACCCCTTTTGCCTGCCGCTCTCCCCGCAGCTTGCAATTGATGTCGCTGCCTCTGAGCTTCGTCGTGATCTCCTCACCTAGGGCGAGGGCCATCTTCCGGAGGCCCTTCCTCTGGTCAGTTCATCCCCCTCTTTCCCTTCCTCCTCCGAACGCCTCGTCCGCCACACGCGAACTGGTTGCCGCACTACCATTTTGGGTATGCTGGGTCTTTTTTGGAGCTCAGTGGGTGCGGCTTCCCTTGCAAAAGGTTTGCCTCTTTTATGTTGGCTAGCTAGCTAGTTGGGGTTTGGGCCTATTGGGCCCTTTTAGAGGGGGCACCGCCCTCATAAAAATTGCTTGCAATATTCAAATAACTTGGGCATGAAACGCGACGGGAGGAAAAGAGAAAATAGGGCTATAAAGTGTGGAATATTAAAGGAGGTTAGAGGAAGATATTAAATCGGAGGATGCGGATAGAAGATAGAGTAAGTCACGCTAGAAATATTTCCATTTCGGAGTAGAAGCATGAATGGGAATGCGTGAGTGTTGATATCTATGCATAGATATCTAAGAGAGAGAGAGAGAGAATTTAGAGTGGAGCTATCCACTTTGAGGCTCTAGCTATTTCTAACCCTTAATTAGTTCACATATGGGTGACTTTATCATCAACTGATGTGGCCCCATCACTCGCCGCAAACCATCTCTTGAGCCCCCTCCTCTTCAGGGCGAATTTCGTCGCCGCCCCGAAGCGTTCTCGTGGGCTCCCTACGTCTGGCCTTTGACCGATCCGTGGATGTCTTGCCACCCTCCCACGCGCAGAGTAGCTGCCTTTCGGGAGCGTTTTGGGCGGGTCTTGGGCACGTAGAGTGGACCTGAGAGTTGATGGGGACTTGAGGTGGGCTACCTGCTCGTCCTATGCACGATTCCCCCTGGCGAGGGACGCGCCGCCTTGGGGGTCGGTCATGTCACCGGCCCTCTCCCTCTAGCGGGGCGGAAATGATCCGGTGCCACGTGGTGTTGGAGAGGGGGCAATCTGGCGAGTGGGGACGAGGGTATGAGGTGGTGTGTGGCGCTAGTGATGTTGGGTGCGATCATACCAGCACTAATGCACCGGATCCCATCAGAACTCCGAAGTTAAGCGTGCTTGGGCGAGAGTAGTACTAGGATGGGTGACCTCCTGGGAAGTCCTCGTGTTGCACCCCTTTTGCCTGCCGCTCTCCCCGCAGCTTGCAATTGATGTCGCTGCCTCTGAGCTTCGTCGTGATCTCCTCACCTAGGGCGAGGGCCATCTTCCGGAGGCCCTTCCTCTGGTCAGTTCATCCCCCTCTTTCCCTTCCTCCTCCGAACGCCTCGTCCGCCACACGCGAACTGGTTGCCGCACTACCATTTTGGGTATGCTGGGTCTTTTTTGGAGCTCAGTGGGTGCGGCTTCCCTTGCAAAAGGTTTGCCTCTTTTATGTTGGCTAGCTAGCTAGTTGGGGTTTGGGCCTATTGGGCCCTTTTAGAGGGGGCACCGCCCTCATAAAAATTGCTTGCAATATTCAAATAACTTGGGCATGAAACGCGACGGGAGGAAAAGAGAAAATAGGGCTATAAAGTGTGGAGTATTAAAGGAGGTTAGAGGAAGATATTAAATCGGAGGATGCGGATAGAAGATAGAGAAAGTCACGCTAGAAATATTTCCATTTCGGAGTAGAAGCATGAATGGGAATGCGTGAGTGTTGATATCTATGCATAGATATCTAAGAGAGAGAGAGGGAGAATTTAGAGTGGAGCTATCCACTTTGAGGCTCTAGCTATTTCTAACCCTTAATTAGTTCACATATGGGTGACTTTATCATCAACTGATGTGGCCCCATCACTCGCCGCAAACCATCTCTTGAGCCCCCTCCTCTTCAGGGCGAATTTCGTCGCCGCCCCGAAGCGTTCTCGTGGGCTCCCTACGTCTGGCCTTTGACCGATCCGTGGATGTCTTGCCACCCTCCTACGCGCAGAGTAGCTGCCTTTCGGGAGCGTTTTGGGCAGGTCTTGGGCACGTAGAGTGGACCTGAGAGTTGATGGGGACTTGAGGTGGGCTACCTGCTCGTCCTATGCACGATTCCCCCTGGCGAGGGACGCGCCGCCTTGGGGGTCGGTCATGTCACCGGCCCTCTCCCTCTAGCGGGGCGGAAATGATCCGGTGCCACGTGGTGTTGGAGAGGGGGCAATCTGGCGAGTGGGGACGAGGGTATGAGGTGGTGTGTGGCGCTAGTGATGTTGGGTGCGATCATACCAGCACTAATGCACCGGATCCCATCAGAACTCCGAAGTTAAGCGTGCTTGGGCGAGAGTAGTACTAGGATGGGTGACCTCCTGGGAAGTCCTCGTGTTGCACCCCTTTTGCCTGCCGCTCTCCCCGCAGCTTGCAATTGATGTCGCTGCCTCTGAGCTTCGTCGTGATCTCCTCACCTAGGGCGAGGGCCATCTTCCGGAGGCCCTTCCTCTGGTCAGTTCATCCCCCTCTTTCCCTTCCTCCTCCGAACGCCTCGTCCGCCACACGCGAACTGGTTGCCGCACTACCATTTTGGGTATGCTGGGTCTTTTTTGGAGCTCAGTGGGTGCGGCTTCCCTTGCAAAAGGTTTGCCTCTTTTATGTTGGCTAGCTAGCTAGTTGGGGTTTGGGCCTATTGGGCCCTTTTAGAGGGGGCACCGCCCTCATAAAAATTGCTTGCAATATTCAAATAACTTGGGCATGAAACGCGACGGGAGGAAAAGAGAAAATAGGGCTATAAAGTGTGGAATATTAAAGGAGGTTAGAGGAAGATATTAAATCGGAGGATGCGGATAGAAGATAGAGAAAGTCACGCTAGAAATATTTCCATTTCGGAGTAGAAGCATGAATGGGAATGCGTGAGTGTTGATATCTATGCATAGATATCTAAGAGAGAGAGAGAGAGAATTTAGAGTGGAGCTATCCACTTTGAGGCTCTAGCTATTTCTAACCCTTAATTAGTTCACATATGGGTGACTTTATCATCAACTGATGTGGCCCCATCACTCGCCGCAAACCATCTCTTGAGCCCCCTCCTCTTCAGGGCGAATTTCGTCGCCGCCCCGAAGCGTTCTCGTGGGCTCCCTACGTCTGGCCTTTGACCGATCCGTGGATGTCTTGCCACCCTCCCACGCGCAGAGTAGCTGCCTTTCGGGAGCGTTTTGGGCAGGTCTTGGGCACGTAGAGTGGACCTGAGAGTTGATGGGGACTTGAGGTGGGCTACCTGCTCGTCCTATGCACGATTCCCCCTGGCGAGGGACGCGCCGCCTTGGGGGTCGGTCATGTCACCGGCCCTCTCCCTCTAGCGGGGCGGAAATGATCCGGTGCCACGTGGTGTTGGAGAGGGGGCAATCTGGCGAGTGGGGACGAGGGTATGAGGTGGTGTGTGGCGCTAGTGATGTTGGGTGCGATCATACCAGCACTAATGCACCGGATCCCATCAGAACTCCGAAGTTAAGCGTGCTTGGGCGAGAGTAGTACTAGGATGGGTGACCTCCTGGGAAGTCCTCGTGTTGCACCCCTTTTGCCTGCCGCTCTCCCCGCAGCTTGCAATTGATGTCGCTGCCTCTGAGCTTCGTCGTGATCTCCTCACCTAGGGCGAGGGCCATCTTCCGGAGGCCCTTCCTCTGGTCAGTTCATCCCCCTCTTTCCCTTCCTCCTCCGAACGCCTCGTCCGCCACACGCGAACTGGTTGCCGCACTATCATTTTGGGTATGCTGGGTCTTTTTTGGAGCTCAGTGGGTGCGGCTTCCCTTGCAAAAGGTTTGCCTCTTTTATGTTGGCTAGCTAGCTAGTTGGGGTTTGGGCCTATTGGGCCCTTTTAGAGGGGGCACCGCCCTCATAAAAATTGCTTGCAATATTCAAATAACTTGGGCATGAAACGCGACGGGAGGAAAAGAGAAAATAGGGCTATAAAGTGTGGAATATTAAAGGAGGTTAGAGGAAGATATTAAATCGGAGGATGCGGATAGAAGATAGAGAAAGTCACGCTAGAAATATTTCCATTTCGGAGTAGAAGCATGAATGGGAATGCGTGAGTGTTGATATCTATGCATAGATATCTAAGAGAGAGAGAGAGAGAATTTAGAGTGGAGCTATCCACTTTGAGGCTCTAGCTATTTCTAACCCTTAATTAGTTCACATATGGGTGACTTTATCATCAACTGATGTGGCCCCATCACTCGCCGCAAACCATCTCTTGAGCCCCCTCCTCTTCAGGGCGAATTTCGTCGCCGCCCCGAAGCGTTCTCGTGGGCTCCCTACGTCTGGCCTTTGACCGATCCGTGGATGTCTTGCCACCCTCCCACGCGCAGAGTAGCTGCCTTTCGGGAGCGTTTTGGGCAGGTCTTGAGCACGTAGAGTGGACCTGAGAGTTGATGGGGACTTGAGGTGGGCTACCTGCTCGTCCTATCCACGATTCCCCCTGGCGAGGGACGCGCCGCCTTGGGGGTCGGTCATGTCACCGGCCCTCTCCCTCTAGCGGGGCGGAAATGATCCGGTGCCACGTGGTGTTGGAGAGGGGGCAATCTGGCGAGTGGGGACGAGGGTATGAGGTGGTGTGTGGCGCTAGTGATGTTGGGTGCGATCATACCAGCACTAATGCACCGGATCCCATCAGAACTCCGAAGTTAAGCGTGCTTGGGCGAGAGTAGTACTAGGATGGGTGACCTCCTGGGAAGTCCTCGTGTTGCACCCCTTTTGCCTGCCGCTCTCCCCGCAGCTTGCAATTGATGTCGCTGCCTCTGAGCTTCGTCGTGATCTCCTCACCTAGGGCGAGGGCCATCTTCCGGAGGCCCTTCCTCTGGTCAGTTCATCCCCCTCTTTCCCTTCCTCCTCCGAACGCCCCGTCCGCCACACGCGAACTGGTTGCCGCACTACCATTTTGGGTATGCTGGGTCTTTTTTGGAGCTCAGTGGGTGCGGCTTCCCTTGCAAAAGGTTTGCCTCTTTTATGTTGGCTAGCTAGCTAGTTGGGGTTTGGGCCTATTGGGCCCTTTTAGAGGGGGCACCGCCCTCATAAAAATTGCTTGCAATATTCAAATAACTTGGGCATGAAACGCGACGGGAGGAAAAGAGAAAATAGGGCTATAAAGTGTGGAATATTAAAGGAGGTTAGAGGAAGATATTAAATCGGAGGATGCGGATAGAAGATAGAGAAAGTCACGCTAGAAATATTTCCATTTCGGAGTAGAAGCATGAATGGGAATGCGTGAGTGTTGATATCTATGCATAGATATCTAAGAGAGAGAGAGAGAGAATTTAGAGTGGAGCTATCCACTTTGAGGCTCTAGCTATTTCTAACCCTTAATTAGTTCACATATGGGTGACTTTATCATCAACTGATGTGGCCCCATCACTCGCCGCAAACCATCTCTTGAGCCCCCTCCTCTTCAGGGCGAATTTCGTCGCCGCCCCGAAGCGTTCTCGTGGGCTCCCTACGTCTGGCCTTTGACCGATCCGTGGATGTCTTGCCACCCTCCCACGCGCAGAGTAGCTGCCTTTCGGGAGCGTTTTGGGCAGGTCTTGGGCACGTAGAGTGGACCTGAGAGTTGATGGGGACTTGAGGTGGGCTACCTGCTCGTCCTATGCACGATTCCCCCTGGCGAGGGACGCGCCGCCTTGGGGGTCGGTCATGTCACCGGCCCTCTCCCTCTAGCGGGGCGGAAATGATCCGGTGCCACGTGGTGTTGGAGAGGGGGCAATCTGGCGAGTGGGGACGAGGGTATGAGGTGGTGTGTGGCGCTAGTGATGTTGGGTGCGATCATACCAGCACTAATGCACCGGATCCCATCAGAACTCCGAAGTTAAGCGTGCTTGGGCGAGAGTAGTACTAGGATGGGTGACCTCCTGGGAAGTCCTCGTGTTGCACCCCTTTTGCCTGCCGCTCTCCCCGCAGCTTGCAATTGATGTCGCTGCCTCTGAGCTTCGTCGTGATCTCCTCACCTAGGGCGGGGGCCATCTTCCGGAGGCCCTTCCTCTGGTCAGTTCATCCCCCTCTTTCCCTTCCTCCTCCGAACGCCCCGTCCGCCACACGCGAACTGGTTGCCGCACTACCATTTTGGGTCTGCTGGGTCTTTTTTGGAGCTCAGTGGGTGCGGCTTCCCTTGCAAAAGGTTTGCCTCTTTTATGTTGGCTAGCTAGCTAGTTGGGGTTTGGGCCTATTGGGCCCTTTTAGAGGGGGCACCGCCCTCATAAAAATTGCTTGCAATATTCAAATAACTTGGGCATGAAACGCGACGGGAGGAAAAGAGAAAATAGGGCTATAAAGTGTGGAATATTAAAGGAGGTTAGAGGAAGATATTAAATCGGAGGATGCGGATAGAAGATAGAGAAAGTCACGCTAGAAATATTTCCATTTCGGAGTAGAAGCATGAATGGGAATGCGTGAGTGTTGATATCTATGCATAGATATCTAAGAGAGAGAGAGAGAGAATTTAGAGTGGAGCTATCCACTTTGAGGCTCTAGCTATTTCTAACCCTTAATTAGTTCACATATGGGTGACTTTATCATCAACTGATGTGGCCCCATCACTCGCCGCAAACCATCTCTTGAGCCCCCTCCTCTTCAGGGCGAATTTCGTCGCCGCCCCGAAGCGTTCTCGTGGGCTCCCTACGTCTGGCCTTTGACCGATCCGTGGATGTCTTGCCACCCTCCCACGCGCAGAGTAGCTGCCTTTCGGGAGCGTTTTGGGCAGGTCTTGGGCACGTAGAGTGGACCTGAGAGTTGATGGGGACTTGAGGTGGGCTACCTGCTCGTCCTATGCACGATTCCCCCTGGCGAGGGACGCGCCGCCTTGGGGGTCGGTCATGTCACCGGCCCTCTCCCTCTAGCGGGGCGGAAATGATCCGGTGCCACGTGGTGTTGGAGAGGGGGCAATCTGGCGAGTGGGGACGAGGGTATGAGGTGGTGTGTGGCGCTAGTGATGTTGGGTGCGATCATACCAGCACTAATGCACCGGATCCCATCAGAACTCCGAAGTTAAGCGTGCTTGGGCGAGAGTAGTACTAGGATGGGTGACCTCCTGGGAAGTCCTCGTGTTGCACCCCTTTTGCCTGCCGCTCTCCCCGCAGCTTGCAATTGATGTCGCTGCCTCTGAGCTTCGTCGTGATCTCCTCACCTAGGGCGAGGGCCATCTTCCGGAGGCCCTTCCTCTGGTCAGTTCATCCCCCTCTTTCCCTTCCTCCTCCGAACGCCCCGTCCGCCACGCGCGAACTGGTTGCCGCACTACCATTTTGGGTATGCTGGGTCTTTTTTGGAGCTCAGTGGGTGCGGCTTCCCTTGCAAAAGGTTTGCCTCTTTTATGTTGGCTAGCTAGCTAGTTGGGGTTTGGGCCTATTGGGCCCTTTTAGAGGGGGCACCGCCCTCATAAAAATTGCTTGCAATATTCAAATAACTTGGGCATGAAACGCGACGGGAGGAAAGGAGAAAATAGGGCTATAAAGTGTGGAATATTAAAGGAGGTTAGAGGAAGATATTAAATCGGAGGATGCGGATAGAAGATAGAGAAAGTCACGCTAGAAATATTTCCATTTCGGAGTAGAAGCATGAGTGGGAATGCGTGAGTGTTGATATCTATGCATAGATATCTAAGAGAGAGAGAGAGAGAATTTAGAGTGGAGCTATCCACTTTGAGGCTCTAGCTATTTCTAACCCTTAATTAGTTCACATATGGGTGACTTTATCATCAACTGATGTGGCCCCATCACTCGCCGCAAACCATCTCTTGAGCCCCCTCCTCTTCAGGGCGAATTTCGTCGCCGCCCCGAAGCGTTCTCGTGGGCTCCCTACGTCTGGCCTTTGACCGATCCGTGGATGTCTTGCCACCCTCCCACGCGCAGAGTAGCTGCCTTTCGGGAGCGTTTTGGGCAGGTCTTGGGCACGTAGAGTGGACCTGAGAGTTGATGGGGACTTGAGGTGGGCTACCTGCTCGTCCTATGCACGATTCCCCCTGGCGAGGGACGCGCCGCCTTGGGGGTCGGTCATGTCACCGGCCCTCTCCCTCTAGCGGGGCGGAAATGATCCGGTGCCACGTGGTGTTGGAGAGGGGGCAATCTGGCGAGTGGGGACGAGGGTATGAGGTGGTGTGTGGCGCTAGTGATGTTGGGTGCGATCATACCAGCACTAATGCACCGGATCCCATCAGAACTCCGAAGTTAAGCGTGCTTGGGCGAGAGTAGTACTAGGATGGGTGACCTCCTGGGAAGTCCTCGTGTTGCACCCCTTTTGCCTGCCGCTCTCCCCGCAGCTTGCAATTGATGTCGCTGCCTCTGAGCTTCGTCGTGATCTCCTCACCTAGGGCGAGGGCCATCTTCCGGAGGCCCTTCCTCTGGTCAGTTCATCCCCCTCTTTCCCTTCCTCCTCCGAACGCCCCGTCCGCCACACGCGAACTGGTTGCCGCACTACCATTTTGGGTATGCTGGGTCTTTTTTGGAGCTCAGTGGGTGCGGCTTCCCTTGCAAAAGGTTTGCCTCTTTTATGTTGGCTAGCTAGCTAGTTGGGGTTTGGGCCTATTGGGCCCTTTTAGAGGGGGCACCGCCCTCATAAAAATTGCTTGCAATATTCAAATAACTTGGGCATGAAACGCGACGGGAGGAAAAGAGAAAATAGGGCTATAAAGTGTGGAATATTAAAGGAGGTTAGAGGAAGATATTAAATCGGAGGATGCGGATAGAAGATAGAGAAAGTCACGCTAGAAATATTTCCATTTCGGAGTAGAAGCATGAATGGGAATGCGTGAGTGTTGATATCTATGCATAGATATCTAAGAGAGAGAGAGAGAGAATTTAGAGTGGAGCTATCCACTTTGAGGCTCTAGCTATTTCTAACCCTTAATTAGTTCACATATGGGTGACTTTATCATCAACTGATGTGGCCCCATCACTCGCCGCAAACCATCTCTTGAGCCCCCTCCTCTTCAGGGCGAATTTCGTCGCCGCCCCGAAGCGTTCTCGTGGGCTCCCTACGTCTGGCCTTTGACCGATCCGTGGATGTCTTGCCACCCTCCCACGCGCAGAGTAGCTGCCTTTCGGGAGCGTTTTGGGCAGGTCTTGGGCACGTAGAGTGGACCTGAGAGTTGATGGGGACTTGAGGTGGGCTACCTGCTCGTCCTATGCACGATTCCCCCTGGCGAGGGACGCGCCGCCTTGGGGGTCGGTCATGTCACCGGCCCTCTCCCTCTAGCGGGGCGGAAATGATCCGGTGCCACGTGGTGTTGGAGAGGGGGCAATCTGGCGAGTGGGGACGAGGGTATGAGGTGGTGTGTGGCGCTAGTGATGTTGGGTGCGATCATACCAGCACTAATGCACCGGATCCCATCAGAACTCCGAAGTTAAGCGTGCTTGGGCGAGAGTAGTACTAGGATGGGTGACCTCCTGGGAAGTCCTCGTGTTGCACCCCTTTTGCCTGCCGCTCTCCCCGCAGCTTGCAATTGATGTCGCTGCCTCTGAGCTTCGTCGTGATCTCCTCACCTAGGGCGAGGGCCATCTTCCGGAGGCCCTTCCTCTGGTCAGTTCATCCCCCTCTTTCCCTTCCTCCTCCGAACGCCCCGTCCGCCACACGCGAACTGGTTGCCGCACTACCATTTTGGGTATGCTGGGTCTTTTTTGGAGCTCAGTGGGTGCGGCTTCCCTTGCAAAAGGTTTGCCTCTTTTATGTTGGCTAGCTAGCTAGTTGGGGTTTGGGCCTATTGGGCCCTTTTAGAGGGGGCACCGCCCTCATAAAAATTGCTTGCAATATTCAAATAACTTGGGCATGAAACGCGACGGGAGGAAAAGAGAAAATAGGGCTATAAAGTGTGGAATATTAAAGGAGGTTAGAGGAAGATATTAAATCGGAGGATGCGGATAGAAGATAGAGAAAGTCACGCTAGAAATATTTCCATTTCGGAGTAGAAGCATGAATGGGAATGCGTGAGTGTTGATATCTATGCATAGATATCTAAGAGAGAGAGAGAGAGAATTTAGAGTGGAGCTATCCACTTTGAGGCTCTAGCTATTTCTAACCCTTAATTAGTTCACATATGGGTGACTTTATCATCAACTGATGTGGCCCCATCACTCGCCGCAAACCATCTCTTGAGCCCCCTCCTCTTCAGGGCGAATTTCGTCGCCGCCCCGAAGCGTTCTCGTGGGCTCCCTACGTCTGGCCTTTGACCGATCCGTGGATGTCTTGCCACCCTCCCACGCGCAGAGTAGCTGCCTTTCGGGAGCGTTTTGGGCAGGTCTTGGGCACGTAGAGTGGACCTGAGAGTTGATGGGGACTTGAGGTGGGCTACCTGCTCGTCCTATGCACGATTCCCCCTGGCGAGGGACGCGCCGCCTTGGGGGTCGGTCATGTCACCGGCCCTCTCCCTCTAGCGGGGCGGAAATGATCCGGTGCCACGTGGTGTTGGAGAGGGGGCAATCTGGCGAGTGGGGACGAGGGTATGAGGTGGTGTGTGGCGCTAGTGATGTTGGGTGCGATCATACCAGCACTAATGCACCGGATCCCATCAGAACTCCGAAGTTAAGCGTGCTTGGGCGAGAGTAGTACTAGGATGGGTGACCTCCTGGGAAGTCCTCGTGTTGCACCCCTTTTGCCTGCCGCTCTCCCCGCAGCTTGCAATTGATGTCGCTGCCTCTGAGCTTCGTCGTGATCTCCTCACCTAGGGCGAGGGCCATCTTCCGGAGGCCCTTCCTCTGGTCAGTTCATCCCCCTCTTTCCCTTCCTCCTCCGAACGCCCCGTCCGCCACACGCGAACTGGTTGCCGCACTACCATTTTGGGTATGCTGGGTCTTTTTTGGAGCTCAGTGGGTGCGGCTTCCCTTGCAAAAGGTTTGCCTCTTTTATGTTGGCTAGCTAGCTAGTTGGGGTTTGGGCCTATTGGGCCCTTTTAGAGGGGGCACCGCCCTCATAAAAATTGCTTGCAATATTCAAATAACTTGGGCATGAAACGCGACGGGAGGAAAAGAGAAAATAGGGCTATAAAGTGTGGAATATTAAAGGAGGTTAGAGGAAGATATTAAATCGGAGGATGCGGATAGAAGATAGAGAAAGTCACGCTAGAAATATTTCCATTTCGGAGTAGAAGCATGAATGGGAATGCGTGAGTGTTGATATCTATGCATAGATATCTAAGAGAGAGAGAGAGAGAATTTAGAGTGGAGCTATCCACTTTGAGGCTCTAGCTATTTCTAACCCTTAATTAGTTCACATATGGGTGACTTTATCATCAACTGATGTGGCCCCATCACTCGCCGCAAACCATCTCTTGAGCCCCCTCCTCTTCAGGGCGAATTTCGTCGCCGCCCCGAAGCGTTCTCGTGGGCTCCCTACGTCTGGCCTTTGACCGATCCGTGGATGTCTTGCCACCCTCCCACGCGCAGAGTAGCTGCCTTTCGGGAGCGTTTTGGGCAGGTCTTGGGCACGTAGAGTGGACCTGAGAGTTGATGGGGACTTGAGGTGGGCTACCTGCTCGTCCTATGCACGATTCCCCCTGGCGAGGGACGCGCCGCCTTGGGGGTCGGTCATGTCACCGGCCCTCTCCCTCTAGCGGGGCGGAAATGATCCGGTGCCACGTGGTGTTGGAGAGGGGGCAATCTGGCGAGTGGGGACGAGGGTATGAGGTGGTGTGTGGCGCTAGTGATGTTGGGTGCGATCATACCAGCACTAATGCACCGGATCCCATCAGAACTCCGAAGTTAAGCGTGCTTGGGCGAGAGTAGTACTAGGATGGGTGACCTCCTGGGAAGTCCTCGTGTTGCACCCCTTTTGCCTGCCGCTCTCCCCGCAGCTTGCAATTGATGTCGCTGCCTCTGAGCTTCGTCGTGATCTCCTCACCTAGGGCGAGGGCCATCTTCCGGAGGCCCTTCCTCTGGTCAGTTCATCCCCCTCTTTCCCTTCCTCCTCCGAACGCCCCGTCCGCCACACGCGAACTGGTTGCCGCACTACCATTTTGGGTATGCTGGGTCTTTTTTGGAGCTCAGTGGGTGCGGCTTCCCTTGCAAAAAGTTTGCCTCTTTTATGTTGGCTAGCTAGCTAGTTGGGGTTTGGGCCTATTGGGCCCTTTTAGAGGGGGCACCGCCCTCATAAAAATTGCTTGCAATATTCAAATAACTTGGGCATGAAACGCGACGGGAGGAAAAGAGAAAATAGGGCTATAAAGTGTGGAATATTAAAGGAGGTTAGAGGAAGATATTAAATCGGAGGATGCGGATAGAAGATAGAGAAAGTCACGCTAGAAATATTTCCATTTCGGAGTAGAAGCATGAATGGGAATGCGTGAGTGTTGATATCTATGCATAGATATCTAAGAGAGAGAGAGAGAGAATTTAGAGTGGAGCTATCCACTTTGAGGCTCTAGCTATTTCTAACCCTTAATTAGTTCACATATGGGTGACTTTATCATCAACTGATGTGGCCCCATCACTCGCCGCAAACCATCTCTTGAGCCCCCTCCTCTTCAGGGCGAATTTCGTCGCCGCCCCGAAGCGTTCTCGTGGGCTCCCTACGTCTGGCCTTTGACCGATCCGTGGATGTCTTGCCACCCTCCCACGCGCAGAGTAGCTGCCTTTCGGGAGCGTTTTGGGCAGGTCTTGGGCACGTAGAGTGGACCTGAGAGTTGATGGGGACTTGAGGTGGGCTACCTGCTCGTCCTATGCACGATTCCCCCTGGCGAGGGACGCGCCGCCTTGGGGGTCGGTCATGTCACCGGCCCTCTCCCTCTAGCGGGGCGGAAATGATCCGGTGCCACGTGGTGTTGGAGAGGGGGCAATCTGGCGAGTGGGGACGAGGGTATGAGGTGGTGTGTGGCGCTAGTGATGTTGGGTGCGATCATACCAGCACTAATGCACCGGATCCCATCAGAACTCCGAAGTTAAGCGTGCTTGGGCGAGAGTAGTACTAGGATGGGTGACCTCCTGGGAAGTCCTCGTGTTGCACCCCTTTTGCCTGCCGCTCTCCCCGCAGCTTGCAATTGATGTCGCTGCCTCTGAGCTTCGTCGTGATCTCCTCACCTAGGGCGAGGGCCATCTTCCGGAGGCCCTTCCTCTGGTCAGTTCATCCCCCTCTTTCCCTTCCTCCTCCGAACGCCCCGTCCGCCACACGCGAACTGGTTGCCGCACTACCATTTTGGGTATGCTGGGTCTTTTTTGGAGCTCAGTGGGTGCGGCTTCCCTTGCAAAAGGTTTGCCTCTTTTATGTTGGCTAGCTAGCTAGTTGGGGTTTGGGCCTATTGGGCCCTTTTAGAGGGGGCACCGCCCTCATAAAAATTGCTTGCAATATTCAAATAACTTGGGCATGAAACGCGACGGGAGGAAAAGAGAAAATAGGGCTATAAAGTGTGGAATATTAAAGGAGGTTAGAGGAAGATATTAAATCGGAGGATGCGGATAGAAGATAGAGAAAGTCACGCTAGAAATATTTCCATTTCGGAGTAGAAGCATGAATGGGAATGCGTGAGTGTTGATATCTATGCATAGATATCTAAGAGAGAGAGAGAGAGAATTTAGAGTGGAGCTATCCACTTTGAGGCTCTAGCTATTTCTAACCCTTAATTAGTTCACATATGGGTGACTTTATCATCAACTGATGTGGCCCCATCACTCGCCGCAAACCATCTCTTGAGCCCCCTCCTCTTCAGGGCGAATTTCGTCGCCGCCCCGAAGCGTTCTCGTGGGCTCCCTACGTCTGGCCTTTGACCGATCCGTGGATGTCTTGCCACCCTCCCACGCGCAGAGTAGCTGCCTTTCGGGAGCGTTTTGGGCAGGTCTTGGGCACGTAGAGTGGACCTGAGAGTTGATGGGGACTTGAGGTGGGCTACCTGCTCGTCCTATGCACGATTCCCCCTGGCGAGGGACGCGCCGCCTTGGGGGTCGGTCATGTCACCGGCCCTCTCCCTCTAGCGGGGCGGAAATGATCCGGTGCCACGTGGTGTTGGAGAGGGGGCAATCTGGCGAGTGGGGACGAGGGTATGAGGTGGTGTGTGGCGCTAGTGATGTTGGGTGCGATCATACCAGCACTAATGCACCGGATCCCATCAGAACTCCGAAGTTAAGCGTGCTTGGGCGAGAGTAGTACTAGGATGGGTGACCTCCTGGGAAGTCCTCGTGTTGCACCCCTTTTGCCTGCCGCTCTCCCCGCAGCTTGCAATTGATGTCGCTGCCTCTGAGCTTCGTCGTGATCTCCTCACCTAGGGCGAGGGCCATCTTCCGGAGGCCCTTCCTCTGGTCAGTTCATCCCCCTCTTTCCCTTCCTCCTCCGAACGCCCCGTCCGCCACACGCGAACTGGTTGCCGCACTACCATTTTGGGTATGCTGGGTCTTTTTTGGAGCTCAGTGGGTGCGGCTTCCCTTGCAAAAGGTTTGCCTCTTTTATGTTGGCTAGCTAGCTAGTTGGGGTTTGGGCCTATTGGGCCCTTTTAGAGGGGGCACCGCCCTCATAAAAATTGCTTGCAATATTCAAATAACTTGGGCATGAAACGCGACGGGAGGAAAAGAGAAAATAGGGCTATAAAGTGTGGAATATTAAAGGAGGTTAGAGGAAGATATTAAATCGGAGGATGCGGATAGAAGATAGAGAAAGTCACGCTAGAAATATTTCCATTTCGGAGTAGAAGCATGAATGGGAATGCGTGAGTGTTGATATCTATGCATAGATATCTAAGAGAGAGAGAGAGAGAATTTAGAGTGGAGCTATCCACTTTGAGGCTCTAGCTATTTCTAACCCTTAATTAGTTCACATATGGGTGACTTTATCATCAACTGATGTGGCCCCATCACTCGCCGCAAACCATCTCTTGAGCCCCCTCCTCTTCAGGGCGAATTTCGTCGCCGCCCCGAAGCGTTCTCGTGGGCTCCCTACGTCTGGCCTTTGACCGATCCGTGGATGTCTTGCCACCCTCCCACGCGCAGAGTAGCTGCCTTTCGGGAGCGTTTTGGGCAGGTCTTGGGCACGTAGAGTGGACCTGAGAGTTGATGGGGACTTGAGGTGGGCTACCTGCTCGTCCTATGCACGATTCCCCCTGGCGAGGGACGCGCCGCCTTGGGGGTCGGTCATGTCACCGGCCCTCTCCCTCTAGCGGGGCGGAAATGATCCGGTGCCACGTGGTGTTGGAGAGGGGGCAATCTGGCGAGTGGGGACAAGGGTATGAGGTGGTGTGTGGCGCTAGTGATGTTGGGTGCGATCATACCAGCACTAATGCACCGGATCCCATCAGAACTCCGAAGTTAAGCGTGCTTGGGCGAGAGTAGTACTAGGATGGGTGACCTCCTGGGAAGTCCTCGTGTTGCACCCCTTTTGCCTGCCGCTCTCCCCGCAGCTTGCAATTGATGTCGCTGCCTCTGAGCTTCGTCGTGATCTCCTCACCTAGGGCGAGGGCCATCTTCCGGAGGCCCTTCCTCTGGTCAGTTCATCCCCCTCTTTCCCTTCCTCCTCCGAACGCCCCGTCCGCCACACGCGAACTGGTTGCCGCACTACCATTTTGGGTATGCTGGGTCTTTTTTGGAGCTCAGTGGGTGCGGCTTCCCTTGCAAAAGGTTTGCCTCTTTTATGTTGGCTAGCTAGCTAGTTGGGGTTTGGGCCTATTGGGCCCTTTTAGAGGGGGCACCGCCCTCATAAAAATTGCTTGCAATATTCAAATAACTTGGGCATGAAACGCGACGGGAGGAAAAGAGAAAATAGGGCTATAAAGTGTGGAATATTAAAGGAGGTTAGAGGAAGATATTAAATCGGAGGATGCGGATAGAAGATAGAGAAAGTCACGCTAGAAATATTTCCATTTCGGAGTAGAAGCATGAATGGGAATGCGTGAGTGTTGATATCTATGCATAGATATCTAAGAGAGAGAGAGAGAGAATTTAGAGTGGAGCTATCCACTTTGAGGCTCTAGCTATTTCTAACCCTTAATTAGTTCACATATGGGTGACTTTATCATCAACTGATGTGGCCCCATCACTCGCCGCAAACCATCTCTTGAGCCCCCTCCTCTTCAGGGCGAATTTCGTCGCCGCCCCGAAGCGTTCTCGTGGGCTCCCTACGTCTGGCCTTTGACCGATCCGTGGATGTCTTGCCACCCTCCCACGCGCAGAGTAGCTGCCTTTCGGGAGCGTTTTGGGCAGGTCTTGGGCACGTAGAGTGGACCTGAGAGTTGATGGGGACTTGAGGTGGGCTACCTGCTCGTCCTATGCACGATTCCCCCTGGCGAGGGACGCGCCGCCTTGGGGGTCGGTCATGTCACCGGCCCTCTCCCTCTAGCGGGGCGGAAATGATCCGGTGCCACGTGGTGTTGGAGAGGGGGCAATCTGGCGAGTGGGGACGAGGGTATGAGGTGGTGTGTGGCGCTAGTGATGTTGGGTGCGATCATACCAGCACTAATGCACCGGATCCCATCAGAACTCCGAAGTTAAGCGTGCTTGGGCGAGAGTAGTACTAGGATGGGTGACCTCCTGGGAAGTCCTCGTGTTGCACCCCTTTTGCCTGCCGCTCTCCCCGCAGCTTGCAATTGATGTCGCTGCCTCTGAGCTTCGTCGTGATCTCCTCACCTAGGGCGAGGGCCATCTTCCGGAGGCCCTTCCTCTGGTCAGTTCATCCCCCTCTTTCCCTTCCTCCTCCGAACGCCCCGTCCGCCACACGCGAACTGGTTGCCGCACTACCATTTTGGGTATGCTGGGTCTTTTTTGGAGCTCAGTGGGTGCGGCTTCCCTTGCAAAAGGTTTGCCTCTTTTATGTTGGCTAGCTAGCTAGTTGGGGTTTGGGCCTATTGGGCCCTTTTAGAGGGGGCACCGCCCTCATAAAAATTGCTTGCAATATTCAAATAACTTGGGCATGAAACGCGACGGGAGGAAAAGAGAAAATAGGGCTATAAAGTGTGGAATATTAAAGGAGGTTAGAGGAAGATATTAAATCGGAGGATGCGGATAGAAGATAGAGAAAGTCACGCTAGAAATATTTCCATTTCGGAGTAGAAGCATGAATGGGAATGCGTGAGTGTTGATATCTATGCATAGATATCTAAGAGAGAGAGAGAGAGAATTTAGAGTGGAGCTATCCACTTTGAGGCTCTAGCTATTTCTAACCCTTAATTAGTTCACATATGGGTGACTTTATCATCAACTGATGTGGCCCCATCACTCGCCGCAAACCATCTCTTGAGCCCCCTCCTCTTCAGGGCGAATTTCGCCGCCGCCCCGAAGCGTTCTCGTGGGCTCCCTACGTCTGGCCTTTGACCGATCCGTGGATGTCTTGCCACCCTCCCACGCGCAGAGTAGCTGCCTTTCGGGAGCGTTTTGGGCAGGTCTTGGGCACGTAGAGTGGACCTGAGAGTTGATGGGGACTTGAGGTGGGCTACCTGCTCGTCCTATGCACGATTCCCCCTGGCGAGGGACGCGCCGCCTTGGGGGTCGGTCATGTCACCGGCCCTCTCCCTCTAGCGGGGCGGAAATGATCCGGTGCCACGTGGTGTTGGAGAGGGGGCAATCTGGCGAGTGGGGACGAGGGTATGAGGTGGTGTGTGGCGCTAGTGATGTTGGGTGCGATCATACCAGCACTAATGCACCGGATCCCATCAGAACTCCGAAGTTAAGCGTGCTTGGGCGAGAGTAGTACTAGGATGGGTGACCTCCTGGGAAGTCCTCGTGTTGCACCCCTTTTGCCTGCCGCTCTCCCCGCAGCTTGCAATTGATGTCGCTGCCTCTGAGCTTCGTCGTGATCTCCTCACCTAGGGCGAGGGCCATCTTCCGGAGGCCCTTCCTCTGGTCAGTTCATCCCCCTCTTTCCCTTCCTCCTCCGAACGCCCCGTCCGCCACACGCGAACTGGTTGCCGCACTACCATTTTGGGTATGCTGGGTCTTTTTTGGAGCTCAGTGGGTGCGGCTTCCCTTGCAAAAGGTTTGCCTCTTTTATGTTGGCTAGCTAGCTAGTTGGGGTTTGGGCCTATTGGGCCCTTTTAGAGGGGGCACCGCCCTCATAAAAATTGCTTGCAATATTCAAATAACTTGGGCATGAAACGCGACGGGAGGAAAAGAGAAAATAGGGCTATAAAGTGTGGAATATTAAAGGAGGTTAGAGGAAGATATTAAATCGGAGGATGCGGATAGAAGATAGAGAAAGTCACGCTAGAAATATTTCCATTTCGGAGTAGAAGCATGAATGGGAATGCGTGAGTGTTGATATCTATGCATAGATATCTAAGAGAGAGAGAGAGAGAATTTAGAGTGGAGCTATCCACTTTGAGGCTCTAGCTATTTCTAACCCTTAATTAGTTCACATATGGGTGACTTTATCATCAACTGATGTGGCCCCATCACTCGCCGCAAACCATCTCTTGAGCCCCCTCCTCTTCAGGGCGAATTTCGTCGCCGCCCCGAAGCGTTCTCGTGGGCTCCCTACGTCTGGCCTTTGACCGATCCGTGGATGTCTTGCCACCCTCCCACGCGCAGAGTAGCTGCCTTTCGGGAGCGTTTTGGGCAGGTCTTGGGCACGTAGAGTGGACCTGAGAGTTGATGGGGACTTGAGGTGGGCTACCTGCTCGTCCTATGCACGATTCCCCCTGGCGAGGGACGCGCCGCCTTGGGGGTCGGTCATGTCACCGGCCCTCTCCCTCTAGCGGGGCGGAAATGATCCGGTGCCACGTGGTATTGGAGAGGGGGCAATCTGGCGAGTGGGGACGAGGGTATGAGGTGGTGTGTGGCGCTAGTGATGtTGGGTGCGATCATACCAGCACTAATGCACCGGATCCCATCAGAACTCCGAAGTTAAGCGTGCTTGGGCGAGAGTAGTACTAGGATGGGTGACCTCCTGGGAAGTCCTCGTGTTGCACCCCTTTTGCCTGCCGCTCTCCCCGCAGCTTGCAATTGATGTCGCTGCCTCTGAGCTTCGTCGTGATCTCCTCACCTAGGGCGAGGGCCATCTTCCGGAGGCCCTTCCTCTGGTCAGTTCATCCCCCTCTTTCCCTTCCTCCTCCGAACGCCCCGTCCGCCACACGCGAACTGGTTGCCGCACTACCATTTTGGGTATGCTGGGTCTTTTTTGGAGCTCATTGGGTGCGGCTTCCCTTGCAAAAGGTTTGCCTCTTTTATGTTGGCTAGCTAGCTAGTTGGGGTTTGGGCCTATTGGGCCCTTTTAGAGGGGGCACCGCCCTCATAAAAATTGCTTGCAATATTCAAATAACTTGGGCATGAAACGCGACGGGAGGAAAAGAGAAAATAGGGCTATAAAGTGTGGAATATTAAAGGAGGTTAGAGGAAGATATTAAATCGGAGGATGCGGATAGAAGATAGAGAAAGTCACGCTAGAAATATTTCCATTTCGGAGTAGAAGCATGAATGGGAATGCGTGAGTGTTGATATCTATGCATAGATATCTAAGAGAGAGAGAGAGAGAATTTAGAGTGGAGCTATCCACTTTGAGGCTCTAGCTATTTCTAACCCTTAATTAGTTCACATATGGGTGACTTTATCATCAACTGATGTGGCCCCATCACTCGCCGCAAACCATCTCTTGAGCCCCCTCCTCTTCAGGGCGAACTTCGTCGCCGCCCCGAAGCGTTCTCGTGGGCTCCCTACGTCTGGCCTTTGACCGATCCGTGGATGTCTTGCCACCCTCCCACGCGCAGAGTAGCTGCCTTTCGGGAGCGTTTTGGGCAGGTCTTGGGCACGTAGAGTGGACCTGAGAGTTGATGGGGACTTGAGGTGGGCTACCTGCTCGTCCTATGCACGATTCCCCCGGGCGAGGGACGCGCCGCCTTGGGGGTCGGTCATGTCACCGGCCCTCTCCCTCGAGCGGGGCGGAAATGATCCGGTGCCACGTGGTGTTGGAGAGGGGGCAATCTGGCGAGTGGGGACGAGGGTATGAGGTGGTGTGTGGCGCTAGTGATGTTGGGTGCGATCATACCAGCACTAATGCACCGGATCCCATCAGAACTCCGAAGTTAAGCGTGCTTGGGCGAGAGTAGTACTAGGATGGGTGACCTCCTGGGAAGTCCTCGTGTTGCACCCCTTTTGCCTGCCGCTCTCCCCGCAGCCTGCAATTGATGTCGCTGCCTCTGAGCTTCGTCGTGATCTCCTCACCTAGGGCGAGGGCCATCTTCCGGAGGCCCTTCCTCTGGTCAGTTCATCCCCCTCTTTCCCTTCCTCCTCCGAACGCCCCGTCCGCCACACGCGAACTGGTTGCCGCACTACCATTTTGGGTATGCTGGGTCTTTTTTGGAGCTCAGTGGGTGCGGCTTCCCTTGCAAAAGGTTTGCCTCTTTTATGTTGGCTAGCTAGCTAGTTGGGGTTTGGGCCTATTGGGCCCTTTTAGAGGGGGCACCGCCCTCATAAAAATTGCTTGCAATATTCAAATAACTTGGGCATGAAACGCGACGGGAGGAAAAGAGAAAATAGGGCTATAAAGTGTGGAATATTAAAGGAGGTTAGAGGAAGATATTAAATCGGAGGATGCGGATAGAAGATAGAGAAAGTCACGCTAGAAATATTTCCATTTCGGAGTAGAAGCATGAATGGGAATGCGTGAGTGTTGATATCTATGCATAGATATCTAAGAGAGAGAGAGAGAGAATTTAGAGTGGAGCTATCCACTTTGAGGCTCTAGCTATTTCTAACCCTTAATTAGTTCACATATGGGTGACTTTATCATCAACTGATGTGGCCCCATCACTCGCCGCAAACCATCTCTTGAGCCCCCTCCTCTTCAGGGCGAATTTCGTCGCCGCCCCGAAGCGTTCTCGTGGGCTCCCTACGTCTGGCCTTTGACCGATCCGTGGATGTCTTGCCACCCTCCCACGCGCAGAGTAGCTGCCTTTCGGGAGCGTTTTGGGCAGGTCTTGGGCACGTAGAGTGGACCTGAGAGTTGATGGGGACTTGAGGTGGGCTACCTGCTCGTCCTATGCACGATTCCCCCTGGCGAGGGACGCGCCGCCTTGGGGGTCGGTCATGTCACCGGCCCTCTCCCTCTAGCGGGGCGGAAATGATCCGGTGCCACGTGGTGTTGGAGAGGGGGCAATCTGGCGAGTGGGGACGAGGGTATGAGGTGGTGTGTGGCGCTAGTGATGTTGGGTGCGATCATACCAGCACTAATGCACCGGATCCCATCAGAACTCCGAAGTTAAGCGTGCTTGGGCGAGAGTAGTACTAGGATGGGTGACCTCCTGGGAAGTCCTCGTGTTGCACCCCTTTTGCCTGCCGCTCTCCCCGCAGCTTGCAATTGATGTCGCTGCCTCTGAGCTTCGTCGTGATCTCCTCACCTAGGGCGAGGGCCATCTTCCGGAGGCCCTTCCTCTGGTCAGTTCATCCCCCTCTTTCCCTTCCTCCTCCGAACGCCCCGTCCGCCACACGCGAACTGGTTGCCGCACTACCATTTTGGGTATGCTGGGTCTTTTTTGGAGCTCAGTGGGTGCGGCTTCCCTTGCAAAAGGTTTGCCTCTTTTATGTTGGCTAGCTAGCTAGTTGGGGTTTGGGCCTATTGGGCCCTTTTAGAGGGGGCACCGCCCTCATAAAAATTGCTTGCAATATTCAAATAACTTGGGCATGAAACGCGACGGGAGGAAAAGAGAAAATAGGGCTATAAAGTGTGGAATATTAAAGGAGGTTAGAGGAAGATATTAAATCGGAGGATGCGGATAGAAGATAGAGAAAGTCACGCTAGAAATATTTCCATTTCGGAGTAGAAGCATGAATGGGAATGCGTGAGTGTTGATATCTATGCATAGATATCTAAGAGAGAGAGAGAGAGAATTTAGAGTGGAGCTATCCACTTTGAGGCTCTAGCTATTTCTAACCCTTAATTAGTTCACATATGGGTGACTTTATCATCAACTGATGTGGCCCCATCACTCGCCGCAAACCATCTCTTGAGCCCCCTCCTCTTCAGGGCGAATTTCGTCGCCGCCCCGAAGCGTTCTCGTGGGCTCCCTACGTCTGGCCTTTGACCGATCCGTGGATGTCTTGCCACCCTCCCACGCGCAGAGTAGCTGCCTTTCGGGAGCGTTTTGGGCAGGTCTTGGGCACGTAGAGTGGACCTGAGAGTTGATGGGGACTTGAGGTGGGCTACCTGCTCGTCCTATGCACGATTCCCCCTGGCGAGGGACGCGCCGCCTTGGGGGTCGGTCATGTCACCGGCCCTCTCCCTCTAGCGGGGCGGAAATGATCCGGTGCCACGTGGTGTTGGAGAGGGGGCAATCTGGCGAGTGGGGACGAGGGTATGAGGTGGTGTGTGGCGCTAGTGATGTTGGGTGCGATCATACCAGCACTAATGCACCGGATCCCATCAGAACTCCGAAGTTAAGCGTGCTTGGGCGAGAGTAGTACTAGGATGGGTGACCTCCTGGGAAGTCCTCGTGTTGCACCCCTTTTGCCTGCCGCTCTCCCCGCAGCTTGCAATTGATGTCGCTGCCTCTGAGCTTCGTCGTGATCTCCTCACCTAGGGCGAGGGCCATCTTCCGGAGGCCCTTCCTCTGGTCAGTTCATCCCCCTCTTTCCCTTCCTCCTCCGAACGCCTCGTCCGCCACACGCGAACTGGTTGCCGCACTACCATTTTGGGTATGCTGGGTCTTTTTTGGAGCTCAGTGGGTGCGGCTTCCCTTGCAAAAGGTTTGCCTCTTTTATGTTGGCTAGCTAGCTAGTTGGGGTTTGGGCCTATTGGGCCCTTTTAGAGGGGGCACCGCCCTCATAAAAATTGCTTGCAATATTCAAATAACTTGGGCATGAAACGCGACGGGAGGAAAAGAGAAAATAGGGCTATAAAGTGTGGAATATTAAAGGAGGTTAGAGGAAGATATTAAATCGGAGGATGCGGATAGAAGATAGAGTAAGTCACGCTAGAAATATTTCCATTTCGGAGTAGAAGCATGAATGGGAATGCGTGAGTGTTGATATCTATGCATAGATATCTAAGAGAGAGAGAGAGAGAATTTAGAGTGGAGCTATCCACTTTGAGGCTCTAGCTATTTCTAACCCTTAATTAGTTCACATATGGGTGACTTTATCATCAACTGATGTGGCCCCATCACTCGCCGCAAACCATCTCTTGAGCCCCCTCCTCTTCAGGGCGAATTTCGTCGCCGCCCCGAAGCGTTCTCGTGGGCTCCCTACGTCTGGCCTTTGACCGATCCGTGGATGTCTTGCCACCCTCCCACGCGCAGAGTAGCTGCCTTTCGGGAGCGTTTTGGGCGGGTCTTGGGCACGTAGAGTGGACCTGAGAGTTGATGGGGACTTGAGGTGGGCTACCTGCTCGTCCTATGCACGATTCCCCCTGGCGAGGGACGCGCCGCCTTGGGGGTCGGTCATGTCACCGGCCCTCTCCCTCTAGCGGGGCGGAAATGATCCGGTGCCACGTGGTGTTGGAGAGGGGGCAATCTGGCGAGTGGGGACGAGGGTATGAGGTGGTGTGTGGCGCTAGTGATGTTGGGTGCGATCATACCAGCACTAATGCACCGGATCCCATCAGAACTCCGAAGTTAAGCGTGCTTGGGCGAGAGTAGTACTAGGATGGGTGACCTCCTGGGAAGTCCTCGTGTTGCACCCCTTTTGCCTGCCGCTCTCCCCGCAGCTTGCAATTGATGTCGCTGCCTCTGAGCTTCGTCGTGATCTCCTCACCTAGGGCGAGGGCCATCTTCCGGAGGCCCTTCCTCTGGTCAGTTCATCCCCCTCTTTCCCTTCCTCCTCCGAACGCCTCGTCCGCCACACGCGAACTGGTTGCCGCACTACCATTTTGGGTATGCTGGGTCTTTTTTGGAGCTCAGTGGGTGCGGCTTCCCTTGCAAAAGGTTTGCCTCTTTTATGTTGGCTAGCTAGCTAGTTGGGGTTTGGGCCTATTGGGCCCTTTTAGAGGGGGCACCGCCCTCATAAAAATTGCTTGCAATATTCAAATAACTTGGGCATGAAACGCGACGGGAGGAAAAGAGAAAATAGGGCTATAAAGTGTGGAGTATTAAAGGAGGTTAGAGGAAGATATTAAATCGGAGGATGCGGATAGAAGATAGAGAAAGTCACGCTAGAAATATTTCCATTTCGGAGTAGAAGCATGAATGGGAATGCGTGAGTGTTGATATCTATGCATAGATATCTAAGAGAGAGAGAGGGAGAATTTAGAGTGGAGCTATCCACTTTGAGGCTCTAGCTATTTCTAACCCTTAATTAGTTCACATATGGGTGACTTTATCATCAACTGATGTGGCCCCATCACTCGCCGCAAACCATCTCTTGAGCCCCCTCCTCTTCAGGGCGAATTTCGTCGCCGCCCCGAAGCGTTCTCGTGGGCTCCCTACGTCTGGCCTTTGACCGATCCGTGGATGTCTTGCCACCCTCCTACGCGCAGAGTAGCTGCCTTTCGGGAGCGTTTTGGGCAGGTCTTGGGCACGTAGAGTGGACCTGAGAGTTGATGGGGACTTGAGGTGGGCTACCTGCTCGTCCTATGCACGATTCCCCCTGGCGAGGGACGCGCCGCCTTGGGGGTCGGTCATGTCACCGGCCCTCTCCCTCTAGCGGGGCGGAAATGATCCGGTGCCACGTGGTGTTGGAGAGGGGGCAATCTGGCGAGTGGGGACGAGGGTATGAGGTGGTGTGTGGCGCTAGTGATGTTGGGTGCGATCATACCAGCACTAATGCACCGGATCCCATCAGAACTCCGAAGTTAAGCGTGCTTGGGCGAGAGTAGTACTAGGATGGGTGACCTCCTGGGAAGTCCTCGTGTTGCACCCCTTTTGCCTGCCGCTCTCCCCGCAGCTTGCAATTGATGTCGCTGCCTCTGAGCTTCGTCGTGATCTCCTCACCTAGGGCGAGGGCCATCTTCCGGAGGCCCTTCCTCTGGTCAGTTCATCCCCCTCTTTCCCTTCCTCCTCCGAACGCCTCGTCCGCCACACGCGAACTGGTTGCCGCACTACCATTTTGGGTATGCTGGGTCTTTTTTGGAGCTCAGTGGGTGCGGCTTCCCTTGCAAAAGGTTTGCCTCTTTTATGTTGGCTAGCTAGCTAGTTGGGGTTTGGGCCTATTGGGCCCTTTTAGAGGGGGCACCGCCCTCATAAAAATTGCTTGCAATATTCAAATAACTTGGGCATGAAACGCGACGGGAGGAAAAGAGAAAATAGGGCTATAAAGTGTGGAATATTAAAGGAGGTTAGAGGAAGATATTAAATCGGAGGATGCGGATAGAAGATAGAGAAAGTCACGCTAGAAATATTTCCATTTCGGAGTAGAAGCATGAATGGGAATGCGTGAGTGTTGATATCTATGCATAGATATCTAAGAGAGAGAGAGAGAGAATTTAGAGTGGAGCTATCCACTTTGAGGCTCTAGCTATTTCTAACCCTTAATTAGTTCACATATGGGTGACTTTATCATCAACTGATGTGGCCCCATCACTCGCCGCAAACCATCTCTTGAGCCCCCTCCTCTTCAGGGCGAATTTCGTCGCCGCCCCGAAGCGTTCTCGTGGGCTCCCTACGTCTGGCCTTTGACCGATCCGTGGATGTCTTGCCACCCTCCCACGCGCAGAGTAGCTGCCTTTCGGGAGCGTTTTGGGCAGGTCTTGGGCACGTAGAGTGGACCTGAGAGTTGATGGGGACTTGAGGTGGGCTACCTGCTCGTCCTATGCACGATTCCCCCTGGCGAGGGACGCGCCGCCTTGGGGGTCGGTCATGTCACCGGCCCTCTCCCTCTAGCGGGGCGGAAATGATCCGGTGCCACGTGGTGTTGGAGAGGGGGCAATCTGGCGAGTGGGGACGAGGGTATGAGGTGGTGTGTGGCGCTAGTGATGTTGGGTGCGATCATACCAGCACTAATGCACCGGATCCCATCAGAACTCCGAAGTTAAGCGTGCTTGGGCGAGAGTAGTACTAGGATGGGTGACCTCCTGGGAAGTCCTCGTGTTGCACCCCTTTTGCCTGCCGCAGCTTTGTGCCGCTCTCCCCGCAGCTTGCAATTGATGTCGCTGCCTCTGAGCTTCGTCGTGATCTCCTCACCTAGGGCGAGGGCCATCTTCCGGAGGCCCTTCCTCTGGTCAGTTCATCCCCCTCTTTCCCTTCCTCCTCCGAACGCCTCGTCCGCCACACGCGAACTGGTTGCCGCACTACCATTTTGGGTATGCTGGGTCTTTTTTGGAGCTCAGTGGGTGCGGCTTCCCTTGCAAAAGGTTTGCCTCTTTTATGTTGGCTAGCTAGCTAGTTGGGGTTTGGGCCTATTGGGCCCTTTTAGAGGGGGCACCGCCCTCATAAAAATTGCTTGCAATATTCAAATAACTTGGGCATGAAACGCGACGGGAGGAAAAGAGAAAATAGGGCTATAAAGTGTGGAATATTAAAGGAGGTTAGAGGAAGATATTAAATCGGAGGATGCGGATAGAAGATAGAGAAAGTCACGCTAGAAATATTTCCATTTCGGAGTAGAAGCATGAATGGGAATGCGTGAGTGTTGATATCTATGCATAGATATCTAAGAGAGAGAGAGAGAGAATTTAGAGTGGAGCTATCCACTTTGAGGCTCTAGCTATTTCTAACCCTTAATTAGTTCACATATGGGTGACTTTATCATCAACTGATGTGGCCCCATCACTCGCCGCAAACCATCTCTTGAGCCCCCTCCTCTTCAGGGCGAATTTCGTCGCCGCCCCGAAGCGTTCTCGTGGGCTCCCTACGTCTGGCCTTTGACCGATCCGTGGATGTCTTGCCACCCTCCCACGCGCAGAGTAGCTGCCTTTCGGGAGCGTTTTGGGCAGGTCTTGGGCACGTAGAGTGGACCTGAGAGTTGATGGGGACTTGAGGTGGGCTACCTGCTCGTCCTATGCACGATTCCCCCTGGCGAGGGACGCGCCGCCTTGGGGGTCGGTCATGTCACCGGCCCTCTCCCTCTAGCGGGGCGGAAATGATCCGGTGCCACGTGGTGTTGGAGAGGGGGCAATCTGGCGAGTGGGGACGAGGGTATGAGGTGGTGTGTGGCGCTAGTGATGTTGGGTGCGATCATACCAGCACTAATGCACCGGATCCCATCAGAACTCCGAAGTTAAGCGTGCTTGGGCGAGAGTAGTACTAGGATGGGTGACCTCCTGGGAAGTCCTCGTGTTGCACCCCTTTTGCCTGCCGCAGCTTTGTGCCGCTCTCCCCGCAGCTTGCAATTGATGTCGCTGCCTCTGAGCTTCGTCGTGATCTCCTCACCTAGGGCGAGGGCCATCTTCCGGAGGCCCTTCCTCTGGTCAGTTCATCCCCCTCTTTCCCTTCCTCCTCCGAACGCCTCGTCCGCCACACGCGAACTGGTTGCCGCACTACCATTTTGGGTATGCTGGGTCTTTTTTGGAGCTCAGTGGGTGCGGCTTCCCTTGCAAAAGGTTTGCCTCTTTTATGTTGGCTAGCTAGCTAGTTGGGGTTTGGGCCTATTGGGCCCTTTTAGAGGGGGCACCGCCCTCATAAAAATTGCTTGCAATATTCAAATAACTTGGGCATGAAACGCGACGGGAGGAAAAGAGAAAATAGGGCTATAAAGTGTGGAATATTAAAGGAGGTTAGAGGAAGATATTAAATCGGAGGATGCGGATAGAAGATAGAGAAAGTCACGCTAGAAATATTTCCATTTCGGAGTAGAAGCATGAATGGGAATGCGTGAGTGTTGATATCTATGCATAGATATCTAAGAGAGAGAGAGAGAGAATTTAGAGTGGAGCTATCCACTTTGAGGCTCTAGCTATTTCTAACCCTTAATTAGTTCACATATGGGTGACTTTATCATCAACTGATGTGGCCCCATCACTCGCCGCAAACCATCTCTTGAGCCCCCTCCTCTTCAGGGCGAATTTCGTCGCCGCCCCGAAGCGTTCTCGTGGGCTCCCTACGTCTGGCCTTTGACCGATCCGTGGATGTCTTGCCACCCTCCCACGCGCAGAGTAGCTGCCTTTCGGGAGCGTTTTGGGCAGGTCTTGGGCACGTAGAGTGGACCTGAGAGTTGATGGGGACTTGAGGTGGGCTACCTGCTCGTCCTATGCACGATTCCCCCTGGCGAGGGACGCGCCGCCTTGGGGGTCGGTCATGTCACCGGCCCTCTCCCTCTAGCGGGGCGGAAATGATCCGGTGCCACGTGGTGTTGGAGAGGGGGCAATCTGGCGAGTGGGGACGAGGGTATGAGGTGGTGTGTGGCGCTAGTGATGTTGGGTGCGATCATACCAGCACTAATGCACCGGATCCCATCAGAACTCCGAAGTTAAGCGTGCTTGGGCGAGAGTAGTACTAGGATGGGTGACCTCCTGGGAAGTCCTCGTGTTGCACCCCTTTTGCCTGCCGCAGCTTTGTGCCGCTCTCCCCGCAGCTTGCAATTGATGTCGCTGCCTCTGAGCTTCGTCGTGATCTCCTCACCTAGGGCGAGGGCCATCTTCCGGAGGCCCTTCCTCTGGTCAGTTCATCCCCCTCTTTCCCTTCCTCCTCCGAACGCCTCGTCCGCCACACGCGAACTGGTTGCCGCACTACCATTTTGGGTATGCTGGGTCTTTTTTGGAGCTCAGTGGGTGCGGCTTCCCTTGCAAAAGGTTTGCCTCTTTTATGTTGGCTAGCTAGCTAGTTGGGGTTTGGGCCTATTGGGCCCTTTTAGAGGGGGCACCGCCCTCATAAAAATTGCTTGCAATATTCAAATAACTTGGGCATGAAACGCGACGGGAGGAAAAGAGAAAATAGGGCTATAAAGTGTGGAATATTAAAGGAGGTTAGAGGAAGATATTAAATCGGAGGATGCGGATAGAAGATAGAGAAAGTCACGCTAGAAATATTTCCATTTCGGAGTAGAAGCATGAATGGGAATGCGTGAGTGTTGATATCTATGCATAGATATCTAAGAGAGAGAGAGAGAGAATTTAGAGTGGAGCTATCCACTTTGAGGCTCTAGCTATTTCTAACCCTTAATTAGTTCACATATGGGTGACTTTATCATCAACTGATGTGGCCCCATCACTCGCCGCAAACCATCTCTTGAGCCCCCTCCTCTTCAGGGCGAATTTCGTCGCCGCCCCGAAGCGTTCTCGTGGGCTCCCTACGTCTGGCCTTTGACCGATCCGTGGATGTCTTGCCACCCTCCCACGCGCAGAGTAGCTGCCTTTCGGGAGCGTTTTGGGCAGGTCTTGGGCACGTAGAGTGGACCTGAGAGTTGATGGGGACTTGAGGTGGGCTACCTGCTCGTCCTATGCACGATTCCCCCTGGCGAGGGACGCGCCGCCTTGGGGGTCGGTCATGTCACCGGCCCTCTCCCTCTAGCGGGGCGGAAATGATCCGGTGCCACGTGGTGTTGGAGAGGGGGCAATCTGGCGAGTGGGGACGAGGGTATGAGGTGGTGTGTGGCGCTAGTGATGTTGGGTGCGATCATACCAGCACTAATGCACCGGATCCCATCAGAACTCCGAAGTTAAGCGTGCTTGGGCGAGAGTAGTACTAGGATGGGTGACCTCCTGGGAAGTCCTCGTGTTGCACCCCTTTTGCCTGCCGCTCTCCCCGCAGCTTGCAATTGATGTCGCTGCCTCTGAGCTTCGTCGTGATCTCCTCACCTAGGGCGAGGGCCATCTTCCGGAGGCCCTTCCTCTGGTCAGTTCATCCCCCTCTTTCCCTTCCTCCTCCGAACGCCTCGTCCGCCACACGCGAACTGGTTGCCGCACTATCATTTTGGGTATGCTGGGTCTTTTTTGGAGCTCAGTGGGTGCGGCTTCCCTTGCAAAAGGTTTGCCTCTTTTATGTTGGCTAGCTAGCTAGTTGGGGTTTGGGCCTATTGGGCCCTTTTAGAGGGGGCACCGCCCTCATAAAAATTGCTTGCAATATTCAAATAACTTGGGCATGAAACGCGACGGGAGGAAAAGAGAAAATAGGGCTATAAAGTGTGGAATATTAAAGGAGGTTAGAGGAAGATATTAAATCGGAGGATGCGGATAGAAGATAGAGAAAGTCACGCTAGAAATATTTCCATTTCGGAGTAGAAGCATGAATGGGAATGCGTGAGTGTTGATATCTATGCATAGATATCTAAGAGAGAGAGAGAGAGAATTTAGAGTGGAGCTATCCACTTTGAGGCTCTAGCTATTTCTAACCCTTAATTAGTTCACATATGGGTGACTTTATCATCAACTGATGTGGCCCCATCACTCGCCGCAAACCATCTCTTGAGCCCCCTCCTCTTCAGGGCGAATTTCGTCGCCGCCCCGAAGCGTTCTCGTGGGCTCCCTACGTCTGGCCTTTGACCGATCCGTGGATGTCTTGCCACCCTCCCACGCGCAGAGTAGCTGCCTTTCGGGAGCGTTTTGGGCAGGTCTTGAGCACGTAGAGTGGACCTGAGAGTTGATGGGGACTTGAGGTGGGCTACCTGCTCGTCCTATCCACGATTCCCCCTGGCGAGGGACGCGCCGCCTTGGGGGTCGGTCATGTCACCGGCCCTCTCCCTCTAGCGGGGCGGAAATGATCCGGTGCCACGTGGTGTTGGAGAGGGGGCAATCTGGCGAGTGGGGACGAGGGTATGAGGTGGTGTGTGGCGCTAGTGATGTTGGGTGCGATCATACCAGCACTAATGCACCGGATCCCATCAGAACTCCGAAGTTAAGCGTGCTTGGGCGAGAGTAGTACTAGGATGGGTGACCTCCTGGGAAGTCCTCGTGTTGCACCCCTTTTGCCTGCCGCAGCTTTGTGCCGCTCTCCCCGCAGCTTGCAATTGATGTCGCTGCCTCTGAGCTTCGTCGTGATCTCCTCACCTAGGGCGAGGGCCATCTTCCGGAGGCCCTTCCTCTGGTCAGTTCATCCCCCTCTTTCCCTTCCTCCTCCGAACGCCTCGTCCGCCACACGCGAACTGGTTGCCGCACTACCATTTTGGGTATGCTGGGTCTTTTTTGGAGCTCAGTGGGTGCGGCTTCCCTTGCAAAAGGTTTGCCTCTTTTATGTTGGCTAGCTAGCTAGTTGGGGTTTGGGCCTATTGGGCCCTTTTAGAGGGGGCACCGCCCTCATAAAAATTGCTTGCAATATTCAAATAACTTGGGCATGAAACGCGACGGGAGGAAAAGAGAAAATAGGGCTATAAAGTGTGGAATATTAAAGGAGGTTAGAGGAAGATATTAAATCGGAGGATGCGGATAGAAGATAGAGAAAGTCACGCTAGAAATATTTCCATTTCGGAGTAGAAGCATGAATGGGAATGCGTGAGTGTTGATATCTATGCATAGATATCTAAGAGAGAGAGAGAGAGAATTTAGAGTGGAGCTATCCACTTTGAGGCTCTAGCTATTTCTAACCCTTAATTAGTTCACATATGGGTGACTTTATCATCAACTGATGTGGCCCCATCACTCGCCGCAAACCATCTCTTGAGCCCCCTCCTCTTCAGGGCGAATTTCGTCGCCGCCCCGAAGCGTTCTCGTGGGCTCCCTACGTCTGGCCTTTGACCGATCCGTGGATGTCTTGCCACCCTCCCACGCGCAGAGTAGCTGCCTTTCGGGAGCGTTTTGGGCAGGTCTTGGGCACGTAGAGTGGACCTGAGAGTTGATGGGGACTTGAGGTGGGCTACCTGCTCGTCCTATGCACGATTCCCCCTGGCGAGGGACGCGCCGCCTTGGGGGTCGGTCATGTCACCGGCCCTCTCCCTCTAGCGGGGCGGAAATGATCCGGTGCCACGTGGTGTTGGAGAGGGGGCAATCTGGCGAGTGGGGACGAGGGTATGAGGTGGTGTGTGGCGCTAGTGATGTTGGGTGCGATCATACCAGCACTAATGCACCGGATCCCATCAGAACTCCGAAGTTAAGCGTGCTTGGGCGAGAGTAGTACTAGGATGGGTGACCTCCTGGGAAGTCCTCGTGTTGCACCCCTTTTGCCTGCCGCAGCTTTGTGCCGCTCTCCCCGCAGCTTGCAATTGATGTCGCTGCCTCTGAGCTTCGTCGTGATCTCCTCACCTAGGGCGAGGGCCATCTTCCGGAGGCCCTTCCTCTGGTCAGTTCATCCCCCTCTTTCCCTTCCTCCTCCGAACGCCTCGTCCGCCACACGCGAACTGGTTGCCGCACTACCATTTTGGGTATGCTGGGTCTTTTTTGGAGCTCAGTGGGTGCGGCTTCCCTTGCAAAAGGTTTGCCTCTTTTATGTTGGCTAGCTAGCTAGTTGGGGTTTGGGCCTATTGGGCCCTTTTAGAGGGGGCACCGCCCTCATAAAAATTGCTTGCAATATTCAAATAACTTGGGCATGAAACGCGACGGGAGGAAAAGAGAAAATAGGGCTATAAAGTGTGGAATATTAAAGGAGGTTAGAGGAAGATATTAAATCGGAGGATGCGGATAGAAGATAGAGAAAGTCACGCTAGAAATATTTCCATTTCGGAGTAGAAGCATGAATGGGAATGCGTGAGTGTTGATATCTATGCATAGATATCTAAGAGAGAGAGAGAGAGAATTTAGAGTGGAGCTATCCACTTTGAGGCTCTAGCTATTTCTAACCCTTAATTAGTTCACATATGGGTGACTTTATCATCAACTGATGTGGCCCCATCACTCGCCGCAAACCATCTCTTGAGCCCCCTCCTCTTCAGGGCGAATTTCGTCGCCGCCCCGAAGCGTTCTCGTGGGCTCCCTACGTCTGGCCTTTGACCGATCCGTGGATGTCTTGCCACCCTCCCACGCGCAGAGTAGCTGCCTTTCGGGAGCGTTTTGGGCAGGTCTTGGGCACGTAGAGTGGACCTGAGAGTTGATGGGGACTTGAGGTGGGCTACCTGCTCGTCCTATGCACGATTCCCCCTGGCGAGGGACGCGCCGCCTTGGGGGTCGGTCATGTCACCGGCCCTCTCCCTCTAGCGGGGCGGAAATGATCCGGTGCCACGTGGTGTTGGAGAGGGGGCAATCTGGCGAGTGGGGACGAGGGTATGAGGTGGTGTGTGGCGCTAGTGATGTTGGGTGCGATCATACCAGCACTAATGCACCGGATCCCATCAGAACTCCGAAGTTAAGCGTGCTTGGGCGAGAGTAGTACTAGGATGGGTGACCTCCTGGGAAGTCCTCGTGTTGCACCCCTTTTGCCTGCCGCAGCTTTGTGCCGCTCTCCCCGCAGCTTGCAATTGATGTCGCTGCCTCTGAGCTTCGTCGTGATCTCCTCACCTAGGGCGAGGGCCATCTTCCGGAGGCCCTTCCTCTGGTCAGTTCATCCCCCTCTTTCCCTTCCTCCTCCGAACGCCTCGTCCGCCACACGCGAACTGGTTGCCGCACTACCATTTTGGGTATGCTGGGTCTTTTTTGGAGCTCAGTGGGTGCGGCTTCCCTTGCAAAAGGTTTGCCTCTTTTATGTTGGCTAGCTAGCTAGTTGGGGTTTGGGCCTATTGGGCCCTTTTAGAGGGGGCACCGCCCTCATAAAAATTGCTTGCAATATTCAAATAACTTGGGCATGAAACGCGACGGGAGGAAAAGAGAAAATAGGGCTATAAAGTGTGGAATATTAAAGGAGGTTAGAGGAAGATATTAAATCGGAGGATGCGGATAGAAGATAGAGAAAGTCACGCTAGAAATATTTCCATTTCGGAGTAGAAGCATGAATGGGAATGCGTGAGTGTTGATATCTATGCATAGATATCTAAGAGAGAGAGAGAGAGAATTTAGAGTGGAGCTATCCACTTTGAGGCTCTAGCTATTTCTAACCCTTAATTAGTTCACATATGGGTGACTTTATCATCAACTGATGTGGCCCCATCACTCGCCGCAAACCATCTCTTGAGCCCCCTCCTCTTCAGGGCGAATTTCGTCGCCGCCCCGAAGCGTTCTCGTGGGCTCCCTACGTCTGGCCTTTGACCGATCCGTGGATGTCTTGCCACCCTCCCACGCGCAGAGTAGCTGCCTTTCGGGAGCGTTTTGGGCAGGTCTTGGGCACGTAGAGTGGACCTGAGAGTTGATGGGGACTTGAGGTGGGCTACCTGCTCGTCCTATGCACGATTCCCCCTGGCGAGGGACGCGCCGCCTTGGGGGTCGGTCATGTCACCGGCCCTCTCCCTCTAGCGGGGCGGAAATGATCCGGTGCCACGTGGTGTTGGAGAGGGGGCAATCTGGCGAGTGGGGACGAGGGTATGAGGTGGTGTGTGGCGCTAGTGATGTTGGGTGCGATCATACCAGCACTAATGCACCGGATCCCATCAGAACTCCGAAGTTAAGCGTGCTTGGGCGAGAGTAGTACTAGGATGGGTGACCTCCTGGGAAGTCCTCGTGTTGCACCCCTTTTGCCTGCCGCAGCTTTGTGCCGCTCTCCCCGCAGCTTGCAATTGATGTCGCTGCCTCTGAGCTTCGTCGTGATCTCCTCACCTAGGGCGAGGGCCATCTTCCGGAGGCCCTTCCTCTGGTCAGTTCATCCCCCTCTTTCCCTTCCTCCTCCGAACGCCTCGTCCGCCACACGCGAACTGGTTGCCGCACTACCATTTTGGGTATGCTGGGTCTTTTTTGGAGCTCAGTGGGTGCGGCTTCCCTTGCAAAAGGTTTGCCTCTTTTATGTTGGCTAGCTAGCTAGTTGGGGTTTGGGCCTATTGGGCCCTTTTAGAGGGGGCACCGCCCTCATAAAAATTGCTTGCAATATTCAAATAACTTGGGCATGAAACGCGACGGGAGGAAAAGAGAAAATAGGGCTATAAAGTGTGGAATATTAAAGGAGGTTAGAGGAAGATATTAAATCGGAGGATGCGGATAGAAGATAGAGAAAGTCACGCTAGAAATATTTCCATTTCGGAGTAGAAGCATGAATGGGAATGCGTGAGTGTTGATATCTATGCATAGATATCTAAGAGAGAGAGAGAGAGAATTTAGAGTGGAGCTATCCACTTTGAGGCTCTAGCTATTTCTAACCCTTAATTAGTTCACATATGGGTGACTTTATCATCAACTGATGTGGCCCCATCACTCGCCGCAAACCATCTCTTGAGCCCCCTCCTCTTCAGGGCGAATTTCGTCGCCGCCCCGAAGCGTTCTCGTGGGCTCCCTACGTCTAGCCTTTGACCGATCCGTGGATGTCTTGCCATCCTCCCACGCGCAGAGTAGCTGCCTTTCGGGAGCGTTTTGGGCAGGTCTT*TGTGTTTTGTTTTTTCTTCGTTGTAAGGAGTGCATTTGTCTCTAGCCCTTTCCTAAGAGTTAGGGTTTGGGTTTTTAACATTTAAGATTAAGAATTCAATATTTATAATTTATGATTTTGAGATTAGGACGAGTTTATAATTTAGGAGTTCATGTCTTGTGAGATTTAACACTTTACTAGCTTAGACACGTTCTAAGACTAATGTGCCTTTTTGGTTGGTGATATGAATCTCATGTGGAGATGTTGGTGATGTTTTATGTTACCTTTCACAAGGCAAGATTTTTTTAAGGATGTTAAATAGAGGGAGGTTGGGACATGTAGCCATTAATGTACACCTATTATTTTGCATTTGTGGAGGCCATAGTTGTAGAATTCCAACTTTTTTAGGGTGCAATGTTTCAGATTTATGATTTAAGATTTAATGCTTAAAGTTCAGAGTTTAAGATTTAGTTTTAAAGTATGGCCTTTTTTAATTTCTACTTAAAGGGTGTAACTCAAGGCTTACCCCCTTTTTAAGAAGCCACCCATCTTAATGCCTCTTTGACCTGAGCATACTTAACTTAAAGGGTGTGATGTATGAAAATTGGTGCCTACTGAAAGGGACACCTAGTTGACCTTCTTAGGAGGTCACTCATCTTAATACAAATCCACTCAACTCCAGAGCTCTTATAAGATCTAATAGTAACATTCCCACTAAGAAGGTTTATGATTTAAGGTTTCAGTATTTTAAGGTTTAGGGCTTAGAGTTATTACATTTTAACATTATAGTTTAAATTTAGGATTAGAGTTTAGAGATAAAAAGTTTTAAGATTTATATACAAAATTTTAAGATTTTTAATTTAATCTTAAGAGTAAGAATTAAGGACTAGGATTTATGATTTATGATTTAAGGCTGCGTCTAGATATTTAGATTTTGGAATTTATATTATTTAAAATTTAGAGTTTATTATTCAAACTTCACTCCGACATCCACCTTTTGGTGAACCCATCTACACTTTTGGTTCTTGACCGCGCAGGCCGAAGAAATCCCACCCATTCGTCCACGCGGATGACAGGAAAGTAACGTCACTGCCCGTGGATAATGTTTTCTTCTTCTTCGTAGCCCTCGTAAAGCGCAGAAGCTGACTGCCAGCCGGTTGCTGTCGGCGTTTCTCTAACGCCGCTTTCTTCCATTTCCGAGAGTTGTGTTCGAAGATGTATGCGTAAAAACCTCGAGCCACCACTGTGCCGCCATGGGTTCTTCTCTCCCGCTGCAGTACTGCCTTTCTCGCTCAATCATCCTTGGTGGTAAGGTGAGCAATGGCTCTCTCTGTTTCTGCTTTCCTCTGTCTGTCCCCGCTTTTCAACCTCCTTTCTTTTCCCGTGCCTGTTGATGAGCATGTGCTCCGCTGGGTTCTCCCCCCTCTTCATTCGTCGATCAGTTGCAGGGGTTTCAAAGCGTGACCACATGGCCGCGTCTCAGTGACAGCAAAAACGGCGTCAATCCTCTGACGCTGGGAAACAAGCGGAGGACATCATCTGCTACTAGAGCTTCGATCAGCACGCCTCCGTATAGGCTATTCGAACCGCCGCATGCGGAGGAGTCATCCTCTGAGGTGCTTCCACCTCCCCTTATCTCCTCTTCATAAAGTTCCAACATTGGCGTTTCTTCTGTGTTCTCGTTAAATCTGGGGAGGAGTCTCTCTTATGCCATTCCTTTGCCCTTATCTTCTCTTGTTTATTTGCATAGCTGGAATCTGCAGACCCGGATTTCTACAAGATTGGGTACGTTAGGAGTGTCCGGGCTTATGGAGTCGAGTTCAAAGAAGGGCCGGATGGATTCGGAGTGTACGCCTCTCGGGATGTTGAACCCCTGCGCAGAGCGAGGGTGAGTTTAGCCTCGACTGATTTTGATTAGAGGATTGTACTTCTCACTAAACGATTTCTCCTTGAAGATCAATTTTCAGAAGCCAGCAAAGTCTACGTCTGCACGTAGCACTTGCTGTGATTTTTGAATTTTCATACACTTAGCTGTGTAAATCTATAAGATATCTTCTGTAGTCTCATTGCTTTCTTAGACTGGGTCTTTCATGATCTGCATAGTCCATTTTCTCTGGCATACTATTTTTTCCATGTATTAAGCTTTCATTGTGTTATTGTGATCATGCTTGGAGGCTGGCGTACATGAGACAAGATATCTTTTATAACCTAGGCCTATGTTATTATCTGGTCGGGGAAACGAGAATCTCACTAGGGACATACGAGCTTCTGAAGAAACCTATTTGGGACTCGTCTCCACTGGTGGACATTTAATGAAAATTTACGAAAAGTGTCACACTACTTGCTTTTCATGGAGCTTCATGAATCCCAACACTAGGGAATGAGTAAATGATGAATTAATCTGGTAATTTGACGGTTTGTAGTGTAAATTTTGGCCGAGCTTAACAAGATACTGCTGTGACCGTCAAATGGGAGTACTGAATGCACTGGTACTATCTATTACATTTAATTTTCAACGTTGTAGAGCATGTGCTATCATCTGTGCCTCAGGATGCTTGGGTGGTCTTAAACTGTGGACGATTTCGAAAATGGGGCCAATGTGGAGTAATGACCTGTTTTTGATTCGAGACTGGGACCAATCTGCTACGGATGCAAGTTTTTATTCTCAGAAGAAACGAGAACTAACATGTCAGTGCTCATCAGCAGAGAAGCATGCTCGAATGAGATTTCATTAATTTCTAATCTTATTTCCAAGCCTATGTCCCCGGACTCTTTTTGTTGAACCCCAGACTATCTCAGTCTGAGCTTCCGGTTTTTGTTAAGTGGTTTCAGTAACCAGACTGGATAAGTGACCTAAAATATTTCCTTGTCAAGTCTTGTAGTATTATCATGCCATTGCTTTCTATTTCGAAGGATTAGATCAGTCATCAGGTAGCAACAAGATGCCACCGTTAGATCTTAGTTTCCATATTTTTATGAATTCTAATATTCAACTCTGTGTTGAGGGTTTGAGTTGGTATGTTCTTAAAATGCAGTGATGGCATGATTAAGTTCCTTTGATTAATGCTTCCTTTTTAATTTTCATGATAATTTTATTATCTTTCCTGATTGATTACGAAAGACATTTACTGGTATTCCAGCTTTTTTAACTGGACTTCATAGTATTCGTTAATTAGACTAAGAATATTATTGAGTTATCAAAAAATTTCTAGAATTTCTTTCGTTCTGGCTTCTACAAGTTCTGAAATTTCATGGAGGATGATTCTGGGTAAATACTTACCAGAAAAAAGGTCAAATATTAATGATTTGAACTATGGAAATGACCAAATCTTCAATTGATAGTTTTACTCTCAGAACAAGCAAAATTTTATGCCATGTGCTCCTTTTGTGTTCACATTCACTTTTTTGTTTTTTTCTGTTTCCTTTTTTTATGCTGTCTCTACTTTCTTCATCCTTCTATTTTATTGGAATAGTTTTCCCTTCACTTTTTGCCATGCTTTCCTTATTTGGTTGGCTATTGGTGAAATGTTTTTTTATCAGATAATAATGGAAATTCCACTGGAGTTGATGCTAACTATCAGCAAGAAGGTTCCTTGGATGTTTCTTCCAGATATTGTGCCAGTTGGTCATCCTATATTTGATATCATCAATTCAACGAATCCCGAGGTAAGCCAATGCTCTATCTGATGATAATATCTTCTTCTTAAGTTAGGTTATGCTACTTTCGTTTTTTCTTAACTTTCCCTAGTCTATTGAAAATTTTCCAGTTTGTAGTAGATATGATAGAATTAAATCTGCGAGGAGAAAAATCATTTCATTTTCATCCCAGAGTCACTCATATTTTTATTTACATTTCTTTTTGCTCAGACCATTCAATGGTTGAAACTTTCGAGATTTTGTTGAGATAGAATACAATTTGTTTGTATTTTTAACCTGTCGGTTTTTATTTATGGACACCATTAATGTTGTCAGTTGCTCTTGTTCATCTGTTTTGAATGAATCATGCTGTCTTCAGATACTATGTTGCAATTGATTTTCTGACTTTGGGGGTATTATTTTTCTGATATTATAGTTACAATGGCCTGTGTTTTTGCTTTTTCGTAGGGTATCGAACAATTAAAGCTAACTCCTTGTAGACGATTGATAAGTTATCATTTTGATGGAATTTAGTCACATGGATGTCCTTGTGATTTTATGGACATTTGAAAACTAGTCATTTGGATATAATTTTATCACATGGATGCTCTTGTGATTTTCCCGCCTACAAATCTGTGTGGAAAATATGCTCCACTGTTTTAGGGTGCATCAGACAATTTTTGTTAGAAAACTTTATTAGAACCTTGTAGAAGCTGTTTCCTTATTAATTTCAAGAAAACTGATTGACATTAGAGCGGACCAAATTTTATTTCAACGATAAATATTTACTGTACGATAATTGTGTATTGTTCTGAAGAATGATAGGAAAAGGTTGAAGGATGTTCTGTTTCATAAATGACTGATTTTCTTCTTCAGCTTGGTGGTGAAATGTATGCACATGGTTTTTTAGTCTGTGACAATCATTCTGAATCATTATGCTCATTTGAAACCATTTTTTAATGGATTGGGTTAAATAAAAAAAATTGCATTTTCGAACTGAAACTTTGGAGGAACTCAAAATGCTTCTAGAGATTTCAGGCTCTTTTTGAGATTGAGAACAGACTCAATGCTGTTGAGTGCTGTGTCACAGGTGGCGAAAGTCCCAAAACCCTGATTCCAAACAGGAAAGTGGAAAGTAAGAGTGACTTGGATTGATTAACATTTTCGGTCTGGTATTTATACCAGGCCCATCAGGGTTGAACCGGACAGGTCTGGTCCAATATGACAATTAAGTGATAAACAACTTATTCTGACAAATATTTCAATATGAAGTGATGGTACTATGCAAGTGTTAGGTTGGAAAAAGTTGAGGGTAGTTGTGGGGACCAAAGGGATGGACCACTGCCTAGGCAATAAGACTACACCGTATCCTTTCAAGTAGTAAAAATTTTATCAAATGATTTGAAGAAACCGAAAAAAAAAAAACTGTGTTCTAATTAAAAAATAAGGACTAAATGAATAATAACTATGAAAATATTAAAGAATAAATTAATAAAGCATTGGCATTTGGATGTATGCCTTAGCGAAGGGAAACAGTGTCAATGCTGATGTATTTGGCTGTGGTATCTTGCAGGTTTCGTTAATGCATCTTTACTCACATGGAGCATGTCTTCGAAACTACAAAAGAACGAACAGAAGAGTTGAGCATTGACAAACCAATTAAGGAATTTTATCCGACTGTAGAAAAACTGCTGGAGTAAGAGTGAAATAATGGACAGAAAAGTTGATCAAAATGTTTTGATCTTTTCAGTGAAAATGGTCAATGTACTTCTAGCTTAGAACTTCAATATTGTCAACACCTGAGGAAATGTTCCCTTAGATGATAATCTAACACTTGTCGGATAACTGAAATTTCTTCTAGGTGTTTGAATTGGGCCTTATTTTGAACAAACTTTAAATAAGCTCCATTAATATGCTGAAAAGTGCCATATATTTAAAACATTGGATTTTGAAGACTGGGTCTCGCTGTTACAGTGCTCAGGCTGTGTTTTATTCAATTTTCTGAGAACTTAATGAAGCAGTATTTTATGATAATTTAGCTTTCTTCTGCTGAGTATTCATTCCAAACTGTTTTTTGAGAAGGGTATTTCACAAAAACTTAGGTCATCTCTGCTACTGGAAACACAGGACACTGTTGCTTTGCGTCAAATTAGATTTTTTTCATGAATAATCGTGTAACTAGTAACATAATTTTGAGAAACATGTAGCAACCTACTGTGTTTTAGTCATTTTGTTAGAATTTAAGTGAATACCTTGTATCACAAATAAAAGGTGGAAATTCATTGCAAGATTTCTGCAGTGACAATAATCATTATCCATAAGTCTGGAAAATTTGTTAAATTGTTTGCAAGAATGAAATAATATACCAAAACTAAAATATTATAAAGTACAAGTCATTTTAAAATTTATTTAGAAAACTATGAAAAAAGAATTTGATTTCAAAGTCATGACACTTTGAGGGCAGTCATCAAATTAATACCTATGAAATTCGTAGTATGCATTGATGTGAGTAAGTTTGCTTGTGTAAATAATATATTTCCTATCATGAATGTGAATAGGTTATTGCTTGTATTTGTTGGAACTCCACGAATCCTATAACACTCCTTGTTAGATGGTATAATTGTATTATTGTAAATGACTAAGGAGATTCTAGGAAAGTGTATGTAGGGTTCATTCTTATTTAATGAATGCTAAAAGTGAATGAACGTTGAGACGGTTTTTCTGCCATTCCTCCCCTCTATCCTCCATGACAAAGGTTTTCTCTCTTCTCATGTGATTGTCTCCTTCCTCTCTTCTACCTTAGTTTACAACATGGTATTAGAGCAAGTAAGGTTAGCTCATTCCATTTTGTTCGAATTCCTTAAGTTATATTTCGTTGATGATTCTCTCAATCAAAGGCACTTGATTTTAGGTTTCCCTAAGTTTAAGATTTATCTTGTCGACATTATTTTGTTGATTGTTGGCTGTCTCGTTTTTATAGAAGGATATTCTCAACTCTATTTATCTGACTTTCAATCTTGGGGGTCTATCTGATTTTTTTCTTTCCGCTACATGACCCTTCATTCCTTTATTTGGCCTCATTTCTCAAAGAGAGGATTGAGAGCTCATCAATTTCTGGCTGAGAATAGGGTGTTTCTTTCTTCTACATGACCCCTCATTCCTCTATTTGACCTCATTTCTCCTAGAGAGGATTTGGAGCAAAGTATTTGGTGGTTGAAGATAGGGTGTCTCTTTCTACTACATAACCCCTCATTCCTTTCTTTGGTCTCATTTTTCCTAGAGAGGACTAGGAGCACAACATTCACTAATTGAGGACGTGTTTTCTCACTAGCTGACTTAGAGTTTCTAGTATCATCACCATCTTCATCGTTGTCTTTCATCAATTTGGAGTTGTTTACCGTCTAGTGTTTTTGTCCATCACATTTAGTGAACCGCCCTCCTCAATACGTCACCCTCTGCCATGCATTCATATGACATCCCCCTTAGTCCTGGGACTAGAGTTTCCAGTCTACACTGCTGCAGTCTATGCAATTTTGTGCCATCCACCCACCATTTGGAGCCATCCCTCTTCTGCCTTTCATTTAGTGCTCTCTCAATCGTGCCAAGATTATCAAGTTCCATTTAATGTCATTCTGCATCATTTTATGTAGTTTGGTACTGTTTCATGAAGTCAGGTATTAAACCATGTTATCATTTGATGAAATAGCTCCATCAAAACTCACCAAAAGAGCTCTTCAACATCGTCATAAAGGGAGTTTCAATAAAGGAGCTGCCACAACGGATCATTAATGGCATGATTGTCAAAGAATTAATCAAGATAAAACTTTTTTAAATTCATCTTGAGGGGGTGTGTTAGATGGTGTAATGGTAATGTTGTAAAGACATGGTATTTCGGCAAAGTGTATGTAGGGTTCATCTTTATTCAAGAGATGTTAAAGGTGAATGAATGTTGAGACAGTTTTTCCGTTGTTCTTCCCCTCTGTCCTCTGGGACAAAGGTTTTCTTTCTCTTCTCATGTGATTGTCTCTTTCCTCCCTTTTACCTTAATTTACAACACTCAGTATGTTATTTCTTCTGTTGTACTGCATGCTTTGTGCTTTTTCTTCACTAAATGTTTCTTCCTTACTAAAGTAATAGAGGTAGGGCACTGGAGCACCTTTAATCATGTTAATATCCAGTATCTTTTCTTCTAGAAATTCTGTTGTACGGTTTACTTTGCCAATGGTAAAATGTAAACTTCAATAAATTAAAGTCAAAATAATGGTGTTCCATTTCCCTTTCGTTAATCTTTGATCTTTTAGCCAAAGAATAATTTGGTTACTTGTAGATTTGGAGTGGTTATTATTATTTAACAAATTGGATGTGTGGATGAGAAAGTTTTTTTTTTTGCATTCTGACACTTTGTTGCCTTTATACAGTCTGACTGGGATCTAAGGTTGGCCTGCCTTCTTCTATTTGCACTTGATTTGGAGGGTAACTTCTGGCAGCTGTATGGAGACTTTTTGCCTAGTCTGGATGAGTGCACCAGCTTGCTTCTTGCTTCAAAGGTTTGATTTATTCTTAGATAAAACGTAATTCCCCTAACTGATAGTAATTTAGATTAGAAAAATGTAGCATTGATGCTGGAATTTTATGTGAAATTTCATTGGTGAATTTACTGAGTCAAAAAATAAAATTTTATGTGATGTATTTTTGCATATCTTGTACTTGTAGATACCCCACCATTTGGAAATTTGTTTTATTTTTCATTTTAATTATTTTTATTTTTGTGCAATATGCATAGGTGGAATCAATACCCTGTTGTAAAGCACGCTTTCAAAAGCTATGGTGTGGTTAGTGTTGGTGCATTATTCAAATTATTTGTAGAGGCATTCTTCAATAGTTTGTGTTCGTGTTCACAGCTAGTATAGAGATTATTGATGGAATTTGATGATCTTATGTAGCACTGTAGGTATTCAATTAGCTTTTAGTCTACCTATTGCAAGATGGCTTTTCAAGGCGTCAACCTAATGTAGGATATGACTTTCAGAAAAAAGGATGGCGAAAGCCTTATGACCTAACTGAAGGACAGGCTATAAAAACTGAGAAAACTCAAAAAAGGCTCAAAATTAGATTATACCCGGATGAACTACTTGAATAATGAGTCAGAGCCAGTTTTTAGTGACTGGCATGTTTGTTAGATTCTTTGAATTTGTTACTCTCAGATAAATTGTGCAGTAGGGTTAGTTTTAGTAACTAATCCTTGTGGATTCATTGGACGTTACTTTGATCTTTGTGGAACTATGGATCATCCAAATCAACTCTTTAAATACAAGGACATATTTTTTATCCTTGTGGCTTCAAGGGTCGTCCAGGAAATCCTTGTAGATTCAAGGATTGGGAAAAATCACGCCAATTGGATTTAAGGTACCATTTTTTTCTTCTTGTACTCCAGATTTGTGTCTTGAGTATTCAGGACTTAAATCATCCATTCTCTATATCTCTTGCTATGTCGATTAGTATGAAAACTAGGTTTCAGGGTCTTCTGGCATCAAGAAGGATTTTATGATCCCTGTGGATTCACGTATCATTCGGACCAATCCTTGTGGATTTGGCCTAATTTCTGTTGTTCAACAAGTCCTATTCATGCCTTGGTATCTACTACTTTAGGGATTCCACCTTTTTGATATTCATTGAGGTCTTGAGTTTTGAGGTAAACCAAAAAATTTTGAGTACACTCGTATCTAATCTTTTACATCTGAAACTTGGTGCTCATTAGCATAATTGCATGTCAGTGGAACGAAATTAGAGCGTGCTGTGCTTGAAGGAGATGAATGTTTCTTCTGTGTATATTCTGTTGCCTTTTTGTTAAACTTCAACTCTTCAACCTTTTGGTTCCACATCACTTTATTTTAAAGTATCACTATAATATACTTCCATAGCCTTTTATAACTCTGAGTCTTTTAACTATTGTCATGACACATCACAGCAGCCTGGGTTTCCTTGATTTTTGCTTTCTATACTATCGTGATTTCTCACAAGAGGGGGGTAAATCTCATCCATAAAATGGAAATAGGATGCTACAGGGTAGAGGAGACAGAACCTCTCACGCTCGAAGTTAGAGGGGAGAGCGGTGGGCTGCCGACTCACTACTTCTCTGAGCCCTACTGTCTCTTAAATGACTAATCCTTACAGCAAGATTCCTAAAAGACCATTATATATTATATATTCCCAAAATACCCTCACTCCTTACGACAATTACTTATTTGAATCTCTTTGTACTACGCTCTTTACTTATTAGCAGTCTTTGTTTTAGCAGTTCATGTTGTCATTGATGCCTATTCATTGTTGACACAAACTTGTTATTGTTGACATGTTTGGTATTGCGGTTGTGTATTTTATGTAT

Supplementary Figure 10. TATA-**binding proteins SpTBP1 and SpTBP2 of *Spirodela polyrhiza*.**

**a** CDS of *SpTBP1* and *SpTBP2*. **b** Deduced amino acid sequences of SpTBP1 (PQ619461) and SpTBP2 (PQ621416). **c** Alignment of amino acid sequences of the TBPs of *Spirodela polyrhiza* (SpTBP1- PQ619461, SpTBP2 - PQ621416), sacred lotus, *Nelumbo nucifera* (NnTBP1 - XP_010276406.1, NnTBP1 -XP_010257701.1), *Arabidopsis thaliana* (AtTBP1 - NP_187953, AtTBP2 - NP_187953), and maize*, Zea mayz* (ZmTBP1 NP_001105318.1, ZmTBP2 - NP_001105319.1).

**a**

***SpTBP1***

ATGGCCGACCAGGTAGAGGGGAGCAGCGAGCCAGTCGATCTTTCCAAGCATCCGTCCGGCATTGTTCCCACGCTACAGAACATTGTTTCAACTGTTAACTTGGACTGCAAGCTAGATCTTAAAGCCATAGCTTTGCAAGCTCGCAATGCCGAGTACAATCCCAAGCGTTTTGCTGCGGTAATTATGCGCATCAGGGAGCCGAAAACAACAGCTCTGATATTCGCTTCTGGAAAAATGGTTTGCACTGGTGCGAAGAGCGAGCAGCAGTCGAAGCTCGCTGCGCGGAAGTATGCTCGCATCATTCAGAAGCTGGGTTTTCCTGCAAAATTCAAGGACTTCAAGATTCAGAATATCGTGGGGTCATGTGACGTCAAGTTCCCCATCAGACTCGAGGGTTTGGCCTATGCACACGGCGCCTTCTCAAGCTACGAACCGGAGTTGTTTCCTGGCCTCATCTACCGGATGAAGCAGCCCAAAATTGTGCTCCTCATCTTCGTCTCCGGCAAGATCGTCCTCACCGGCGCCAAGGTGCGGGAAGAGACGTACACAGCCTTTGAAAACATATTCCCCGCCCTCTCCGAGTTCAGAAAAGTCCAGCAGTGA

***SpTBP2***

ATGGCAGACCAAGTTATGGAAGGAAGCAGTCAGCCTGTTGATTTTTCGAAGCATCCGTCAGGGATTATTCCCACACTTCAGAATATTGTTTCTACCGTTAACTTAGACTGCAAGTTGGATCTTAAAGCCATAGCATTACAAGCACGTAATGCAGAGTACAACCCCAAGCGTTTTGCTGCAGTTATCATGAGAATCAGGGAACCCAAAACGACAGCTTTAATTTTTGCATCTGGAAAGATGGTTTGTACAGGTGCAAAAAGTGAACAACAGTCAAAACTTGCAGCACGAAAGTATGCTCGGATCATTCAGAAGCTTGGTTTCCCCGCCAAATTTAAGGACTTCAAGGTCCAGAATATTGTTGGCTCTTGTGATGTCAAATTTCCTATTAGGCTGGAGGGTCTGGCTTATTCTCATGGTGCTTTCTCAAGTTATGAACCAGAGTTGTTCCCTGGTCTGATCTATCGAATGAAACAACCAAAGATTGTGCTGCTGATCTTCGTTTCGGGGAAGATTGTGCTCACTGGAGCGAAGGTGAGAGAGGAAACCTATACGGCATTCGAGAACATATACCCAGTGTTGACAGAGTTCAGGAAAAGCCAGCAATGGTATGCGTCTTGTTTCTGCATGGAATTTTTCTCGAGCTCTTTTACACGTGGATCTACGTTCCCACTAACGCCATATAAAAAGCACAGCAGAAATAAAGTATCTTTGCTTTGA

**b**

**SpTBP1**

MADQVEGSSEPVDLSKHPSGIVPTLQNIVSTVNLDCKLDLKAIALQARNAEYNPKRFAAVIMRIREPKTTALIFASGKMVCTGAKSEQQSKLAARKYARIIQKLGFPAKFKDFKIQNIVGSCDVKFPIRLEGLAYAHGAFSSYEPELFPGLIYRMKQPKIVLLIFVSGKIVLTGAKVREETYTAFENIFPALSEFRKVQQ*

**SpTBP2**

MADQVMEGSSQPVDFSKHPSGIIPTLQNIVSTVNLDCKLDLKAIALQARNAEYNPKRFAAVIMRIREPKTTALIFASGKMVCTGAKSEQQSKLAARKYARIIQKLGFPAKFKDFKVQNIVGSCDVKFPIRLEGLAYSHGAFSSYEPELFPGLIYRMKQPKIVLLIFVSGKIVLTGAKVREETYTAFENIYPVLTEFRKSQQWYASCFCMEFFSSSFTRGSTFPLTPYKKHSRNKVSLL*

**c**

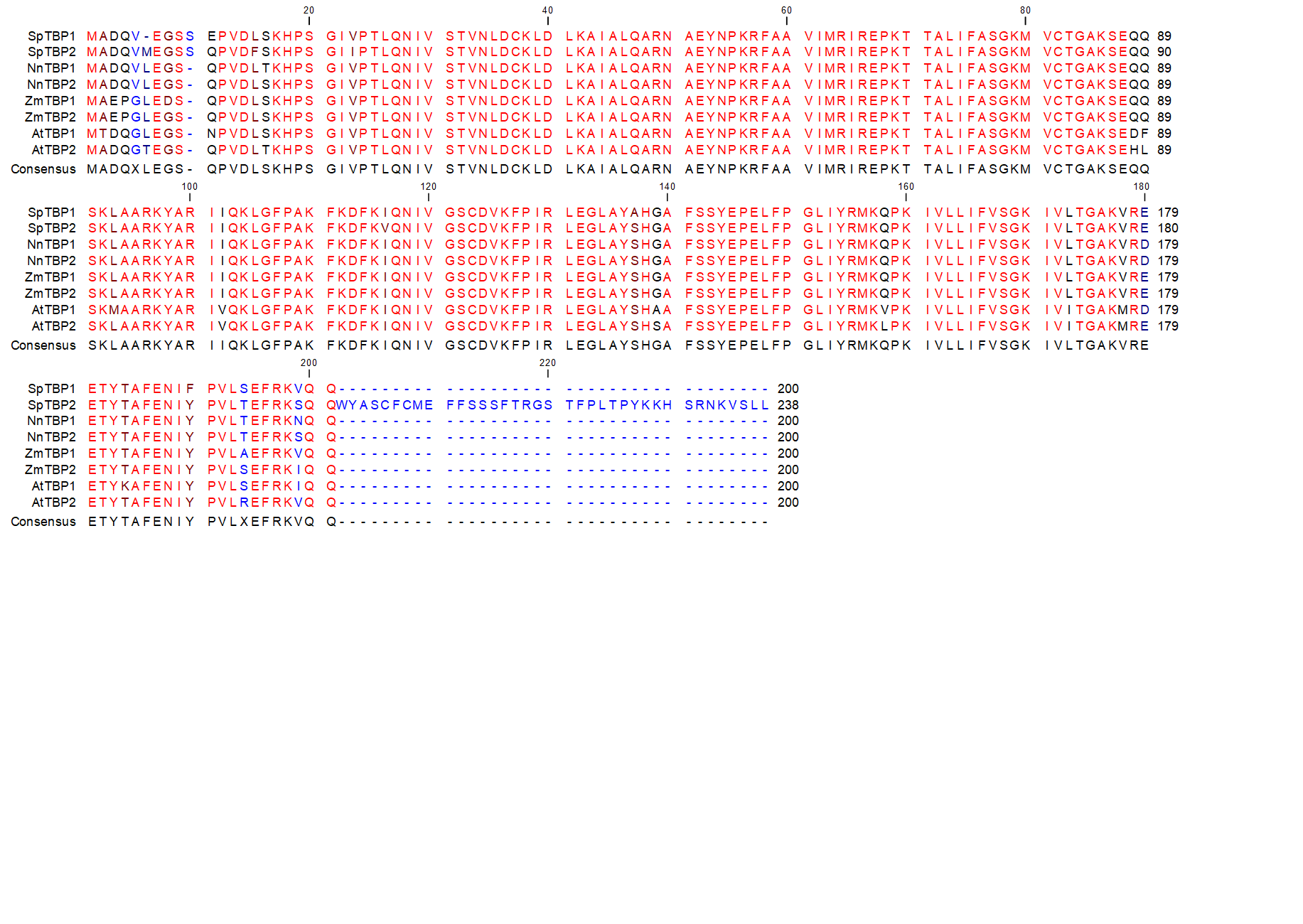

**Supplementary Table 1.** **Signals volume and intensity measurements after FISH using an Alexa594-labeled probe specific for Sp5S-S on fifteen different nuclei.**

| **Samples Measurements** | **Smaller signal 1** | | **Larger signal 2** | |
| --- | --- | --- | --- | --- |
|  | **Volume (µm³)** | **Intensity** | **Volume (µm³)** | **Intensity** |
| 1 | 0.0525 | 1.03E+07 | 0.1060 | 2.11E+07 |
| 2 | 0.0396 | 1.06E+07 | 0.0894 | 2.62E+07 |
| 3 | 0.0465 | 1.77E+07 | 0.1140 | 4.16E+07 |
| 4 | 0.0658 | 2.20E+07 | 0.1750 | 4.42E+07 |
| 5 | 0.0684 | 2.14E+07 | 0.1010 | 3.62E+07 |
| 6 | 0.0653 | 1.97E+07 | 0.0893 | 3.10E+07 |
| 7 | 0.0312 | 1.10E+07 | 0.0531 | 1.79E+07 |
| 8 | 0.1290 | 2.71E+07 | 0.1360 | 3.81E+07 |
| 9 | 0.0477 | 1.84E+07 | 0.0743 | 3.12E+07 |
| 10 | 0.0410 | 1.07E+07 | 0.0964 | 2.65E+07 |
| 11 | 0.0363 | 1.18E+07 | 0.1120 | 3.96E+07 |
| Sum | 0.6233 | 1.81E+08 | 1.1465 | 3.54E+08 |
| **Average** | 0.06 | 1.64E+07 | 0.10 | 3.21E+07 |

**Supplementary Table 2.** **Estimation of FISH data obtained with Atto488-labelled probe specific for Sp5S-L on fifteen independent nuclei.**

| **Samples Measurements** | **Smaller signal 1** | | **Larger signal 2** | |
| --- | --- | --- | --- | --- |
|  | **Volume (µm³)** | **Intensity sum** | **Volume (µm³)** | **Intensity sum** |
| 1 | 0.1500 | 2.42E+07 | 0.2380 | 3.79E+07 |
| 2 | 0.0463 | 7.45E+06 | 0.1250 | 1.60E+07 |
| 3 | 0.0226 | 2.73E+06 | 0.1090 | 2.56E+07 |
| 4 | 0.0494 | 1.10E+07 | 0.1380 | 3.33E+07 |
| 5 | 0.0346 | 4.07E+06 | 0.1130 | 1.66E+07 |
| 6 | 0.1210 | 1.03E+07 | 0.2030 | 2.47E+07 |
| 7 | 0.0455 | 8.70E+06 | 0.1300 | 2.99E+07 |
| 8 | 0.0284 | 4.28E+06 | 0.1890 | 4.00E+07 |
| 9 | 0.0371 | 1.01E+07 | 0.0942 | 2.52E+07 |
| 10 | 0.0400 | 7.03E+06 | 0.1640 | 3.64E+07 |
| 11 | 0.0343 | 6.60E+06 | 0.1970 | 4.63E+07 |
| 12 | 0.0263 | 4.63E+06 | 0.1550 | 3.16E+07 |
| 13 | 0.0526 | 1.36E+07 | 0.1660 | 3.62E+07 |
| 14 | 0.0346 | 3.73E+06 | 0.1930 | 3.03E+07 |
| 15 | 0.0312 | 3.69E+06 | 0.0315 | 1.76E+07 |
| Sum | 0.7539 | 1.22E+08 | 2.2457 | 4.48E+08 |
| **Average** | 0.05 | 8.14E+06 | 0.15 | 2.98E+07 |

**Supplementary Table 3. Analyzed extra-long reads containing shorter 5S rDNA units of Sp5S-S type.**

|  | **Length, bp** | **Upstream sequence, bp** | **rDNA copies** | **Downstream sequence, bp** | **ON read ID** |
| --- | --- | --- | --- | --- | --- |
| 1 | 57,788 | 1,690 | 40 | 33,897 | 3ab0b3f8-9e42-499a-a733-e7f7ef5a88e3 |
| 2 | 40,102 | 19.247 | 36 | **-** | 9e7ce0b7-2939-4dbb-bae9-bd122f577594 |
| 3 | 29,933 | 17,310 | 25 | **-** | 185db047-daf5-4418-bfc1-3146a719f062 |
| 4 | 25,619 | 2,965 | 45 | **-** | 4086360e-5052-4bcb-a8cf-bef2420e8378 |
| 5 | 32,380 | **-** | 65 | **-** | fe9ebfe0-b4ab-4f96-bb7f-b8aefc7a02d8 |
| 6 | 16,061 | **-** | 32 | **-** | 4c73f83e-0b3e-4ffc-a967-fec8a2b45b5f |
| 7 | 16,027 | **-** | 32 | **-** | 0941502f-db94-4747-a66c-eac550034760 |
| 8 | 21,176 | **-** | 42 | **-** | bd6dd058-e66d-4a56-bdb7-ceb6cf75d7c8 |
| 9 | 22,147 | **-** | 44 | **-** | 4b63a220-c21e-4e7d-a3c0-158ddc218cfd |
| 10 | 20,371 | **-** | 40 | **-** | 826afbef-7b6b-4a9b-ae18-2d68a34ce955 |
| 11 | 19,244 | **-** | 38 | **-** | 04312a98-50fa-4e3c-b8f3-c3441df8df10 |
| 12 | 21,765 | **-** | 41 | 877 | 3aaec8ea-153c-4961-bb09-709bfcc853d0 |
| 13 | 22,552 | - | 30 | 7,650 | 64274095-ccaa-41ef-938a-92bfa4de3b5c |
| 14 | 54,000 | - | 39 | 34,923 | tig00000154 |
| 15 | 38,563 | - | 10 | 34,269 | e01ffd98-937c-4aaf-b945-ccb8d9b38713 |
| 16 | 16,912 |  | 31 | 1,500 | 354677d5-b0d5-4ab9-8bc4-a1473377246e |
| 17 | 35296 |  | 18 | 26,202 | 82a2bb86-1f26-4371-bcf5-24a910bada4c |
| 18 | 24,538 |  | 24 | 12,466 | 8c50c172-9709-469e-88f6-22205bc2644a |
| 19 | 23,868 | **-** | 26 | 10,433 | 46c6ca65-0ff9-4d66-9851-44c98cf7823b |
| 20 | 22,252 | **-** | 6 | 19,404 | 53de415d-56a3-4fbd-a07c-535553365f4e |
| 21 | 21,427 | **-** | 17 | 12,288 | bbdd3995-2fc7-471e-a4e5-3315fa7556d4 |
| 22 | 25,746 | **-** | 9 | 20,856 | b543dff1-3c45-4343-a961-d9cc74f8e565 |

**Supplementary Table 4. Extra-long ONT reads containing Sp5S-L type 5S rDNA units.**

|  | **Length, bp** | **Upstream sequence, bp** | **Minor cluster, rDNA copies,** | **Major cluster, rDNA copies** | **Downstream sequence, bp** | **ON read ID** |
| --- | --- | --- | --- | --- | --- | --- |
| 1 | 85,978 | 44,910 | 2 | 38 | - | 0a173a95-c7c9-4fe4-abba-d5044ff12423 |
| 2 | 50,300 | 25,724 | 2 | 23 | - | b289bf08-e82b-4bcf-9f3c-5671fe2dd9dd |
| 3 | 75,841 | 13,408 | 2 | 57 | - | 3d770410-1945-4622-afd8-7a3257e79119 |
| 4 | **73,409** | 15,456 | 2 | 51 | - | dfdf443c-0e79-4afe-b46b-540bd8a6b145 |
| 5 | 46,592 | 40,647 | 2 | 5 | - | 102846b4-1a31-48da-94df-ee234e45947c |
| 6 | 28,925 | 17,429 | 1 | 11 | - | f879b4a1-d6ad-4e5b-99b5-d9d8d51357a0 |
| 7 | 53,598 | 39,088 | 1 | 12 | - | e048 9576-770f-4f88-9aef-2667b331eef5 |
| 8 | 26,992 | 16,113 | 1 | 10 | - | 17bfb 185-203a-49b9-b54b-07ca1354cf60 |
| 9 | 69,203 | 53,047 | 1 | 15 | - | 70c01d50-5ccf-4b6f-b163-e6aae90aab56 |
| 10 | 52,251 | 29,060 | 1 | 20 | - | c72d9c80-f704-4d3d-b030-25b9f213fe3c |
| 11 | 48,555 | 8,416 | - | 35 |  | d5727ca2-07e3-4691-bdb2-217b0be2e207 |
| 12 | 63,701 | - | - | 56 | - | d3ee0938-b6c9-4e89-a57b-b0eb83cac377 |
| 13 | 68,265 | - | - | 60 | - | e9d7a5e2-5a84-4615-a68c-f902b4cce5d4 |
| 14 | 39,765 | - | - | 35 | - | f2730a53-09b1-41ca-a286-8e0b18c96061 |
| 15 | 32,245 |  |  | 28 |  | 9b99f11a-694e-4f96-a7b2-14f8ba1cc468 |
| 16 | 32,452 | - | - | 28 | - | a3c916e5-1eb5-47e0-be72-f67549367915 |
| 17 | 44,618 | - | - | 39 | - | fa414864-fe20-4119-834b-ba5f1ffd6b16 |
| 18 | 51,239 | - | - | 32 | 14.650 | e8f2492e-3138-4f7f-bf33-7c0c23f3e617 |
| 19 | 58,280 | - | - | 29 | 25.072 | 975741bb-49d8-49ec-9870-ba5698e90901 |
| 20 | 17,431 | - | - | 12 | 5,198 | 098e8e1d-8ef5-4e08-84d5-d8dd3cc555c5 |
| 21 | 65,647 | - | - | 46 | 11,153 | 5cf44779-0cda-4a54-a602-c253f27d29a6 |
| 23 | 51,690 | - | - | 22 | 28,634 | 9cbd7e91-966d-4889-8c77-419a380015e9 |
| 24 | 19,405 | - | - | 9 | 9,415 | 11f73350-8620-4e29-8e50-640083d27c8d |
| 25 | 26,044 | - | - | 10 | 14,785 | ee291f06-0920-4936-be67-d37632f988a7 |
| 26 | 32,227 | - | - | 13 | 17,388 | e78851b4-75a2-4974-926b-8fbcd52a105f |
| 27 | 19,540 | - | - | 3 | 16,402 | 6cde006c-07ad-4b42-91e4-c3cb1c80bf26 |
| 28 | 33,836 | - | - | 27 | 5,910 | f7270481-6b72-4e0b-a23b-b3f24c080e9d |
| 29 | 34,874 | - | - | 16 | 17,167 | 2c785a69-5173-46aa-92d6-fdb85bebd299 |

**Supplementary Table 5.** **List of primers.**

| **Primer ID** | **Sequence, 5’→3’** | **Direction** | **Target** | **Used for** |
| --- | --- | --- | --- | --- |
| 5S-F1 | CTTGGGCGAGAGTAGTACTAGG | Forward | 5S rDNA | Cloning 5S rDNA units |
| 5S-F1 | CACGCTTAACTTCGGAGTTCTG | Reverse |  | Cloning 5S rDNA units |
| cn-5S-for | GGGTGCGATCATACCAGCAC | Forward | 5S rDNA | Copy number estimation |
| cn-5S-rev | GGGTGCAACACGAGGACTTC | Reverse |  | Copy number estimation |
| ActinF1 | TGTTTTCCCAAGTATCGTC | Forward | Actin | Copy number estimation |
| ActinR1 | TCCCAGTTGGTGACGATT | Reverse |  | Copy number estimation |
| 5SshcnF1 | CCCCTGCTCTGGTCTCTTGT | Forward | Type-S IGS | FISH probe preparation |
| 5SshcnR1 | CCCTCGAAAGATAGTTAGCC | Reverse |  | FISH probe preparation |
| 5SlgcnF2 | GACGGGAGGAAAAGAGAAAA | Forward | Type-L IGS | FISH probe preparation |
| 5SlcnR2 | TCCCATTCATGCTTCTACT | Reverse |  | FISH probe preparation |
| TBP1chr4F | GGAATCTGAGAAAAACGAGTGG | Forward | TBP1 | Cloning TBP1 |
| TBP1chr4R | ATTTCACTGCTGGACTTTTC | Reverse |  | Cloning TBP1 |
| TBP2F | CCATGGCAGACCAAGTTATGGAAGG | Forward | TBP2 | Cloning TBP2 |
| TBP2R | CTCTAGATCATTTGGCAATCCCTATTATGAGC | Reverse |  | Cloning TBP2 |
| 045I-5S-UpF1 | TCTTCTGCCTTGTACCTATT | Forward | ChrSp6 locus | Cloning upstream locus border |
| 045I-5S-downR1 | ACCCACATGTTAATAACATCAC | Reverse |  | Cloning  downstream locus border |
| 019I-5S-MinF3 | AAGGGAAGCGCATCCACATAG | Forward | ChrSp13 locus | Cloning minor  upstream locus border |
| 019I-5S-MinR3 | TATATCTCCTCACCCAAGGTG | Reverse |  | Cloning minor  downstream locus border |
| 019I-5S-MaUPF2 | CTAAATACCCACTTTATTCC | Forward | ChrSp13 locus | Cloning major  upstream locus border |
| 019I-5S-MaDownR2 | TAGGAAAGGGCTAGAGACAA | Reverse |  | Cloning major  downstream locus border |
